# Novel insights on obligate symbiont lifestyle and adaptation to chemosynthetic environment as revealed by the giant tubeworm genome

**DOI:** 10.1101/2021.09.04.458960

**Authors:** André Luiz de Oliveira, Jessica Mitchell, Peter Girguis, Monika Bright

## Abstract

The mutualism between the giant tubeworm *Riftia pachyptila* and its endosymbiont *Candidatus* Endoriftia persephone has been extensively researched over the past 40 years. However, the lack of the host whole genome information has impeded the full comprehension of the genotype/phenotype interface in *Riftia*. Here we described the high-quality draft genome of *Riftia*, its complete mitogenome, and tissue-specific transcriptomic data. The *Riftia* genome presents signs of reductive evolution, with gene family contractions exceeding expansions. Expanded gene families are related to sulphur metabolism, detoxification, anti-oxidative stress, oxygen transport, immune system, and lysosomal digestion, reflecting evolutionary adaptations to the vent environment and endosymbiosis. Despite the derived body plan, the developmental gene repertoire in the gutless tubeworm is extremely conserved with the presence of a near intact and complete Hox cluster. Gene expression analyses establishes that the trophosome is a multi-functional organ marked by intracellular digestion of endosymbionts, storage of excretory products and haematopoietic functions. Overall, the plume and gonad tissues both in contact to the environment harbour highly expressed genes involved with cell cycle, programmed cell death, and immunity indicating a high cell turnover and defence mechanisms against pathogens. We posit that the innate immune system plays a more prominent role into the establishment of the symbiosis during the infection in the larval stage, rather than maintaining the symbiostasis in the trophosome. This genome bridges four decades of physiological research in *Riftia*, whilst simultaneously provides new insights into the development, whole organism functions and evolution in the giant tubeworm.

## Main

The discovery of the giant tubeworm *Riftia pachyptila* Jones, 1981 at deep-sea hydrothermal vents on the Galapagos Spreading centre in 1977 (Corliss et al. 1979) has initiated the onset of a continuous torrent of studies (Childress and Fisher 1992; Nelson and Fisher 1995; Stewart and Cavanaugh 2006; Bright and Lallier 2010; Childress and Girguis 2011; Hilário et al. 2011). With its enormous size (Fisher et al. 1988a; Hessler et al. 1988; Shank et al. 1998), rapid cell proliferation (Pflugfelder et al. 2009), seemingly fast growth (Lutz et al. 1994; Lutz et al. 2001), but short life (Klose et al. 2015) one of the most puzzling findings was the lack of a digestive system in an animal with a highly unusual body plan (Jones 1981). Descriptions of mouth- and gutless pogonophoran relatives go back a century (Caullery 1914). The first vestimentiferans *Lamellibrachia barhami* Webb, 1969 and *L. luymesi* van der Land and Nørrevang, 1975 were described already a few years earlier than *Riftia.* However, it was the discovery of *Riftia,* thriving in an apparently poisonous hydrothermal vent environment, which sparked the discovery of the first-described chemosynthetic animal-microbe symbiosis (Cavanaugh et al. 1981); an association in which *Riftia*, without a mouth or a gut, relies on the sulphide oxidizing chemoautotrophic symbionts for nutrition (Cavanaugh et al. 1981; Felbeck 1981; Rau 1981a; Rau 1981b) (Arp and Childress 1981; Arp and Childress 1983).

Despite the fact that neither the animal host, nor the symbiont, nor the intact association are amenable to long-term cultivation, *Riftia* is easily one of the best studied deep-sea animals which have consistently led to major discoveries (reviewed by Bright and Lallier 2010). Crucial was the development of various devices to measure chemical and physical parameters directly in the deep sea to understand the abiotic conditions under which this tubeworm thrives at vigorous diffuse vent flow (Hessler et al. 1988; Shank et al. 1998; Luther et al. 2001; Le Bris et al. 2003; Mullineaux et al. 2003; Le Bris, Govenar, et al. 2006; Le Bris, Rodier, et al. 2006). Unprecedented and equally important was the development of high-pressure flow-through systems to simulate in situ conditions in the lab (Quetin and Childress 1980; Girguis et al. 2000).There has been probably no deep-sea animal with more resourceful experimental approaches applied *in situ* and *ex situ* than *Riftia,* e.g. catheterised tubeworms under flow-through pressure (Felbeck and Turner 1995), artificial insemination and developmental studies under pressure (Marsh et al. 2001), predation experiments with mesh cages *in situ* (Micheli et al. 2002), hydraulically actuated collection devices of tubeworm aggregations (Hunt et al. 2004; Govenar et al. 2005), artificial plastic tube deployments (Govenar and Fisher 2007), pressurized experiments (Goffredi et al. 1997; Shillito et al. 1999; Girguis et al. 2000; Girguis et al. 2002), and finally, various *in situ* settlement devices for tubeworm larvae (Mullineaux et al. 2000; Nussbaumer et al. 2006; Mullineaux et al. 2020). These innovative experiments associated with four decades of research taught us about many aspects of *Riftia’*s evolution and biology.

After many microanatomical studies accompanied by heated, highly controversial phylogenetic discussions, the question of who the closest relatives of *Riftia* are was ultimately solved by traditional cladistic and novel molecular analyses (Fauchald and Rouse 1997; McHugh 1997; Halanych et al. 2001; Rouse 2001; Schulze 2002). They showed that vestimentiferans are lophotrochozoan polychaetae worms within Annelida (Fig. 1A) (Polychaeta, Siboglinidae, Vestimentifera) (Pleijel et al. 2009). Similar to many other polychaetes, *Riftia* is gonochoristic with internal fertilization and undergoes a biphasic life cycle with a pelagic phase including indirect development through spiral cleavage and a trochophore larvae (Marsh et al. 2001). The benthic phase is marked by the uptake of the symbiont into the metatrochophore larvae and growth into an adult, which completely reduces its mouth, gut, and anus. Instead, a unique mesodermal nutritional organ, the trophosome, functionally replaces the digestive system (Nussbaumer et al. 2006; Bright et al. 2013). The adult body is organized into four distinct regions, the obturacular region, the vestimentum, the trunk and the opisthosoma (Fig. 1B). The anterior obturacular region of the animal projects a vascularised branchial plume, which is responsible for the sequestration of nutrients and gas exchange, followed by the vestimentum, a muscular head region enclosing the heart, brain, the excretory organ, and the gonopores. The trunk region, the single elongated first segment, harbours the trophosome and the gonads. The posterior part, the opisthosoma, contains a typical segmented annelid region with serially arranged chaetae (Bright et al. 2013). It is so far unknown how this unusual body plan lacking the entire digestive system is reflected in their developmental genes and signalling pathways. Gutless parasitic tapeworms, e.g. have lost many developmental genes including all ParaHox genes (Tsai et al. 2013).

**Figure 1.**
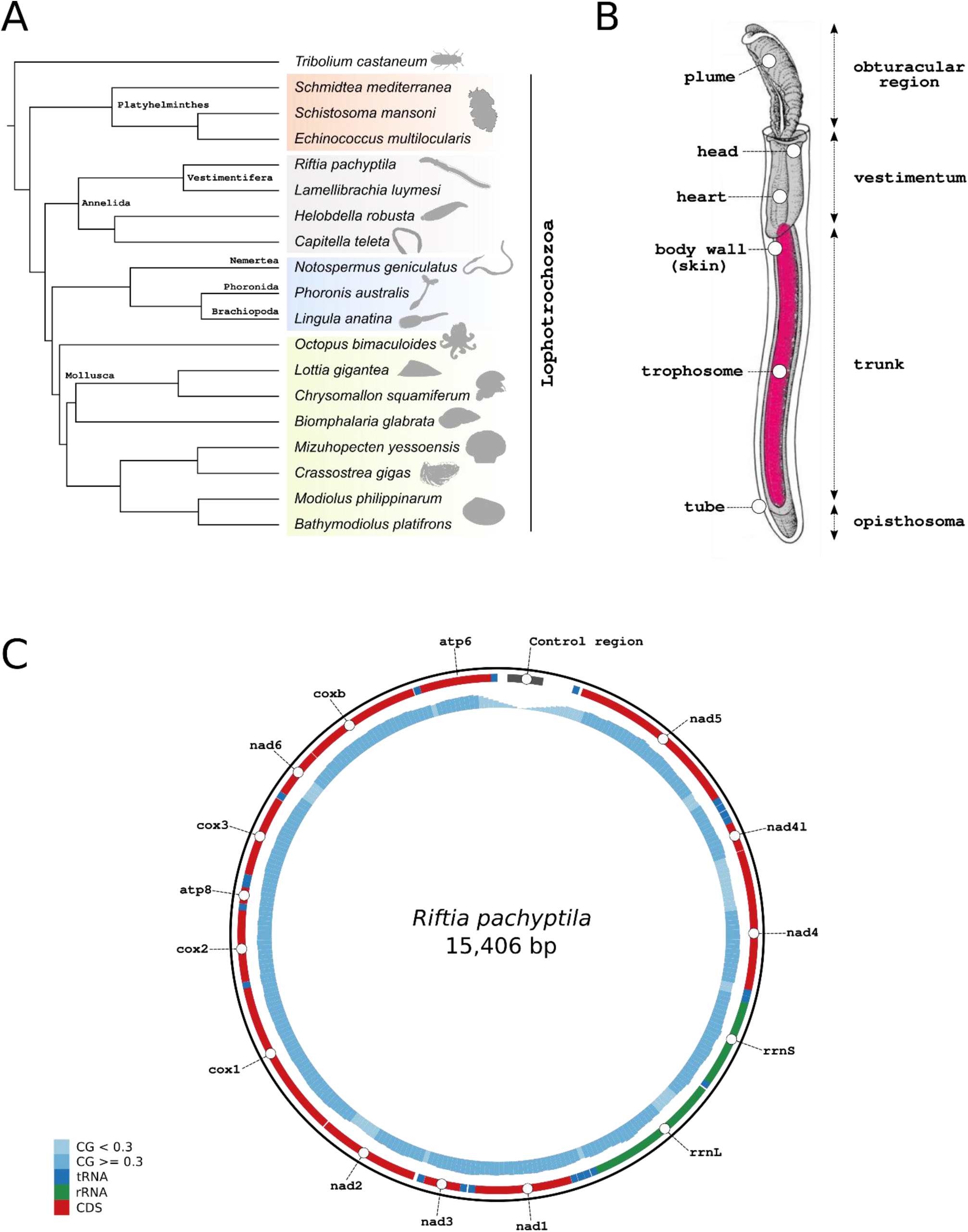
Overview of *Riftia pachyptila* body plan anatomy, phylogenetic placement and mitochondrial genome. **A**, Phylogetic placement of *Riftia pachyptila* within Lophotrochozoa. *Riftia* together with *Lamellibrachia* forms the clade Vestimentifera, a clade of marine animals living in chitinous tubes and lacking a digestive tract. Animal silhouettes were download from http://phylopic.org/. Tree topology was obtained through phylogenomic analysis. **B**, Schematic drawing of *Riftia pachyptila* adult. The first part of the body, the obturacular region, contains the highly vascularised plume, whereas the head, heart and gonads are located in the second body part, the vestimentum. The trunk region and third body part harbors the trophosome (organ that houses the symbiotic bacteria), body wall (skin). The posterior part, the opisthosoma is the fourth and last body region of the tubeworm. Schematic drawing was modified from Nussbaumer et al. (2006). **C**, Schematic representation of *Riftia pachyptila* mitochondrial genome, including the complete control region. CG-content and tRNA genes are represented by the blue histograms and boxes, respectively.

The trophosome of *Riftia*, a soft multi-lobed and highly vascular tissue, houses a polyclonal endosymbiotic population dominated by one genotype of *Candidatus* Endoriftia persephone, a chemoautotrophic gammaproteobacteria (Robidart et al. 2008; Gardebrecht et al. 2012; Polzin et al. 2019) that oxidizes sulphur compounds via oxygen and nitrate and, in turn, harnesses that energy to fix dissolved inorganic carbon (or DIC, which includes carbon dioxide and bicarbonate) to organic matter. Briefly, the trophosome is far removed from the external environment, so the host presumably provides all of the inorganic nutrients to the symbionts. This primarily occurs via the highly vascular brachial plume, which takes up oxygen and hydrogen sulphide (H_2_S) from the external environment and transports these to the trophosome via a complex and unique complement of haemoglobins (Arp and Childress 1981; Arp and Childress 1983; Arp et al. 1987; Zal et al. 1996; Zal et al. 1997; Bailly et al. 2002; Flores et al. 2005). DIC is also taken up by *Riftia*, which is unusual as carbon dioxide is an animal respiratory waste product. However, in this case the worm must provide additional DIC to the symbionts for *net* carbon fixation, and does so by accumulating DIC in the blood (e.g. Goffredi et al. 1997; Goffredi et al. 1999). Moreover, physiological studies have shown that *Riftia* also takes up nitrate (also unusual for an animal), and in turn the symbionts reduce it to organic nitrogen (e.g. Hentschel et al. 1993; Girguis et al, 2000). In return, the host is nourished through the symbiont releasing organic matter and symbiont digestion, which occurs prior to bacteriocyte death in the periphery of the trophosome lobules (Felbeck 1985; Hand 1987; Felbeck and Jarchow 1998; Bright et al. 2000; Hinzke et al. 2019). Despite four decades of research, key questions about trophosome function remain, including but not limited to A) which of the two nutritional modes is more important (organic matter release or symbiont digestion; Bright et al. 2000) and B) the mechanisms that underlie organic nitrogen synthesis and distribution between the symbionts and the host.

Despite the highly derived annelid body plan, symbiotic lifestyle, and over forty years of extensive physiological research, whole genome information of *Riftia* has been lacking. Here, we generated a high-quality genome draft and distinct tissue-specific transcriptomes of the giant gutless tubeworm *Riftia*. By analysing the genome and transcriptomes of *Riftia* in a comparative framework, we highlight many evolutionary adaptations related to the obligate symbiotic lifestyle and survival in the deep-sea hydrothermal vent environment. The *Riftia* genome, together with a transcriptome and proteome study (Hinzke et al. 2019), a transcriptome study on the close relative *Ridgeia piscesae* (Nyholm et al. 2012), the genomic resources available for another close relative *Lamellibrachia luymesi* (Li et al. 2019) (short *Lamellibrachia*), and an extensive body of research broadens our understanding of one of the most conspicuous models for host-symbiont interaction and of the biology of Vestimentifera. Most importantly, we show that the developmental gene repertoire is conserved, and that besides the well-known nutritional aspect of the trophosome, its mesodermal origin brought an inherited suite of functions such as, haematopoiesis, endosomal digestion of endosymbionts, and storage of excretory products likely adapted to serve host-symbiont physiological interactions. While the innate immune system apparently is little upregulated in the presence of the symbiont, it is highly active in the remaining body directly exposed, or connected through openings to, to the environment.

## Results and discussion

### *Riftia* represents the most complete annelid genome to date including a complete mitogenome

To assess the whole genome content of the giant tubeworm (Jones 1981), we sequenced a single individual from the hydrothermal vent site Tica, East Pacific Rise 9° 50’ N region, with ∼87-fold coverage using Pacific Biosciences Sequel system (Supplementary Figures 1-3; Supplementary Table 1). We found the haploid genome size (560,7Mb with a N50 length of ∼2,8Mb) to be smaller than previous genome-size estimates (Bonnivard et al. 2009) (Table 1- Supplementary Figure 4). The *Riftia* GC value is 40.49%, and the repeat content accounts for 29.99% of the total length of the genome with most of the repetitive landscape dominated by interspersed and unclassified lineage-specific elements (35.2%) (Supplementary Figure 5). After genome post-processing, we identified a total of 25,984 protein coding genes with homologue, transcriptome, *ab-initio,* and gene expression evidence. The BUSCO4 (Simão et al. 2015) genome completeness score is 99.37%. These numbers render *Riftia* the most complete annelid genome to date (Simakov et al. 2013; Li et al. 2019; Martín-Durán et al. 2020) (Supplementary Figure 6).

The complete reconstruction of siboglinid mitochondrial genomes including the AT-rich control region has been notoriously difficult (Li et al. 2015). In this case, we were able to obtain it due to deep long read sequencing. The 15,406 bp circular mitochondrial genome contains all expected 13 coding sequence genes, two ribosomal RNA genes and the 22 tRNAs, typical of bilaterian mitogenomes (Fig. 1C – Supplementary Figure 7) (Boore 1999). In contrast to two other *Riftia* reference mitogenomes (Jennings and Halanych 2005; Li et al. 2015), we recovered the full control region (D-loop), yielding a mitochondrial genome longer than those previously reported. The gene order and the number of genes are conserved among all three *Riftia* and other siboglinids reference mitogenomes, though there are size differences that are most likely due to the incomplete nature of previously published genomes.

### The developmental gene repertoire in gutless *Riftia* is conserved

Because of the lack of molecular information on the development of cell types and the evolution of the vestimentiferan body plan, we identified and annotated a suite of key developmental genes and signalling pathway-related genes in the giant tubeworm genome. We found that key genes involved in the development of the digestive tract in metazoans (Hejnol and Martín-Durán 2015; Nielsen et al. 2018), such as *goosecoid, brachyury*, *foxA* and all three ParaHox genes, *xlox*, *cdx* and *gsx*, present in the *Riftia* and *Lamellibrachia* genomes (Supplementary Figure 8-10). The conservation of these genes in vestimentiferans is apparently not only crucial for developmental processes but also serves the microphagous nutrition in settled larvae until nourishment by the symbionts takes over in juveniles (Nussbaumer et al. 2006).

The Hox cluster (∼578kb in size – Fig. 2A), homeodomain-containing transcription factors with roles in anterior-posterior axial identity in metazoans (Pearson et al. 2005; Duboule 2007), is nearly intact and complete in the giant tubeworm genome (Supplementary Figures 8-9, Supplementary note 2). We did not identify *hox7* in *Riftia*, indicating a secondary loss of this gene in the giant tubeworm, a pattern also observed in other lophotrochozoan representatives such as phoronids (Luo et al. 2018) and bivalves (Gerdol et al. 2015; Calcino et al. 2019). *Hox7*, *lox*2 and *lox5* are missing from *Lamellibrachia* genome suggesting a possible loss of the central Hox cluster elements (Fig. 2B) (Li et al. 2019). The Hox-like elements, homeotic genes equally important for body plan specification and developmental processes, *gbx*, *evx*, *mox*, *mnx, en* and *dlx* were also found in the giant tubeworm genome. *Engrailed* (*En*) and *even-skipped* (*Evx*) have 2 and 4 copies, respectively (Supplementary Figure 8).

**Figure 2.**
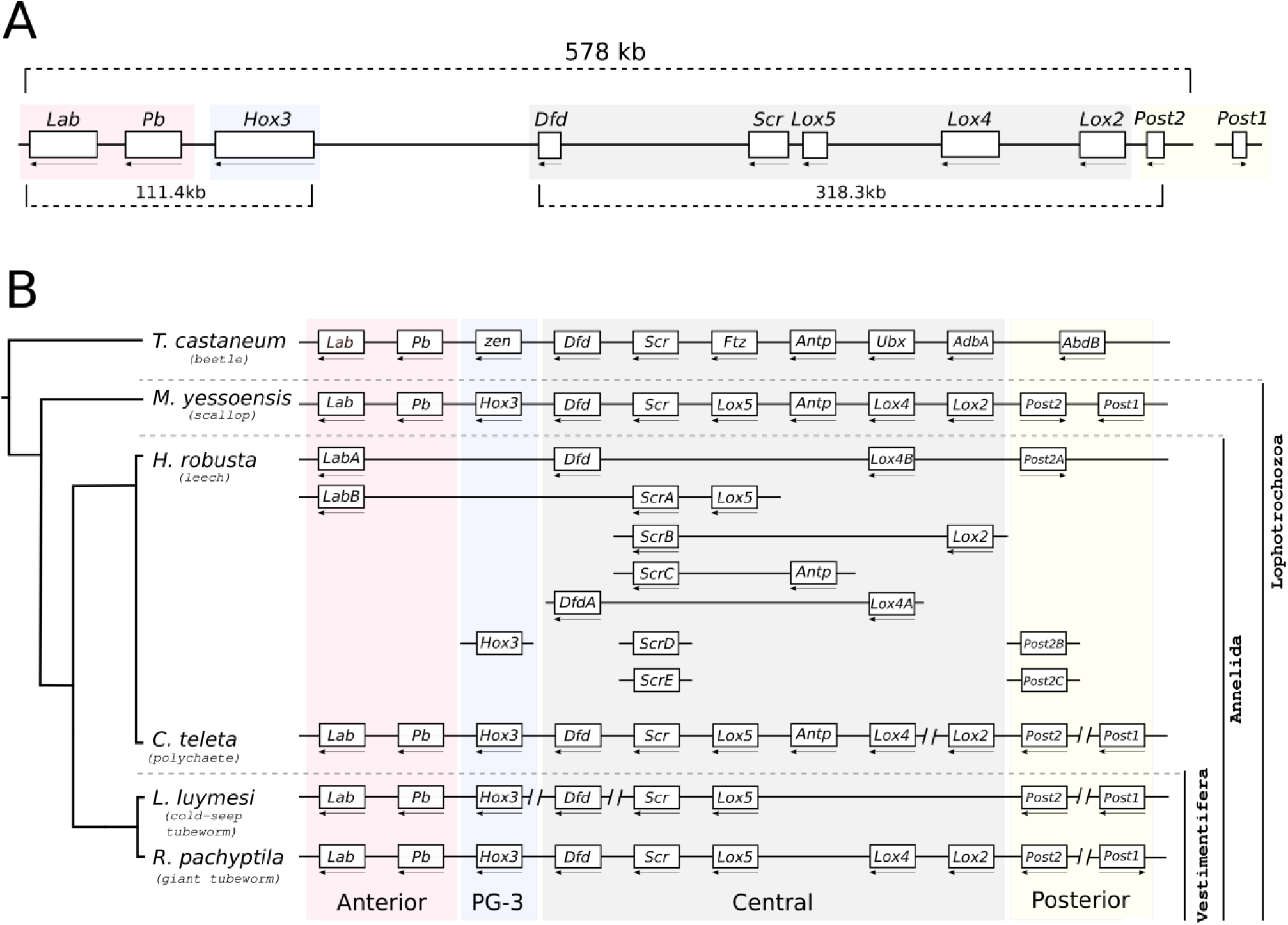
*The* Hox gene complement of *Riftia pachyptila* and selected metazoans. **A**, Hox cluster organisation in the genome of *Riftia pachyptila*. Nine out of the ten Hox genes are located in one single genomic scaffold. *Hox7* is missing from the giant tubeworm genome. Arrows indicate direction of transcription. Only the longest gene model is shown. **B**, Hox cluster present in selected metazoans. *Riftia* presents the most intact Hox cluster among annelids. Part of the central Hox class is missing from the cold-seep tubeworm *Lamellibrachia*. *Helobdella* and *Capitella* cluster are adapt from Simakov et al. (2013), whereas *Mizuhopecten* cluster is from Wang et al. (2017).

Few signalling pathways are required to control cell-to-cell interactions and produce the plethora of cell types and tissues in Metazoa (Pires-daSilva and Sommer 2003) among them TGFβ, Wnt, Notch and Hedgehog (Moustakas and Heldin 2009; Ingham et al. 2011; Holstein 2012; Massagué 2012; Niehrs 2012; Gazave et al. 2017) (Supplementary Figures 11-15) The *Riftia* genome contains 14 TGFβ genes, including *nodal* and its antagonist *lefty*, the latter previously assumed to be a deuterostome innovation (Simakov et al. 2015). *Notch* and *hedgehog* are present as single copy genes in the *Riftia* genome as well as in *Lamellibrachia*, however, the *notch* receptor *jagged* is missing from both tubeworms. *Jagged* is present in the annelids *Capitella teleta*, *Helobdella robusta* and *Platynereis dumerilii* (Gazave et al. 2017), suggesting a secondary loss in Vestimentifera. Patched and dispatched genes, membrane receptors for the hedgehog ligand (Ingham et al. 2011) are present in *Riftia* with the dispatched genes expanded in vestimentiferans. In *Riftia*, we identified the 12 expected Wnt ligands (*Wnt3* has been shown to be lost in the Protostomia lineage) and their receptors frizzled, smoothened and sFRP (Holstein 2012). There is a genetic linkage of *Wnt1*,*6, 9* and 10 in *Riftia* akin to the gastropod *Lottia gigantea* and the fruit fly *Drosophila melanogaster* (Cho et al. 2010), reaffirming the ancient protostomian ancestral conserved linkage. The remaining eight Wnt genes in *Riftia* are disorganized on eight different scaffolds. Overall, despite the highly derived body plan, *Riftia* presents a deep conservation of the developmental gene toolkit akin to many distinct bilaterian animals.

### The *Riftia* genome is characterised by reductive evolution

Multiple lines of evidence point to a relatively small genome, with gene family contractions exceeding expansions in *Riftia*, indicative of reductive evolution. The giant tubeworm genome is ∼168Mb smaller than *Lamellibrachia,* its relative from cold hydrocarbon seeps whose genome is ∼688 Mb with a N50 of 373kb (Li et al. 2019)). The difference can be attributed to the increased number of repeat elements and protein coding genes in the cold seep tubeworm (38,998 gene models and repetitive content of 36.92%; Supplementary Note 1; Supplementary Figure 16).

To identify clusters of orthologous genes shared among the two vestimentiferans *Riftia* and *Lamellibrachia,* the polychaete *Capitella teleta* (herein called *Capitella*), and the clitellid *Helobdella robusta* (herein called *Helobdella*) (Simakov et al. 2013), we employed tree-based orthology inferences (Emms and Kelly 2019). The annelid core genome, the collection of orthogroups shared among the four annelids, contains 6,349 cluster of orthologous genes. Less than half of them represent the vestimentiferan core genome (2,883 orthogroups) shared between *Riftia* and *Lamellibrachia*. Interestingly, the number of shared orthogroups between the *Riftia-Helobdella* (17) and -*Capitella* (116) pairs are smaller than those between *Lamellibrachia*-*Helobdella* (89) and - *Capitella* (349) pairs, indicating that *Riftia* contains a more derived gene repertoire than its close relative *Lamellibrachia*.

To further investigate the important processes of gene losses and gains, known to shape animal evolution (Fernández and Gabaldón 2020; Guijarro-Clarke et al. 2020), identify expanded protein domains, taxonomically restricted genes, and positively selected genes in *Riftia*, we employed multi-level comparative approaches involving statistical analysis, taxon rich orthology inferences (N=36), and sensitive similarity searches (Supplementary Table 2; Supplementary Note 3). The *Riftia* genome shows a net reduction of gene numbers with only 734 expanded but 1,897 contracted gene families, whereas the evolutionary history of *Lamellibrachia* is characterised by gene gains. Notably, the average expansion value of gene families in *Riftia* is the lowest among the four selected annelids herein analysed (Supplementary Figure 17). A total of 8,629 lineage-specific genes (∼33.21% of the total) were identified in *Riftia*. Compared to the giant tubeworm, *Lamellibrachia* contains more lineage-specific genes (10,262 – 26.31% of the total).

The contracted gene families are not restricted to any specific biological process, as revealed by our gene ontology (GO) term enrichment analysis (N=18) (Supplementary Figure 18). Rather it appears that the giant tubeworm genome is undergoing a broad reduction in gene content (i.e., reductive evolution). Among the contracted gene families are genes controlling the transcriptional machineries (Supplementary Figure 19; Supplementary Table 3). Transcription factors (TFs) are proteins with sequence specific DNA-binding domains that control gene transcription and tissue identity (Schmitz et al. 2016). To gain understanding into the repertoire of TFs in *Riftia*, we annotated and classified genes in the tubeworm genome present in five major groups of TFs (bzip, homeobox, nuclear factor, bHLH and zinc-finger) with sensitive similarity searches. The giant tubeworm presents the lowest number of TFs within the analysed annelids (414), supporting our gene family analysis (discussed below). The cold-seep tubeworm genome contains a similar complement size as *Riftia* (423), with *Capitella* (551) and *Helobdella* (568) presenting a higher number of TF genes, comparatively. These results point to pervasive TF losses in the Vestimentifera lineage (Supplementary note 3).

### Expanded and lineage specific gene families in *Riftia*

Despite overall genome reduction, the *Riftia* genome exhibits, there is also a variety of expanded gene families (Supplementary note 3; Supplementary Figure 20; Supplementary Table 4). These expanded families are enriched with GO terms associated with sulphur metabolism, membrane transport and detoxification of xenobiotic, e.g. foreign substances (xenobiotic transmembrane transporter activity, galactosylceramide sulfotransferase activity, CoA-transferase activity) (Gamage et al. 2006), detoxification of hydrogen peroxide as anti-oxidative stress response (glutathione catabolic and biosynthetic processes) (Espinosa-Diez et al. 2015), neurotransmitter- and ion channel-related functions (sodium symporter activity), oxygen transport (oxygen binding, haemoglobin complex), endosomal degradation (lysozyme activity), and secretion of chitin (chitin binding, protein glycosylation) (discussed with more details later).

Genes involved in the production of extracellular components of vestimentiferans such as the cuticle and the basal matrixes as well as the tube and chaetae (Gardiner and Jones 1994) were found in expanded families of *Riftia* (as well as *Lamellibrachia*), some of which are specific to either *Riftia* or *Lamellibrachia* (Supplementary note 3; Supplementary Figures 21-24; Supplementary Tables 5-6). The *Riftia* genome contains expanded protein domains related to several high-molecular mass proteins such as laminin, nidogen and collagen. These proteins are part of extracellular matrix secreted basally from epithelia, also known to regulate cellular activity and growth in other animals (Timpl and Brown 1996). In *Riftia*, extensive short collagen fibres are found below the epidermis, extending between muscles cells, and building the matrix of the obturaculum. In addition, long helically arranged collagen fibres are the main component of the cuticle apically secreted from the epidermis (Gardiner and Jones 1994). Importantly, many genes involved in chitin production, a biopolymer part of the hard protective tube secreted from pyriform glands of the vestimentum, trunk, body wall, and opisthosoma (Gardiner and Jones 1993), are taxonomically restricted to the *Riftia* lineage. Expectedly, we identified in the vestimentum and body wall tissues of *Riftia* several tissue-specific genes (TSGs) involved in the chitin metabolism responsible for the tube production as well as dissolution (Supplementary Figures 25-26). Although the specific gland type responsible for dissolution of tube material has yet to be identified, we suggest that in *Riftia* the straight tube, that can reach up to three metres in length and five centimetres in diameter (Gaill and Hunt 1986; Grassle 1987; Fisher et al. 1988b), can only widen in diameter to accommodate growth of the worm when tube material is dissolved and newly secreted, which agrees with the distribution of many TSG involved with tube biosynthesis. Overall, our findings of these expanding gene families as well as gene expression patterns underline the importance of chitin in *Riftia,* which is considered one of the fasted growing invertebrates (Lutz et al. 1994. Lutz et al. 2001). In order to achieve these high growth rates, *Riftia* needs to both digest and remodulate its own tube with astonishing speed.

Furthermore, the multi-level comparative analyses revealed an enrichment of GO terms in the lineage specific *Riftia* genes involved with the control of the chromosome condensation and nucleosome assembly, and positively selected genes related to tumour suppression (*PIN2/TERF1*-interacting telomerase inhibitor) and transcription initiation (*TFIIB*- and *-D*) (Roeder 1996; Zhou and Lu 2001). Interestingly, in *Lamellibrachia smad4* (Li et al. 2019), which is a tumour suppressor and transcription factor, is under positive selection, suggesting a common vestimentiferan evolutionary adaption responsible for controlling the chromatin-remodelling events and the extraordinarily cell proliferation rates in these two tubeworms (Supplementary Table 7; Supplementary note 3) (Pflugfelder et al. 2009).

The protein annotation of the rapidly evolving expanded gene families in *Riftia* identified members of the complement system involved in innate immunity and self-, non-self-recognition (sushi repeat domain-containing protein) (Kirkitadze and Barlow 2001). *Riftia* contains the greatest number of sushi-domain containing proteins among lophotrochozoans, presenting a total of 42 copies which are organised either in genomic clusters or dispersed as single elements throughout the genome (Supplementary Figure 4; Supplementary Figures 27). Of these, only 40 are shared with the cold seep tubeworm *Lamellibrachia*, pointing to a lineage-specific expansion at the base of Vestimentifera. Sushi genes, a common component of haemocytes (i.e., immune cells with phagocytic function (Pila et al. 2016), have been implicated in the mediation of the host-symbiont tolerance in the bobtail squid *Euprymna scolopes –* bioluminescent *Aliivibrio fischeri* association (McAnulty and Nyholm 2017). Although the rapid evolution of these proteins in *Riftia* and *Lamellibrachia* suggests similar evolutionary adaptations to the tubeworm/endosymbiont mutualism, the absence of any significant expressions in adult tissues rather point to their involvement in recognition of the symbiont during transmission in the larval stage or to potential pathogen recognition upregulated upon exposure.

### Substrate transport for energy conservation and biosynthesis is supported by lineage-specific adaptations and parallel evolutionary events in *Riftia*

As an adaptation to the sulphidic vent environment, and in support of a symbiotic lifestyle, the respiratory pigments in *Riftia*, and other vestimentiferans such as *Lamellibrachia*, bind non-competitively and reversibly to oxygen and sulphide, simultaneously providing a key substrate for chemosynthesis by the symbionts while also averting the sulphidic inhibition of the hosts’ mitochondrial oxidative chain reactions (Arp and Childress 1983; Terwilliger et al. 1985). Our previous gene family evolution analysis identified an expansion of haemoglobins in the giant tubeworm genome compared to other non-vestimentiferan lophotrochozoans (Supplementary Figure 21; Supplementary Table 6; Fig. 3A). Additionally, a recent genome study found a massive expansion of β1-haemoglobin in the cold seep tubeworm *Lamellibrachia* (Li et al. 2019). To gain better insights in the evolution of Hb and linker genes in the Vestimentifera lineage, we employed thorough comparative genomics, phylogenetics, domain composition and gene quantification analyses. The genomic arrangement of the giant tubeworm Hb genes indicates that they were originated through a series of tandem duplications, totalling seven distinct genomic clusters (Fig. 3B). We annotated 26 extracellular Hbs and six linker genes in the *Riftia* genome (Supplementary Figures 28-29; Supplementary Note 4). Twenty-two Hb genes were phylogenetically placed in the β1-Hb group, surpassing previous estimates of the β1-Hb complement in the giant tubeworm (Bailly et al. 2002; Sanchez et al. 2007; Hinzke et al. 2019). α2- and β2-Hbs are found as single copy genes, whereas α1-Hb group contains two paralogous genes. The sulphide-binding ability of the *Riftia* Hbs is associated with the occurrence of free cysteine residues in one α2 and one β2 Hb genes (Bailly et al. 2002), as well as the formation of persulphide groups on linker chains (Zal et al. 1998; Bailly et al. 2002). Our results show that seven additional paralogous genes belonging to the β1-Hb group contain the putative free-cysteine residues, which were confirmed through multiple sequence alignments and homology model generation (Supplementary Figures 30-31, Supplementary note 4). Additionally, it has been hypothesized that zinc ions, rather than free-cysteine residues, are responsible for the H2S binding and transport on vestimentiferan α2 chains (Flores et al. 2005). We identified the three conserved histidine residues (B12, B16 and G9), predicted to bind zinc moieties, in *Riftia* Hb genes. However, we observed variations within the *Lamellibrachia* α2 genes. A broader comparison of α2-Hb genes belonging to different annelid taxa challenged the hypothesis of zinc sulphide-binding mechanisms for H2S in siboglinids and vestimentiferans (Li et al. 2019). Our results, solely based on the conservation of histidine residues, corroborate Flores et al. (2005) hypothesis that zinc residues may be involved in the sequestration and transport of hydrogen sulphide at least on the giant tubeworm.

**Figure 3.**
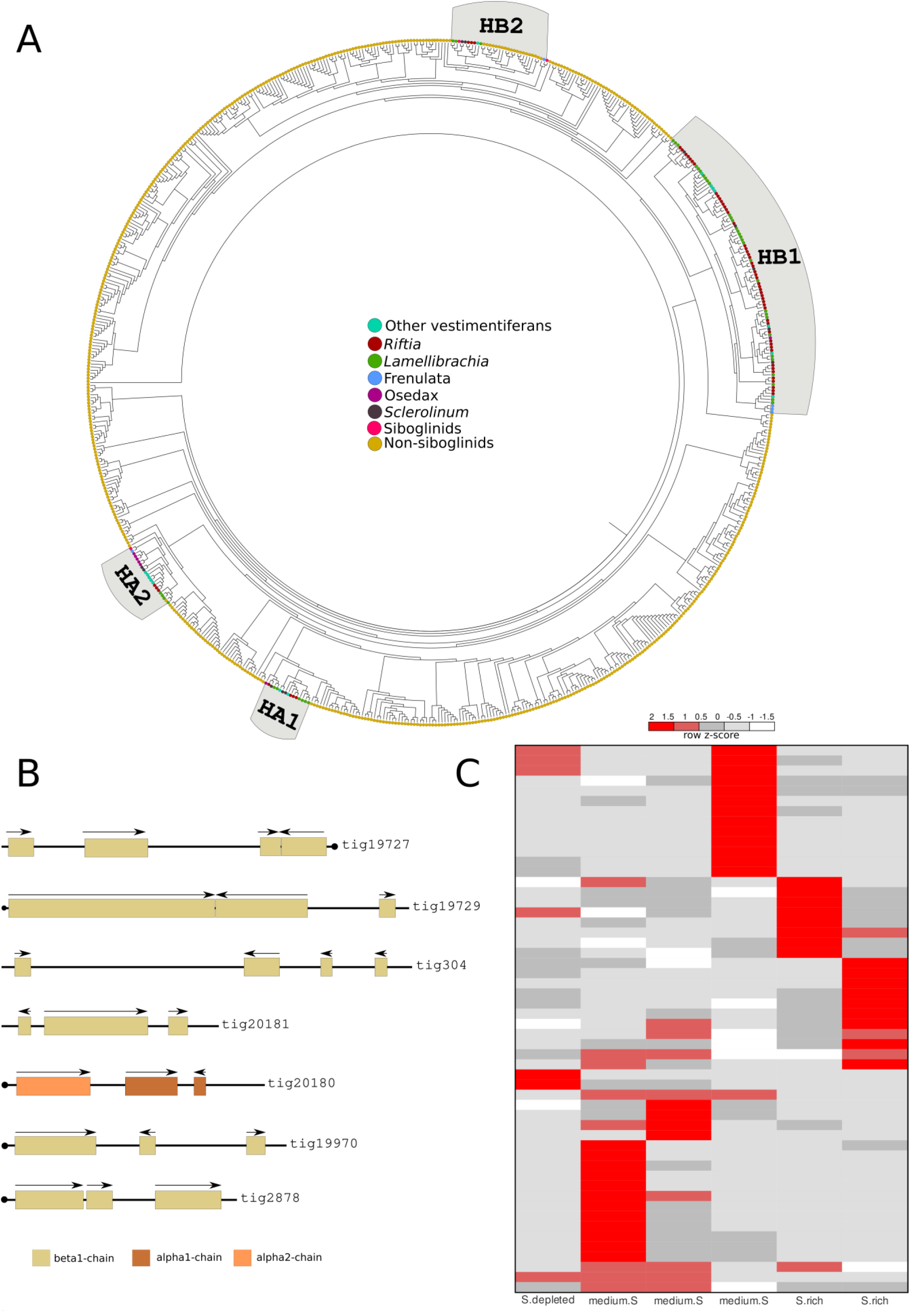
Expanded haemoglobin complement in *Riftia pachyptila.* **A**, Midpoint rooted phylogeny of 693 *Riftia*, annelid and metazoan haemoglobin genes, using Paiva et al. (2019) as backbone. Coloured circles correspond to different annelid taxa and metazoans. **B**, Seven genomic clusters of haemoglobin genes in *Riftia* genome. Arrows indicate the direction of transcription. Scaffolds with a circle on the end indicate the presence of Hb genes in the terminal end of the scaffolds. Only the longest gene models are shown. Colours represent the different haemoglobin chains. Two β1- and one β2-Hbs genes are located in three separate scaffolds (tig3224, 19723 and 19768). **C**, Heat map expression of haemoglobins in the trophosome under three experimental conditions: medium sulphide (medium.S), sulphide rich (S.rich) and sulphide depleted (S.depleted).

To investigate the gene expression dynamics of the newly and previously identified Hb paralogs in *Riftia*, we analysed published transcriptomes sampled from *Riftia’s* trophosomes containing sulphur-rich to sulphur-depleted symbionts (Hinzke et al. 2019). Hb gene expression showed great variation, indicating a more specialised role of the Hbs according to the environmental chemical fluctuations in the unstable deep-vent ecosystem (Fig. 3C, Supplementary note 4; Supplementary Table 8). Taken together, these results suggest a more complex system coordinating oxygen-sulphide sequestration and distribution in the giant tubeworm tissues. The Hb complement of *Riftia* and *Lamellibrachia* are similar and unique among annelids and lophotrochozoans, in respect to gene numbers and distribution, indicating a Vestimentifera synapomorphy.

As the endosymbionts require carbon dioxide (CO_2_) for fixing inorganic carbon, the transport of CO_2_ and the conversion of its alternative forms (e.g., bicarbonate; HCO3^-^) is mediated by another class of enzymes, the carbonic anhydrases (CAs) (Shively et al. 1998; Cian et al. 2003). We found ten carbonic anhydrase genes in the *Riftia* genome, from which seven are tandemly arrayed in two genomic clusters (Supplementary Figure 32). A similar CA complement in *Riftia* (nine genes) was found in a recent study (Hinzke et al. 2019). To better understand the diversity of CA genes we analysed tissue-specific transcriptomes and found at least five CA genes are membrane bound with three of them moderately/highly expressed in the trophosome, indicating that HCO3^-^ conversion to CO2 and diffusion across the bacteriocyte membrane might be a common process in the trophosome, as suggested previously (Sanchez et al. 2007; Bright and Lallier 2010; Hinzke et al. 2019). Tandem duplications and tissue-specific CA expression linked to the intracellular supply of CO_2_ to endosymbionts have been recently reported in deep-sea bivalves (Ip et al. 2021), showing remarkable resemblance to our findings. Taken together, our results show that the transport of essential compounds to the chemoautotrophic endosymbionts and the maintenance of the mutualistic relationship is driven by lineage-specific and parallel evolutionary events.

### Trait and gene loss is compensated by the endosymbionts

The loss of the digestive system requires nourishment through the symbiont. The mechanisms of carbon transfer between the endosymbiont and *Riftia* were shown to be through the fast release of fixed carbon from the symbiont and uptake into host tissue, as well as, through symbiont digestion prior death of the bacteriocytes (Felbeck 1985; Hand 1987; Felbeck and Jarchow 1998; Bright and Lallier 2010; Hinzke et al. 2019). We found corroborating evidence for the uptake of released organic carbon from the symbiont based on the enrichment of GO terms and tissue specificity of succinate-semialdehyde complex genes and nuclear-encoded proteins of the inner mitochondrial membranes (including the tricarboxylate mitochondrial carrier responsible for the transport of succinate) (Majd et al. 2018) in the trophosome (Supplementary Figure 33; Supplementary Table 9). These results suggest an increased movement of cytosolic succinate through the mitochondrial membrane, possibly increasing the ATP production via the oxidative metabolism. These findings corroborate previous findings and support the involvement of this molecule for nourishment in *Riftia* from its endosymbiont (Felbeck and Jarchow 1998).

Evidence of digestion was revealed with tissue-specific transcriptome analyses which allowed us to identify the genes involved in the successive stages of lysosomal-associated degradation of symbionts (Supplementary Figure 33; Supplementary Tables 8). The expression of genes associated with endosomal activity, the expression of several lysosomal-associated hydrolases (Supplementary Figures 34-35), vacuolar ATPases, and small Ras-related GTPases (rab genes) (Supplementary Figure 36; Supplementary Note 5) in the trophosome, is indicative of lysosomal-associated degradation of symbionts, as suggested earlier in ultrastructural studies, which describe the presence of primary lysosomes and symbionts in different lytic stages in *Riftia* (Bright et al. 2000; Bright and Sorgo 2003; Hinzke et al. 2019). In addition, we detected specific genes in the trophosome which are associated with actin cytoskeleton dynamics (ARP2/3) known to be essential for endosomal dynamics (Kast and Dominguez 2017) (Supplementary Figure 33; Supplementary Table 9). Recent *de novo* tissue-specific transcriptomes and gene expression quantification of the host and symbiont support the digestive route of nutrition in *Riftia*/Endoriftia symbiosis (Hinzke et al., 2019).

Furthermore, genes involved in the transport of fatty acyl units into the mitochondrial matrix (e.g., carnitine/acylcarnitine carrier) (Indiveri et al. 2011), tricarboxylic acid (TCA) cycle, oxidative phosphorylation, antioxidant systems (i.e., superoxide dismutase II genes and methionine sulfoxide reductases) (Supplementary Figure 38), and key players of the fatty acid β-oxidation (Fig. 4A, B and C), showed tissue specificity in the trophosomal tissue (Supplementary Table 9; Supplementary note 5). Fatty acid β-oxidation is a central and deeply conserved energy-yielding process that fuels the TCA cycle and oxidative phosphorylation (Houten et al. 2016). As *Riftia* relies solely on its endosymbionts for sustenance, the metabolism of fatty acids in the trophosome is certainly linked to the bacterial digestion in this tissue, which is corroborated by a previous proteomic study (Hinzke et al. 2019). Altogether, the results point to different modes of nutrient transfer in the trophosome involving the translocation of released nutrients from symbiont to host through succinate, and the digestion of the symbionts by lysosomal enzymes followed by the degradation of fatty acids using the mitochondrial β-oxidation pathway.

**Figure 4.**
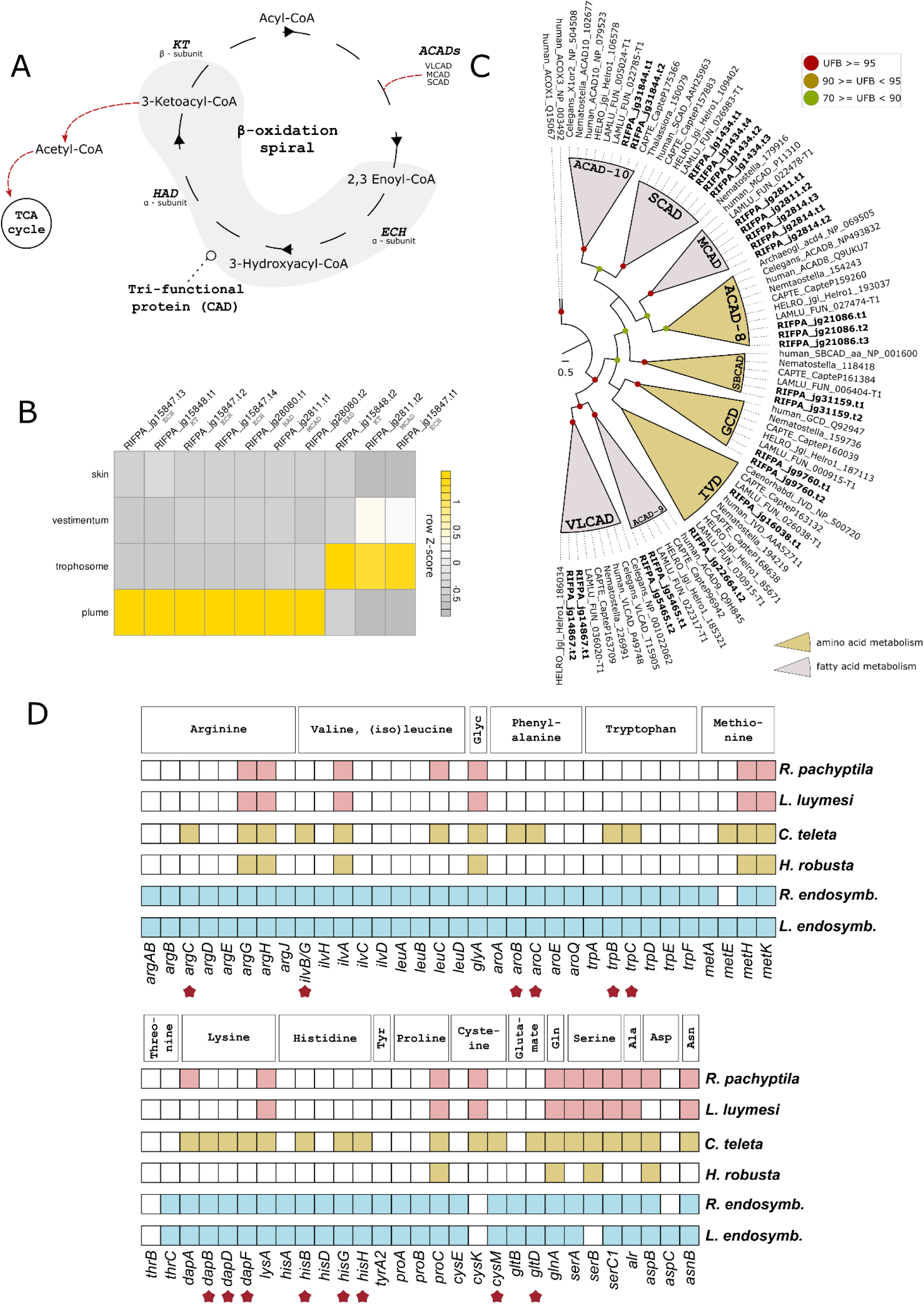
Amino acid and fatty acid biosynthesis in *Riftia***A,** A schematic representation of mitochondrial fatty acid β-oxidation (FAO). The fatty acid degradation is performed in four enzymatic steps involving the membrane bound mitochondrial trifunctional protein and acyl-CoA dehydrogenases. The resulting acetyl-CoA is further oxidised in the TCA cycle. **B**, Expression profile of FAO genes. Colour coding reflects the expression patterns based on row Z-score calculations. The FAO pathway is activated in the trophosome and plume tissues. **C,** Maximum-likelihood phylogenetic tree inference of the ACAD genes using 1000 rapid bootstrap replicates. The branch support values are represented by the coloured circles in the tree nodes. Accession numbers for NCBI database are displayed after the species names and homologs were retrieved from a previous study (Swigoňová et al. 2009). *Capitella, Helobdella* and *Lamellibrachia* gene identification are derived from the publicly available annotated genomes. **D**, Key enzymes related to amino acid biosynthesis identified in *Riftia*, selected annelids and two tubeworm endosymbiont genomes. Identification of the genes was performed with KEGG and reconfirmed with similarity searches against publicly protein dabases. *Riftia* and *Lamellibrachia* lack many amino acid biosynthesis genes indicating nutritional dependence on their endosymbionts. Stars represent genes present in the free-living polychaete *Capitella* and Endoriftia but absent in *Riftia* (based on Li et al. (2019) scheme).

To further explore the extent to which degree *Riftia* is dependent on its endosymbiont for nutrition, we screened the genome of giant tubeworm and selected annelids for key enzymes related to amino acid biosynthesis (Supplementary Table 10). We found that *Riftia*, together with cold-seep tubeworm *Lamellibrachia* and the parasitic leech *Helobdella*, lacks many key enzymes related to amino acid biosynthesis when compared to close free-living polychaete relative *Capitella* (Fig. 4D). Genes involved with amino acid biosynthesis are constitutively expressed across the tubeworm tissues, with enzymes related to arginine and glycine metabolism highly expressed in the trophosome (Supplementary Figure 39). These findings suggest that loss of key enzymes in mutualistic vestimentiferans as well as a parasitic leech may be due to the beneficial and parasitic relationships, respectively allowing for compensated gene loss, compared to free-living polychaetes.

Overall, endosomal-associated digestion of endosymbionts seems to be a hallmark of intracellular digestion accomplished in the mesodermal trophosome of vestimentiferans, such as *Riftia* and *Lamellibrachia* (Nussbaumer et al. 2006; Hinzke et al. 2019; Li et al. 2019) (see also below). This process serves the host nutrition as well as the control of the symbiont population density during host growth, known from many other symbioses (Angela E. Douglas 2010). In addition, the symbiont provides the host with released organic carbon. While we do not know yet which partner controls this mode of nutritional translocation, both the evolutionary adaptation of endosymbiont digestion in a mesodermal tissue as well as carbon release contributed to trait loss in one partner compensated by the other (Ellers et al. 2012), and consequently has made *Riftia* obligatorily associated with its symbiont.

### Haematopoiesis operates in the trophosome of *Riftia*

Haematopoiesis, the production of blood cells and pigments is still a poorly understood process in vestimentiferans. The heart body, a mesodermal tissue in the dorsal blood vessel of vestimentiferans has been hypothesized to be the site of haemoglobin biosynthesis (Schulze 2002). The presence of many TSGs in the trophosome related to 5-aminolevulinate synthase, porphyrin metabolism, and metal ion binding indicate that this tissue harbours the enzymatic machinery necessary for haem biosynthesis. Haem is an integral part of haemoglobin molecules, which is synthesized in a seven multistep pathway that begins and ends in the mitochondrion. To fully characterise the haem biosynthesis pathway in the giant tubeworm, we screened the *Riftia* genome for the presence of the seven universal enzymes required to synthesize the haem (Supplementary Figure 40; Supplementary note 5). The giant tubeworm contains all the seven enzymes present as single copy in its genome with recognisable orthologs in the annelids *Lamellibrachia*, *Capitella* and *Helodbdella.* Gene expression analysis showed that the key enzymes present in the haem biosynthetic pathway are moderately/highly expressed in the trophosome, supporting the GO enrichment analysis (Supplementary Figure 40; Supplementary Figure 33). The haem biosynthesis in the trophosome is further corroborated by the presence of TSGs in this tissue related to phosphoserine aminotransferase and the mitochondrial coenzyme A transporter, which act as an important co-factor in the final step of haem synthesis and in the transport of coenzyme A into the mitochondria, respectively (Schneider et al. 2000; Fiermonte et al. 2009). These findings confirm the involvement of the mesodermal trophosome in haemoglobin metabolism and suggests that this organ is the site haematopoiesis. In the frenulate *Oligobrachia mashikoi*, the visceral mesoderm also strongly expresses globin subunits based on *in-situ* hybridisation and semi-quantitative RT-PCR (Nakahama et al. 2008), but in this siboglinid the visceral mesoderm is organized as simple peritoneum surrounding the endodermal trophosome (Southward 1993).

### Excretory products are stored in the trophosome of *Riftia*

The finding of TSGs in the trophosome related to the biosynthesis of nitrogen-containing compounds and in the transport of ornithine (Supplementary Tables 9 and 11) agrees with the high levels of uric acid and urease activity in this host tissue, as previously reported (Cian et al. 2000; Minic and Hervé 2003).To explore the nitrogen metabolism pathways in *Riftia*, we identified and quantified the gene expression of several enzymes related to the purineolytic/uricolytic, purine/pyrimidine, taurine, and the polyamine pathways, as well as the urea and ammonia cycles (Supplementary Figures 41-45; Supplementary Note 6).

Most of the identified genes are found as single copy in the giant tubeworm genome (Supplementary Note 6; Supplementary Figures 41-45), however, we identified the presence of three chromosomal clusters harbouring glutamine synthetase, cytoplasmatic taurocyamine kinase and xanthine dehydrogenase/oxidase genes (Fig 5A). These enzymes are involved in the ammonia, urea and uricolytic pathways, respectively. *Riftia* and *Lamellibrachia* contain the highest number of glutamine synthetase genes in the herein investigated lophotrochozoan genomes (Fig. 5B). Seven out of the nine glutamine synthetase genes present in *Riftia* belong to the group I and the remaining to the group II (Fig. 5C). Interestingly, only annelid orthologs are found to be phylogenetically close to the prokaryotic group I, indicating a secondary loss of this genes in the remaining lophotrochozoan lineages (see Supplementary Figure 43 for the expanded version of the phylogenetic tree). An expanded set of lengsin genes, an ancient class I glutamine synthetase family (Wyatt et al. 2006), is present in Vestimentifera (seven copies in *Riftia* and 13 in *Lamellibrachia*). Some members of the newly identified lengsins are also highly expressed in the trophosome, suggesting that these enzymes might play a role in mitigating toxicity of urea, ammonia, and other nitrogenous compounds (Wyatt et al. 2006).

**Figure 5.**
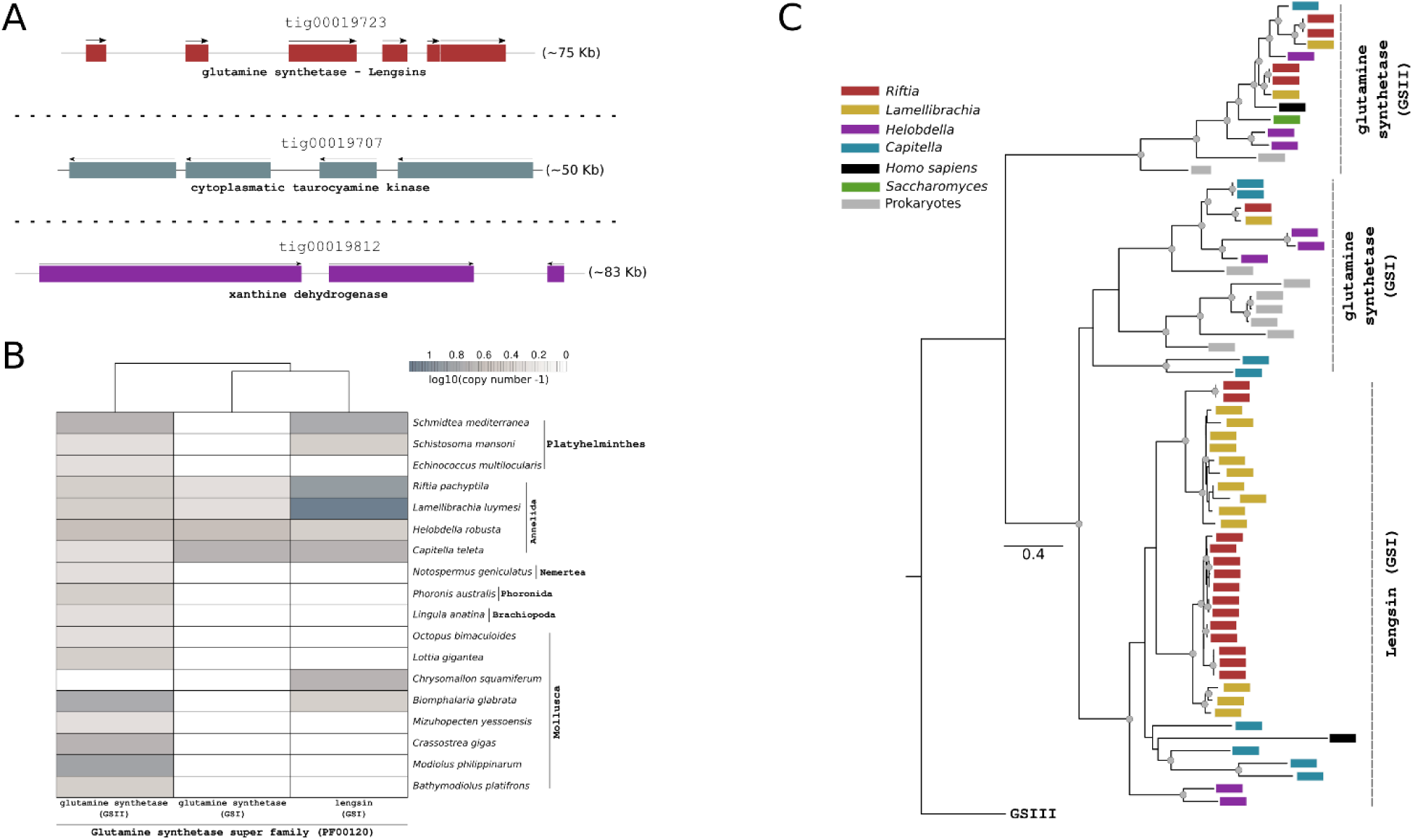
Genomic clusters of important genes related to the nitrogen metabolism in *Riftia*, and distribution of phylogeny of glutamine synthetase genes among selected metazoans. **A,** Genomic organisation of important genes related to nitrogen metabolism in *Riftia*. Arrows indicate the direction of transcription. **B,** Distribution of glutamine synthase-related genes in the giant tubeworm, closely related annelids, and selected lophotrochozoans. **C,** Maximum likelihood phylogenetic tree inference of members of the glutamine synthetase superfamily using 1000 ultrafast bootstrap replicates. Coloured boxes correspond to different annelids, vertebrates, yeast, and prokaryotes.

The identification of five cytoplasmatic taurocyamine kinase genes, four organised in a genomic cluster (Fig. 5A), surpasses previous reports (in which only one cytoplasmatic gene was identified; Supplementary Figure 44) (Uda et al. 2005). In accordance with a recent study (Hinzke et al. 2019) and contrasting previous biochemicals investigations on the *de novo* pyrimidine and polyamine biosynthesis in *Riftia* (Minic et al. 2001; Minic and Hervé 2003), we identified the trifunctional CAD protein in the genome of the tubeworm reinforcing the notion that *Riftia* can catalyse the first steps of the pyrimidine synthesis independently of its endosymbionts (Supplementary Note 6). These results are not unforeseen, since during the aposymbiotic phase (i.e., *Riftia*’s fertilised egg until the settled larva) pyrimidine metabolism plays a fundamental role in the development and growth of the animal.

We also found that the key enzymes of the uricolytic pathway and urea cycle are highly active in the trophosome (Supplementary Figure 41; Supplementary Table 8). These results are consistent with enzymatic/light micrograph studies, which show that the trophosome contains high concentration of ammonia, urea, creatinine, and uric acid crystals in the periphery of the lobules (Cian et al. 2000), and with a more recent transcriptomic and metaproteomic study (Hinzke et al. 2019). Surprisingly, we only identified in closed (De Oliveira, in review) and previous endosymbiont genome drafts the subunit-A of the urea transporter (*urtA*), with the four remaining subunits missing (*urtBCDE*) (Veaudor et al. 2019). Since all five subunits are required for a proper function of the urea transporter, these results challenge the idea of an active shuttle of urea from the host to the endosymbiont (Robidart et al. 2008).

### Cell proliferation and cell death interplay with innate immunity in *Riftia*

To better understand how fast growth (Lutz et al. 1994) fuelled by high proliferation rates (Pflugfelder et al. 2009) and innate immunity act in *Riftia* in tissues exposed to the environment and in the endosymbiont-housing trophosome, we characterised the key molecular components, and their gene expressions, of important pathways related to cell cycle signalling (Supplementary Figure 46), Toll-like receptor/MyD88 (Supplementary Figures 47-49), as well as the apoptotic (Supplementary Figures 50-57) and autophagic (i.e. macroautophagy) cell death events (Supplementary Figures 58). The *Riftia* genome similar to other investigated lophotrochozoans and closely related annelids harbours all the key components of these conserved pathways (Supplementary Note 7) (G. Zhang et al. 2012; Sun et al. 2017; Luo et al. 2018; Li et al. 2019; Ip et al. 2021). We did not identify any extensive remodelling (i.e., gene family expansions and contractions) of the immune (with the exception of sushi genes) and programmed cell death components in the giant tubeworm genome, as shown to be important in the maintenance of host-symbiont interactions in deep-sea mussels and clams (Sun et al. 2017; Ip et al. 2021).

Overall, *Riftia’s* gonad and plume tissues are highly active in cell proliferation and programmed cell death. Subject to potential pathogen infections through the gonopore opening and direct contact to the vent water (Jones 1981), respectively, these tissues show the entire suite of genes involved in the innate immunity recognition with TLRs, downstream cellular immune responses, as well as apoptosis, autophagy and endosomal-related genes. These results were additionally supported by the GO enrichment analyses in the female gonad and plume tissues (Supplementary Figures 59-60). The trophosome, in contrast, despite the remarkably high bacterial population density (Powell and Somero 1986; Bright and Sorgo 2003) does not show any striking upregulation of TLR for endosymbiont recognition, nor cell proliferation, nor programmed cell death pathways (at least not in the classical sense; see Hintzke et al., 2019). Instead, we found few moderately/highly expressed genes present in the immune system (*irak2* and 4, *tab1, tak1*, *mkk3/6*), cell cycle (cyclin A, B2, D2, *cdk4*), apoptotic (*cas2*, *cas8*, *birc8*), and autophagic (*becn1*, *atg2b-7-8-16*) pathways in the trophosome of *Riftia*. Interestingly, a previous study suggested that immune-related genes were significantly more expressed in the trophosome in relation to other symbiont-free tissues in the siboglinid *Ridgeia piscesae*, positing a more important role of the immune system in the host-endosymbiont homeostasis (Nyholm et al. 2012).

Few other individual components of the innate immune system, i.e., bactericidal permeability-increasing proteins and pattern recognition receptors, have been implicated in symbiont population control in tubeworms (Nyholm et al. 2012; Hinzke et al. 2019). However, based on our broad gene expression analyses, we argue that the host immune system does not play a major role in taming the endosymbiont population in the trophosome, as previously suggested (Hinzke et al. 2019). Furthermore, immunohistochemical and ultrastructural cell cycle analyses identified apoptotic and proliferative events in the trophosome (Pflugfelder et al. 2009), indicating that despite the overall low expression of gene markers related to these pathways described herein, these events occur in this tissue. In which extent these different pathways interact to shape the host/symbiont interactions and to maintain tissue homeostasis remains to be shown, however, it is clear that multiple and not mutually exclusive programmed cell-death, immune-related, and proliferative events (Supplementary note 7) are acting on the trophosome.

### From phenotype to genotype and back

After 40 years of intensive research, we are now finally able to integrate the obtained genome and tissue-specific transcriptome information with the current body of knowledge on the phenotype to better understand the genotype-phenotype interplay in the giant tubeworm. The *Riftia pachyptila* genome is characterized by reductive evolution with broad gene family contractions exceeding gene family expansions. Compared to the close relative *Lamelibrachia luymesi* (*Lamellibrachia* live at longer-lived and less physiologically-taxing hydrocarbon seeps), *Riftia* exhibits a more derived gene repertoire for important traits related to symbiosis and the highly disturbed and stressful hydrothermal vent habitat they inhabit in the deep sea.

The mutualism between *Riftia* and its symbiont has not transited from individuality of symbiotic partners to a new integrated organism (Szathmáry and Smith 1995) because it lacks mutual dependency (Kiers and West 2015; West et al. 2015). *Riftia* is, in fact, one of the few examples known in which dependency is asymmetric with a facultative horizontally transmitted symbionts, which have the capacity to live with or without the host. The *Riftia* host, however, is obliged to partner with the symbiont or else they cannot thrive. Therefore, the host’s fitness is strictly tied to the persistence of this association over ecological and evolutionary time scales. The genome data now clearly shows the peculiarities and divergencies in *Riftia’s* genotype compared to closely related free-living annelids and other lophotrochozoans, as well as which evolutionary adaptations of the host genotype ensure the maintenance of the association.

We found that despite the drastic morphological remodelling during its early development leading to the mouth-, gutless adult animal, *Riftia* retained the highly conserved developmental gene repertoire present in other lophotrochozoans and distant related animals. These results can be interpreted as counterintuitive considering that the adult body plan alone provides little unambiguous evidence of the vestimentiferan phylogenetic relationship. These animals were initially compared to deuterostomes (Caullery 1914) and considered related to hemichordates (Beklemishev 1944) as well as protostomes, but so unique that new phyla were erected to accommodate them (i.e., Pogonophora and Vestimentifera) (reviewed by (Rouse 2001; Pleijel et al. 2009)). The conservation of the developmental gene toolkit probably reflects the developmental constraints into the necessary to go step by step through deterministic stereotypic spiral cleavage and larval development (Nielsen 2004). Akin to other polychaetes, the endoderm is necessary not only later for feeding functions, also seen in the metatrochophore larvae prior symbiont infection (Nussbaumer et al. 2006), but also to develop most mesodermal tissue.

Combining the genomic information with tissue-specific transcriptomes allows us to hypothesize that the mesodermal trophosome (Nussbaumer et al. 2006; Bright et al. 2013) (Nussbaumer et al. 2006; Bright et al. 2013) is a multi-functional organ with ancestral inherited functions such as haematopoiesis. This trait, we hypothesize belongs to the functional repertoire known from mesodermal chloragogen (extravasal tissue surrounding the gut and blood vessels) derived from the visceral mesoderm in annelids like the trophosome in vestimentiferans (Nussbaumer et al. 2006; Bright et al. 2013). In fact, van der Land and Nørrevang suggested already in 1975, long before the symbionts were detected, that the trophosome in *Lammelibrachia luymesi* is the nutritive chloragogen tissue (Van der Land 1975). Although, overall knowledge is fragmentary it has been suggested that haematopoiesis in annelids is carried out by visceral as well as somatic mesoderm (Hartenstein 2006; Grigorian and Hartenstein 2013). In various polychaete species it was localized in particular in the (extravasal) chloragogen tissue, the (intravasal) heart body (Potswald 1969; Friedman and Weiss 1980; Braunbeck and Dales 1984; Fischer 1993) or the somatic peritoneum (Eckelbarger 1976). Our data unambiguously support the production of haemoglobin in the trophosome. Whether coelomocytes, known to be the immunocompetent cells of eucoelomates (Vetvicka and Sima 2009) including annelids (Dales 1964; Salzet et al. 2006; Cuvillier-Hot et al. 2014), and the haemocytes also develop from trophosomal tissue appears to be likely but remains to be verified.

The trophosome, however, further shows adaptations to new functions such as the well-known intracellular digestion through endosomal-like maturation of symbiosomes, as well as the processing of ammonia and storage of nitrogen waste analogous to the vertebrate liver. Most aquatic invertebrates, including annelids, are virtually ammoniotelic secreting ammonia (Larsen et al. 2011). Surprisingly, *Riftia* employs an additional ureotelic metabolism similar to terrestrial invertebrates and vertebrates converting toxic ammonia to urea/and or uric acid. Specifically, we found the entire set of genes for a complete urea cycle known to detoxify ammonia in the *Riftia* genome, with most of them upregulated in the trophosome. Therefore, we hypothesize that the trophosome share similar functions to the liver of vertebrates: Instead of secreting nitrogenous waste products through kidneys like in vertebrates or nephridia in annelids, the trophosome was found to store large amounts of uric acid and urea. Uric acid and urea can be utilized as a bioavailable source of N via the catabolic arm of the urea cycle yielding NH_4_ and CO_2_. Given the lack of urea transporters in the symbiont’s genome, and the presence of active ureases in the trophosome host tissue, this suggests that both the synthesis and breakdown of uric acid and urea is under host control.

What factor(s) might lead to the evolution of this physiological capacity to sequester and metabolize urea and uric acid? It has been shown in other symbioses that the exchange of bioavailable N between symbiotic partners plays an important role in recycling bioavailable N, such as in coral-dinoflagellate symbiosis that show an almost complete retention of bioavailable N (Tanaka et al. 2018). At many deep-sea vents, including those where *Riftia* thrive, bioavailable N is limited as ammonium and free amino acids are found in pM concentrations (Johnson et al. 1988). Moreover, *Riftia* are unable to ingest particulate matter so they cannot derive nitrogen from detritus. However, an abundant source of N is nitrate, which is found in deep seawater and can be reduced to ammonium by some microbes (Girguis et al. 2000). Previous studies (Hentschel et al. 1993; Girguis et al. 2000) found that *Riftia* take up nitrate from their environment, and the symbionts reduce nitrate to ammonium for symbiont and host growth and biosynthesis. However, the *Riftia* host’s ability to produce urea means that if can sequester bioavailable N that is only available to the host. At first glance, limiting symbiont access to N might be considered a way to control symbiont growth, as seen in cnidarian-Symbiodiniaceae (Xiang et al. 2020). This latter scenario, however, seems unlikely as there is ample bioavailable N (in the form of ammonium) throughout the trophosome in both freshly collected and experimentally tested worms (De Cian et al. 2000; Girguis et al. 2000). Rather, it seems plausible that *Riftia’*s production of urea allows the host to store and sequester N in a stable, largely nontoxic form. Whether urea is mobilized and provided to the host and symbionts during time of low N availability has yet to be experimentally tested, but this physiological capacity is another example of the remarkable adaptions found within that host, which allow it to modulate the rapid environmental changes found at vents and continue to provide for its own and the symbionts’ metabolic demands.

While the physiological and evolutionary aspects of tubeworm endosymbiosis have been sufficiently addressed over the past 40 years, the molecular mechanisms regulating host and symbiont interactions in siboglinids are still not fully understood. An immuno-centric view has been explored to explain the maintenance and regulation of the endosymbiont population in the giant tubeworm trophosome (Nyholm et al. 2012; Hinzke et al. 2019). Our results, contrary to the expectations, indicate that genes involved with the innate immune responses are downregulated in the trophosome (e.g., Toll-like receptor/MyD88) or in adult tubeworm tissues (e.g., sushi). These results suggest that the innate immune system plays a more prominent role into the establishment of the symbiosis during the infection in the larval stage, rather than preservation of the mutualism during the juvenile/adult life cycle. The control of the endosymbiont population in the trophosome is mainly achieved by the upregulation of endosomal and lysosomal hydrolases resulting in the active digestion of the endosymbionts (a “mowing” process as described by Hinzke et al. 2019).

The giant tubeworm genome establishes a unique and unprecedent hallmark bridging more than four decades of physiological research in *Riftia*, whilst it simultaneously provides new insights into the development, whole organism function and evolution of one of the most studied models for metazoan-symbiont interaction. We envisage that the resources generated herein foster many hypothesis-driven research pointing towards a more complete understanding of the genotype/phenotype interface in the *Riftia* and closely related taxa.

## Methods

A detailed methods section is available in Supplementary Material and methods and in Supplementary Figure 61. A brief overview of the bioinformatics pipeline follows below.

### Biological material and sequencing

*Riftia* genome DNA was obtained from a piece of vestimentum tissue belonging to single worm collected at the hydrothermal vent site Tica, East Pacific Rise (Alvin dive 4839, 9° 50.398 N, 104° 17.506 W, 2514 m depth, 2016) (Supplementary Figures 1, 2). PacBio libraries were generated with Sequel technology using the purified *Riftia* DNA. Tissue-specific transcriptomes were obtained from two specimens collected at Guaymas Basin, one female from the vent site Rebecca’s Roost (SuBastian dive 231, 27° 0.645 N, 111° 24.418 W, 2012 m depth, 2019) and one male from a vent site close to Big Pagoda (SuBastian dive 233, 27° 0.823 N, 111° 24.663 W, 2015 m depth, 2019) (Supplementary Figures 1, 2). The eight stranded paired-end tissue-specific transcriptomes (2×150 pb) were sequenced using Illumina NovaSeq SP technology.

### Genome assembly and processing

The five *Riftia* PacBio libraries were mapped against a custom database built with *Riftia* mitochondrial genome and its complete Endoriftia genome using minimap v2.17-r941 (Li 2018). Genome assembly was performed with canu v1.8 (Koren et al. 2017) with optimised parameters. Genome pre-processing, polishing, haplotig removal and contamination screening, was performed with arrow v2.3.3 (https://github.com/pacificbiosciences/genomicconsensus), purge_dups (https://github.com/dfguan/purge_dups) and blobtools v1.1.1 (Laetsch and Blaxter 2017), respectively. Mitochondrial and endosymbiont genome assemblies were executed with flye v2.5 (Kolmogorov et al. 2019). Annotation of the mitochondrial genome was performed with MITOS2 and GeSeq (Bernt et al. 2013; Tillich et al. 2017).

### Transcriptome assembly and processing

The removal of adapter sequences and quality filtering of reads from the raw transcriptome databases was performed with bbduk v38.42 (https://sourceforge.net/projects/bbmap/). *De novo* and genome-guided transcriptome assemblies were performed with transabyss v.2.0.1 and STAR v2.7.1a – Stringtie v2.0.6, respectively (Robertson et al. 2010; Dobin et al. 2013; Kovaka et al. 2019). Possible Endoriftia contamination was removed from the transcriptomes using blastn v2.8.1+ (Camacho et al. 2009). A global *de novo* transcriptome was generated with corset and Lace (https://github.com/Oshlack/Lace) (Davidson and Oshlack 2014).

### Gene prediction and annotation

The repeat landscape of *Riftia* genome was identified combining a custom giant tubeworm RepeatModeler v2.0 library followed by the masking of the repetitive elements with RepeatMasker v4.0.9 (A.F.A Smit, R. Hubley & P. Green, *RepeatMasker Open-4.0*). *Ab initio* gene prediction was performed with Augustus v3.3.3 aided with hint files (Stanke and Morgenstern 2005; Hoff and Stanke 2018). Only gene models with homology, orthology and gene expression evidence were kept. Protein annotation was performed with Interproscan v5.39-77.0, RNAscan-SE 2.0.5, signalP v5.0b and pfam_scan.pl (Jones et al. 2014; Lowe and Chan 2016; Almagro Armenteros et al. 2019).

### Identification of gene toolkits in *Riftia*

*Riftia* protein sequences were searched against well-curated catalog of developmental genes, amino and fatty acid biosynthesis, endocytosis-, apoptosis, autophagy- and immune-related genes using blastp v2.8.1+ and KEGG Automatic Annotation server (https://www.genome.jp/kegg/kaas/) (Moriya et al. 2007). Additionally, protein domain information was retrieved from pfam_scan.pl and Interpro results. Homology of the identified genes was confirmed through phylogenetic inferences using iqtree v1.6.11 combining ModelFinder, tree search, 1000 ultra-fast bootstrap and SH-aLRT test replicates (Nguyen et al. 2015; Kalyaanamoorthy et al. 2017; Hoang et al. 2018). The protein diagrams were drawn using IBS v.1.0.3 software (Liu et al. 2015), and the clustered heatmaps generated with the R package pheatmap (v1.0.12). Quantification of the gene expression levels was performed with kallisto v0.46.1 (Bray et al. 2016).

### Orthology, gene family analysis and positively selected genes

To assess *Riftia, Lamellibrachia* and Annelida lineage-specific genes orthology inferences using selected non-bilaterian, deuterostome, lophotrochozoan, ecdysozoans, representatives (N=36) were performed with Orthofinder v2.3.8 (Emms and Kelly 2019). To identify statistically significant gene family expansions/contractions in *Riftia* compared to other lophotrochozoans a second round of orthology was performed using 18 lophotrochozoan representatives and *Tribolium castaneum* as outgroup. Finally, to identify the gene family core within Annelida a last instance of orthofinder v2.3.8 was invoked using the *C. teleta, H. robusta, L. luymesi* and *R. pachyptila.* Only the longest isoform for each gene was used in the analysis. Non-synonymous (Ka) and synonymous (Ks) substitution rates were calculated with the stand-alone version of KaKs_calculator v.2 and HyPhy v. 2.5.15 (Pond et al. 2005; Wang et al. 2010; Z. Zhang et al. 2012). Only single-copy genes (1:1 orthologs) without any inconsistencies between the nucleotide and protein sequences were used in the analyses. Contracted and expanded gene families in the giant tubeworm genome were identified using CAFE v4.2.1 (De Bie et al. 2006; Han et al. 2013) using a calibrated starting tree produced by Phylobayes v4.1b (Lartillot et al. 2013). The contracted/expanded gene families were annotated with Interproscan v5.39-77.0 and the enrichment analysis for Gene Ontology was performed with topGO v2.36.0 using Fisher’s exact test against the *R. pachyptila* background (i.e., complete set of *Riftia* genes) coupled with weight01 algorithm. Rapidly evolving gene families in *Riftia* were annotated using PANTHER HMM scoring tool v2.2 with PANTHER_hmmscore database v15 (Mi et al. 2017). Protein domain contractions and expansions were found using iterative two-tailed Fisher’s exact (Supplementary File 2) test applied to pfam_scan.pl results. The obtained p-values were corrected using Benjamini and Hochberg method (Benjamini and Hochberg 1995) and only domains with a significant p-value of < 0.01 were further investigated.

### Haemoglobin evolution

The predicted *Riftia* haemoglobin (Hb) protein sequences were interrogated for the presence of the globin domain (PF00042) with hmmalign v3.1b2 (Mistry et al. 2013) and proteins without a hit were excluded from the analyses. Manual inspection and characterisation of the signature diagnostic residues/motifs in the haemoglobin chain and linker sequences were performed following previous works (Belato et al. 2019). Phylogenetic analyses were carried out as described in “Identification of gene toolkits in *Riftia*”. The resulting trees were midpoint rooted using Figtree (http://tree.bio.ed.ac.uk/software/figtree/). Additionally, to investigate the haemoglobin gene expression across different environmental conditions (sulphur rich, sulphur depleted and medium) we downloaded six publicly available trophosome transcriptomes from SRA (https://www.ncbi.nlm.nih.gov/sra) (accession numbers: SRR8949066 to SRR8949071). The transcriptome libraries were pre-processed as described in “Transcriptome assembly and processing”. *Riftia* Hb sequence was modelled using the Prime program implemented in the Schrödinger Drug Discovery (v2020.2) software suite. All illustrations of structures were made with PyMol v2.4 (https://pymol.org/2/).

### Comparative tissue-specific transcriptome

The *Riftia* transcriptome libraries were pseudoaligned against the merged filtered AUGUSTUS gene models with kallisto v.0.46.1 (Bray et al. 2016) to collect the gene expression data expressed as TPM counts (transcripts per million). Normalisation within and across tissues were independently performed before calculating the tissue specificity tau values (see https://rdrr.io/github/roonysgalbi/tispec/f/vignettes/). To mitigate possible sex-specific differences in the gene expression levels, tau calculations were performed using only the tubeworm female tissues. The absolutely tissue-specific genes (genes expressed only in a single tissue defined by a tau value of 1) were submitted to enrichment analyses for Gene Ontology with topGO as mentioned in “Orthology, gene family analysis and positively selected genes”.

## Data availability

The raw long and short reads used to generate the draft genome and tissue-specific transcriptomes, respectively, are available in the SRA database under the BioProject number PRJNA754493 (Supplementary Table 12).

## Acknowledgments

We thank Christian Baranyi (University of Vienna) and Jennifer Delaney (Harvard University) for the technical support on the RNA/DNA extraction and purification protocols, respectively. We also thank the Vienna BioCenter Core Facilities and the Genomics Center of the University of Minnesota for the generation of *Riftia*’s “omics” data. Andre Luiz de Oliveira (ALO) thanks Thales Kronenberger (Universität Klinikum Tübingen) for the help with the haemoglobin 3D homology modelling. ALO also thanks Andrew Calcino and Salvador Espada Hinojosa for their constructive conversations during the development of this work, and finally Bruna Yuri Pinheiro Imai for her continuous support to the first author. This work was funded by the Austrian FWF project P 31543 granted to M.B.

## Author contributions

ALO, MB and PG designed the project. ALO generated, implemented, and executed all the bioinformatic pipelines. All data analysis was performed by ALO with input from MB and PG, and JM. ALO wrote the first complete draft of this manuscript and all authors read, commented on, and approved the final version.

## Additional Information

### Supplementary tables

Supplementary Table 1 – *Riftia* genome and transcriptome pre-processing and annotation (.xlsx document). **A-B**, Overview of the *Riftia* PacBio libraries and genome statistics for the different pre-processed genome drafts. **C**, RepeatMasker results indicating the distribution of repeat elements in the *Riftia* genome. **D**, Overview of the eight tissue-specific *Riftia* libraries, trimming statistics and *de novo* assemblies. **E**, Mapping statistics of the individual tissue-specific transcriptomes using StringTie (full length transcripts x draft genome) and Bowtie2 (transcriptome library X *de novo* transcriptome) .**F**, Proteome prediction and BUSCO4 scores of the combined *de novo* and reference-based transcriptomes. The predicted proteomes were also mapped against the nr database.

Supplementary Table 2 – Databases used in the orthoFinder analysis (.xlsx document). **A**, Metazoan databases used in the orthoFinder analysis. **B**, Phyletic distribution of the orthogroups found in selected annelid genomes.

Supplementary Table 3 – Overview and quantification of transcription factor families in selected metazoans. Protein domain identification was performed with pfamscan. General overview of the transcription factors identified in annelids, molluscs, flatworms, phoronids, brachiopods and nemerteans.

Supplementary Table 4 – Gene family analysis with CAFE. **A**, Average expansions rates calculated by CAFE using lophotrochozoan orthogroups (N=18). Annelid values are highlighted in light red. **B-D**, Expanded, contracted and rapidly evolving gene families identified by CAFE in the giant tubeworm genome. Genes were annotated with Panther scoring tool. GO enrichment analyses using lineage-specific genes were performed with topGO in the three distinct groups. The enriched GO terms found in the three main ontologies are shown (Biological process, molecular function, and cellular component). In **D**, light purple and light red rows indicated expanded and contracted rapidly evolving orthogroups, respectively.

Supplementary Table 5 – Lineage-specific gene annotation and GO enrichment analyses. **A-C**, *Riftia*-, *Lamellibrachia*- and Siboglinidae-specific genes obtained through orthology analyses. Genes were annotatated with Panther scoring tool. GO enrichment analyses using lineage-specific genes were performed with topGO in the three distinct groups. The enriched GO terms found in the three main ontologies are shown (Biological process, molecular function, and cellular component).

Supplementary Table 6 – Contracted and expanded PFAM analysis in selected lophotrochozoans using two-tailed Fisher’s exact test with Bonferroni correction. **A-D**, PFAM domain quantification using four different sets of organisms. **A-** All lophotrochozoans and PFAM domains. **B –** All lophotrochozoans without transposase- and DUF-associated domains. **C** – All lophotrochozoans without transposase- and DUF-associated domains, except the siboglinids *Riftia* and *Lamellibrachia*. **D-** All lophotrochozoans without transposase- and DUF-associated domains, except deep-vent symbiotic animals. **E1-2**, Two-tailed Fisher’s exact test with Bonferroni correction using PFAM domains of *Riftia*/*Lamellibrachia,* and *Riftia*/average non-siboglinid lophotrochozoans pairs. **F,** Two-tailed Fisher’s exact test with Bonferroni correction using PFAM domains of *Lamellibrachia* and average non siboglinid lophotrochozoans. **G1-4**, Pairwise two-tailed Fisher’s exact test with Bonferroni correction between deep-sea symbiotic animals (*Riftia*, *Lamellibrachia*, *Bathymodiolus*, *Chrysomallon*) and the average non-deep-sea-symbiotic lophotrochozoans. **H**, Overlapping contracted/expanded domains found in symbiotic deep-sea symbiotic lophotrochozoans.

Supplementary Table 7 – Positively selected genes identified by HyPhy and KaKs calculator. **A-B**, Positively selected genes using two distinct methods. Gray rows correspond to positively selected genes identified in both methods. Annotation of positively selected genes was performed with Panther scoring tool.

Supplementary Table 8 – Gene expression quantification of selected proteins and protein families. **A-Y**, TPM (transcripts per million) values of selected proteins and protein families found in the eight different tissue-specific transcriptomes of *Riftia*. Colour scale (red – minimum; yellow – percentile 50; green -max) depicts the TPM values of genes found in the different *Riftia* tissues.

Supplementary Table 9 – Tau specific genes and GO enrichment analyses for the female tissue-specific transcriptomes (.xlsx document). Annotation of tau genes was performed with Panther scoring tool. GO enrichment analyses using tau specific genes were performed with topGO in the three distinct groups. The enriched GO terms found in the three main ontologies are shown (Biological process, molecular function, and cellular component).

Supplementary Table 10 – Distribution of important enzymes related to the biosynthesis of amino acids in *Riftia, Lamellibrachia* and their endosymbionts. For comparison other two annelids were included in the analysis (*Capitella*, *Hellobdela*). **A,** Distribution of key enzymes related to amino acids biosynthesis based on the KEGG pathways. Colour scale (red – minimum; yellow – percentile 50; green -max) depicts the number of genes found in the different annelid genomes. **B,** Gene identifier of key enzymes related to the biosynthesis of amino acids in the *Capitella*, *Helobdella*, *Riftia* and *Lamellibrachia* genomes. **C**, Distribution of Endoriftia genes found in the secretion system type II pathway, as presented in KEGG.

Supplementary Table 11 – Mitochondrial carrier proteins identified in the *Riftia* genome and their PANTHER and blastp annotations. Genes highlighted in grey are highly expressed in the trophosome tissue.

Supplementary Table 12 – SRA accession numbers for the genomic and transcriptomic data generated in this study.

### Supplementary files

Supplementary File 1: Supplementary information (.pdf).

Supplementary File 2: Rscript files, CAFE codes, *Riftiás* gene model created with Augustus and the giant tubeworm repeat database generated with RepeatModeler and RepeatMasker(.zip).

