## Supplementary File 1 for "Novel insights on obligate symbiont lifestyle and adaptation to chemosynthetic environment as revealed by the giant tubeworm genome"

André Luiz de Oliveira[1], Jessica Mitchell[2], Peter Girguis[2], Monika Bright[1]
[1] Department of Functional and Evolutionary Ecology, University of Vienna, Austria
[2] Department of Organismic and Evolutionary Biology, Harvard University, Cambridge, MA, USA

### Supplementary information

#### SUPPLEMENTARY FIGURES

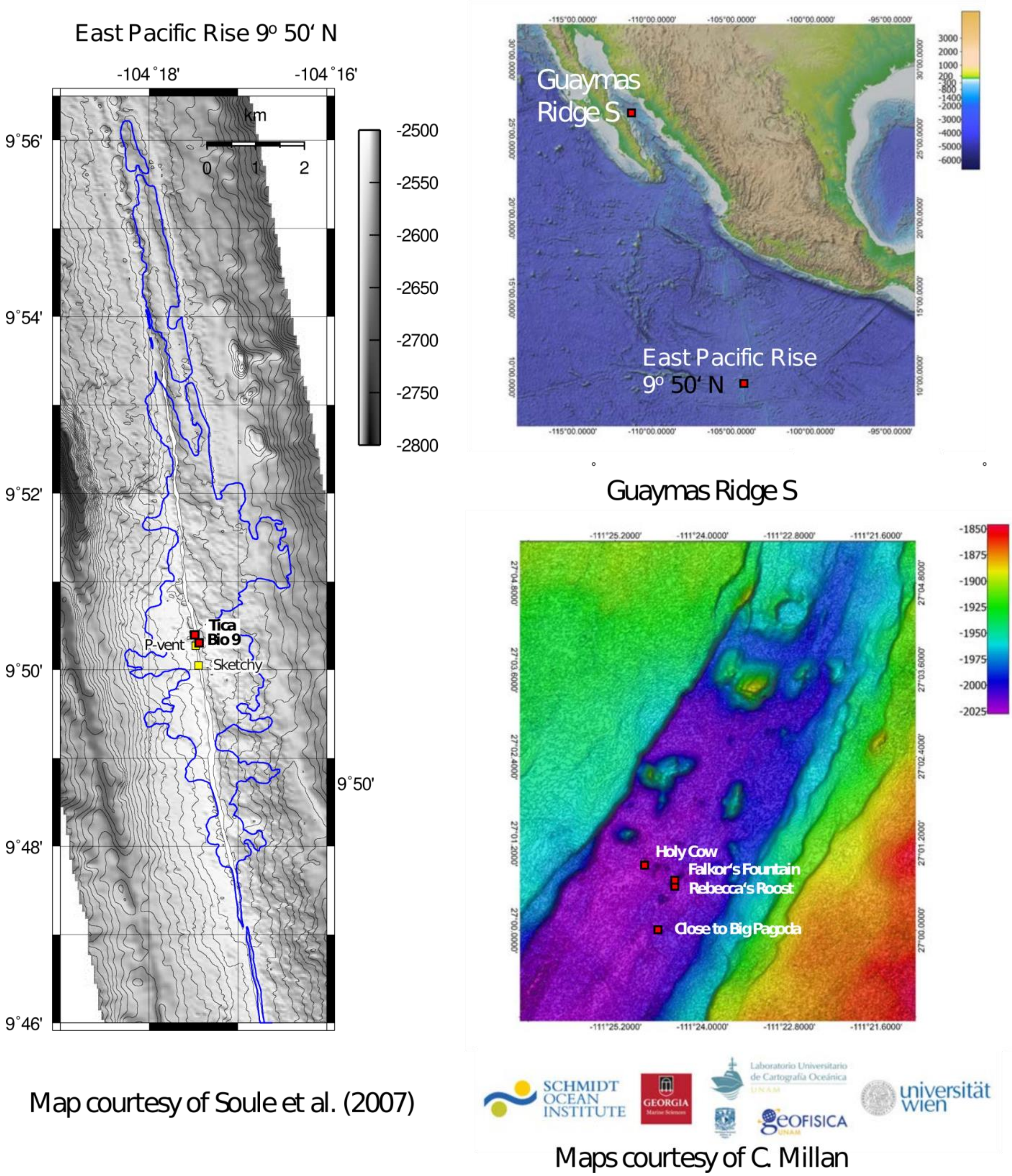

**Supplementary figure 1 | Maps of *Riftia* sampling sites.** The *Riftia* genome samples were collected from Tica vent site, located along the East Pacific Rise at ~9.50°N; *Riftia* transcriptomic samples came from vent sites Rebecca's Roost and Big Pagoda, Guaymas Basin.

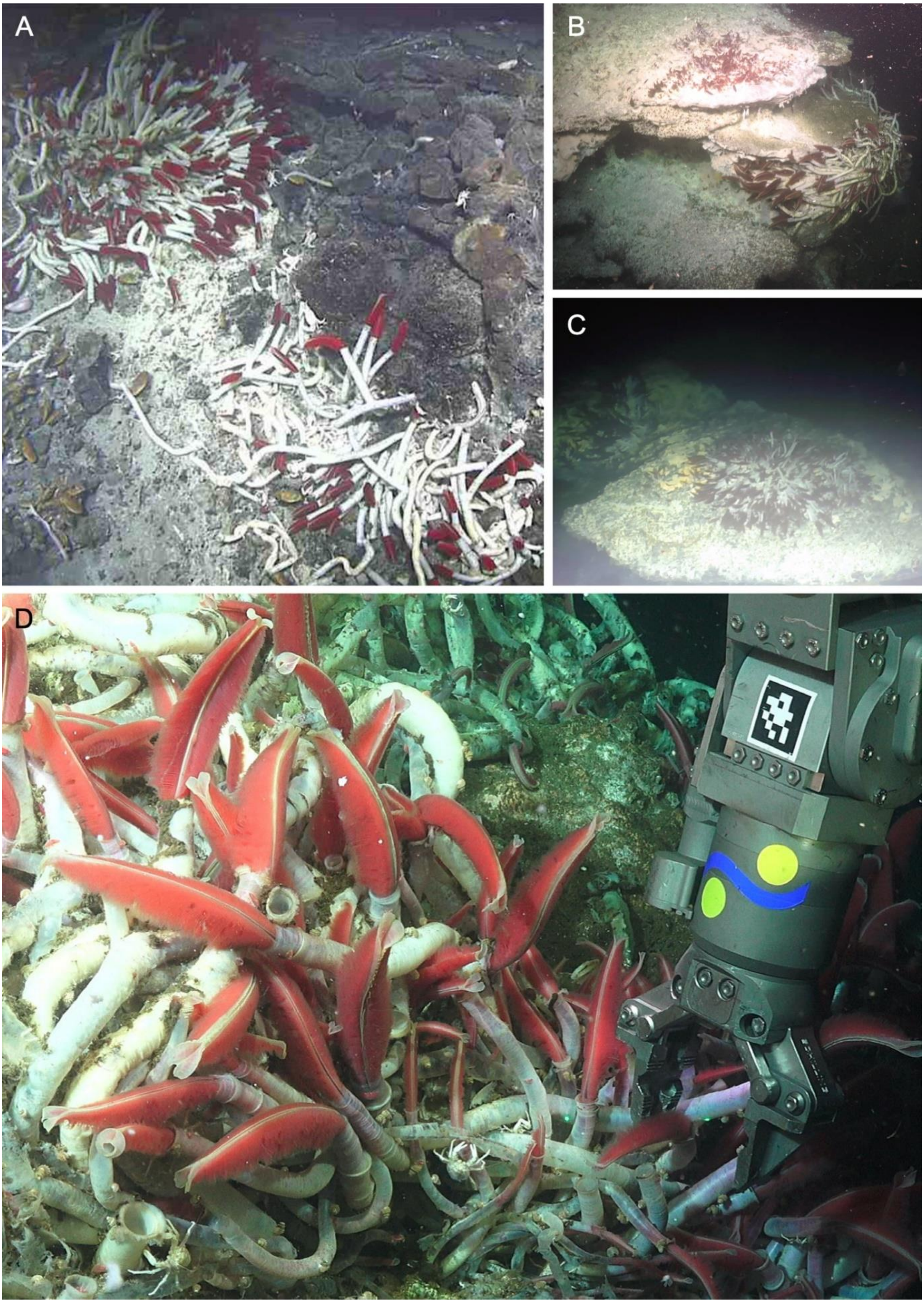

**Supplementary figure 2 | Vent sites. A** the Tica vent site, East Pacific Rise (*HOV Alvin* dive #4839, 2016). **B** Rebecca's Roost vent site, Guaymas Basin, 2019 (*ROV SuBastian* dive #231, 2019). **C** and **D** sites close to Big Pagoda, Guaymas Basin, 2019 (*ROV SuBastian* dive #233, 2019).

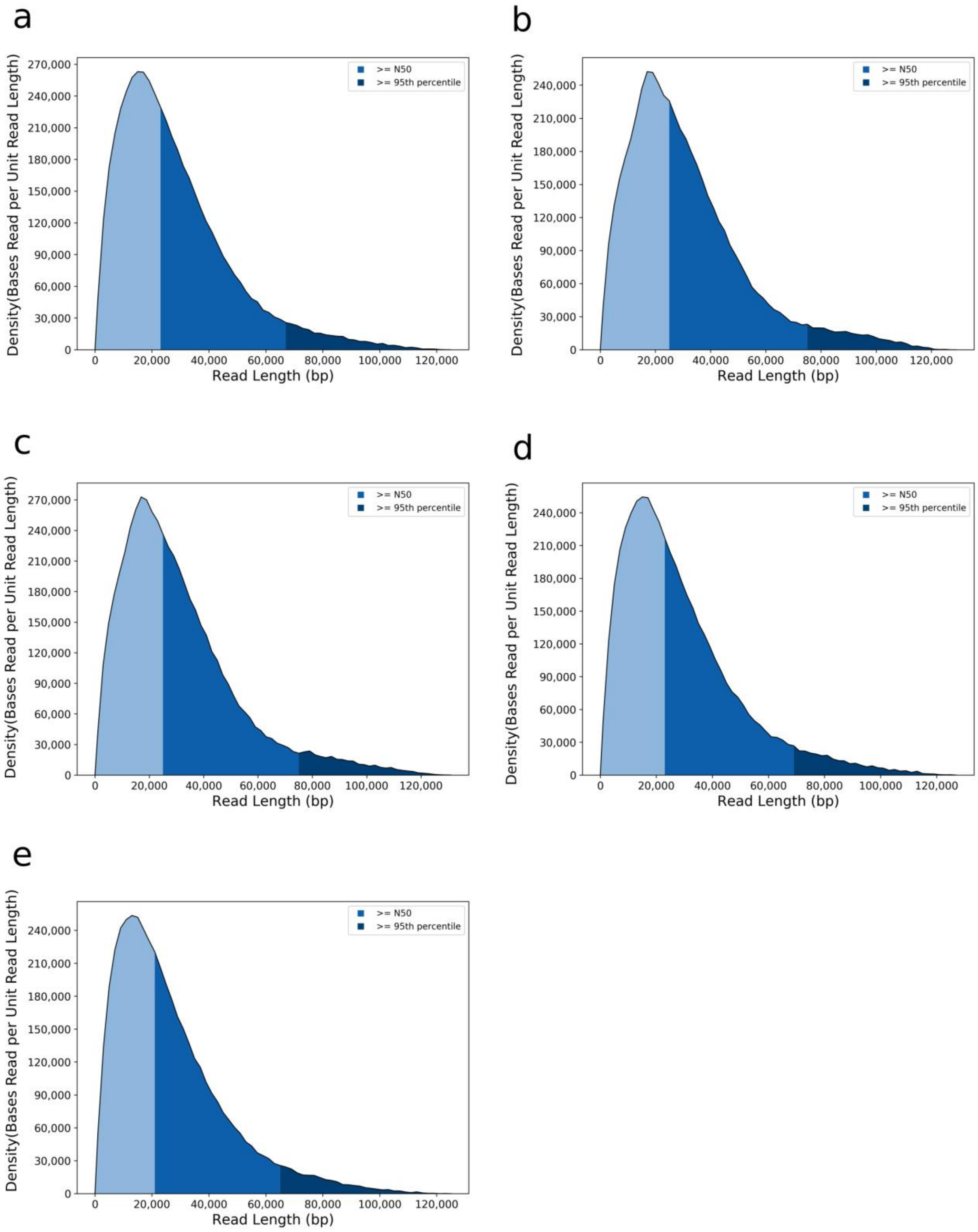

**Supplementary figure 3 | Long read sequencing of *Riftia* genome. a-e,** Pacific Biosciences™ Sequel sequencing performance of the five *Riftia* genomic libraries. The density (number of sequencing bases per read length) distribution is shown for each of the five libraries. Shades of blue indicate the number of sequenced bases present in reads smaller (light blue) and longer (blue, dark blue) than the N50 read value. The dark blue area corresponds the top 5% of the sequenced bases present in the longest read lengths. Note that half of the sequenced bases in all libraries are present in reads > 20kb.

A

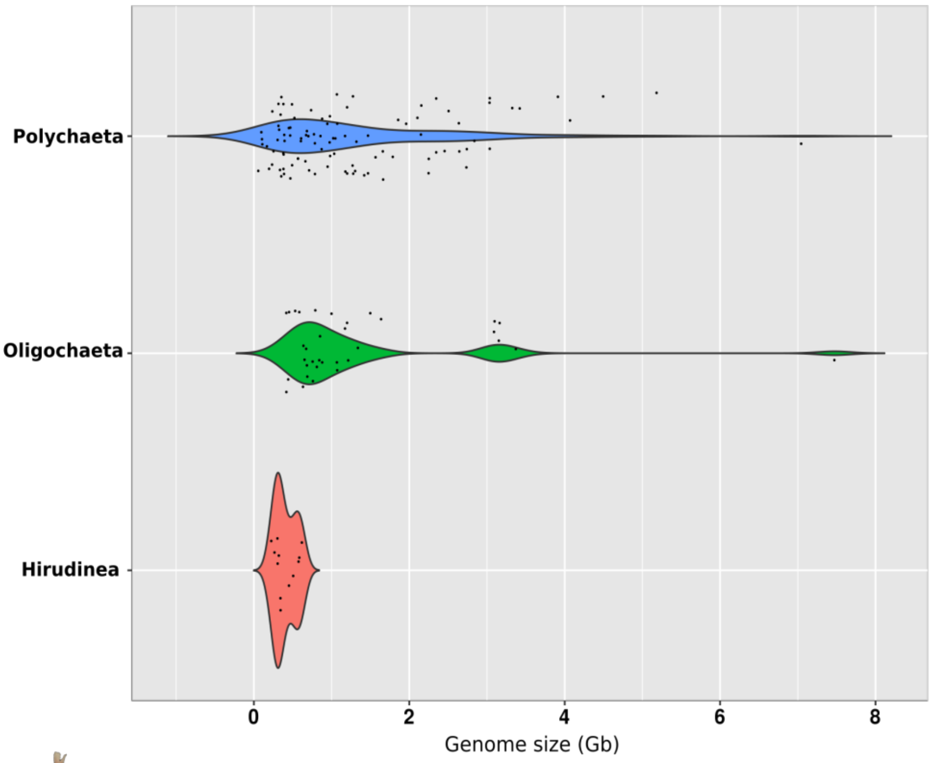

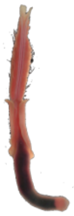

| Species | Genome size (Mb) | # of genes | Repeats | References |
|---|---|---|---|---|
| *R. pachyptila* | 772.62 | not available | not available | Bonnivard et al., 2009 |
| *R. pachyptila* | 625.92 | not available | not available | Dixon et al., 2001 |
| *R. pachyptila* | 560.78 | 25,983 | 29.99% | This study |

B

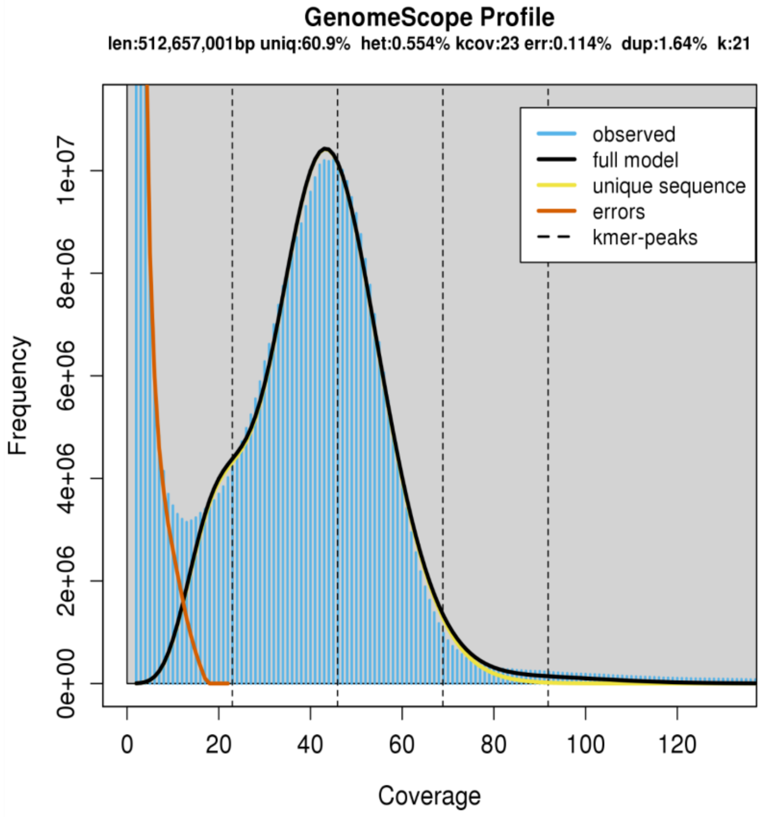

**Supplementary Figure 4 | *Riftia* genome size. A**, Violin plots of annelid genomes obtained from the Genome Size Database (http://www.genomesize.com/). Haploid DNA contents (C-values, in picograms) were converted to base pairs following Dolezel formula (Doležel et al., 2003). Kernel probability density values for the different annelid class-level taxa are shown (width of the plot). Individual data points, i.e., annelid genome sizes, are represented by black dots randomly scattered over the graph. The violin plots depict the range of the genome sizes (horizontal axis) and their frequencies (vertical axis) in the different annelid class-level taxa. B, GenomeScope estimations of genome heterozygosity, repeat content and size based on Canu (Koren et al. 2017) output results using a kmer-based statistical approach (http://qb.cshl.edu/genomescope/). The *Riftia* genome size estimation from the current study is markedly smaller than from the previously published studies.

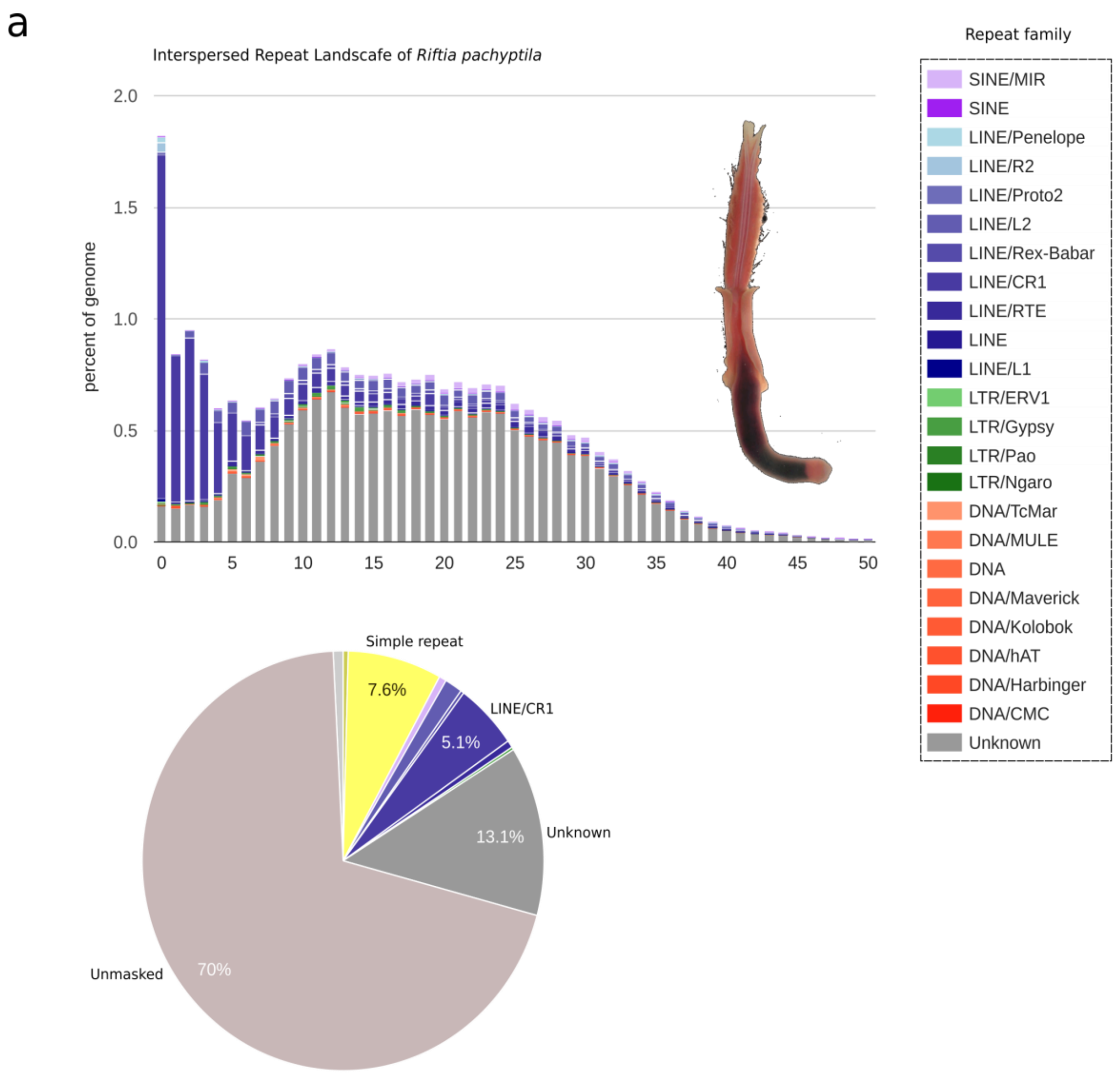

b

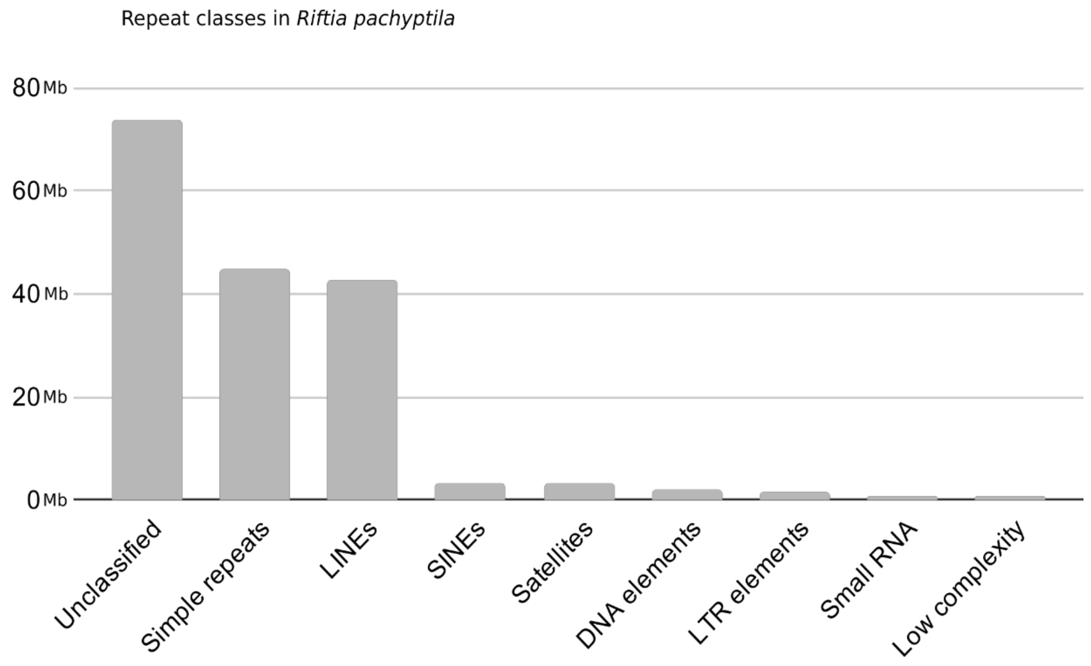

**Supplementary Figure 5 | *Riftia* interspersed repeat landscape. A,** The interspersed repeat landscape of *Riftia pachyptila* based on Kimura distances. The percentage of transposable elements in the tubeworm genome (y-axis) are clustered based on the Kimura values (x-axis). The pie chart shows the fraction of the *Riftia* genome covered by the different repeat families. Note that the repeat landscape is dominated by unclassified lineage-specific elements (i.e., unknown class). **B,** The number of repeat elements (in megabases) assigned to the main repeat classes in *Riftia.*

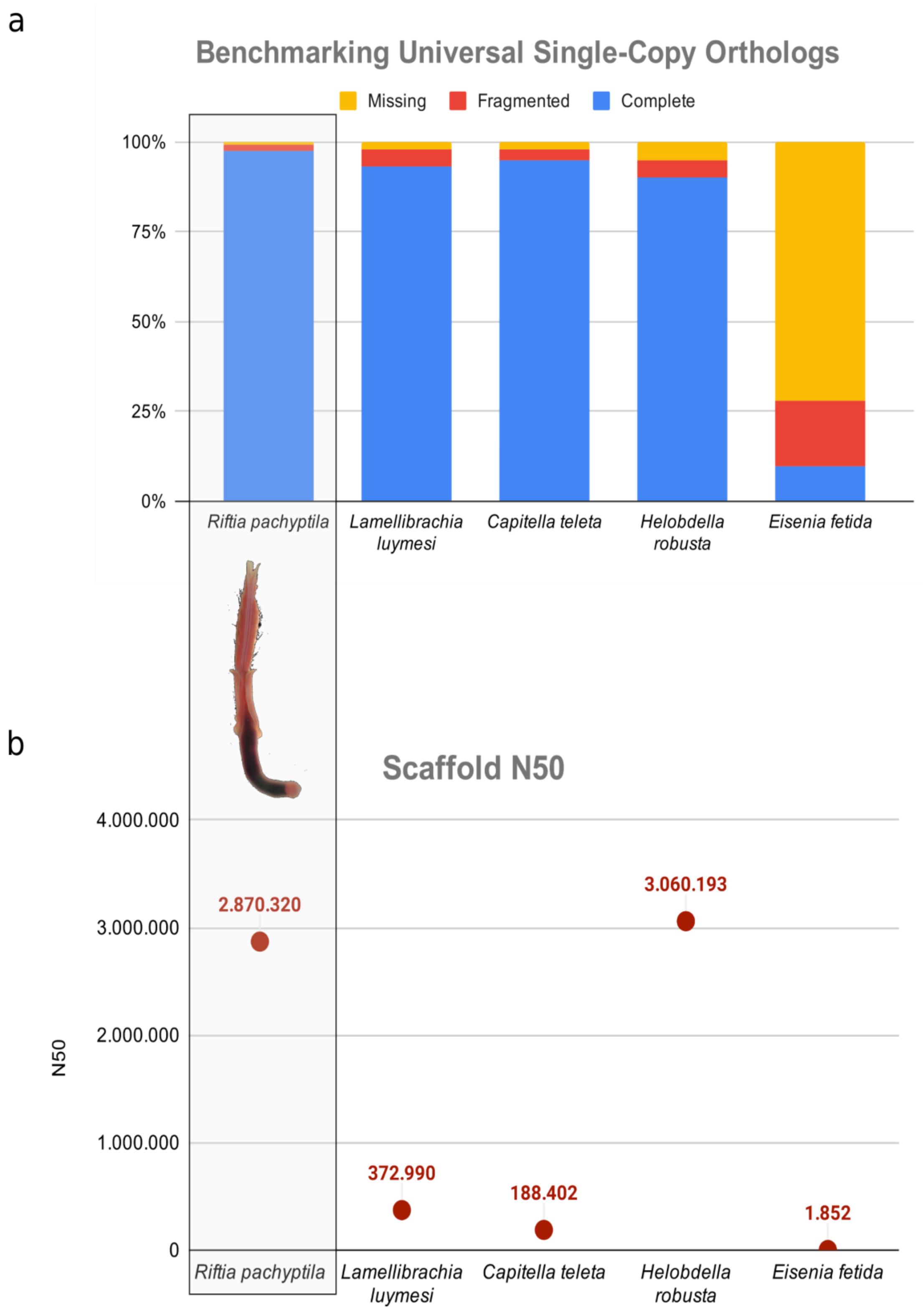

**Supplementary Figure 6 | Genome assembly assessment. A,** Metazoan BUSCO4 genome assembly scores of *Riftia pachyptila* and other four publicly available annelid genomes (Simakov et al. 2013; Paul et al. 2018; Li et al. 2019). **B,** scaffold $N_{50}$ values of *Riftia* and four other publicly available annelid genomes. The giant tubeworm genome presents the most complete genome up to date. The scaffold $N_{50}$ length of *Riftia* is similar to the leech *Helobdella robusta*.

a

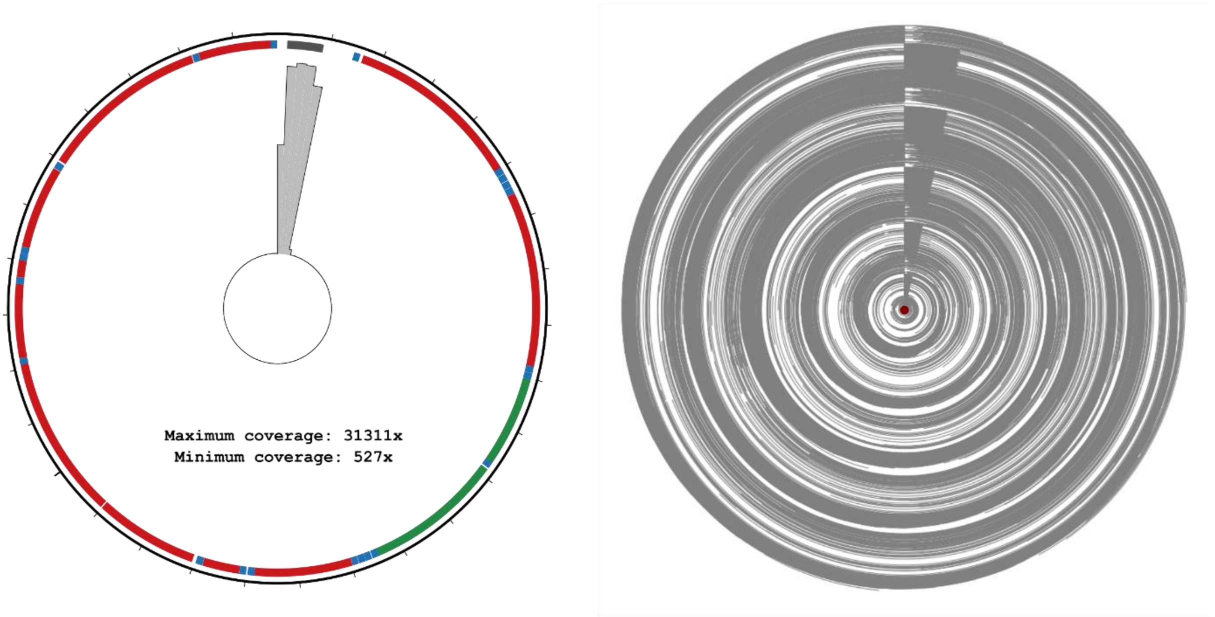

b

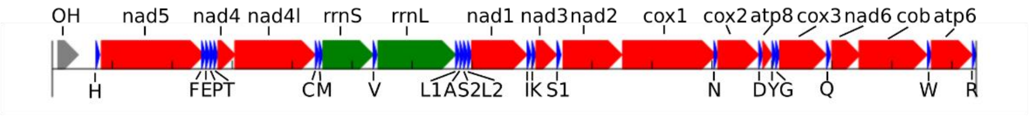

c

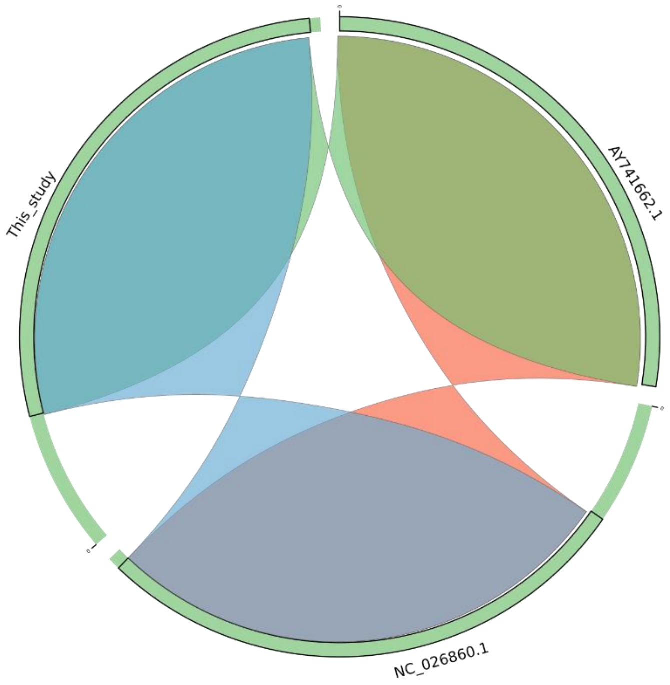

**Supplementary figure 7 | Mitochondrial genome. A**, Coverage plots of the PacBio long reads across the mitochondrial genome produced by circos and ConcatMap tools ([https://github.com/darylgohl/ConcatMap](https://github.com/darylgohl/ConcatMap)). A total of 43,009 long reads mapped against the closed *Riftia* mitogenome with coverage levels ranging from 31,311x (matching the repetitive D-loop region)

to 527x. **B**, Linear representation of the *R. pachyptila* mitochondrial genome showing the 13 coding sequencing regions, 2 rRNA and 22 tRNA genes. **C**, Large scale synteny blocks among the closed giant tubeworm mitochondrial genome provided in this study and other two publicly available references (AY741662 and NC_026860). The three mitochondrial reference genomes are virtually identical (>99% identity) with most of the mismatches located in the control region (D-loop). No rearrangements nor break of synteny were identified.

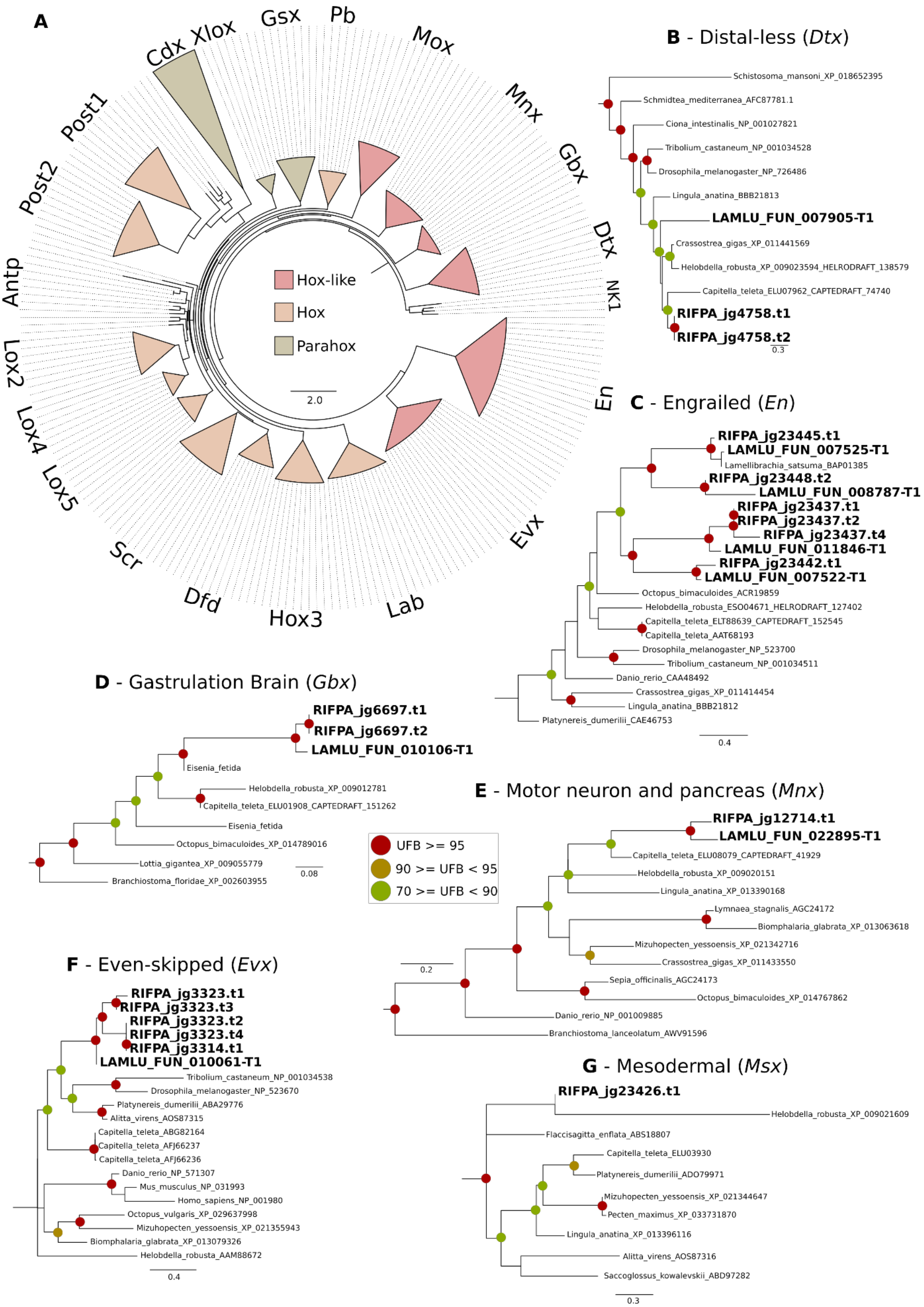

**Supplementary figure 8 | Phylogeny of Hox, ParaHox and Hox-like genes**. **A,** Maximum-likelihood phylogenetic tree inference of the homebox genes belonging to Hox, ParaHox and Hox-like families. The different ANTP genes were collapsed for an easier visualization. The different colours represent Hox, ParaHox and Hox-like gene members. NK1 genes were used as outgroup. **B-G,** Expanded tree inferences of Hox-like ANTP genes. The branch support values are represented by the coloured circles in the tree nodes. Red circles represent ultrafast bootstrap values >= 95. Yellow circles represent ultrafast bootstrap values >= 90 and < 95. Green circles represent ultrafast bootstrap values < 90 and >= 70. Ultrafast bootstrap values smaller than 70 are not shown. Accession numbers for NCBI database are displayed after the species names.

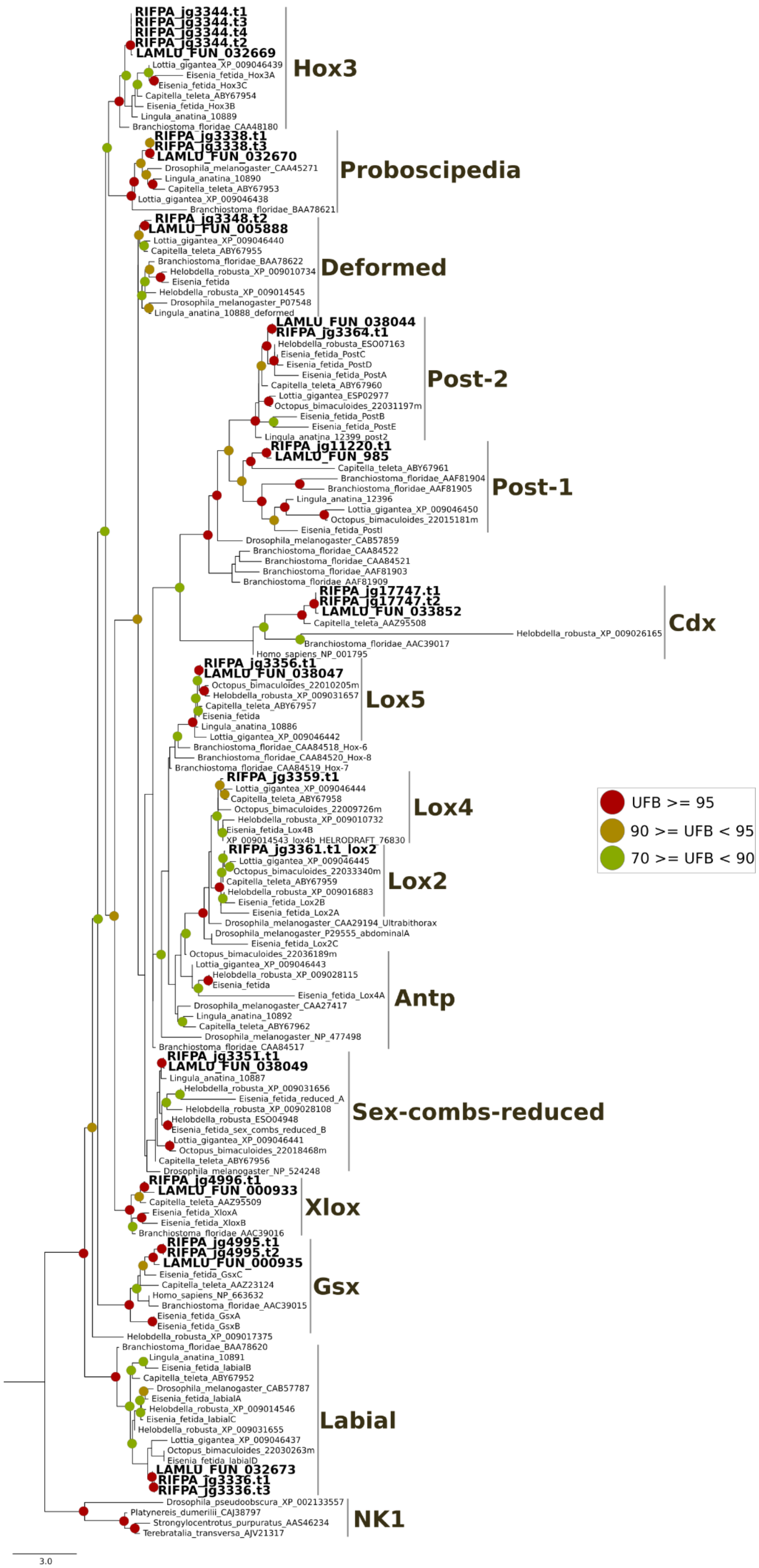

RIFPA_jg3344.t1
RIFPA_jg3344.t3
RIFPA_jg3344.t4
RIFPA_jg3344.t2
LAMLU_FUN_032669
Lottia_gigantea_XP_009046439
Eisenia_fetida_Hox3A
Eisenia_fetida_Hox3C
Capitella_teleta_ABY67954
Eisenia_fetida_Hox3B
Lingula_anatina_10889
Branchiostoma_floridae_CAA48180
Hox3
RIFPA_jg3338.t1
RIFPA_jg3338.t3
LAMLU_FUN_032670
Drosophila_melanogaster_CAA45271
Lingula_anatina_10890
Capitella_teleta_ABY67953
Lottia_gigantea_XP_009046438
Branchiostoma_floridae_BAA78621
Proboscipedia
RIFPA_jg3348.t2
LAMLU_FUN_005888
Lottia_gigantea_XP_009046440
Capitella_teleta_ABY67955
Branchiostoma_floridae_BAA78622
Helobdella_robusta_XP_009010734
Eisenia_fetida
Helobdella_robusta_XP_009014545
Drosophila_melanogaster_P07548
Lingula_anatina_10888_deformed
Deformed
LAMLU_FUN_038044
RIFPA_jg3364.t1
Helobdella_robusta_ESO07163
Eisenia_fetida_PostC
Eisenia_fetida_PostD
Eisenia_fetida_PostA
Capitella_teleta_ABY67960
Lottia_gigantea_ESP02977
Octopus_bimaculoides_22031197m
Eisenia_fetida_PostB
Eisenia_fetida_PostE
Lingula_anatina_12399_post2
Post-2
RIFPA_jg11220.t1
LAMLU_FUN_985
Capitella_teleta_ABY67961
Branchiostoma_floridae_AAF81904
Branchiostoma_floridae_AAF81905
Lingula_anatina_12396
Lottia_gigantea_XP_009046450
Octopus_bimaculoides_22015181m
Eisenia_fetida_PostI
Post-1
Drosophila_melanogaster_CAB57859
Branchiostoma_floridae_CAA84522
Branchiostoma_floridae_CAA84521
Branchiostoma_floridae_AAF81903
Branchiostoma_floridae_AAF81909
RIFPA_jg17747.t1
RIFPA_jg17747.t2
LAMLU_FUN_033852
Capitella_teleta_AAZ95508
Helobdella_robusta_XP_009026165
Branchiostoma_floridae_AAC39017
Homo_sapiens_NP_001795
Cdx
RIFPA_jg3356.t1
LAMLU_FUN_038047
Octopus_bimaculoides_22010205m
Helobdella_robusta_XP_009031657
Capitella_teleta_ABY67957
Eisenia_fetida
Lingula_anatina_10886
Lottia_gigantea_XP_009046442
Branchiostoma_floridae_CAA84518_Hox-6
Branchiostoma_floridae_CAA84520_Hox-8
Branchiostoma_floridae_CAA84519_Hox-7
Lox5
RIFPA_jg3359.t1
Lottia_gigantea_XP_009046444
Capitella_teleta_ABY67958
Octopus_bimaculoides_22009726m
Helobdella_robusta_XP_009010732
Eisenia_fetida_Lox4B
XP_009014543_lox4b_HELRODRAFT_76830
Lox4
RIFPA_jg3361.t1_lox2
Lottia_gigantea_XP_009046445
Octopus_bimaculoides_22033340m
Capitella_teleta_ABY67959
Helobdella_robusta_XP_009016883
Eisenia_fetida_Lox2B
Eisenia_fetida_Lox2A
Lox2
Drosophila_melanogaster_CAA29194_Ultrabithorax
Drosophila_melanogaster_P29555_abdominalA
Eisenia_fetida_Lox2C
Octopus_bimaculoides_22036189m
Lottia_gigantea_XP_009046443
Helobdella_robusta_XP_009028115
Eisenia_fetida
Eisenia_fetida_Lox4A
Drosophila_melanogaster_CAA27417
Lingula_anatina_10892
Capitella_teleta_ABY67962
Drosophila_melanogaster_NP_477498
Branchiostoma_floridae_CAA84517
Antp
RIFPA_jg3351.t1
LAMLU_FUN_038049
Lingula_anatina_10887
Helobdella_robusta_XP_009031656
Eisenia_fetida_reduced_A
Helobdella_robusta_XP_009028108
Helobdella_robusta_ESO04948
Eisenia_fetida_sex_combs_reduced_B
Lottia_gigantea_XP_009046441
Octopus_bimaculoides_22018468m
Capitella_teleta_ABY67956
Drosophila_melanogaster_NP_524248
Sex-combs-reduced
RIFPA_jg4996.t1
LAMLU_FUN_000933
Capitella_teleta_AAZ95509
Eisenia_fetida_XloxA
Eisenia_fetida_XloxB
Branchiostoma_floridae_AAC39016
Xlox
RIFPA_jg4995.t1
RIFPA_jg4995.t2
LAMLU_FUN_000935
Eisenia_fetida_GsxC
Capitella_teleta_AAZ23124
Homo_sapiens_NP_663632
Branchiostoma_floridae_AAC39015
Eisenia_fetida_GsxA
Eisenia_fetida_GsxB
Gsx
Helobdella_robusta_XP_009017375
Branchiostoma_floridae_BAA78620
Lingula_anatina_10891
Eisenia_fetida_labialB
Capitella_teleta_ABY67952
Drosophila_melanogaster_CAB57787
Eisenia_fetida_labialA
Helobdella_robusta_XP_009014546
Eisenia_fetida_labialC
Helobdella_robusta_XP_009031655
Lottia_gigantea_XP_009046437
Octopus_bimaculoides_22030263m
Eisenia_fetida_labialD
LAMLU_FUN_032673
RIFPA_jg3336.t1
RIFPA_jg3336.t3
Labial
Drosophila_pseudoobscura_XP_002133557
Platynereis_dumerilii_CAJ38797
Strongylocentrotus_purpuratus_AAS46234
Terebratalia_transversa_AJV21317
NK1
3.0
UFB >= 95
90 >= UFB < 95
70 >= UFB < 90

**Supplementary figure 9 | Phylogeny of Hox and ParaHox genes**. Maximum-likelihood phylogenetic tree inference of the Hox and ParaHox genes using 1000 ultrafast bootstrap replicates. The branch support values are represented by the coloured circles in the tree nodes. Red circles represent ultrafast bootstrap values >= 95. Yellow circles represent ultrafast bootstrap values >= 90 and < 95. Green circles represent ultrafast bootstrap values < 90 and >= 70. Ultrafast bootstrap values smaller than 70 are not shown. *Riftia pachyptila* contains 10 out of the 11 lophotrochozoan Hox genes (*Hox7* is missing) and all the three ParaHox (*Cdx*, *Xlox* and *Gsx*). NK1 genes were used as outgroup. Accession numbers for NCBI database are displayed after the species names.

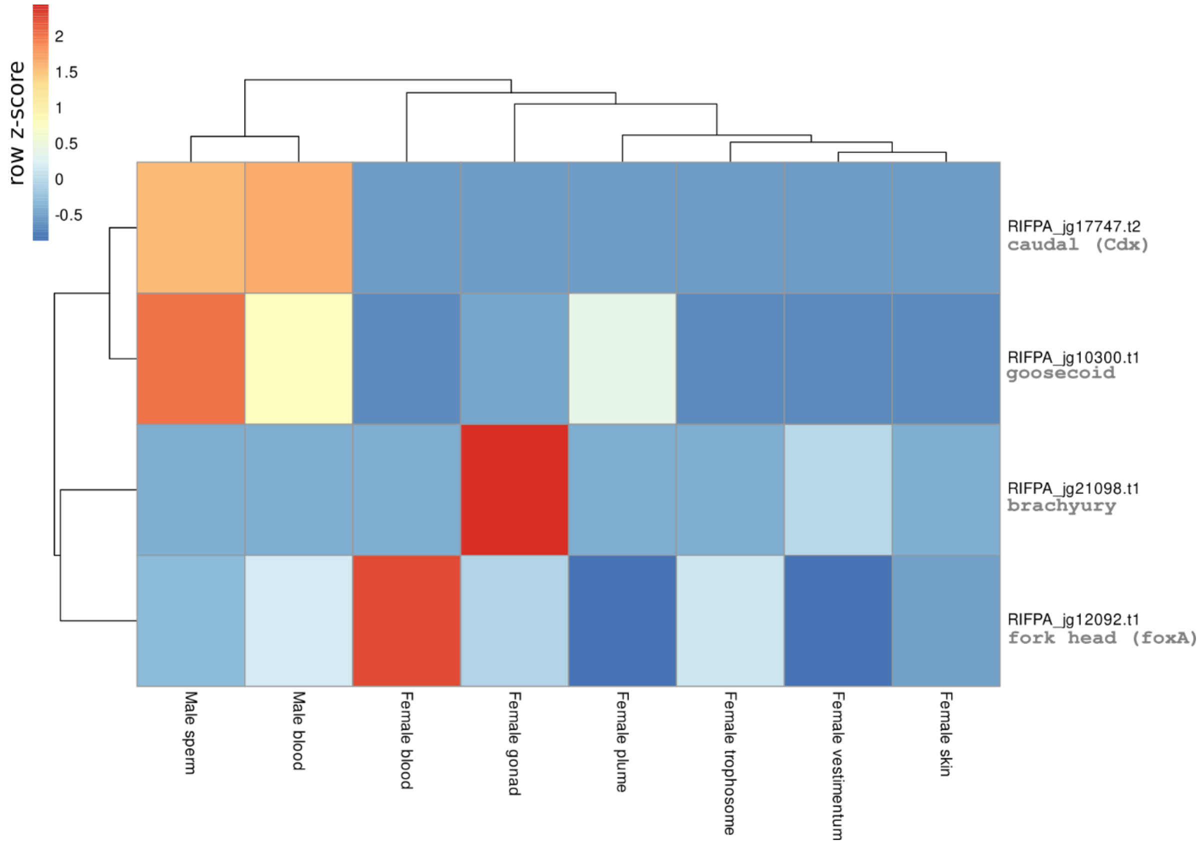

**Supplementary figure 10 | Expression of selected genes involved in the development of the digestive tract.** Expression profile of select genes involved in the development of the digestive tract in *Riftia pachyptila*. Colour coding reflects the expression patterns based on row Z-score calculations. The Parahox genes *Xlox* and *Gsx*, responsible for the patterning of the digestive tract in metazoans, did not show any expression on the eight adult *Riftia* tissues. All developmental genes involved in the digestive tract patterning are lowly expressed in the adult tissues.

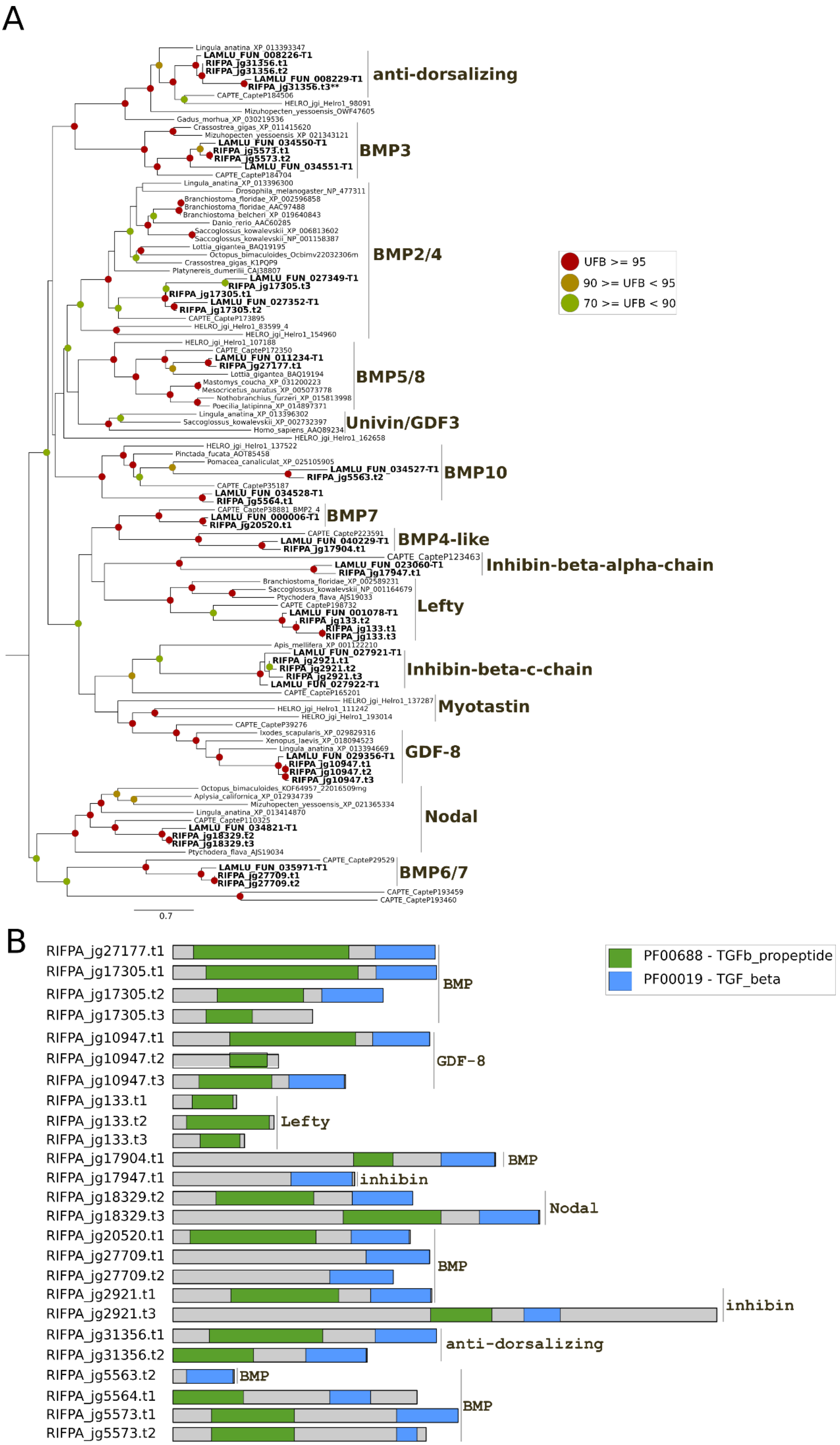
A
anti-dorsalizing
BMP3
BMP2/4
BMP5/8
Univin/GDF3
BMP10
BMP7
BMP4-like
Inhibin-beta-alpha-chain
Lefty
Inhibin-beta-c-chain
Myotastin
GDF-8
Nodal
BMP6/7
UFB >= 95
90 >= UFB < 95
70 >= UFB < 90
0.7
B
PF00688 - TGFb_propeptide
PF00019 - TGF_beta
RIFPA_jg27177.t1
RIFPA_jg17305.t1
RIFPA_jg17305.t2
RIFPA_jg17305.t3
BMP
RIFPA_jg10947.t1
RIFPA_jg10947.t2
RIFPA_jg10947.t3
GDF-8
RIFPA_jg133.t1
RIFPA_jg133.t2
RIFPA_jg133.t3
Lefty
RIFPA_jg17904.t1
BMP
RIFPA_jg17947.t1
inhibin
RIFPA_jg18329.t2
RIFPA_jg18329.t3
Nodal
RIFPA_jg20520.t1
RIFPA_jg27709.t1
RIFPA_jg27709.t2
BMP
RIFPA_jg2921.t1
RIFPA_jg2921.t3
inhibin
RIFPA_jg31356.t1
RIFPA_jg31356.t2
anti-dorsalizing
RIFPA_jg5563.t2
BMP
RIFPA_jg5564.t1
RIFPA_jg5573.t1
RIFPA_jg5573.t2
BMP

**Supplementary figure 11 | Phylogeny of TGF-beta genes. A,** Mid-rooted maximum-likelihood phylogenetic tree inference of the TGF-beta genes using 1000 ultrafast bootstrap replicates. The branch support values are represented by the coloured circles in the tree nodes. Red circles represent ultrafast bootstrap values >= 95. Yellow circles represent ultrafast bootstrap values >= 90 and < 95. Green circles represent ultrafast bootstrap values < 90 and >= 70. Ultrafast bootstrap values smaller than 70 are not shown. **B,** Domain composition of TGF-beta ligands based on PFAM database. Note that *lefty* gene does not contain the typical TGF_beta domain characteristic of the TGF-beta family.** no TGFb_propeptide and TGF_beta protein domains could be identified in this protein. Accession numbers for NCBI database are displayed after the species names. *Capitella, Helobdella* and *Lamellibrachia* gene identification are derived from the publicly available annotated genomes.

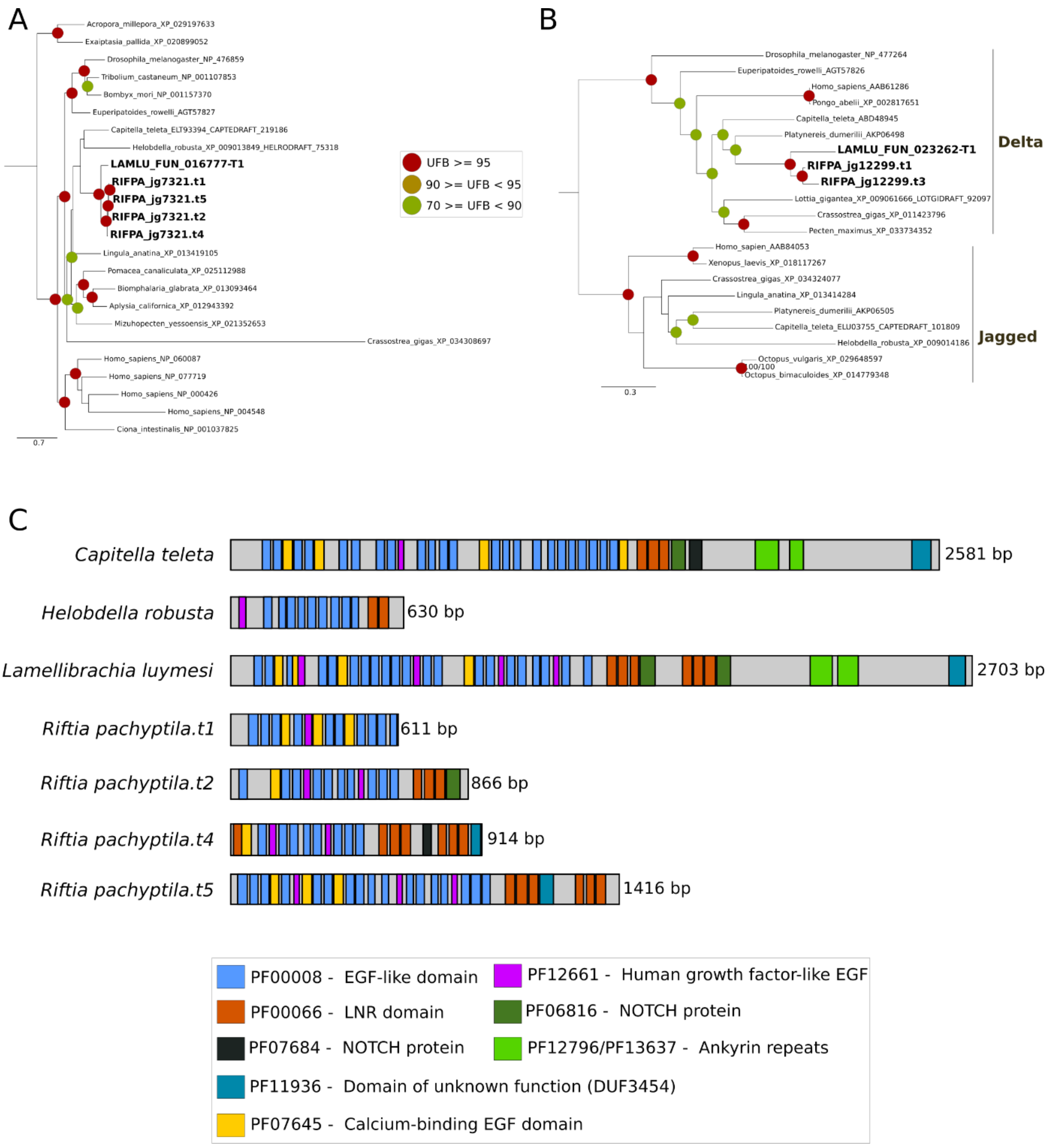

**Supplementary figure 12 | Phylogeny of Notch ligand and receptor genes. A-B,** Mid-rooted maximum-likelihood phylogenetic tree inference of the Notch ligand and receptor genes using 1000

ultrafast bootstrap replicates. The branch support values are represented by the coloured circles in the tree nodes. Red circles represent ultrafast bootstrap values >= 95. Yellow circles represent ultrafast bootstrap values >= 90 and < 95. Green circles represent ultrafast bootstrap values < 90 and >= 70. Ultrafast bootstrap values smaller than 70 are not shown. **B,** Domain composition of Notch ligands based on PFAM database. Accession numbers for NCBI database are displayed after the species names. *Capitella, Helobdella* and *Lamellibrachia* gene identification are derived from the publicly available annotated genomes.

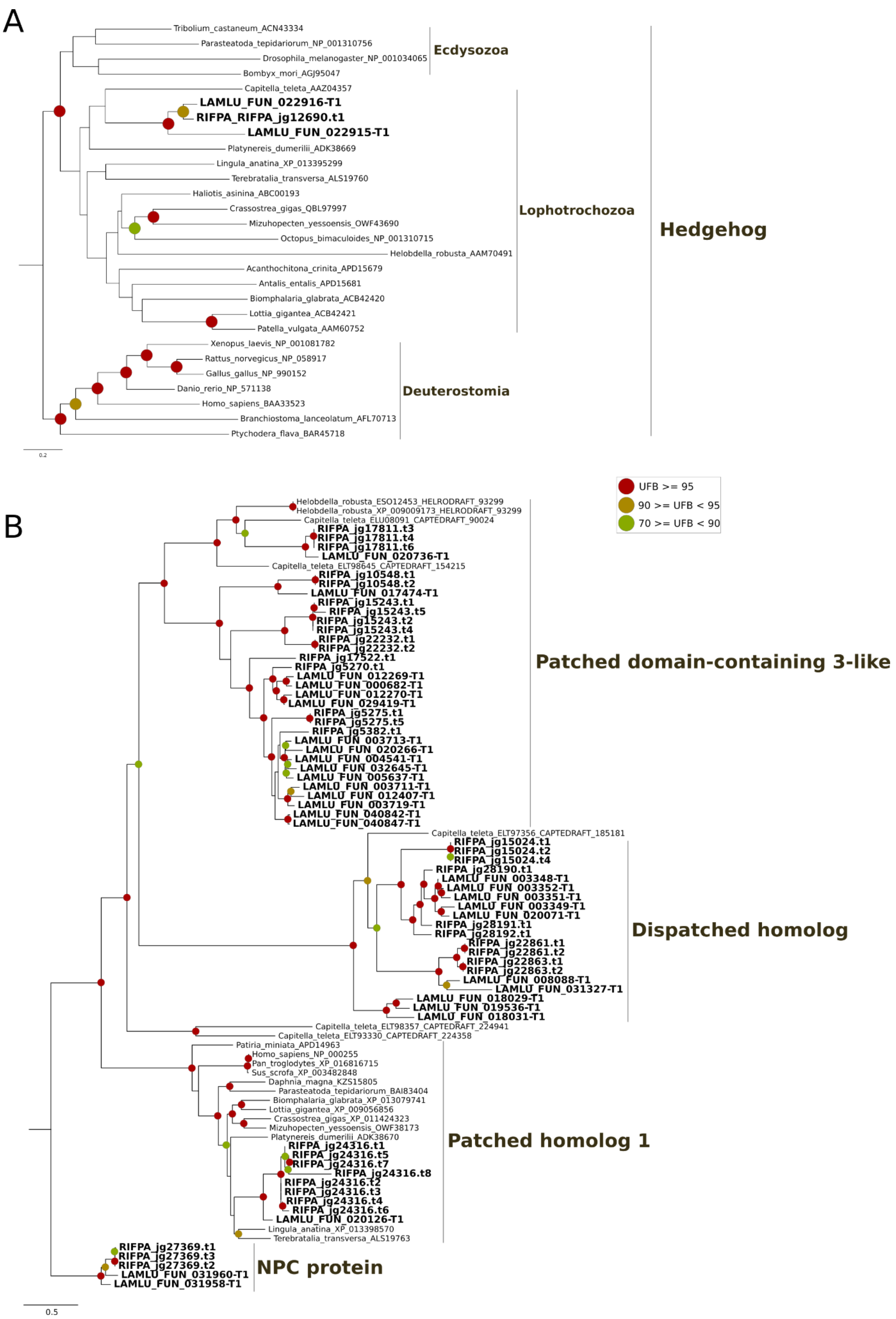

**Supplementary figure 13 | Phylogeny of Hedgehog ligand and receptor genes. A-B,** Maximum-likelihood phylogenetic tree inference of the Hedgehog ligand and receptor genes (*patched* and *dispatched*) using 1000 ultrafast bootstrap replicates. Dispatched receptors are expanded in the two tubeworm genomes. The branch support values are represented by the coloured circles in the tree nodes. Red circles represent ultrafast bootstrap values >= 95. Yellow circles represent ultrafast bootstrap values >= 90 and < 95. Green circles represent ultrafast bootstrap values < 90 and >= 70. Ultrafast bootstrap values smaller than 70 are not shown. Tubeworm Niemann-Pick C (NPC) proteins were used as outgroup. Accession numbers for NCBI database are displayed after the species names. *Capitella, Helobdella* and *Lamellibrachia* gene identification are derived from the publicly available annotated genomes.

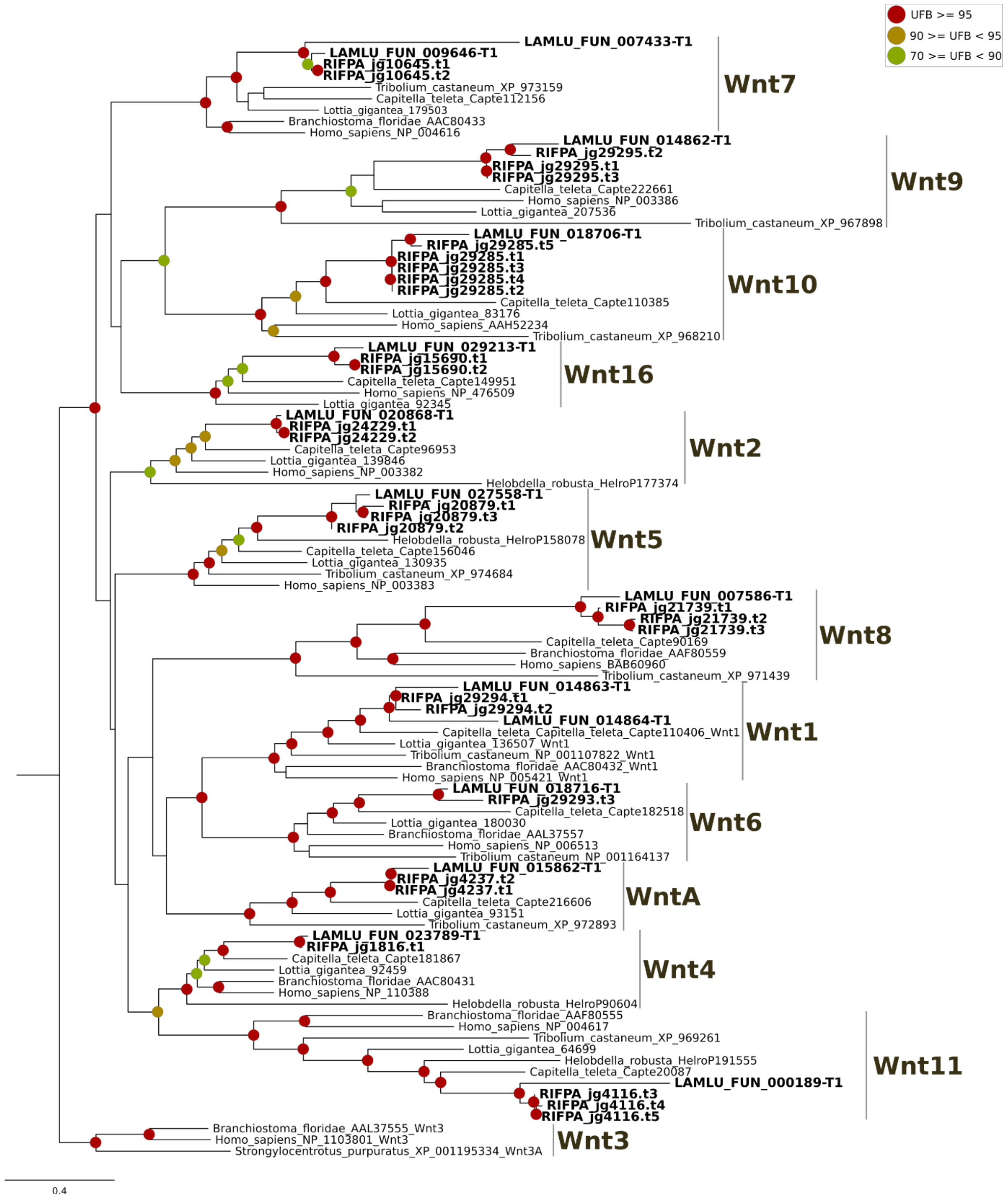

**Supplementary figure 14 | Phylogeny of Wnt ligands**. Maximum-likelihood phylogenetic tree inference of the Wnt ligand genes using 1000 ultrafast bootstrap replicates. *Riftia* contains all expected Wnt genes. The branch support values are represented by the coloured circles in the tree nodes. Red circles represent ultrafast bootstrap values >= 95. Yellow circles represent ultrafast bootstrap values >= 90 and < 95. Green circles represent ultrafast bootstrap values < 90 and >= 70. Ultrafast bootstrap values smaller than 70 are not shown. Deuterostome *Wnt3* genes were used as outgroup. Accession numbers for NCBI database are displayed after the species names. *Capitella, Helobdella* and *Lamellibrachia* gene identification are derived from the publicly available annotated genomes.

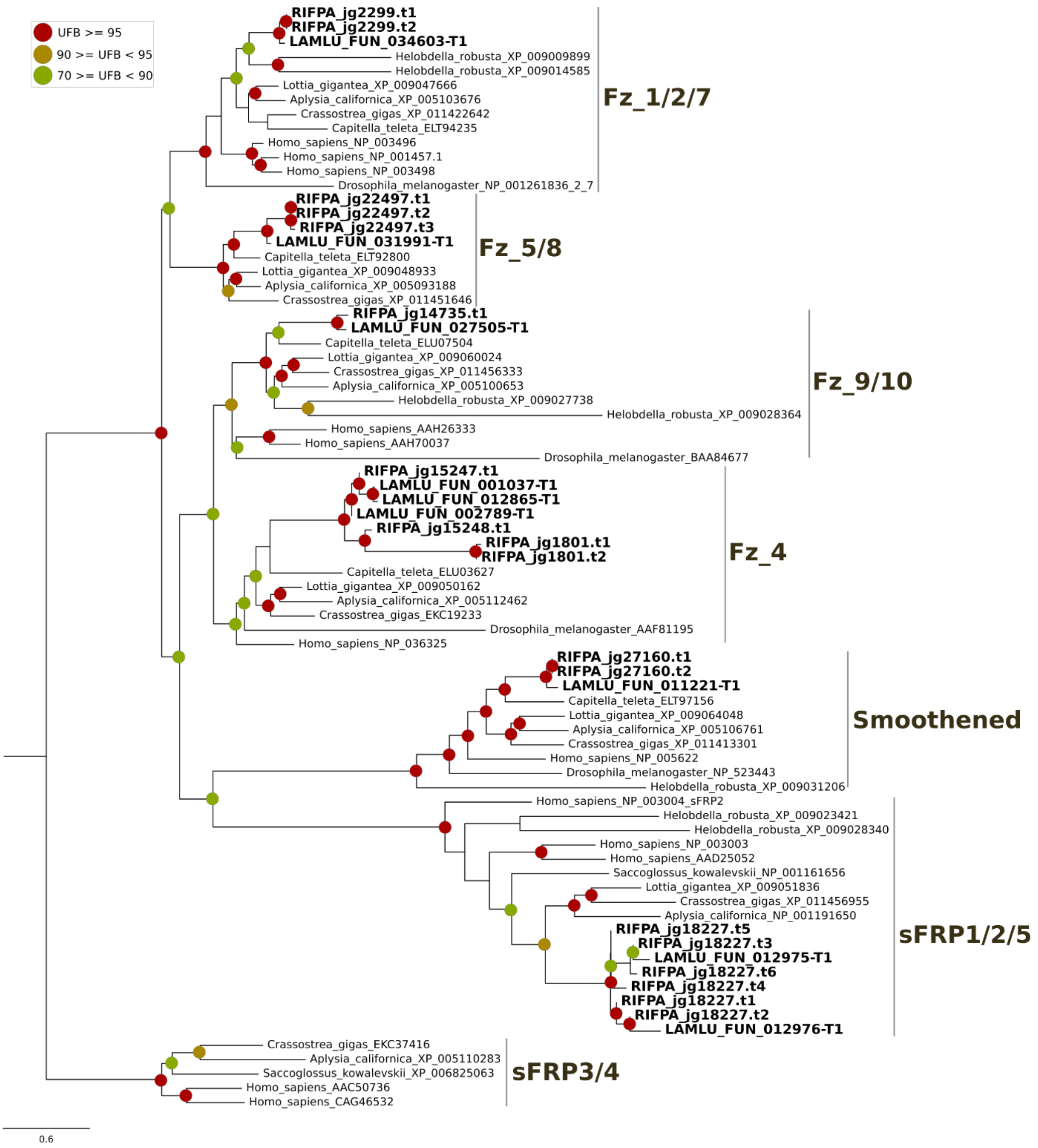

**Supplementary figure 15 | Phylogeny of Wnt receptor genes (Frizzled)**. Maximum-likelihood phylogenetic tree inference of the Wnt receptor genes using 1000 ultrafast bootstrap replicates. The branch support values are represented by the coloured circles in the tree nodes. Red circles represent ultrafast bootstrap values >= 95. Yellow circles represent ultrafast bootstrap values >= 90 and < 95. Green circles represent ultrafast bootstrap values < 90 and >= 70. Ultrafast bootstrap values smaller than 70 are not shown. SFRP3/4 receptor genes were used as outgroup. Accession numbers for NCBI database are displayed after the species names.

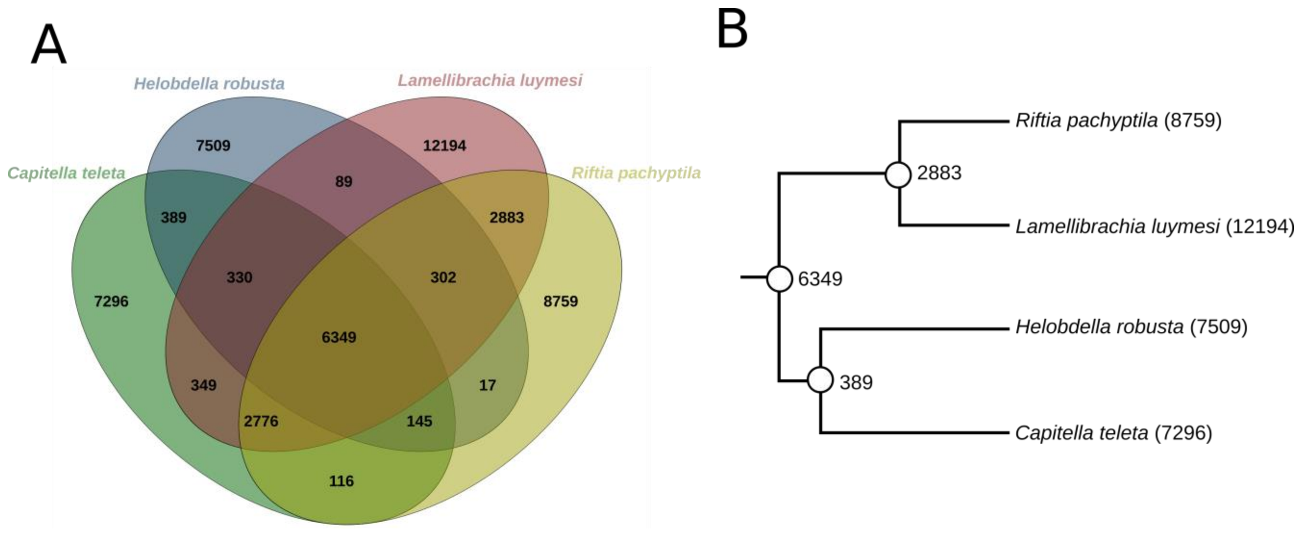

**Supplementary figure 16 | Shared orthogroups in Annelida.** **A**, Shared ortholog groups among *Capitella*, *Helobdella* and the two tubeworms *Lamellibrachia* and *Riftia*. The number of shared groups between *Riftia* and each of the two non-vestimentiferan worms *Helobdella* and *Capitella* is lower than the *Lamellibrachia-Helobdella* and *Lamellibrachia-Capitella* pairs indicating a more derived gene repertoire in the giant tubeworm. **B**, Annelida phylogenetic tree with the shared orthogroups mapped into the nodes. Numbers between in parenthesis represent the total number lineage-specific genes, i.e., orphan genes.

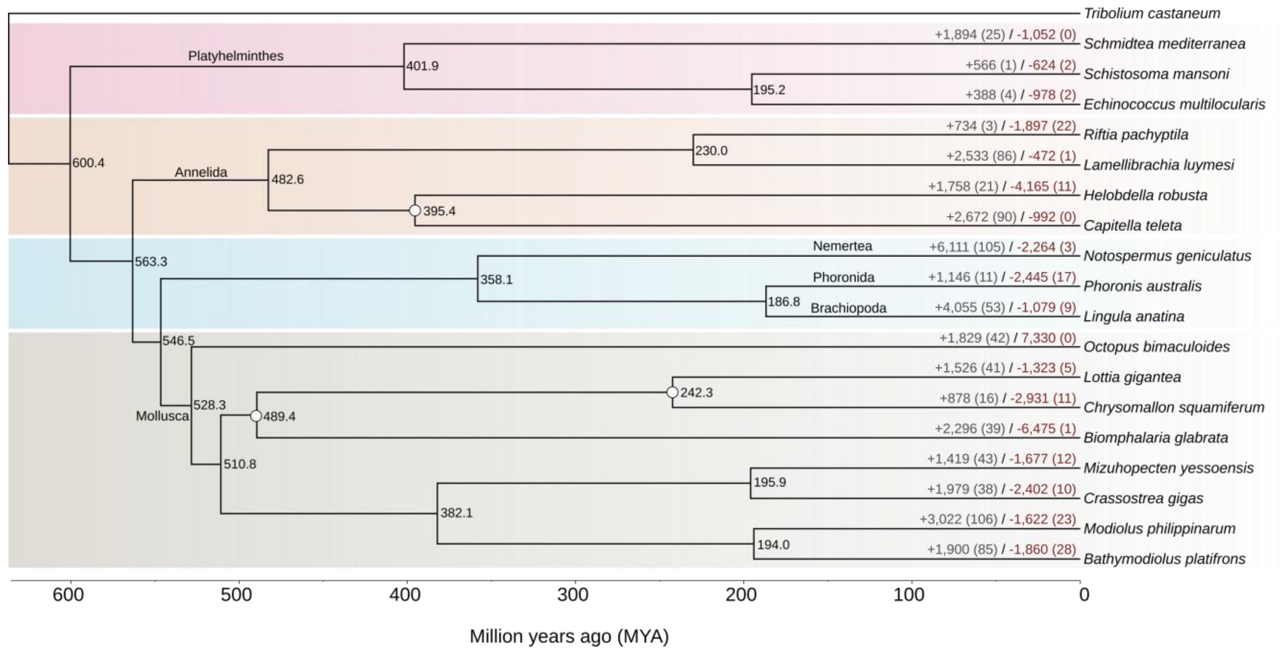

**Supplementary figure 17 | Gene gain and loss in selected lophotrochozoans.** Phylogenetic tree generated with PhyloBayes using available calibrations based on fossil records[10] (white circles in the nodes). The ultrametric tree was obtained after 31,665 rounds discarding the initial 7,916 rounds as burn-in (25%). Gene family evolutionary history of gain and loss was analyzed with CAFE. The number of gene family gains and losses are marked by the plus (+) and minus (-) signs. *Riftia* genome shows a net reduction of gene numbers.

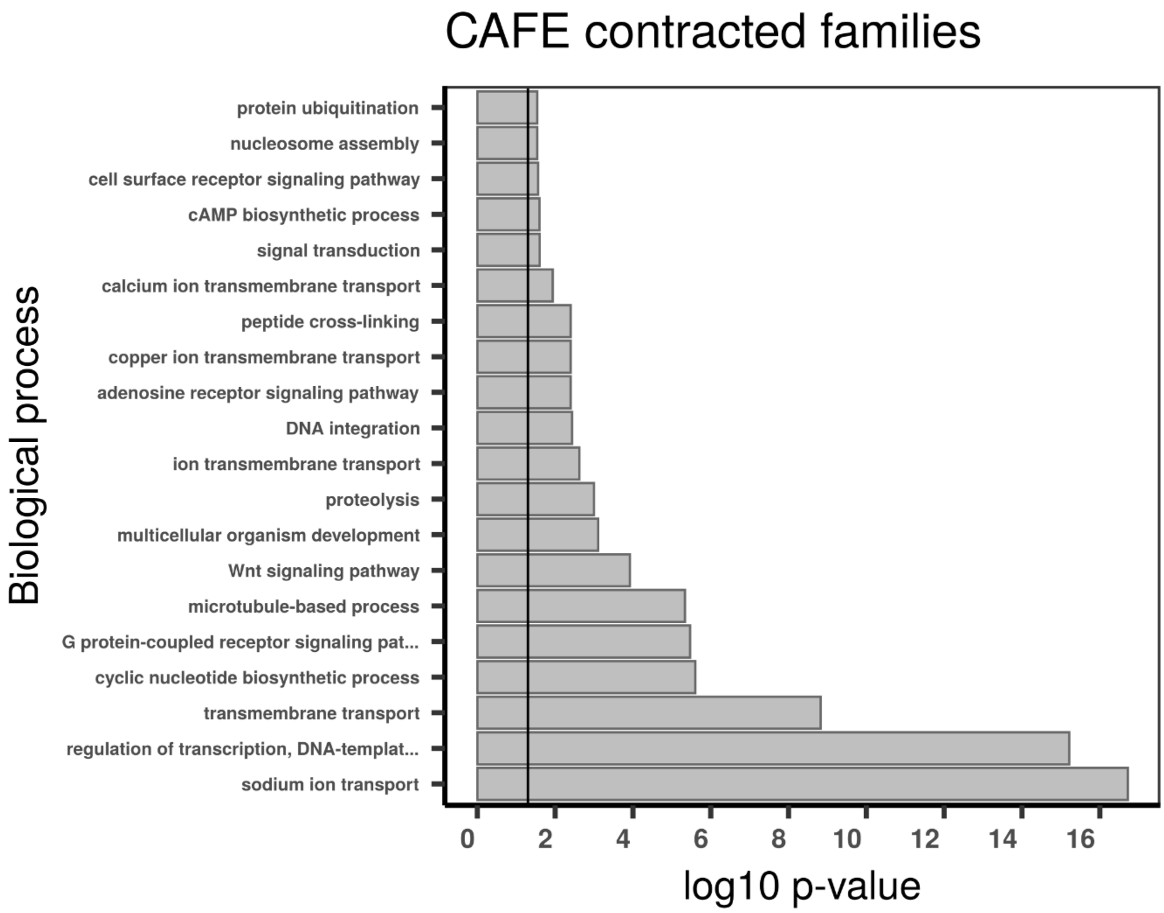

CAFE contracted families
Biological process
protein ubiquitination
nucleosome assembly
cell surface receptor signaling pathway
cAMP biosynthetic process
signal transduction
calcium ion transmembrane transport
peptide cross-linking
copper ion transmembrane transport
adenosine receptor signaling pathway
DNA integration
ion transmembrane transport
proteolysis
multicellular organism development
Wnt signaling pathway
microtubule-based process
G protein-coupled receptor signaling pat...
cyclic nucleotide biosynthetic process
transmembrane transport
regulation of transcription, DNA-templat...
sodium ion transport
0
2
4
6
8
10
12
14
16
log10 p-value

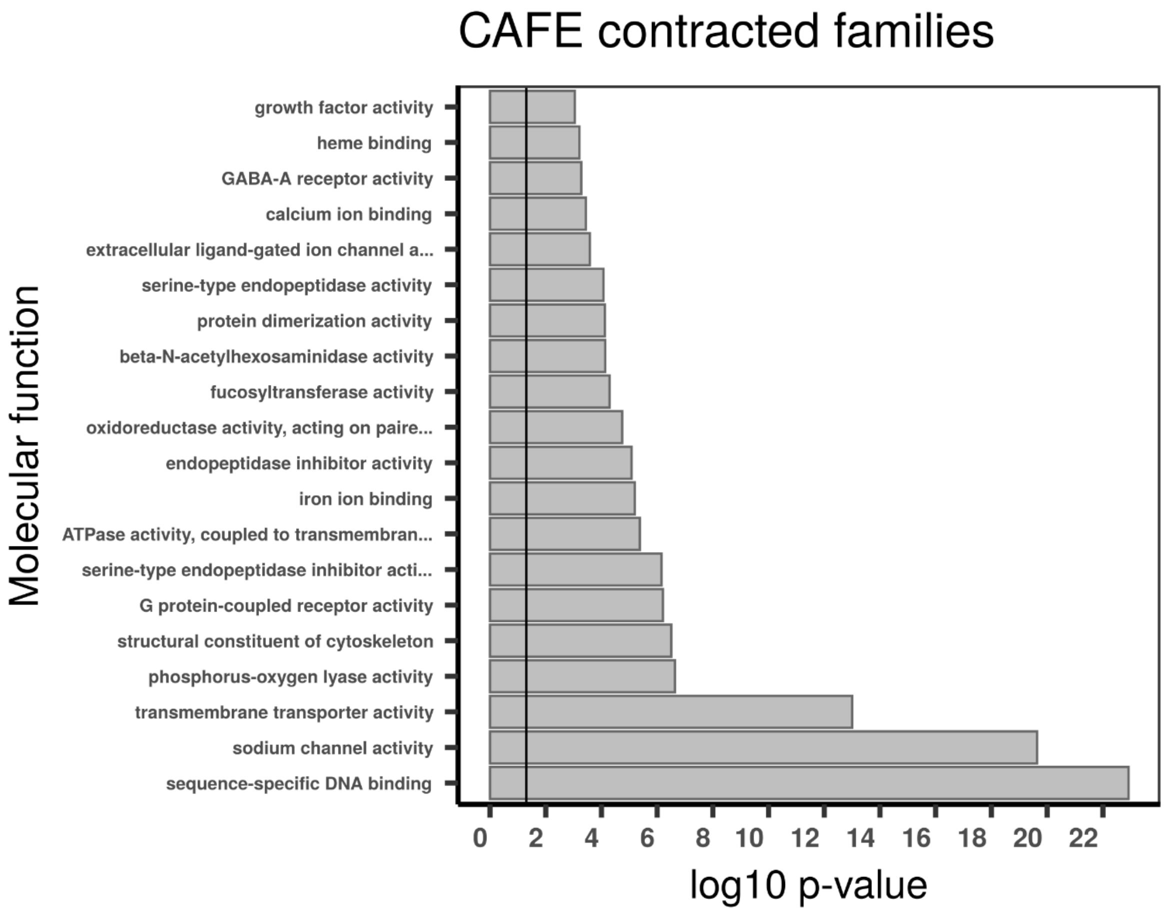

CAFE contracted families
Molecular function
growth factor activity
heme binding
GABA-A receptor activity
calcium ion binding
extracellular ligand-gated ion channel a...
serine-type endopeptidase activity
protein dimerization activity
beta-N-acetylhexosaminidase activity
fucosyltransferase activity
oxidoreductase activity, acting on paire...
endopeptidase inhibitor activity
iron ion binding
ATPase activity, coupled to transmembran...
serine-type endopeptidase inhibitor acti...
G protein-coupled receptor activity
structural constituent of cytoskeleton
phosphorus-oxygen lyase activity
transmembrane transporter activity
sodium channel activity
sequence-specific DNA binding
0
2
4
6
8
10
12
14
16
18
20
22
log10 p-value

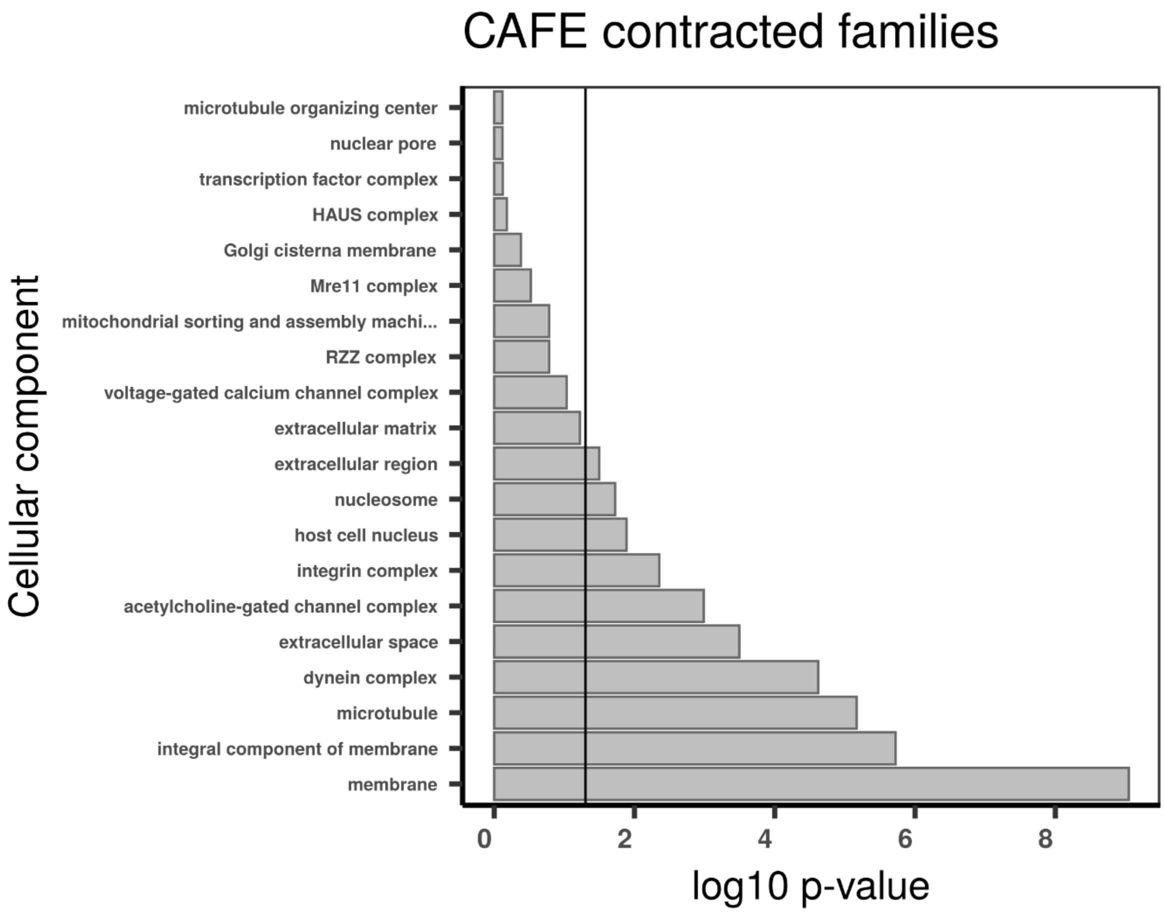

**Supplementary figure 18 | Gene set enrichment analysis with topGO using CAFE contract families in *Riftia***. Gene ontology (GO) enrichment analyses for CAFE contracted families. The graphs correspond to the three domains of ontologies: biological process (BP), molecular function (MF) and cellular component (CC). The selected genes were analysed for enrichment in specific GO categories using the TopGO program against the background (all coding sequence genes). Y-axis corresponds to enriched GO terms found in the respective domains (BP, MF and CC). X-axis correspond to the log function of Fisher p-values obtained for each one of the enriched terms. The back line denotes a p-value = 0.05. P-values greater than 1,30 (log 0.05) indicate statistically significant enriched term. The contracted gene families are not restricted to any specific biological process or molecular function, suggesting that the giant tubeworm genome is undergoing a broad reduction in gene content (i.e., reductive evolution).

A

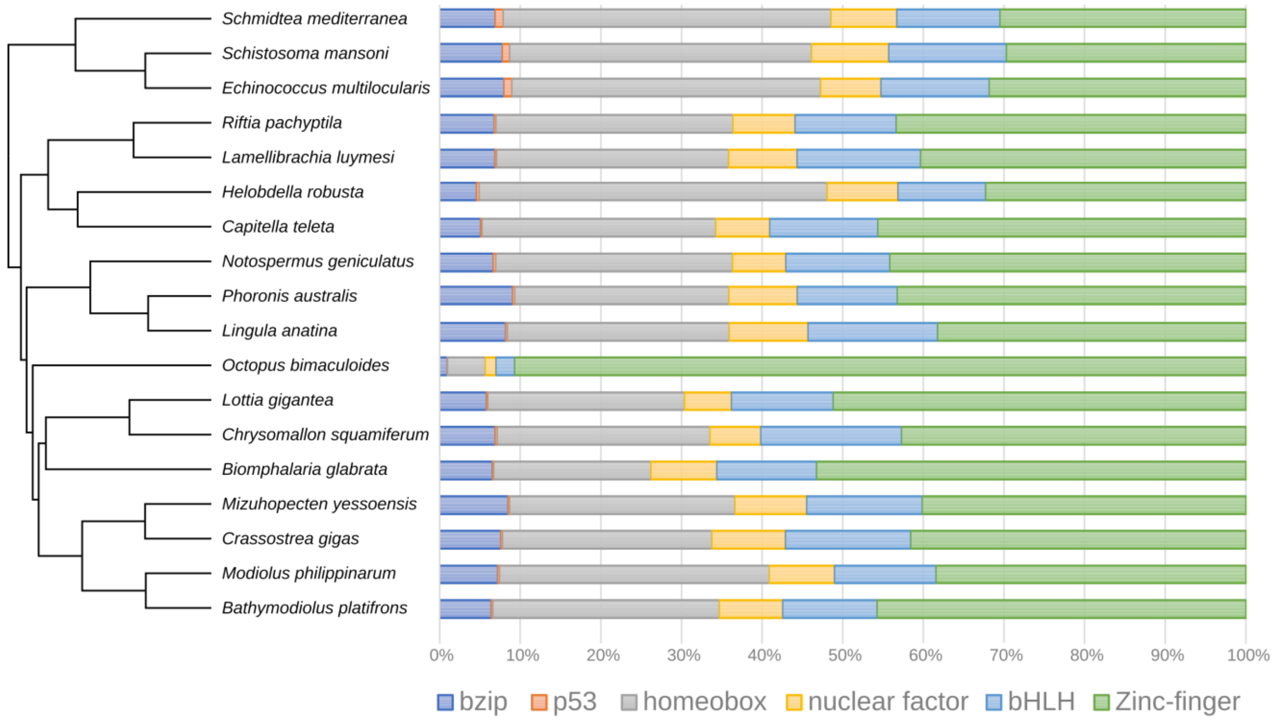

B

| Distribution of the transcription factor PFAM domains among four annelids | | | | | |
|---|---|---|---|---|---|
| **PFAM** | **Domain** | ***R. pachyptila*** | ***L. luymesi*** | ***C. teleta*** | ***H. robusta*** |
| PF00170 | bZIP_1 | 11 | 13 | 10 | 14 |
| PF07716 | bZIP_2 | 12 | 10 | 15 | 9 |
| PF03131 | bZIP_Maf | 5 | 6 | 3 | 3 |
| PF00870 | P53 | 1 | 1 | 1 | 2 |
| PF00046 | Homeodomain | 108 | 103 | 132 | 229 |
| PF05920 | Homeobox_KN | 12 | 12 | 28 | 14 |
| PF04218 | CENP-B_N | 2 | 7 | 0 | 3 |
| PF00105 | zf-C4 | 32 | 36 | 37 | 50 |
| PF00010 | HLH | 52 | 65 | 74 | 62 |
| PF00096 | zf-C2H2 | 10 | 15 | 33 | 24 |
| PF05605 | zf-Di19 | 4 | 3 | 4 | 5 |
| PF12171 | zf-C2H2_jaz | 5 | 4 | 6 | 5 |
| PF12759 | zf-C2H2_2 | 2 | 2 | 2 | 2 |
| PF12874 | zf-met | 7 | 7 | 6 | 4 |
| PF13465 | zf-H2C2_2 | 127 | 115 | 169 | 133 |
| PF13909 | zf-H2C2_5 | 16 | 16 | 23 | 8 |
| PF13912 | zf-C2H2_6 | 5 | 5 | 6 | 2 |
| PF13913 | zf-C2HC_2 | 4 | 4 | 3 | 1 |
| **Total** | | 415 | 424 | 552 | 570 |

**Supplementary figure 19 | Distribution of transcription factors among lophotrochozoans . A,** Percent stacked barchart of the six major classes of transcription factors (TFs) among lophotrochozoans. *Riftia pachyptila* TF complement is similar to other lophotrochozoans. *Octopus bimaculoides* complement is composed of a massive expansion of zinc-finger elements, as previously described[9]. **B,** Distribution of the transcription factor PFAM domains among four annelids. *Riftia* TF complement is the lowest among the annelids.

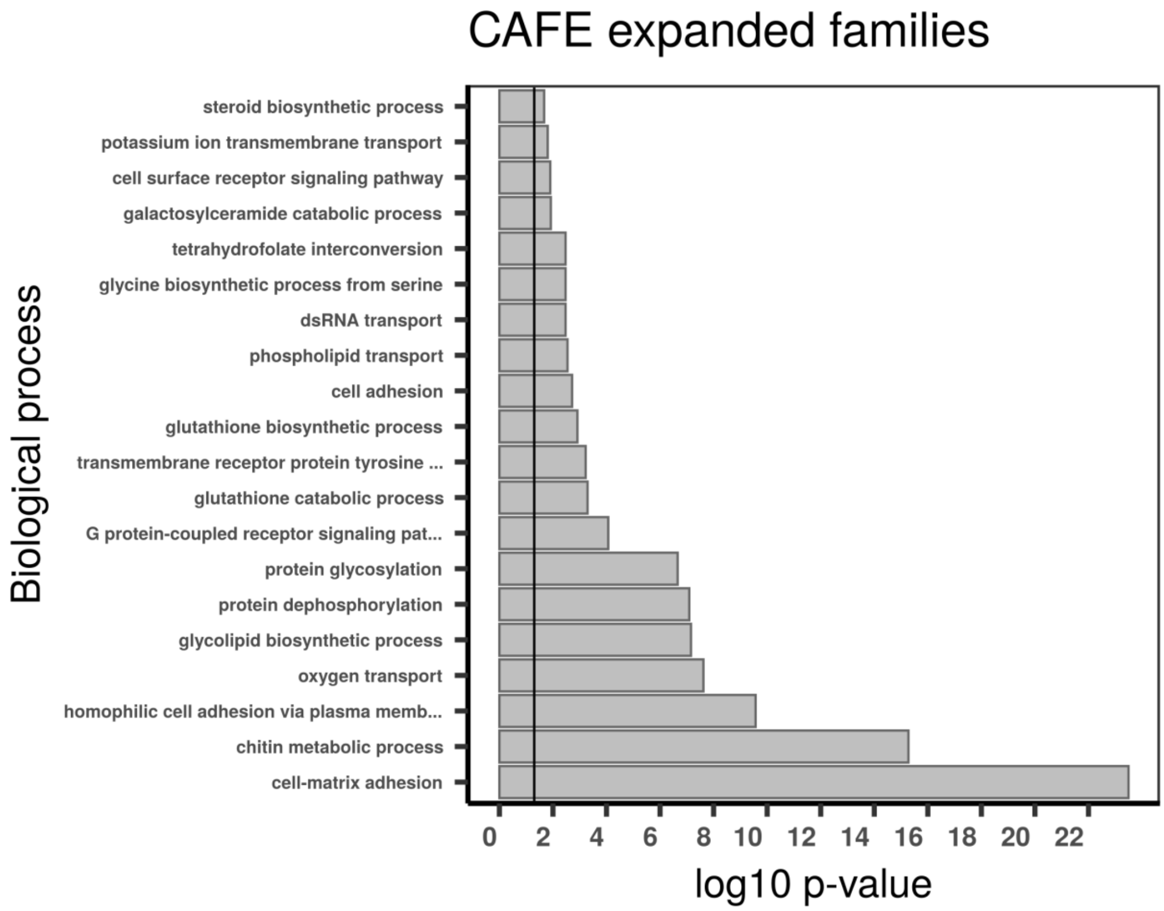

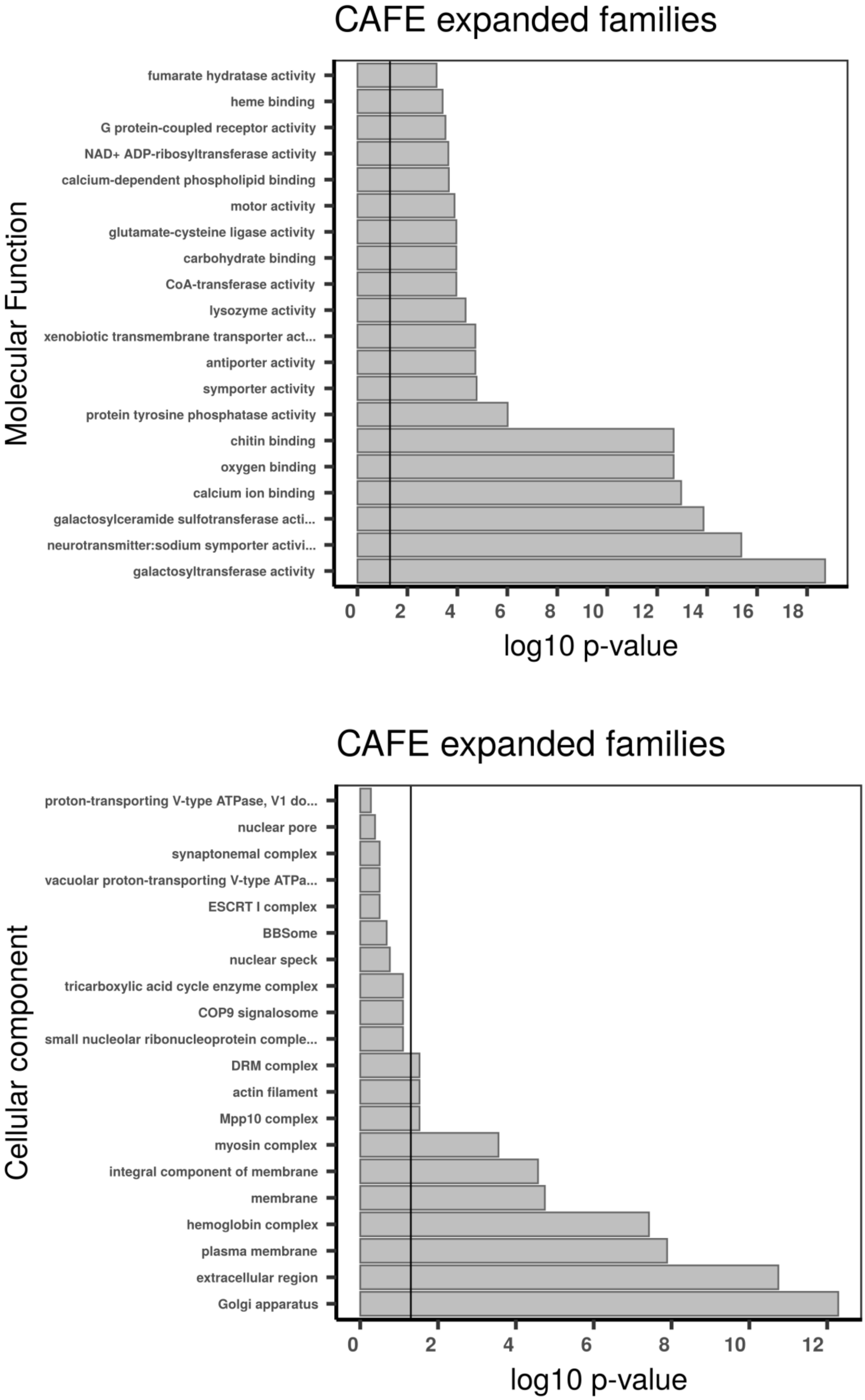

**Supplementary figure 20 | Gene set enrichment analysis with topGO using CAFE expanded families in *Riftia*.** Gene ontology (GO) enrichment analyses for CAFE expanded families. The graphs correspond to the three domains of ontologies: biological process (BP), molecular function (MF) and cellular component (CC). The selected genes were analysed for enrichment in specific GO categories using the TopGO program against the background (all coding sequence genes). Y axis corresponds to enriched GO terms found in the respective domains (BP, MF and CC). X axis correspond to the log function of Fisher p-values obtained for each one of the enriched terms. The back line denotes a p-value = 0.05. P-values greater than 1,30 (log 0.05) indicate statistically significant enriched term. Genes involved in sulphur metabolism and detoxification, anti-oxidative stress, oxygen transport, neurotransmitter- and ion channel-related functions, digestion (lysozyme activity) and secretion of chitinous material are expanded in the *Riftia pachyptila* tubeworm genome.

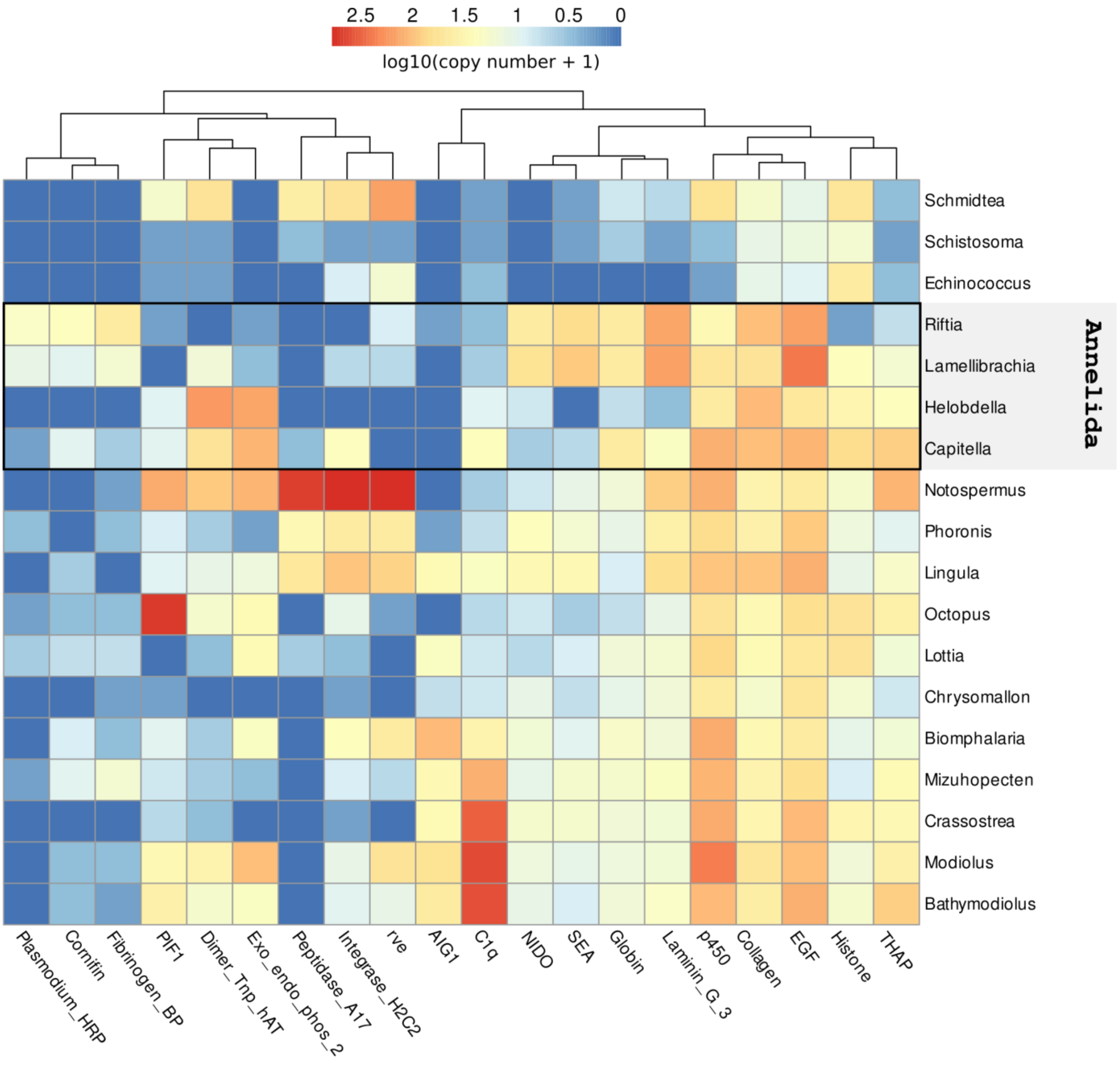

**Supplementary figure 21 | Distribution of *Riftia* expanded/contracted PFAM domains among selected lophotrochozoans.** Log distribution of the number of *Riftia* expanded/contracted PFAM domains among flatworms (*Schmidtea*, *Schistosoma*, *Echinococcus*), *molluscs* (*Octopus*, *Lottia*, *Chrysomallon*, *Biomphalaria*, *Mizuhopecten*, *Crassotrea*, *Modiolus*, *Bathymodiolus*), brachiopods (*Lingula*), nemerteans (*Notospermus*) and phoronids (*Phoronis*). PFAM domains were considered enriched or contracted in *Riftia* using two-tailed Fisher exact test (p-value < 0.01 after correction)

against the outgroup average (i.e., lophotrochozoans minus *Riftia*). Multiple protein domains in a single given gene were counted only once. Expanded domains in *Riftia* include globin, NIDO, Laminin_G_3, EGF and collagen PFAMs.

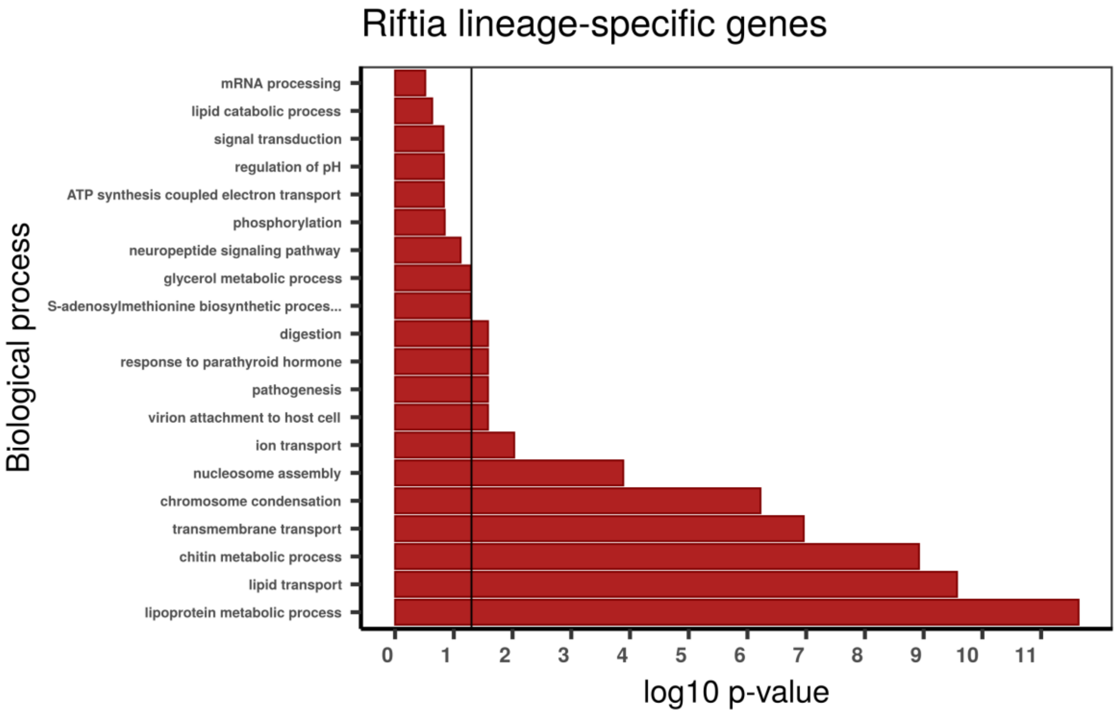

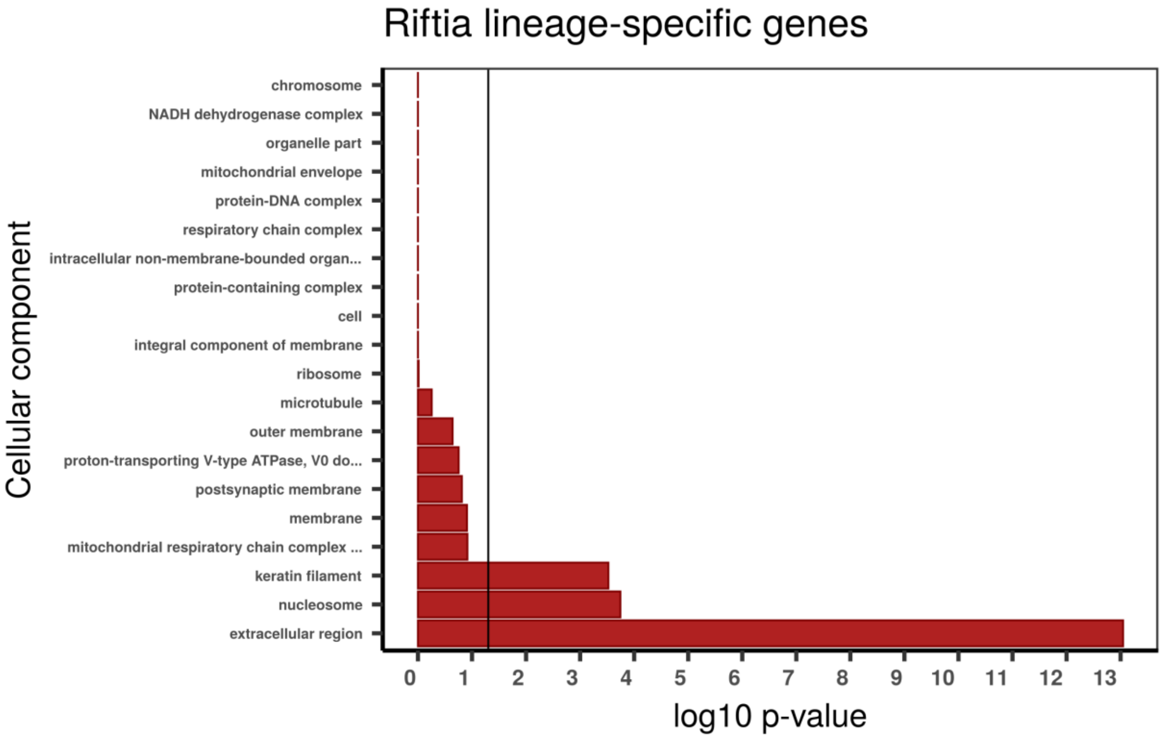

**Supplementary figure 22 | Gene set enrichment analysis with topGO using *Riftia pachyptila* lineage-specific genes**. Gene ontology (GO) enrichment analyses for *Riftia pachyptila* lineage-specific genes. The graphs correspond to the three domains of ontologies: biological process (BP), molecular function (MF) and cellular component (CC). The selected genes were analysed for enrichment in specific GO categories using the TopGO program against the background (all coding sequence genes). Y-axis corresponds to enriched GO terms found in the respective domains (BP, MF and CC). X-axis correspond to the log function of Fisher p-values obtained for each one of the enriched terms. The back line denotes a p-value = 0.05. P-values greater than 1,30 (log 0,05) indicate statistically significant enriched term. Genes involved in cell cycle and chitin production and secretion are enriched in *Riftia*.

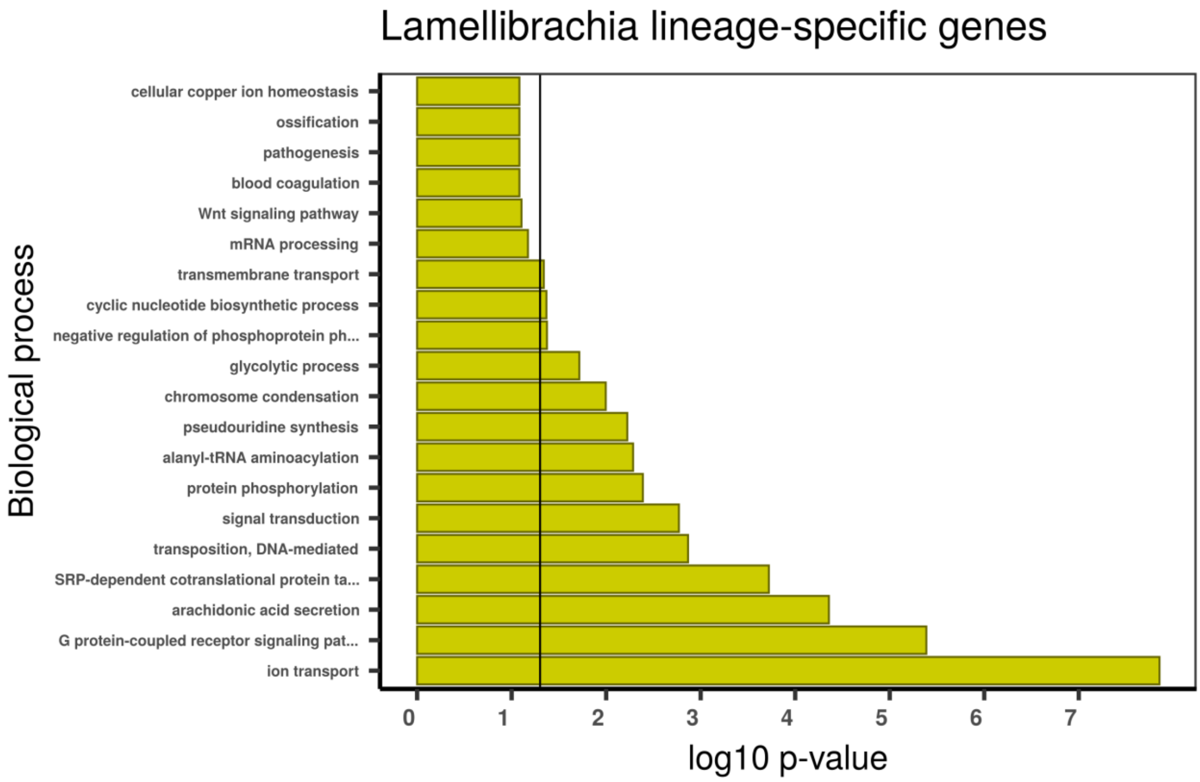

Lamellibrachia lineage-specific genes
Biological process
cellular copper ion homeostasis
ossification
pathogenesis
blood coagulation
Wnt signaling pathway
mRNA processing
transmembrane transport
cyclic nucleotide biosynthetic process
negative regulation of phosphoprotein ph...
glycolytic process
chromosome condensation
pseudouridine synthesis
alanyl-tRNA aminoacylation
protein phosphorylation
signal transduction
transposition, DNA-mediated
SRP-dependent cotranslational protein ta...
arachidonic acid secretion
G protein-coupled receptor signaling pat...
ion transport
0 1 2 3 4 5 6 7
log10 p-value

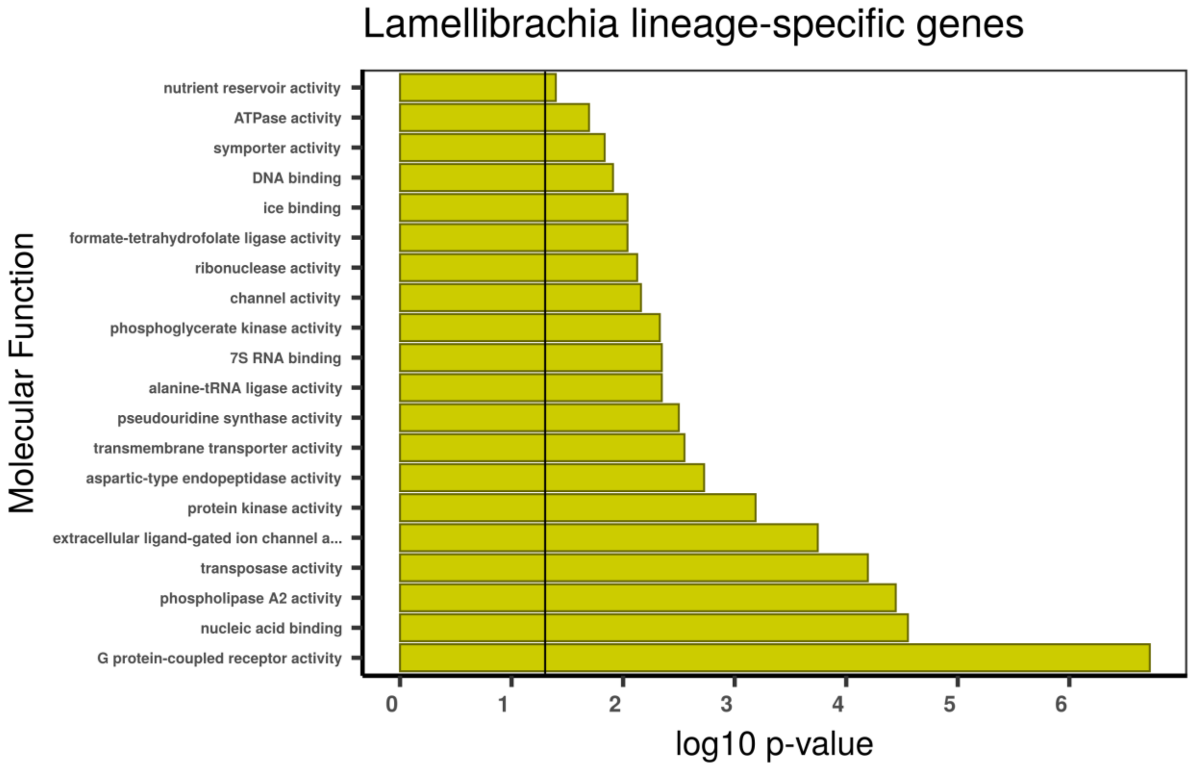

Lamellibrachia lineage-specific genes
Molecular Function
nutrient reservoir activity
ATPase activity
symporter activity
DNA binding
ice binding
formate-tetrahydrofolate ligase activity
ribonuclease activity
channel activity
phosphoglycerate kinase activity
7S RNA binding
alanine-tRNA ligase activity
pseudouridine synthase activity
transmembrane transporter activity
aspartic-type endopeptidase activity
protein kinase activity
extracellular ligand-gated ion channel a...
transposase activity
phospholipase A2 activity
nucleic acid binding
G protein-coupled receptor activity
0 1 2 3 4 5 6
log10 p-value

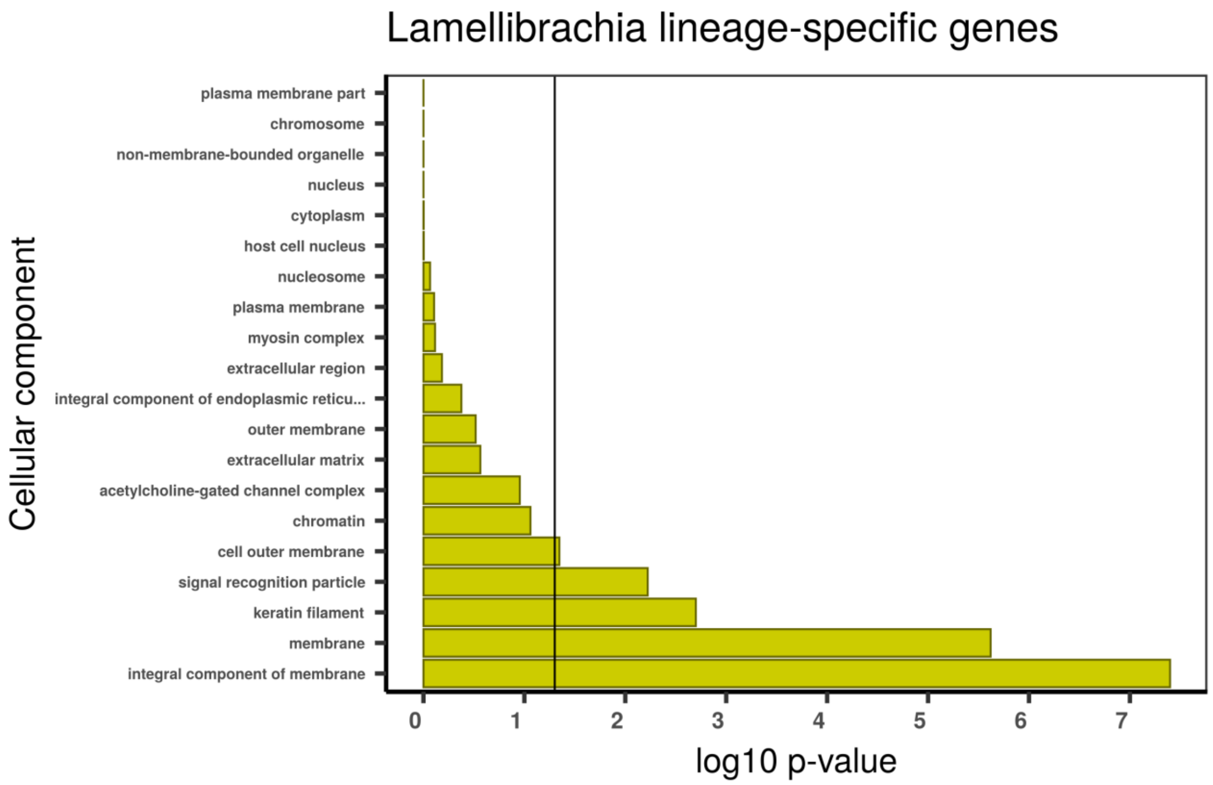

**Supplementary figure 23 | Gene set enrichment analysis with topGO using *Lamellibrachia luymesi* lineage-specific genes**. Gene ontology (GO) enrichment analyses for *Lamellibrachia luymesi* lineage-specific genes. The graphs correspond to the three domains of ontologies: biological process (BP), molecular function (MF) and cellular component (CC). The selected genes were analysed for enrichment in specific GO categories using the TopGO program against the background (all coding sequence genes). Y-axis corresponds to enriched GO terms found in the respective domains (BP, MF and CC). X-axis correspond to the log function of Fisher p-values obtained for each one of the enriched terms. The back line denotes a p-value = 0.05. P-values greater than 1,30 (log 0,05) indicate statistically significant enriched term. Genes involved in transposase activity and cell cycle are enriched in *Lamellibrachia*.

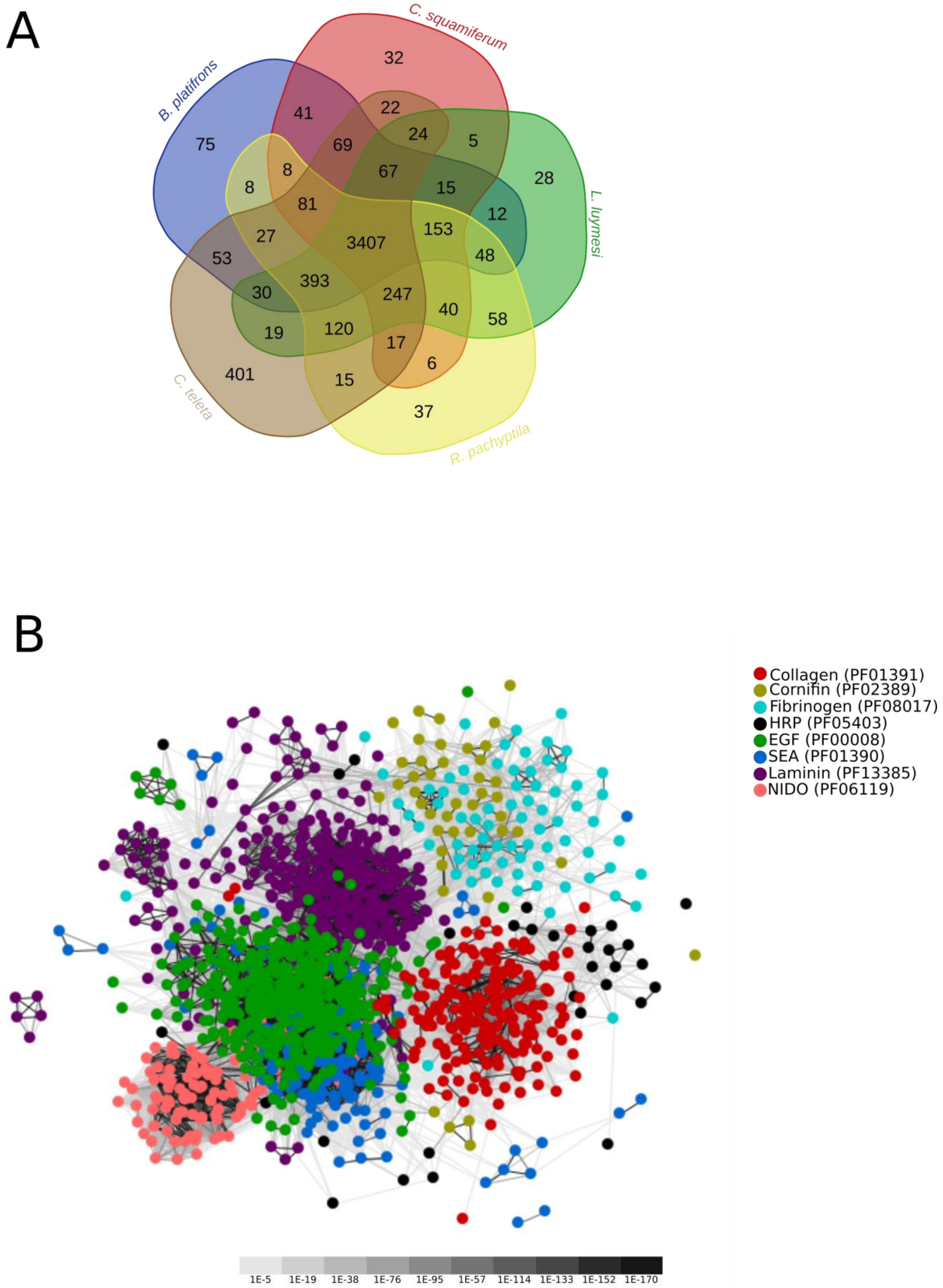

**Supplementary figure 24 | PFAM functional domains shared among selected deep vent lophotrochozoans and cluster analysis of selected *Riftia* expanded domains.** Venn diagram depicting the shared PFAM functional domains among tubeworms (*Riftia* and *Lamellibrachia*), one bivalve (*Bathymodiolus platifrons*), one gastropod (*Chrysomallon squamiferum*) and the polychaete *Capitella teleta.* **B,** 2D-cluster map of collagen, cornifin, fibrinogen, HRP, EGF, SEA, Laminin and NIDO domain-containing proteins in *Riftia*. Colours are based on the different PFAM domain. Edges correspond to blastp connections of p-value < 1e-06. The high connectivity of the cluster is probably due the presence of small repetitive protein domains in the genes (EGF, collagen), rather than evolutionary relatedness.

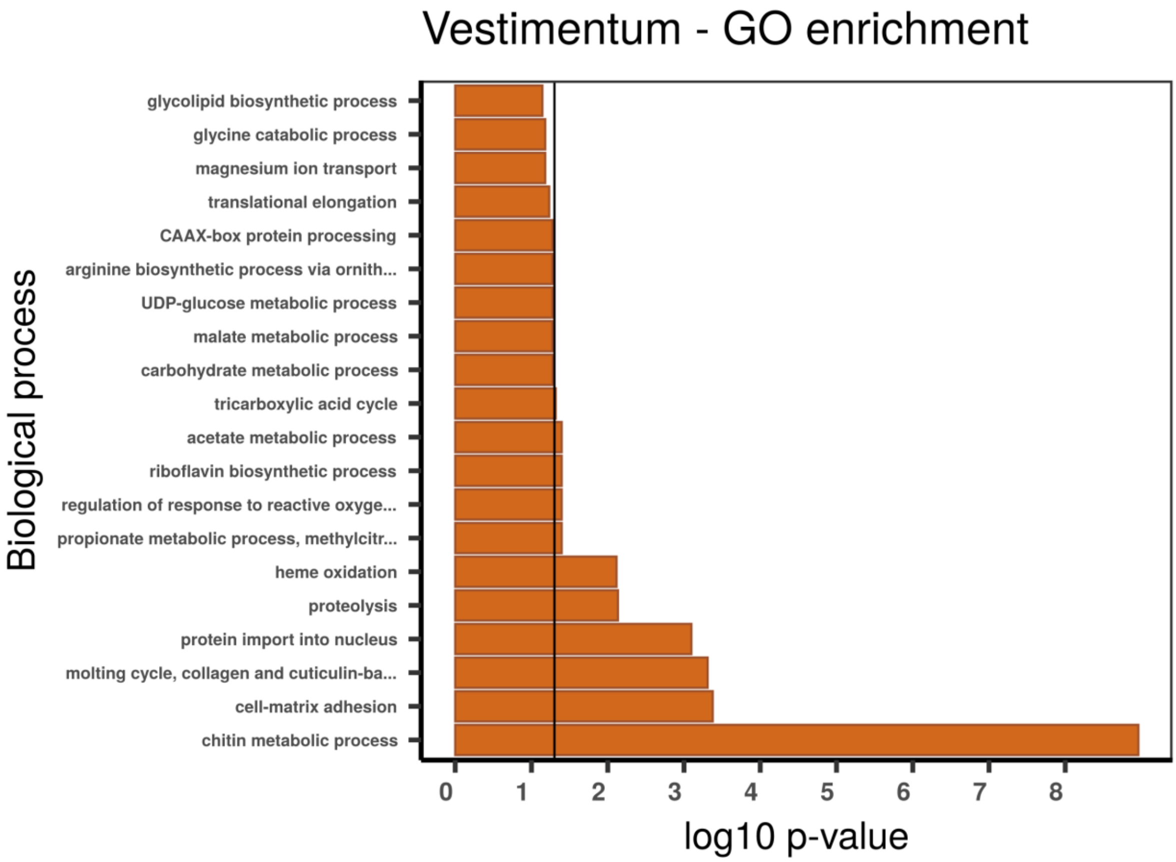

Vestimentum - GO enrichment
Biological process
glycolipid biosynthetic process
glycine catabolic process
magnesium ion transport
translational elongation
CAAX-box protein processing
arginine biosynthetic process via ornith...
UDP-glucose metabolic process
malate metabolic process
carbohydrate metabolic process
tricarboxylic acid cycle
acetate metabolic process
riboflavin biosynthetic process
regulation of response to reactive oxyge...
propionate metabolic process, methylcitr...
heme oxidation
proteolysis
protein import into nucleus
molting cycle, collagen and cuticulin-ba...
cell-matrix adhesion
chitin metabolic process
0 1 2 3 4 5 6 7 8
log10 p-value

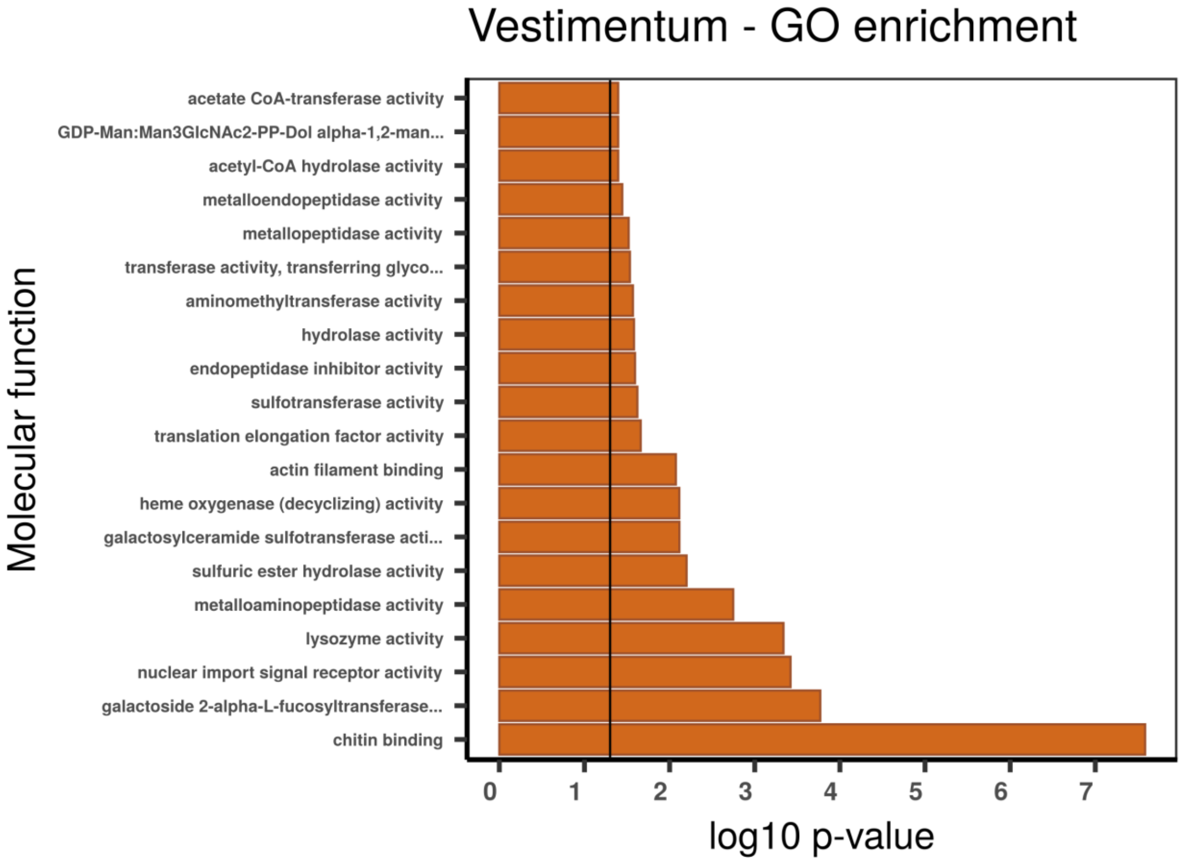

Vestimentum - GO enrichment
Molecular function
acetate CoA-transferase activity
GDP-Man:Man3GlcNAc2-PP-Dol alpha-1,2-man...
acetyl-CoA hydrolase activity
metalloendopeptidase activity
metallopeptidase activity
transferase activity, transferring glyco...
aminomethyltransferase activity
hydrolase activity
endopeptidase inhibitor activity
sulfotransferase activity
translation elongation factor activity
actin filament binding
heme oxygenase (decyclizing) activity
galactosylceramide sulfotransferase acti...
sulfuric ester hydrolase activity
metalloaminopeptidase activity
lysozyme activity
nuclear import signal receptor activity
galactoside 2-alpha-L-fucosyltransferase...
chitin binding
0 1 2 3 4 5 6 7
log10 p-value

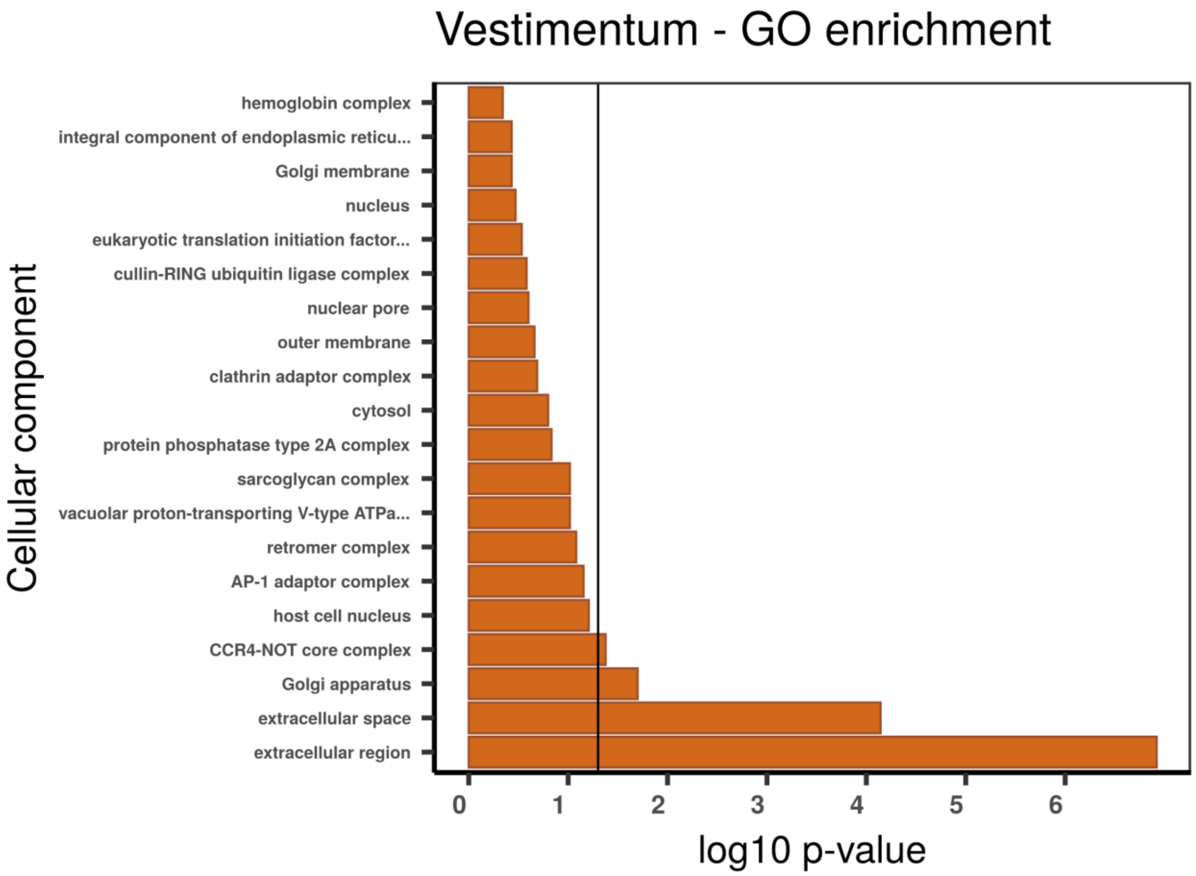

**Supplementary figure 25 | Gene set enrichment analysis with topGO using absolutely vestimentum specific TAU genes**. Gene ontology (GO) enrichment analyses for absolutely vestimentum specific TAU genes. The graphs correspond to the three domains of ontologies: biological process (BP), molecular function (MF) and cellular component (CC). The selected genes were analysed for enrichment in specific GO categories using the TopGO program against the background (all coding sequence genes). Y axis corresponds to enriched GO terms found in the respective domains (BP, MF and CC). X axis correspond to the log function of Fisher p-values obtained for each one of the enriched terms. The back line denotes a p-value = 0.05. P-values greater than 1,30 (log 0,05) indicate statistically significant enriched term. Genes involved chitin metabolism are differentially expressed in the vestimentum

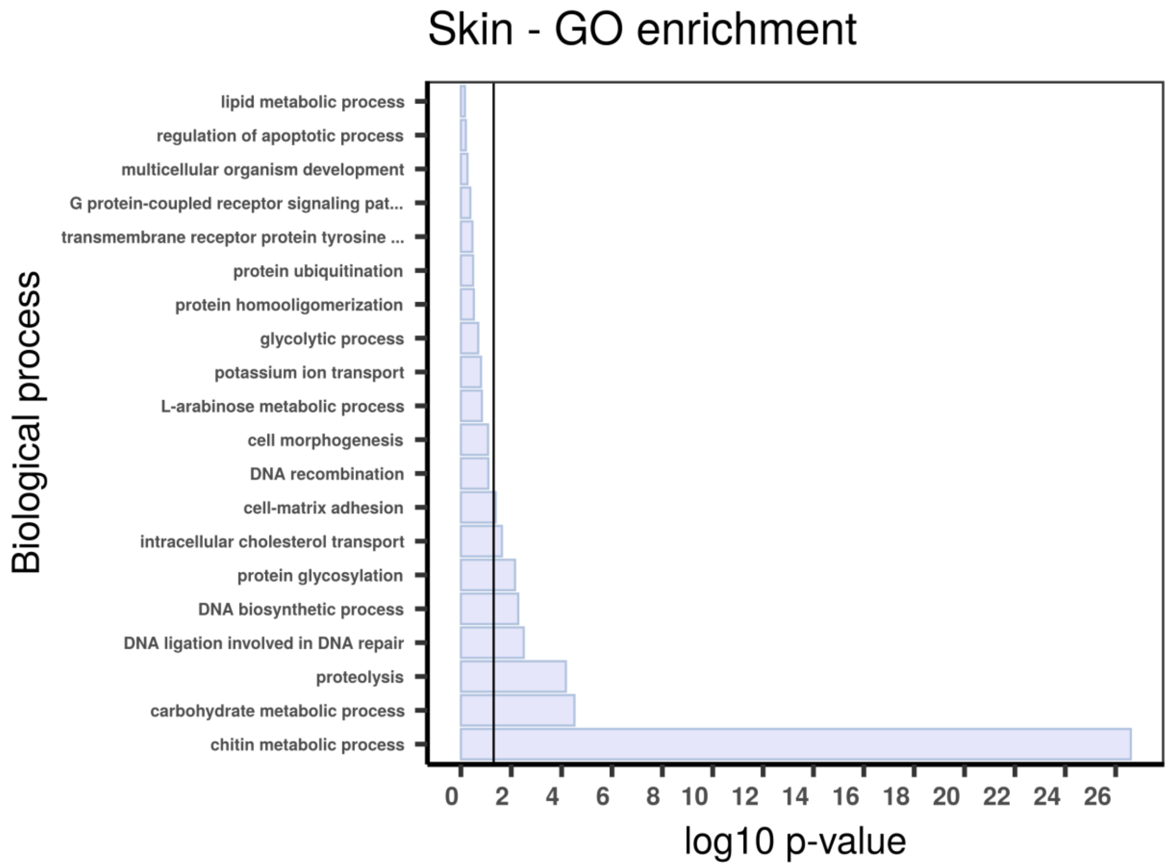

Skin - GO enrichment
Biological process
lipid metabolic process
regulation of apoptotic process
multicellular organism development
G protein-coupled receptor signaling pat...
transmembrane receptor protein tyrosine ...
protein ubiquitination
protein homooligomerization
glycolytic process
potassium ion transport
L-arabinose metabolic process
cell morphogenesis
DNA recombination
cell-matrix adhesion
intracellular cholesterol transport
protein glycosylation
DNA biosynthetic process
DNA ligation involved in DNA repair
proteolysis
carbohydrate metabolic process
chitin metabolic process
0 2 4 6 8 10 12 14 16 18 20 22 24 26
log10 p-value

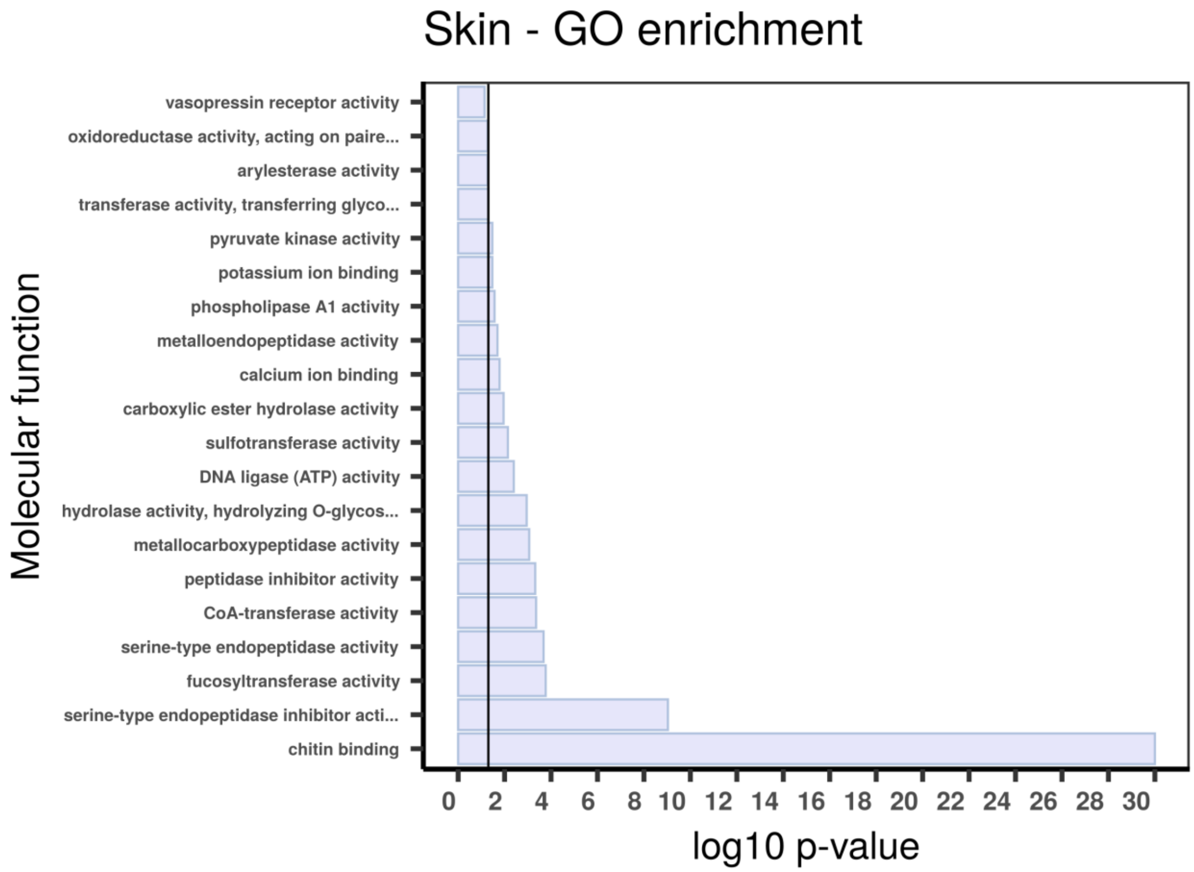

Skin - GO enrichment
Molecular function
vasopressin receptor activity
oxidoreductase activity, acting on paire...
arylesterase activity
transferase activity, transferring glyco...
pyruvate kinase activity
potassium ion binding
phospholipase A1 activity
metalloendopeptidase activity
calcium ion binding
carboxylic ester hydrolase activity
sulfotransferase activity
DNA ligase (ATP) activity
hydrolase activity, hydrolyzing O-glycos...
metallocarboxypeptidase activity
peptidase inhibitor activity
CoA-transferase activity
serine-type endopeptidase activity
fucosyltransferase activity
serine-type endopeptidase inhibitor acti...
chitin binding
0 2 4 6 8 10 12 14 16 18 20 22 24 26 28 30
log10 p-value

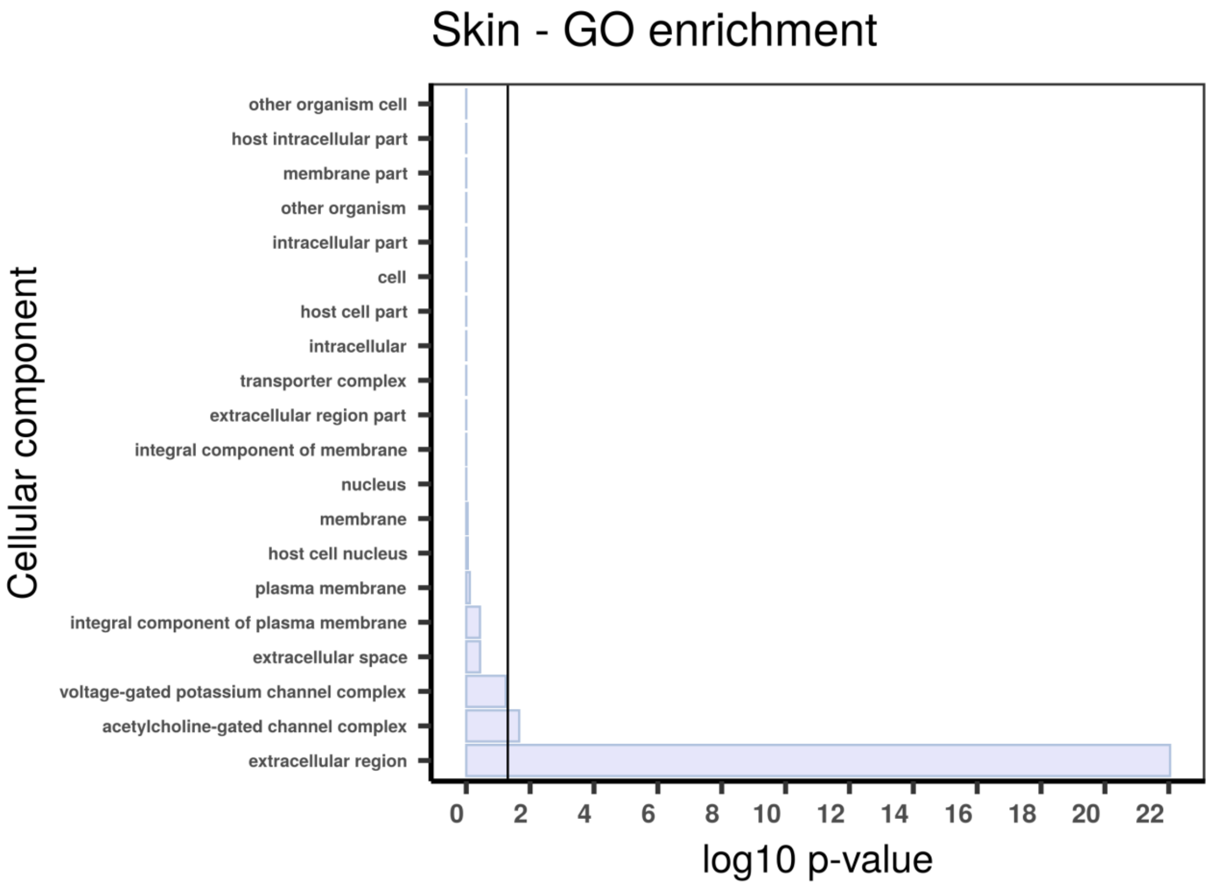

**Supplementary figure 26 | Gene set enrichment analysis with topGO using absolutely body wall (skin) specific TAU genes.** Gene ontology (GO) enrichment analyses for absolutely skin (body wall) specific TAU genes. The graphs correspond to the three domains of ontologies: biological process (BP), molecular function (MF) and cellular component (CC). The selected genes were analysed for enrichment in specific GO categories using the TopGO program against the background (all coding sequence genes). Y axis corresponds to enriched GO terms found in the respective domains (BP, MF and CC). X axis correspond to the log function of Fisher p-values obtained for each one of the enriched terms. The back line denotes a p-value = 0.05. P-values greater than 1,30 (log 0,05) indicate statistically significant enriched term. Genes involved chitin metabolism are differentially expressed in the body wall (skin).

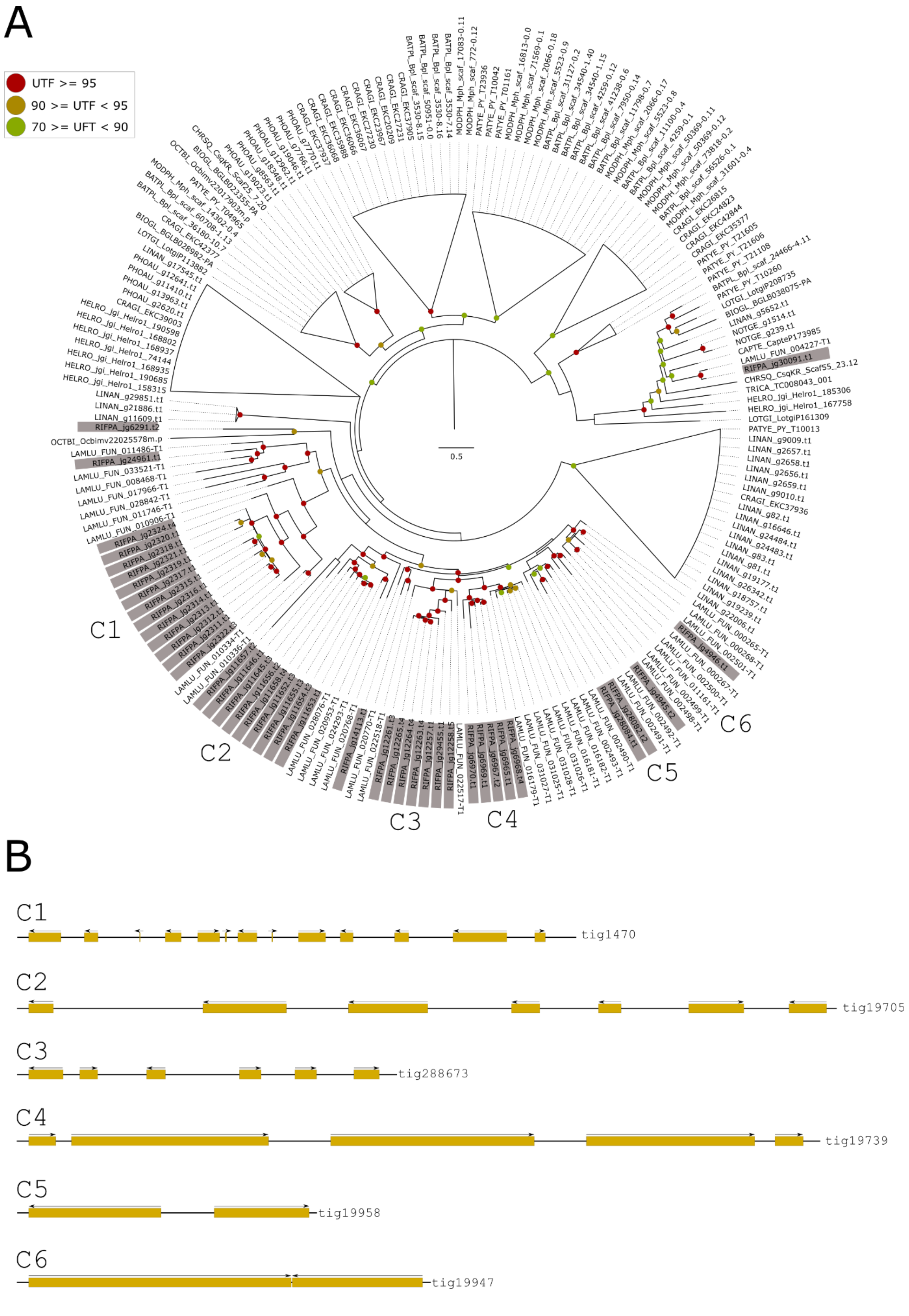

**Supplementary figure 27 | Phylogeny and genomic organisation of sushi genes in *Riftia pachyptila*. A,** Maximum-likelihood phylogenetic tree inference of the sushi genes using 1000 ultrafast bootstrap replicates. The branch support values are represented by the coloured circles in the tree nodes. Red circles represent ultrafast bootstrap values >= 95. Yellow circles represent ultrafast bootstrap values >= 90 and < 95. Green circles represent ultrafast bootstrap values < 90 and >= 70. Ultrafast bootstrap values smaller than 70 are not shown. Gene identification is derived from the publicly available annotated lophotrochozoan genomes. *Riftia* contains the greatest number of sushi genes in the analysed lophotrochozoans. It is possible to note an independent expansion of sushi genes in the base of Vestimentifera (*Lamellibrachia* + *Riftia*). **B**, Genomic organisation of sushi genes in *Riftia* genome. Sushi genes are organised in six different genomic clusters possibly originated through a series of tandem duplications.

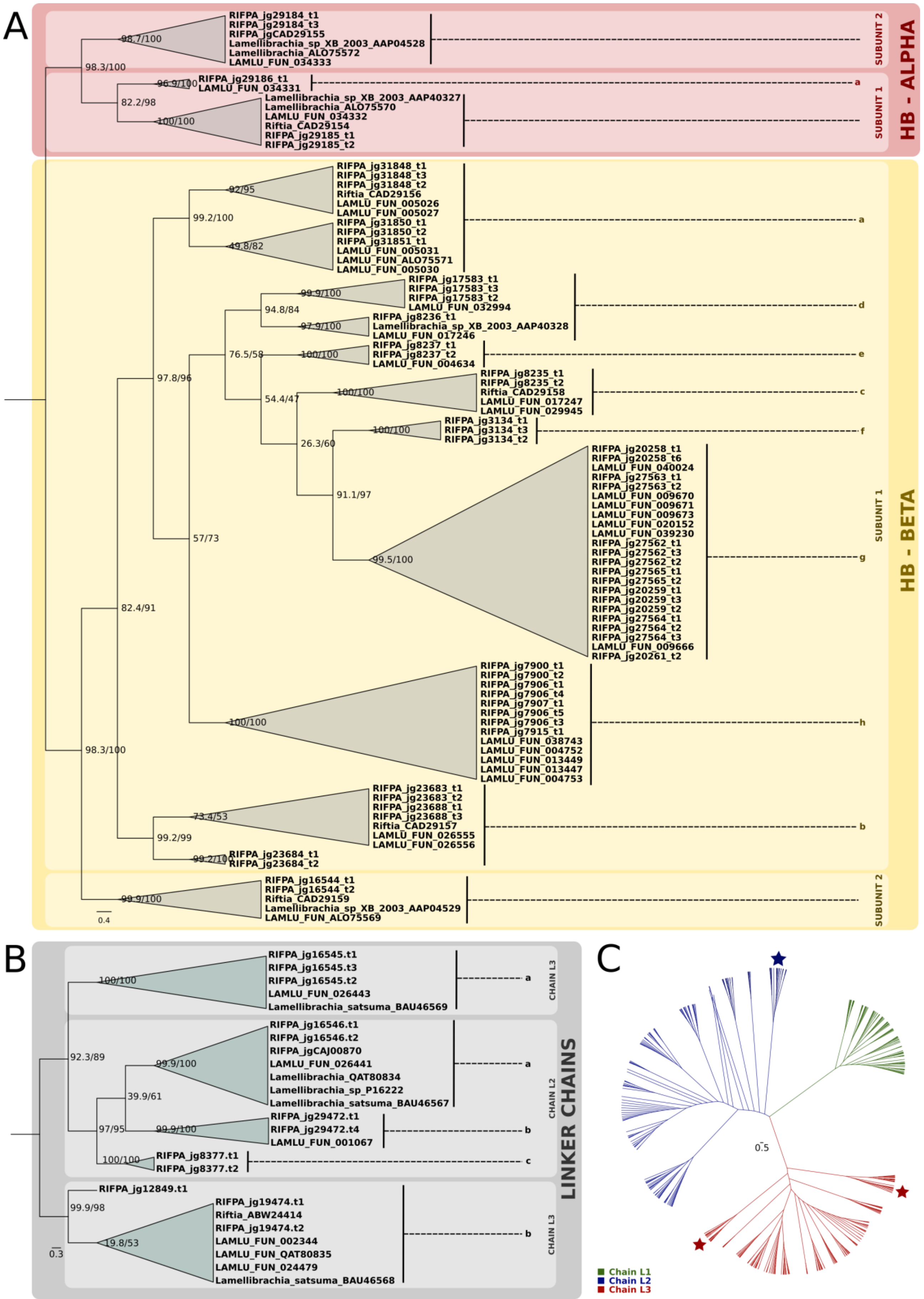

A
HB - ALPHA
SUBUNIT 2
SUBUNIT 1
RIFPA_jg29184_t1
RIFPA_jg29184_t3
RIFPA_jgCAD29155
Lamellibrachia_sp_XB_2003_AAP04528
Lamellibrachia_ALO75572
LAMLU_FUN_034333
98.7/100
98.3/100
96.9/100
RIFPA_jg29186_t1
LAMLU_FUN_034331
82.2/98
100/100
Lamellibrachia_sp_XB_2003_AAP40327
Lamellibrachia_ALO75570
LAMLU_FUN_034332
Riftia_CAD29154
RIFPA_jg29185_t1
RIFPA_jg29185_t2
a
HB - BETA
RIFPA_jg31848_t1
RIFPA_jg31848_t3
RIFPA_jg31848_t2
Riftia_CAD29156
LAMLU_FUN_005026
LAMLU_FUN_005027
RIFPA_jg31850_t1
RIFPA_jg31850_t2
RIFPA_jg31851_t1
LAMLU_FUN_005031
LAMLU_FUN_ALO75571
LAMLU_FUN_005030
92/95
99.2/100
49.8/82
RIFPA_jg17583_t1
RIFPA_jg17583_t3
RIFPA_jg17583_t2
LAMLU_FUN_032994
RIFPA_jg8236_t1
Lamellibrachia_sp_XB_2003_AAP40328
LAMLU_FUN_017246
99.9/100
94.8/84
97.9/100
d
RIFPA_jg8237_t1
RIFPA_jg8237_t2
LAMLU_FUN_004634
76.5/58
100/100
e
RIFPA_jg8235_t1
RIFPA_jg8235_t2
Riftia_CAD29158
LAMLU_FUN_017247
LAMLU_FUN_029945
97.8/96
54.4/47
100/100
c
RIFPA_jg3134_t1
RIFPA_jg3134_t3
RIFPA_jg3134_t2
100/100
26.3/60
f
RIFPA_jg20258_t1
RIFPA_jg20258_t6
LAMLU_FUN_040024
RIFPA_jg27563_t1
RIFPA_jg27563_t2
LAMLU_FUN_009670
LAMLU_FUN_009671
LAMLU_FUN_009673
LAMLU_FUN_020152
LAMLU_FUN_039230
RIFPA_jg27562_t1
RIFPA_jg27562_t3
RIFPA_jg27562_t2
RIFPA_jg27565_t1
RIFPA_jg27565_t2
RIFPA_jg20259_t1
RIFPA_jg20259_t3
RIFPA_jg20259_t2
RIFPA_jg27564_t1
RIFPA_jg27564_t2
RIFPA_jg27564_t3
LAMLU_FUN_009666
RIFPA_jg20261_t2
91.1/97
57/73
99.5/100
g
82.4/91
RIFPA_jg7900_t1
RIFPA_jg7900_t2
RIFPA_jg7906_t1
RIFPA_jg7906_t4
RIFPA_jg7907_t1
RIFPA_jg7906_t5
RIFPA_jg7906_t3
RIFPA_jg7915_t1
LAMLU_FUN_038743
LAMLU_FUN_004752
LAMLU_FUN_013449
LAMLU_FUN_013447
LAMLU_FUN_004753
100/100
h
98.3/100
RIFPA_jg23683_t1
RIFPA_jg23683_t2
RIFPA_jg23688_t1
RIFPA_jg23688_t3
Riftia_CAD29157
LAMLU_FUN_026555
LAMLU_FUN_026556
73.4/53
99.2/99
99.2/100
RIFPA_jg23684_t1
RIFPA_jg23684_t2
b
RIFPA_jg16544_t1
RIFPA_jg16544_t2
Riftia_CAD29159
Lamellibrachia_sp_XB_2003_AAP04529
LAMLU_FUN_ALO75569
99.9/100
0.4
B
LINKER CHAINS
CHAIN L3
CHAIN L2
CHAIN L3
RIFPA_jg16545.t1
RIFPA_jg16545.t3
RIFPA_jg16545.t2
LAMLU_FUN_026443
Lamellibrachia_satsuma_BAU46569
100/100
a
92.3/89
RIFPA_jg16546.t1
RIFPA_jg16546.t2
RIFPA_jgCAJ00870
LAMLU_FUN_026441
Lamellibrachia_QAT80834
Lamellibrachia_sp_P16222
Lamellibrachia_satsuma_BAU46567
99.9/100
39.9/61
a
RIFPA_jg29472.t1
RIFPA_jg29472.t4
LAMLU_FUN_001067
97/95
99.9/100
b
100/100
RIFPA_jg8377.t1
RIFPA_jg8377.t2
c
RIFPA_jg12849.t1
99.9/98
RIFPA_jg19474.t1
Riftia_ABW24414
RIFPA_jg19474.t2
LAMLU_FUN_002344
LAMLU_FUN_QAT80835
LAMLU_FUN_024479
Lamellibrachia_satsuma_BAU46568
19.8/53
b
0.3
C
0.5
Chain L1
Chain L2
Chain L3

**Supplementary figure 28 | Phylogeny of Riftia and Lamellibrachia globin and linker genes**. **A,** Mid-rooted maximum-likelihood phylogenetic tree inference of the haemoglobin genes using 1000 ultrafast bootstrap replicates. Note the expansion of hemoglobin β1 genes in *Lamellibrachia* and *Riftia pachyptila*. **B,** Mid-rooted maximum-likelihood phylogenetic tree inference of the linker genes using 1000 ultrafast bootstrap replicates. **C** Unrooted maximum-likelihood phylogenetic tree inference of linker genes in a wide range of annelids and metazoans. *Riftia* linker genes belong to L3 and L2 family. *Lamellibrachia* gene identification is derived from the public annotated genome.

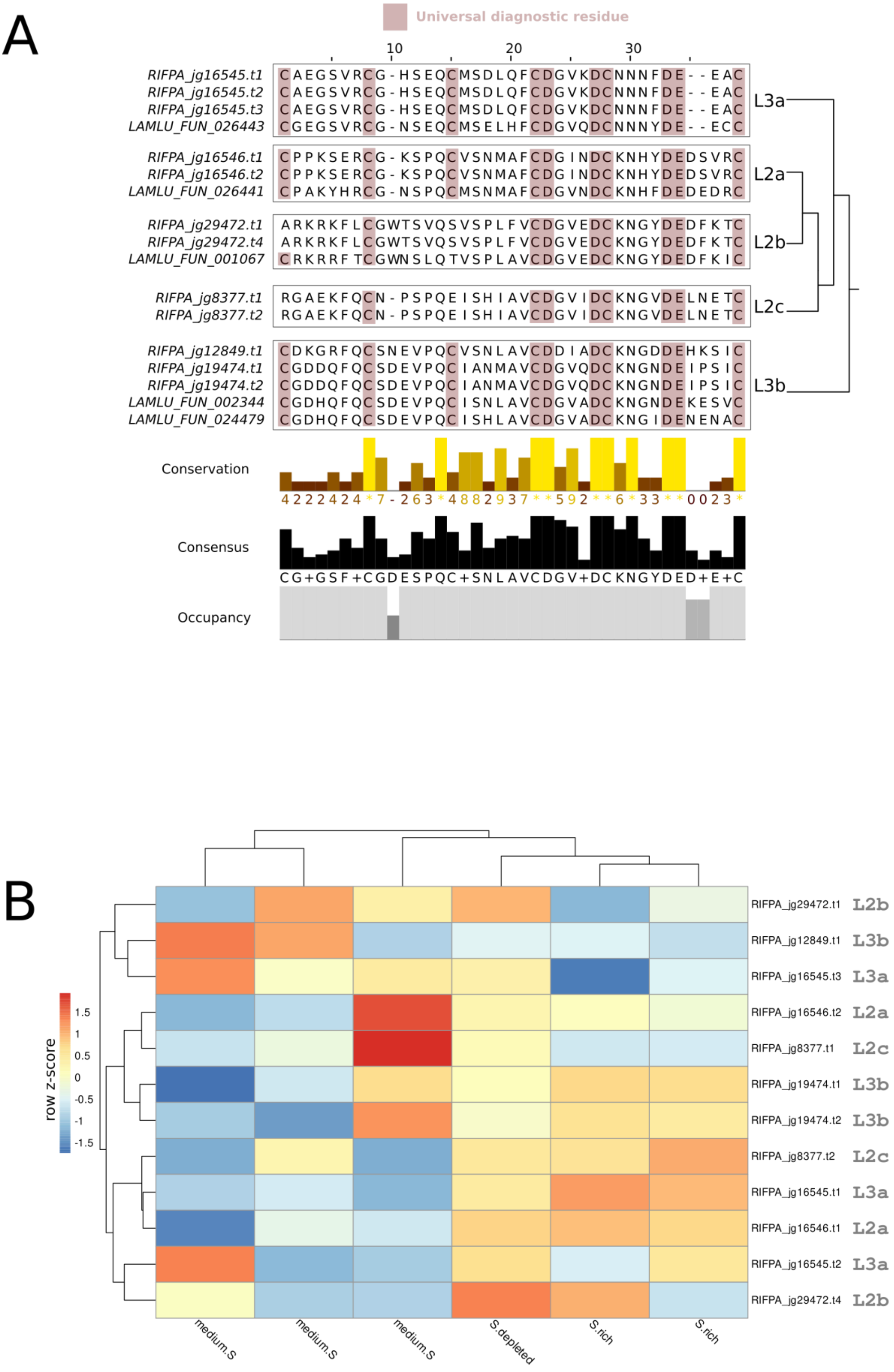

**Supplementary figure 29 | Multiple sequence alignments and gene expression of *Riftia* linker genes A,** Multiple sequence alignment, identification and characterisation of the signature diagnostic residues/motifs in the linker chains. Dark red boxes correspond to universal diagnostic residues.

Histogram plots correspond to conservation and occupancy of each multiple sequence alignment column. **B,** Expression profile of linker genes in the trophosome *Riftia pachyptila*. Medium-Sulphide, Sulphide-Depleted and Sulphide-rich correspond to different environmental conditions. Colour coding reflects the expression patterns based on row Z-score calculations. Linker genes are constitutively expressed in all trophosome tissues, irrespective of the experimental conditions.

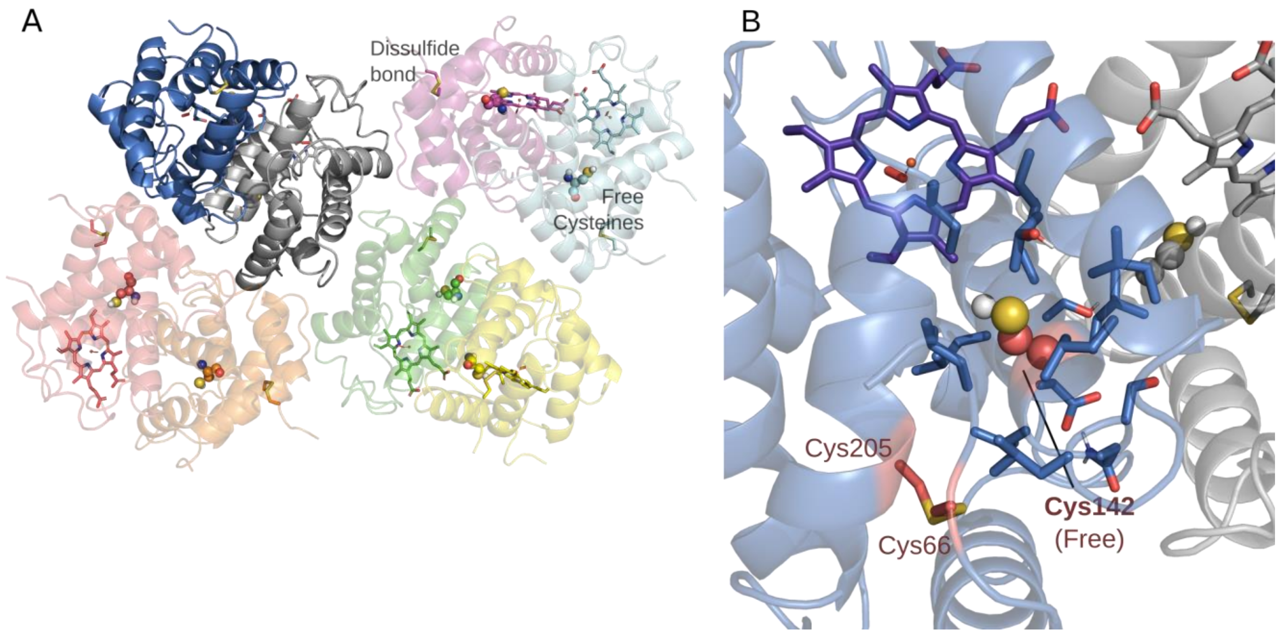

**Supplementary figure 30| Homology model generation for Riftia haemoglobin. A,** Haemoglobin octomeric structure of *Riftia* gene RIFPA_jg20259.t3 belonging to the β1c-Hb paralog group. **B,** Schematic representation of the putative free cysteine residue, hypothesized to bind toxic hydrogen sulphide ($H_2S$).

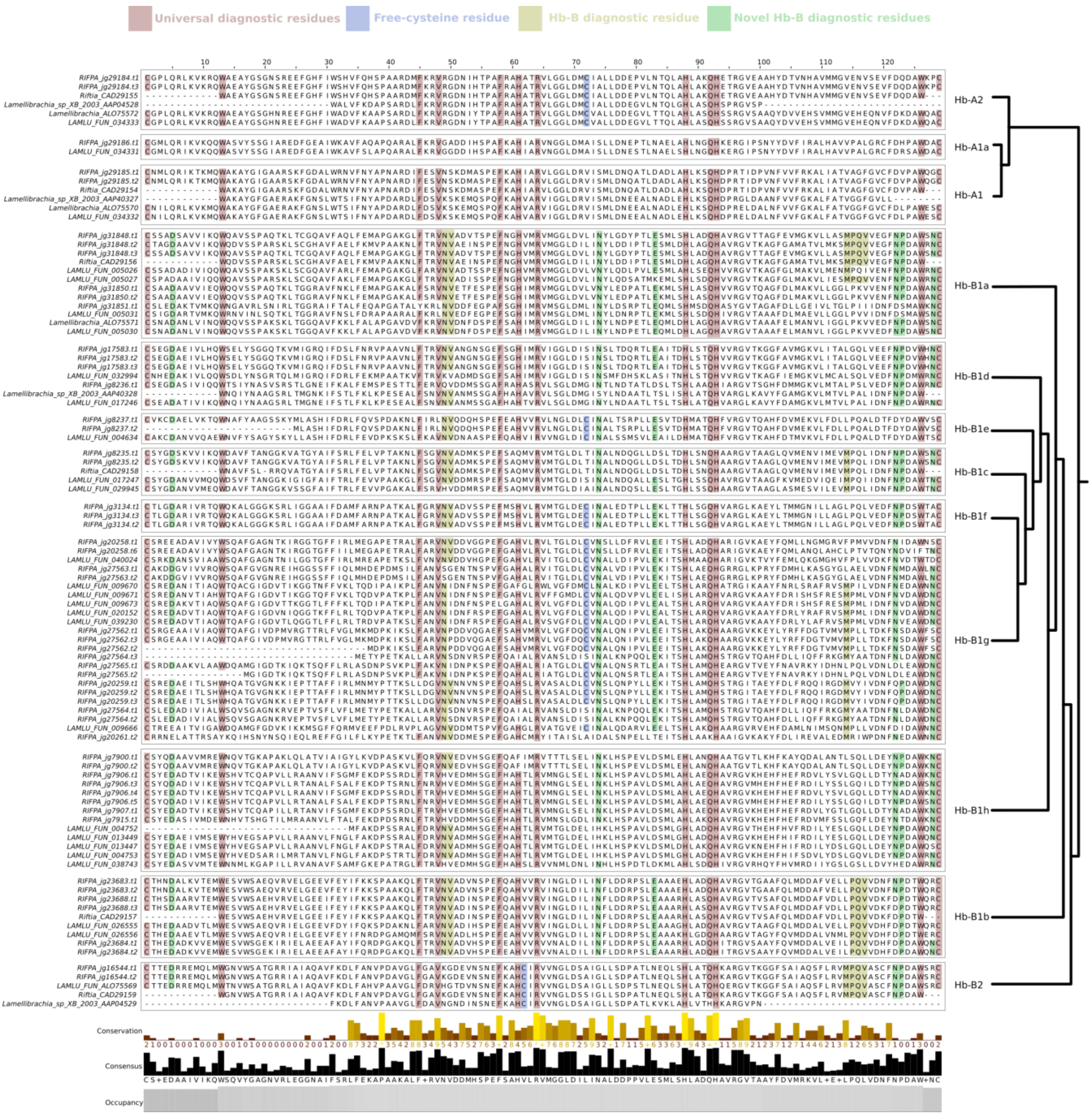

**Supplementary figure 31 | Multiple sequence alignment of *Riftia* and *Lamellibrachia* haemoglobin genes.** Multiple sequence alignment, identification, and characterisation of the signature diagnostic residues/motifs in the haemoglobin chains. Dark red boxes correspond to universal diagnostic residues; light purple to putative free cysteine residues; light yellow HB-β diagnostic residues; and finally, light green correspond to novel β chain residues identified in this study. Histogram plots correspond to conservation and occupancy of each multiple sequence alignment column. Putative free cysteine residues were identified in many newly described β1 genes.

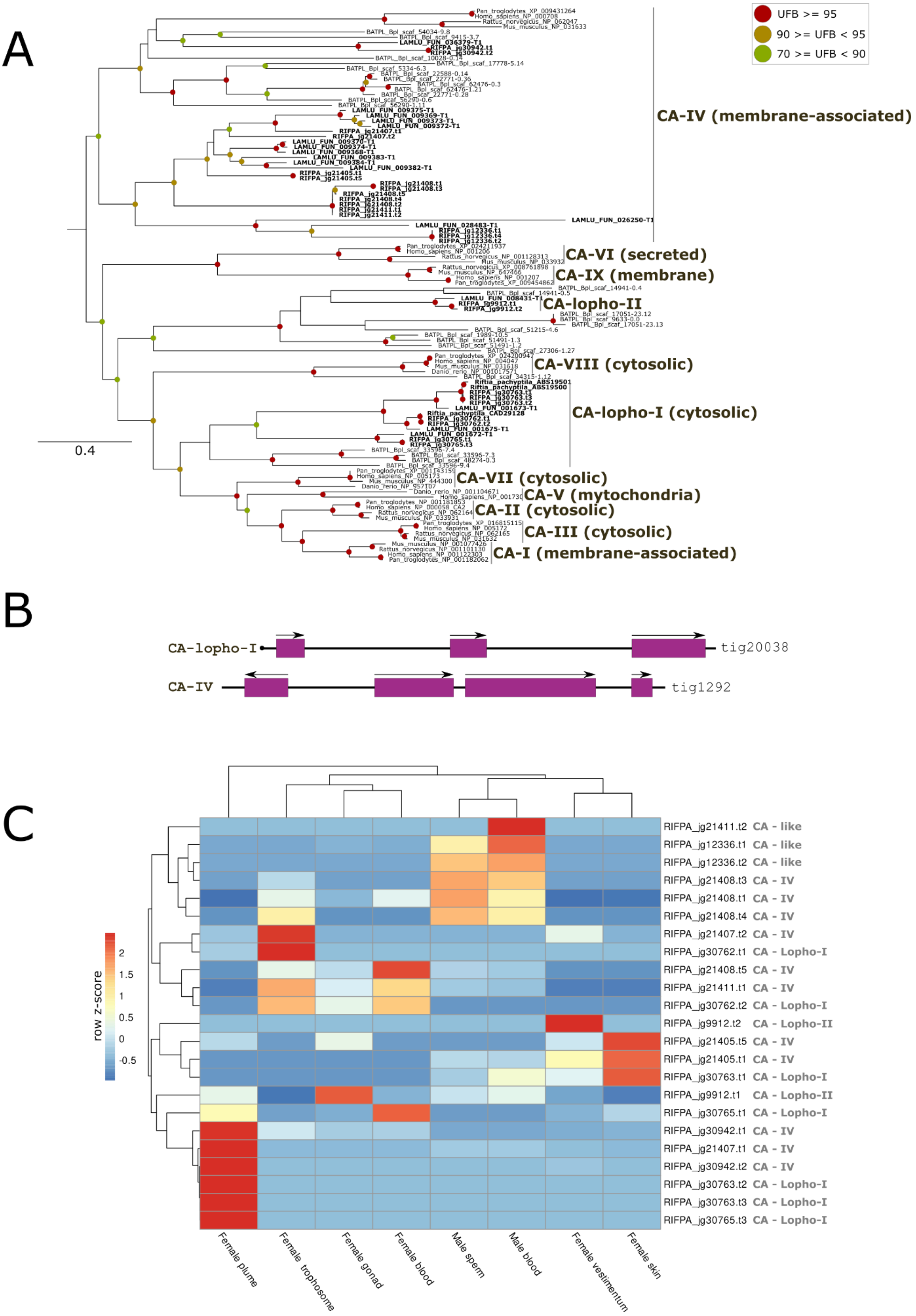

**Supplementary figure 32 | Carbonic anhydrase genes in *Riftia pachyptila*. A**, Mid-rooted maximum-likelihood phylogenetic tree inference of the carbonic anhydrase genes using 1000 ultrafast bootstrap replicates. Red circles represent ultrafast bootstrap values >= 95. Yellow circles represent ultrafast bootstrap values >= 90 and < 95. Green circles represent ultrafast bootstrap values < 90 and >= 70. Ultrafast bootstrap values smaller than 70 are not shown. Gene identification is derived from the public annotated genome. **B,** Genomic organisation of seven CA gene in the giant tubeworm genome. **C,** Expression profile of carbonic anhydrase genes in eight different adult tissues of *Riftia pachyptila.*

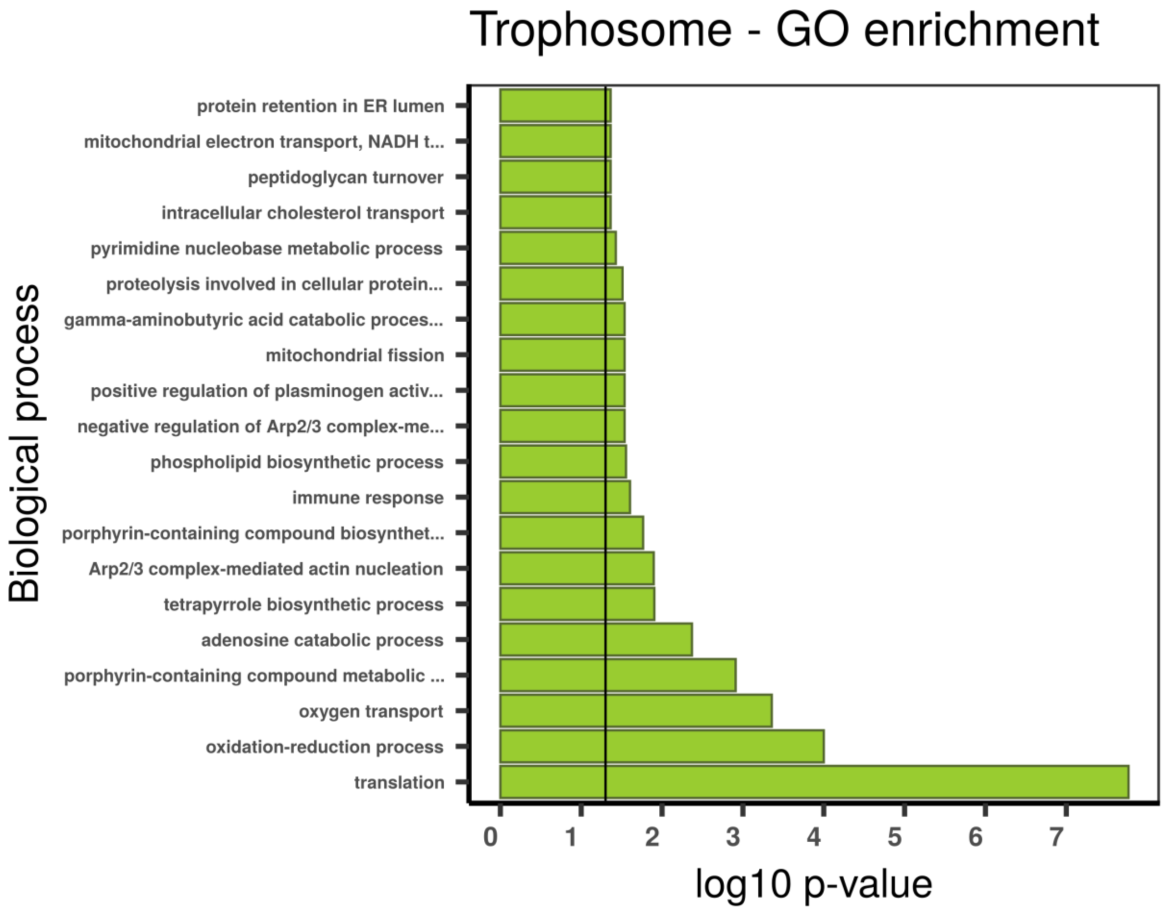

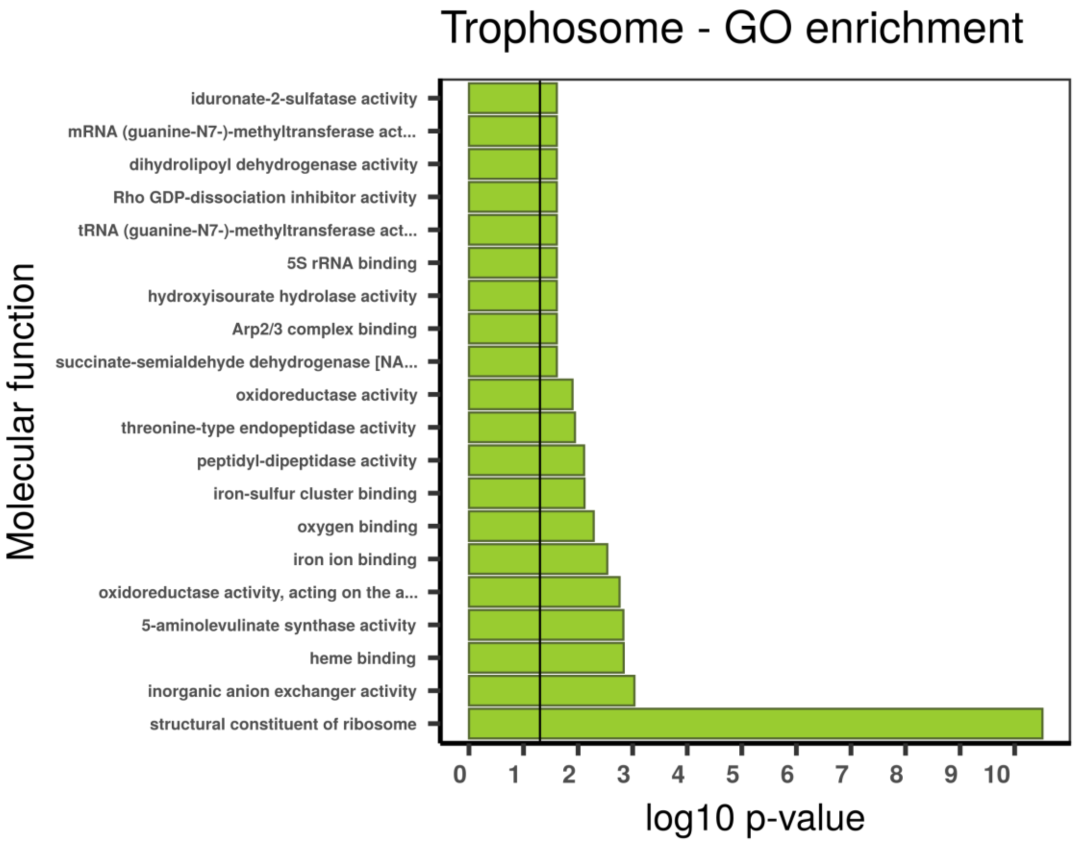
Trophosome - GO enrichment
Molecular function
iduronate-2-sulfatase activity
mRNA (guanine-N7-)-methyltransferase act...
dihydrolipoyl dehydrogenase activity
Rho GDP-dissociation inhibitor activity
tRNA (guanine-N7-)-methyltransferase act...
5S rRNA binding
hydroxyisourate hydrolase activity
Arp2/3 complex binding
succinate-semialdehyde dehydrogenase [NA...
oxidoreductase activity
threonine-type endopeptidase activity
peptidyl-dipeptidase activity
iron-sulfur cluster binding
oxygen binding
iron ion binding
oxidoreductase activity, acting on the a...
5-aminolevulinate synthase activity
heme binding
inorganic anion exchanger activity
structural constituent of ribosome
0 1 2 3 4 5 6 7 8 9 10
log10 p-value

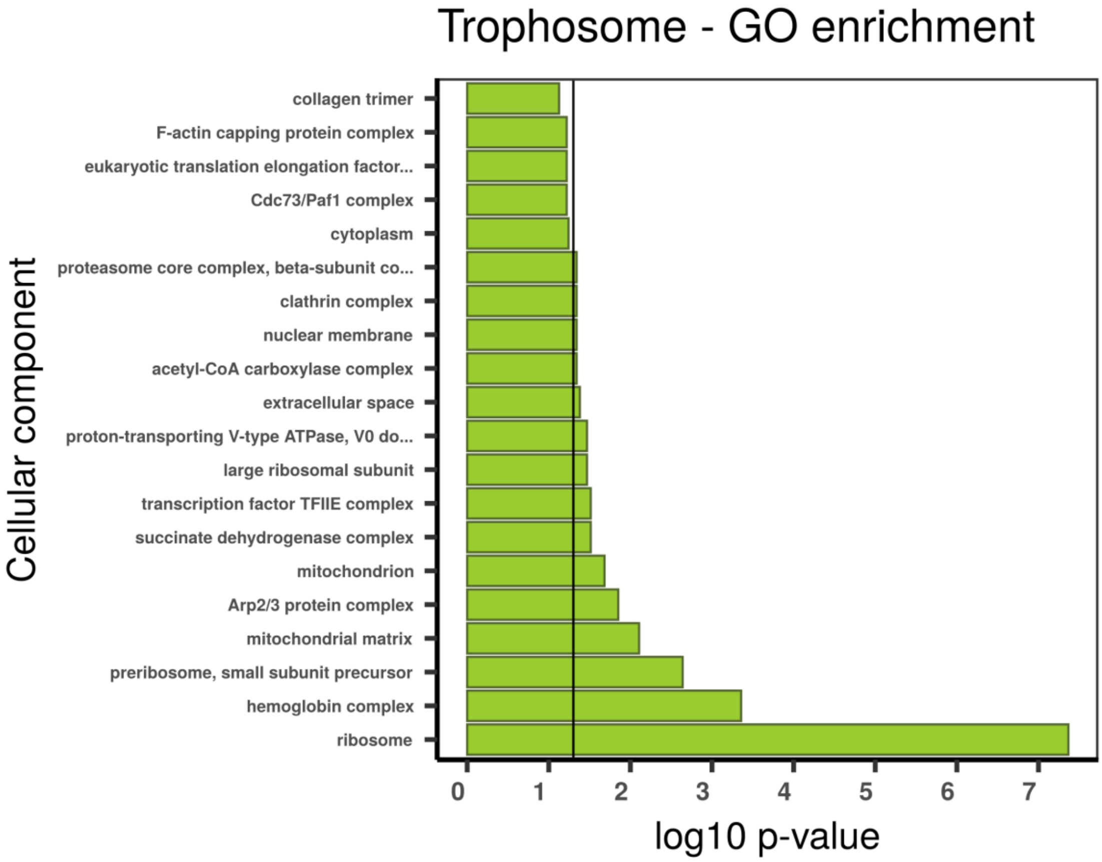
Trophosome - GO enrichment
Cellular component
collagen trimer
F-actin capping protein complex
eukaryotic translation elongation factor...
Cdc73/Paf1 complex
cytoplasm
proteasome core complex, beta-subunit co...
clathrin complex
nuclear membrane
acetyl-CoA carboxylase complex
extracellular space
proton-transporting V-type ATPase, V0 do...
large ribosomal subunit
transcription factor TFIIE complex
succinate dehydrogenase complex
mitochondrion
Arp2/3 protein complex
mitochondrial matrix
preribosome, small subunit precursor
hemoglobin complex
ribosome
0 1 2 3 4 5 6 7
log10 p-value

**Supplementary figure 33 | Gene set enrichment analysis with topGO using absolutely trophosome specific TAU genes.** Gene ontology (GO) enrichment analyses for absolutely trophosome specific TAU genes. The graphs correspond to the three domains of ontologies: biological process (BP), molecular function (MF) and cellular component (CC). The selected genes were analysed for enrichment in specific GO categories using the TopGO program against the background (all coding sequence genes). Y axis corresponds to enriched GO terms found in the respective domains (BP, MF and CC). X axis correspond to the log function of Fisher p-values obtained for each one of the enriched terms. The back line denotes a p-value = 0.05. P-values greater than 1,30 (log 0,05) indicate statistically significant enriched term. Genes involved chitin metabolism are differentially expressed in the body wall (skin).

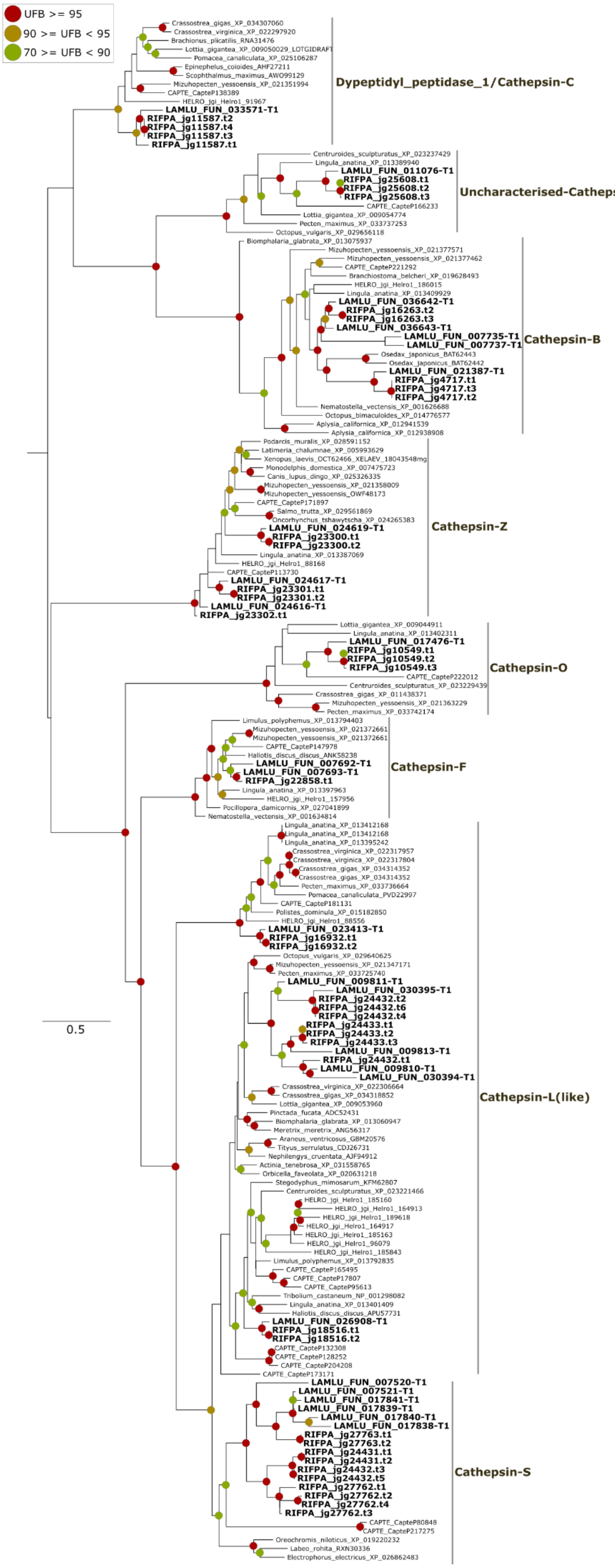

UFB >= 95
90 >= UFB < 95
70 >= UFB < 90
Crassostrea_gigas_XP_034307060
Crassostrea_virginica_XP_022297920
Brachionus_plicatilis_RNA31476
Lottia_gigantea_XP_009050029_LOTGIDRAFT
Pomacea_canaliculata_XP_025106287
Epinephelus_coioides_AHF27211
Scophthalmus_maximus_AWO99129
Mizuhopecten_yessoensis_XP_021351994
CAPTE_CapteP138389
HELRO_jgi_Helro1_91967
LAMLU_FUN_033571-T1
RIFPA_jg11587.t2
RIFPA_jg11587.t4
RIFPA_jg11587.t3
RIFPA_jg11587.t1
Dypeptidyl_peptidase_1/Cathepsin-C
Centruroides_sculpturatus_XP_023237429
Lingula_anatina_XP_013389940
LAMLU_FUN_011076-T1
RIFPA_jg25608.t1
RIFPA_jg25608.t2
RIFPA_jg25608.t3
CAPTE_CapteP166233
Lottia_gigantea_XP_009054774
Pecten_maximus_XP_033737253
Octopus_vulgaris_XP_029656118
Uncharacterised-Cathepsin
Biomphalaria_glabrata_XP_013075937
Mizuhopecten_yessoensis_XP_021377571
Mizuhopecten_yessoensis_XP_021377462
CAPTE_CapteP221292
Branchiostoma_belcheri_XP_019628493
HELRO_jgi_Helro1_186015
Lingula_anatina_XP_013409929
LAMLU_FUN_036642-T1
RIFPA_jg16263.t2
RIFPA_jg16263.t3
LAMLU_FUN_036643-T1
LAMLU_FUN_007735-T1
LAMLU_FUN_007737-T1
Cathepsin-B
Osedax_japonicus_BAT62443
Osedax_japonicus_BAT62442
LAMLU_FUN_021387-T1
RIFPA_jg4717.t1
RIFPA_jg4717.t3
RIFPA_jg4717.t2
Nematostella_vectensis_XP_001626688
Octopus_bimaculoides_XP_014776577
Aplysia_californica_XP_012941539
Aplysia_californica_XP_012938908
Podarcis_muralis_XP_028591152
Latimeria_chalumnae_XP_005993629
Xenopus_laevis_OCT62466_XELAEV_18043548mg
Monodelphis_domestica_XP_007475723
Canis_lupus_dingo_XP_025326335
Mizuhopecten_yessoensis_XP_021358009
Mizuhopecten_yessoensis_OWF48173
CAPTE_CapteP171897
Salmo_trutta_XP_029561869
Oncorhynchus_tshawytscha_XP_024265383
LAMLU_FUN_024619-T1
RIFPA_jg23300.t1
RIFPA_jg23300.t2
Lingula_anatina_XP_013387069
HELRO_jgi_Helro1_88168
CAPTE_CapteP113730
LAMLU_FUN_024617-T1
RIFPA_jg23301.t1
RIFPA_jg23301.t2
LAMLU_FUN_024616-T1
RIFPA_jg23302.t1
Cathepsin-Z
Lottia_gigantea_XP_009044911
Lingula_anatina_XP_013402311
LAMLU_FUN_017476-T1
RIFPA_jg10549.t1
RIFPA_jg10549.t2
RIFPA_jg10549.t3
CAPTE_CapteP222012
Centruroides_sculpturatus_XP_023229439
Crassostrea_gigas_XP_011438371
Mizuhopecten_yessoensis_XP_021363229
Pecten_maximus_XP_033742174
Cathepsin-O
Limulus_polyphemus_XP_013794403
Mizuhopecten_yessoensis_XP_021372661
Mizuhopecten_yessoensis_XP_021372661
CAPTE_CapteP147978
Haliotis_discus_discus_ANK58238
LAMLU_FUN_007692-T1
LAMLU_FUN_007693-T1
RIFPA_jg22858.t1
Lingula_anatina_XP_013397963
HELRO_jgi_Helro1_157956
Pocillopora_damicornis_XP_027041899
Nematostella_vectensis_XP_001634814
Cathepsin-F
Lingula_anatina_XP_013412168
Lingula_anatina_XP_013412168
Lingula_anatina_XP_013395242
Crassostrea_virginica_XP_022317957
Crassostrea_virginica_XP_022317804
Crassostrea_gigas_XP_034314352
Crassostrea_gigas_XP_034314352
Pecten_maximus_XP_033736664
Pomacea_canaliculata_PVD22997
CAPTE_CapteP181131
Polistes_dominula_XP_015182850
HELRO_jgi_Helro1_88556
LAMLU_FUN_023413-T1
RIFPA_jg16932.t1
RIFPA_jg16932.t2
Octopus_vulgaris_XP_029640625
Mizuhopecten_yessoensis_XP_021347171
Pecten_maximus_XP_033725740
LAMLU_FUN_009811-T1
LAMLU_FUN_030395-T1
RIFPA_jg24432.t2
RIFPA_jg24432.t6
RIFPA_jg24432.t4
RIFPA_jg24433.t1
RIFPA_jg24433.t2
RIFPA_jg24433.t3
LAMLU_FUN_009813-T1
RIFPA_jg24432.t1
LAMLU_FUN_009810-T1
LAMLU_FUN_030394-T1
Crassostrea_virginica_XP_022306664
Crassostrea_gigas_XP_034318852
Lottia_gigantea_XP_009053960
Pinctada_fucata_ADC52431
Biomphalaria_glabrata_XP_013060947
Meretrix_meretrix_ANG56317
Araneus_ventricosus_GBM20576
Tityus_serrulatus_CDJ26731
Nephilengys_cruentata_AJF94912
Actinia_tenebrosa_XP_031558765
Orbicella_faveolata_XP_020631218
Stegodyphus_mimosarum_KFM62807
Centruroides_sculpturatus_XP_023221466
HELRO_jgi_Helro1_185160
HELRO_jgi_Helro1_164913
HELRO_jgi_Helro1_189618
HELRO_jgi_Helro1_164917
HELRO_jgi_Helro1_185163
HELRO_jgi_Helro1_96079
HELRO_jgi_Helro1_185843
Limulus_polyphemus_XP_013792835
CAPTE_CapteP165495
CAPTE_CapteP17807
CAPTE_CapteP95613
Tribolium_castaneum_NP_001298082
Lingula_anatina_XP_013401409
Haliotis_discus_discus_APU57731
LAMLU_FUN_026908-T1
RIFPA_jg18516.t1
RIFPA_jg18516.t2
CAPTE_CapteP132308
CAPTE_CapteP128252
CAPTE_CapteP204208
CAPTE_CapteP173171
Cathepsin-L(like)
LAMLU_FUN_007520-T1
LAMLU_FUN_007521-T1
LAMLU_FUN_017841-T1
LAMLU_FUN_017839-T1
LAMLU_FUN_017840-T1
LAMLU_FUN_017838-T1
RIFPA_jg27763.t1
RIFPA_jg27763.t2
RIFPA_jg24431.t1
RIFPA_jg24431.t2
RIFPA_jg24432.t3
RIFPA_jg24432.t5
RIFPA_jg27762.t1
RIFPA_jg27762.t2
RIFPA_jg27762.t4
RIFPA_jg27762.t3
CAPTE_CapteP80848
CAPTE_CapteP217275
Oreochromis_niloticus_XP_019220232
Labeo_rohita_RXN30336
Electrophorus_electricus_XP_026862483
Cathepsin-S
0.5

**Supplementary figure 34 | Phylogeny of cathepsins.** Mid-rooted maximum-likelihood phylogenetic tree inference of the cathepsin genes using 1000 ultrafast bootstrap replicates. The branch support values are represented by the coloured circles in the tree nodes. Red circles represent ultrafast bootstrap values >= 95. Yellow circles represent ultrafast bootstrap values >= 90 and < 95. Green circles represent ultrafast bootstrap values < 90 and >= 70. Ultrafast bootstrap values smaller than 70 are not shown. Accession numbers for NCBI database are displayed after the species names. *Capitella, Helobdella* and *Lamellibrachia* gene identification are derived from the publicly available annotated genomes.

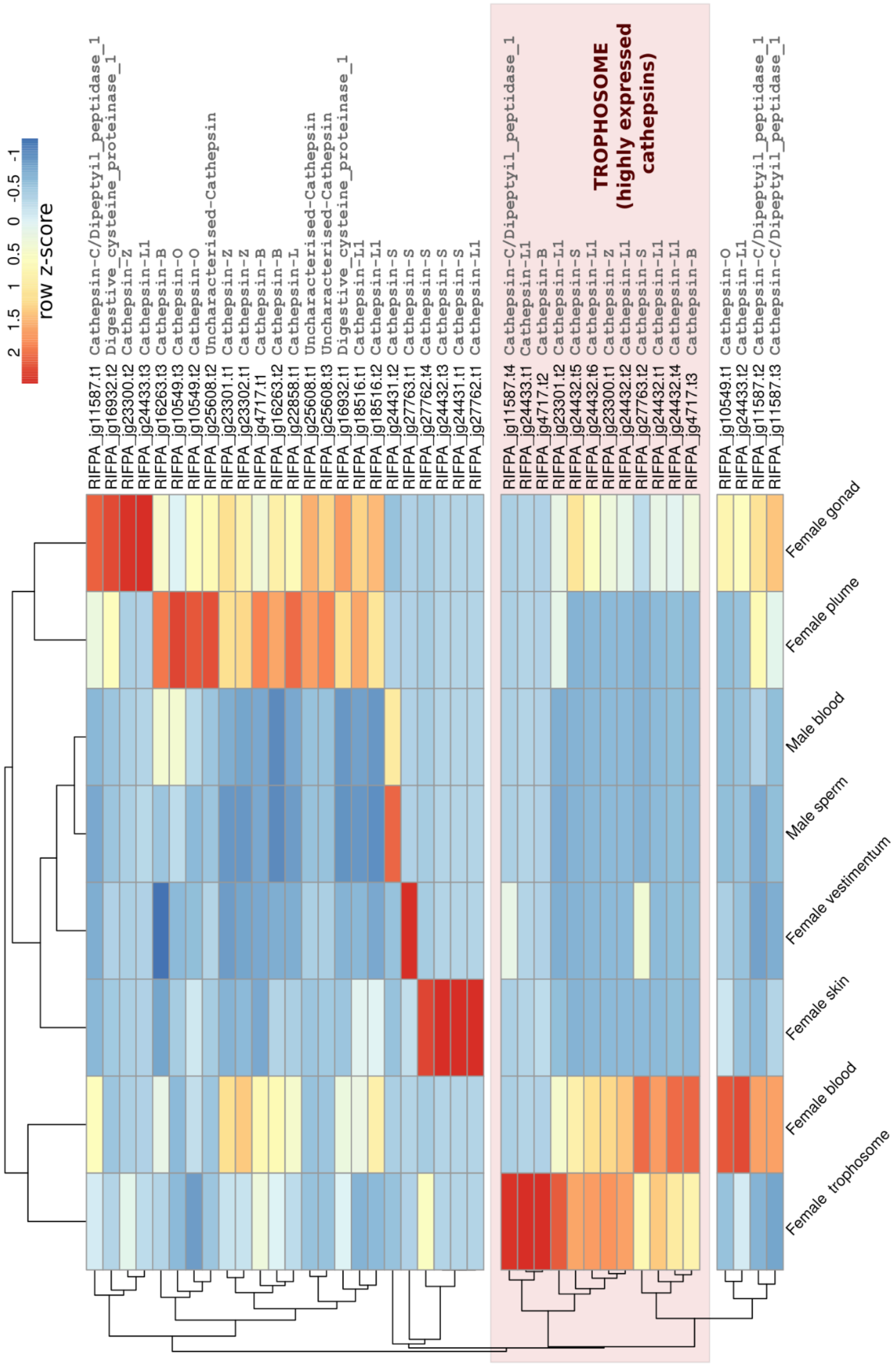

**Supplementary figure 35 | Gene expression of cathepsins.** Gene expression of cathepsin genes in eight adult tissues of *Riftia pachyptila*. Colour coding reflects the expression patterns based on row Z-score calculations. Cathepsins highly expressed in the trophosome are highlighted in light red.

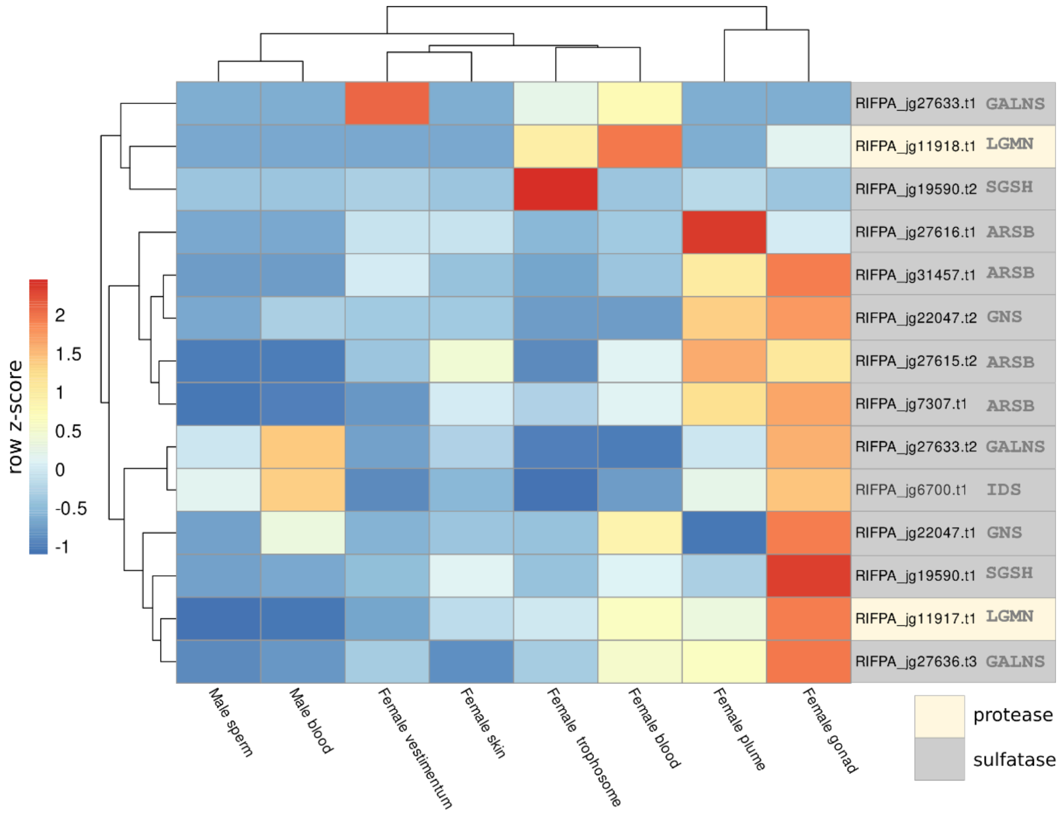

row z-score
2
1.5
1
0.5
0
-0.5
-1
RIFPA_jg27633.t1 GALNS
RIFPA_jg11918.t1 LGMN
RIFPA_jg19590.t2 SGSH
RIFPA_jg27616.t1 ARSB
RIFPA_jg31457.t1 ARSB
RIFPA_jg22047.t2 GNS
RIFPA_jg27615.t2 ARSB
RIFPA_jg7307.t1 ARSB
RIFPA_jg27633.t2 GALNS
RIFPA_jg6700.t1 IDS
RIFPA_jg22047.t1 GNS
RIFPA_jg19590.t1 SGSH
RIFPA_jg11917.t1 LGMN
RIFPA_jg27636.t3 GALNS
Male sperm
Male blood
Female vestimentum
Female skin
Female trophosome
Female blood
Female plume
Female gonad
protease
sulfatase

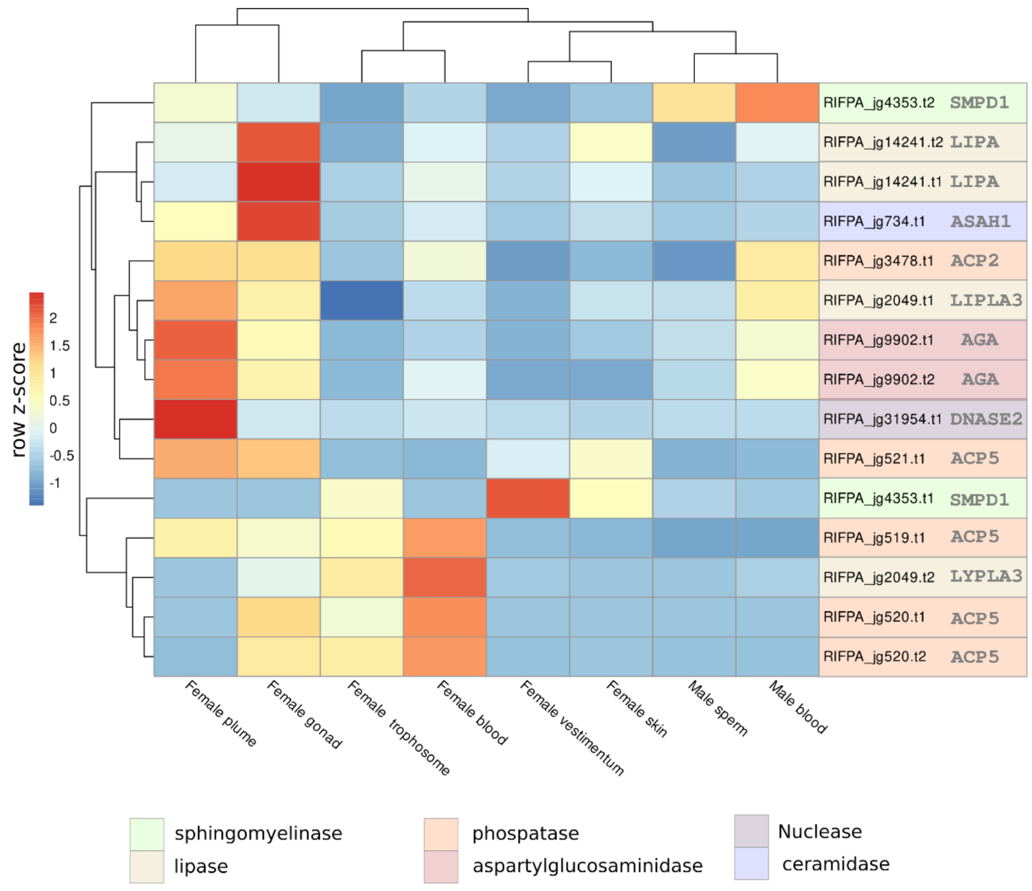

row z-score
2
1.5
1
0.5
0
-0.5
-1
RIFPA_jg4353.t2 SMPD1
RIFPA_jg14241.t2 LIPA
RIFPA_jg14241.t1 LIPA
RIFPA_jg734.t1 ASAH1
RIFPA_jg3478.t1 ACP2
RIFPA_jg2049.t1 LIPLA3
RIFPA_jg9902.t1 AGA
RIFPA_jg9902.t2 AGA
RIFPA_jg31954.t1 DNASE2
RIFPA_jg521.t1 ACP5
RIFPA_jg4353.t1 SMPD1
RIFPA_jg519.t1 ACP5
RIFPA_jg2049.t2 LYPLA3
RIFPA_jg520.t1 ACP5
RIFPA_jg520.t2 ACP5
Female plume
Female gonad
Female trophosome
Female blood
Female vestimentum
Female skin
Male sperm
Male blood
sphingomyelinase
lipase
phospatase
aspartylglucosaminidase
Nuclease
ceramidase

**Supplementary figure 36 | Gene expression of lysosomal-associated hydrolases.** Gene expression of lysosomal-associated hydrolase genes based on KEGG results. Genes identified on the KEGG orthology lysosome pathway (Lysosome 04142 – PATH:ko04142) were used to calculate the gene expression across the adult tubeworm tissues. Colour coding reflects the expression patterns based on row Z-score calculations.

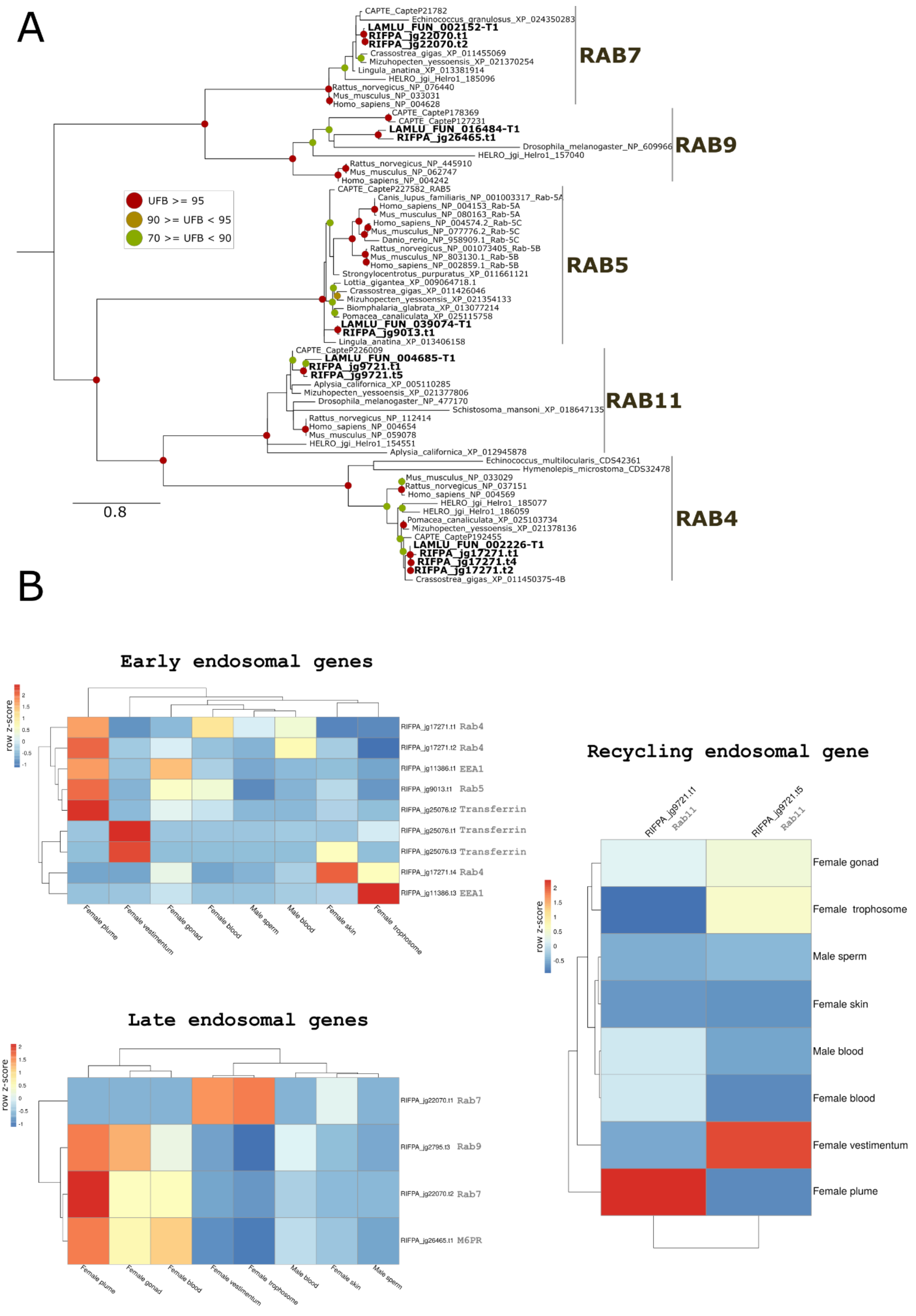

**Supplementary figure 37 | Phylogeny and gene expression of endosomal genes**. A, Mid-rooted maximum-likelihood phylogenetic tree inference of the endosomal genes using 1000 ultrafast bootstrap replicates. The branch support values are represented by the coloured circles in the tree nodes. Red circles represent ultrafast bootstrap values >= 95. Yellow circles represent ultrafast bootstrap values >= 90 and < 95. Green circles represent ultrafast bootstrap values < 90 and >= 70. Ultrafast bootstrap values smaller than 70 are not shown. Accession numbers for NCBI database are displayed after the species names. *Capitella*, *Helobdella* and *Lamellibrachia* gene identification are derived from the publicly available annotated genomes. B, Expression profile of early, late and recycling endosomal genes. Colour coding reflects the expression patterns based on row Z-score calculations.

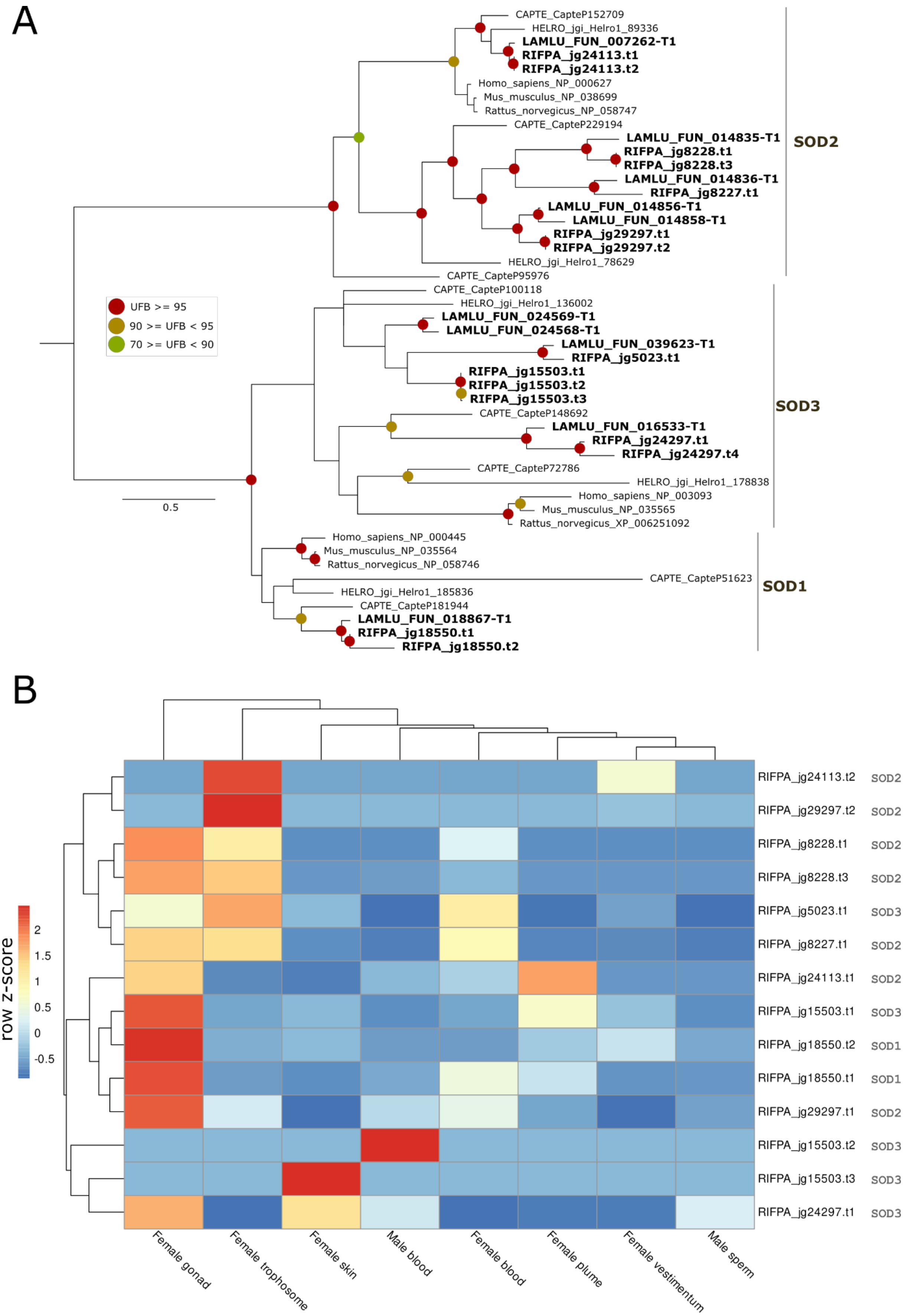

**Supplementary figure 38 | Phylogeny and gene expression of SOD genes**. **A.** Mid-rooted maximum-likelihood phylogenetic tree inference of the SOD genes using 1000 ultrafast bootstrap replicates. The branch support values are represented by the coloured circles in the tree nodes. Red circles represent ultrafast bootstrap values >= 95. Yellow circles represent ultrafast bootstrap values >= 90 and < 95. Green circles represent ultrafast bootstrap values < 90 and >= 70. Ultrafast bootstrap values smaller than 70 are not shown. Accession numbers for NCBI database are displayed after the species names. *Capitella, Helobdella* and *Lamellibrachia* gene identification are derived from the publicly available annotated genomes. **B,** Expression profile of SOD genes. Colour coding reflects the expression patterns based on row Z-score calculations. SOD genes are highly expressed in the trophosome indicative of oxidative stress.

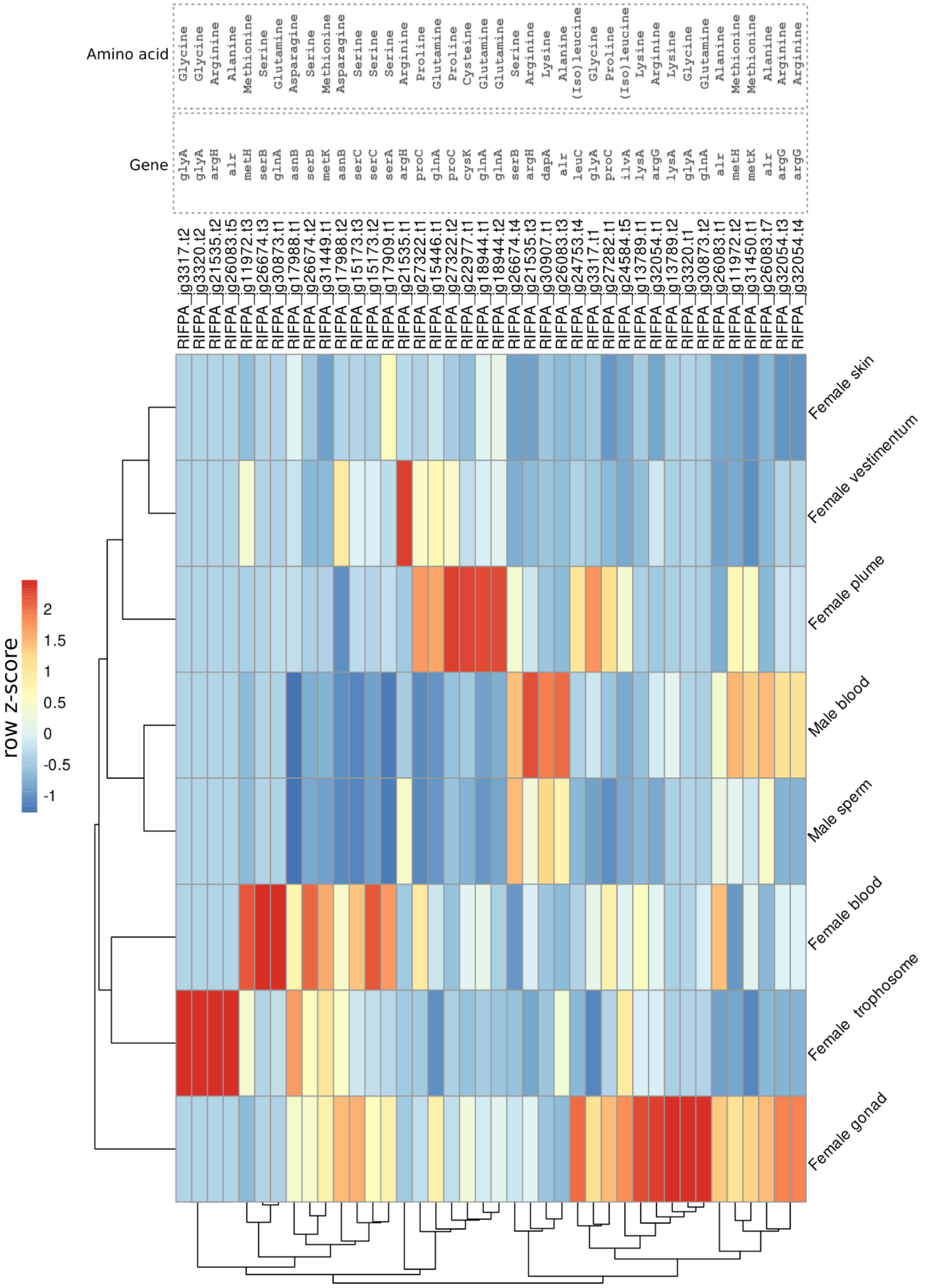

**Supplementary figure 39 | Gene expression of enzymes related to amino acid biosynthesis.** Gene expression of key enzymes related to amino acid biosynthesis genes based on KEGG results (check Supplementary Table 10). Colour coding reflects the expression patterns based on row Z-score calculations. Enzymes related to arginine and glycine metabolism are highly expressed in the trophosome.

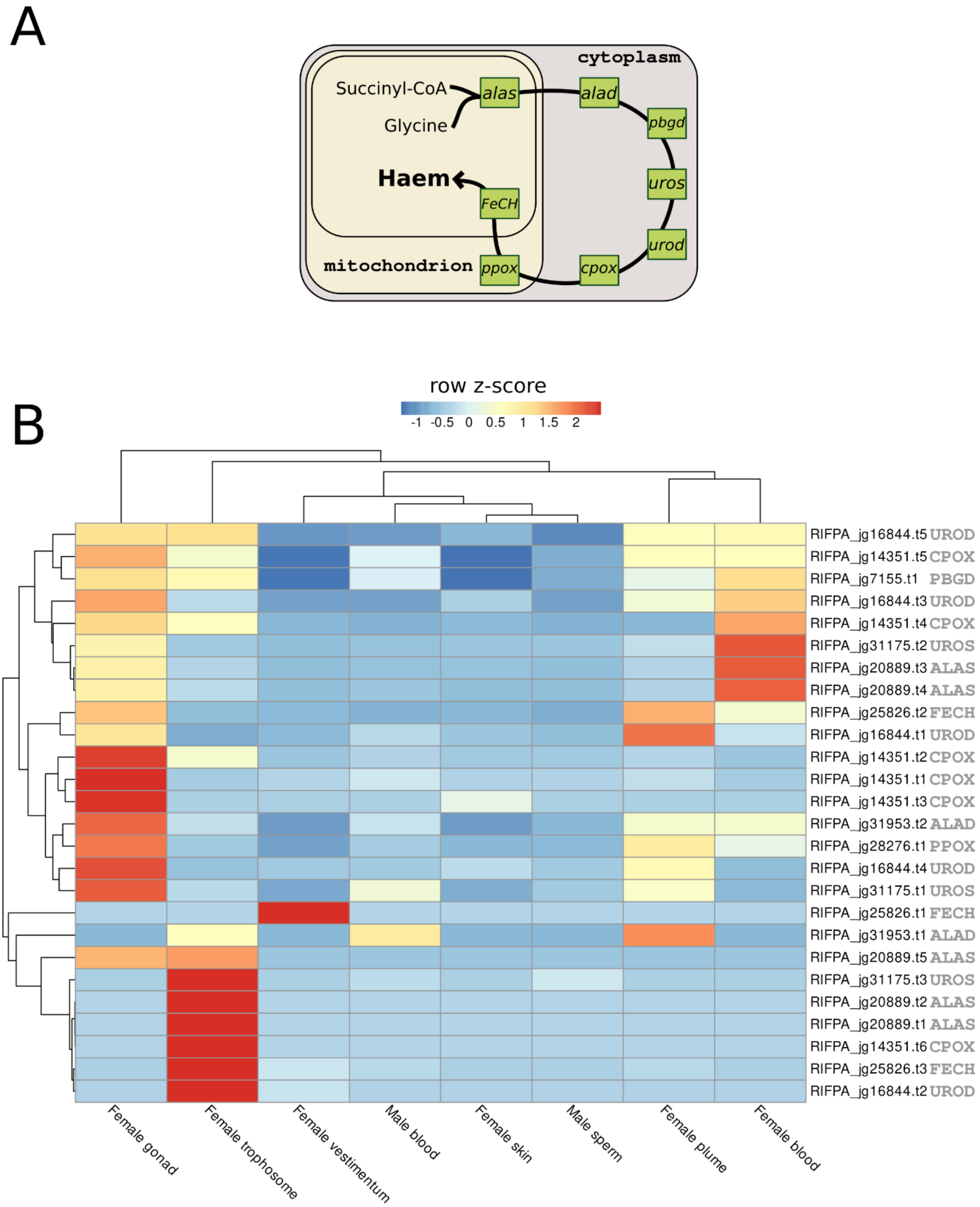

**Supplementary figure 40 | Gene expression of enzymes related to haem biosynthesis. A**, Scheme representing the seven conserved enzymes required for haem synthesis (adapted from Kořený et al., 2013). All enzymes were identified in the giant tubeworm genome. **B,** Gene expression of key enzymes related haem synthesis. Colour coding reflects the expression patterns based on row Z-score calculations. Enzymes related to haem synthesis are highly expressed in the trophosome suggesting that this tissue is involved with haematopoiesis in *Riftia*.

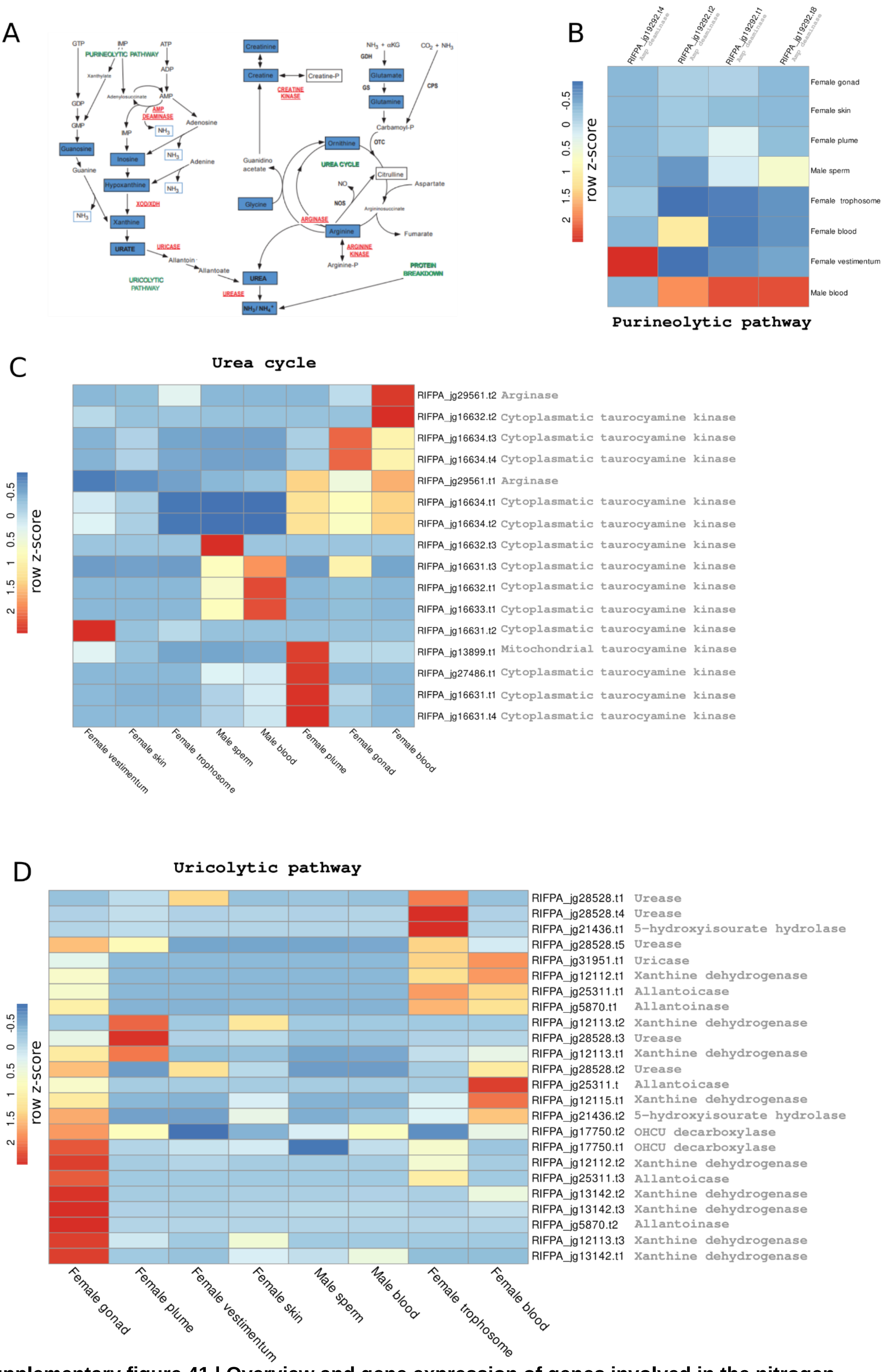

**Supplementary figure 41 | Overview and gene expression of genes involved in the nitrogen metabolism in *Riftia*. A,** Nitrogen metabolism and pathways leading to the main nitrogeneous endproducts (obtained from Cian et al., 2000). **B – D,** Expression profile of different key enzymes related to the purineolytic, uricolytic and urea pathways. Colour coding reflects the expression patterns based on row Z-score calculations. Genes involved in the uricolytic pathway are highly expressed in the trophosome.

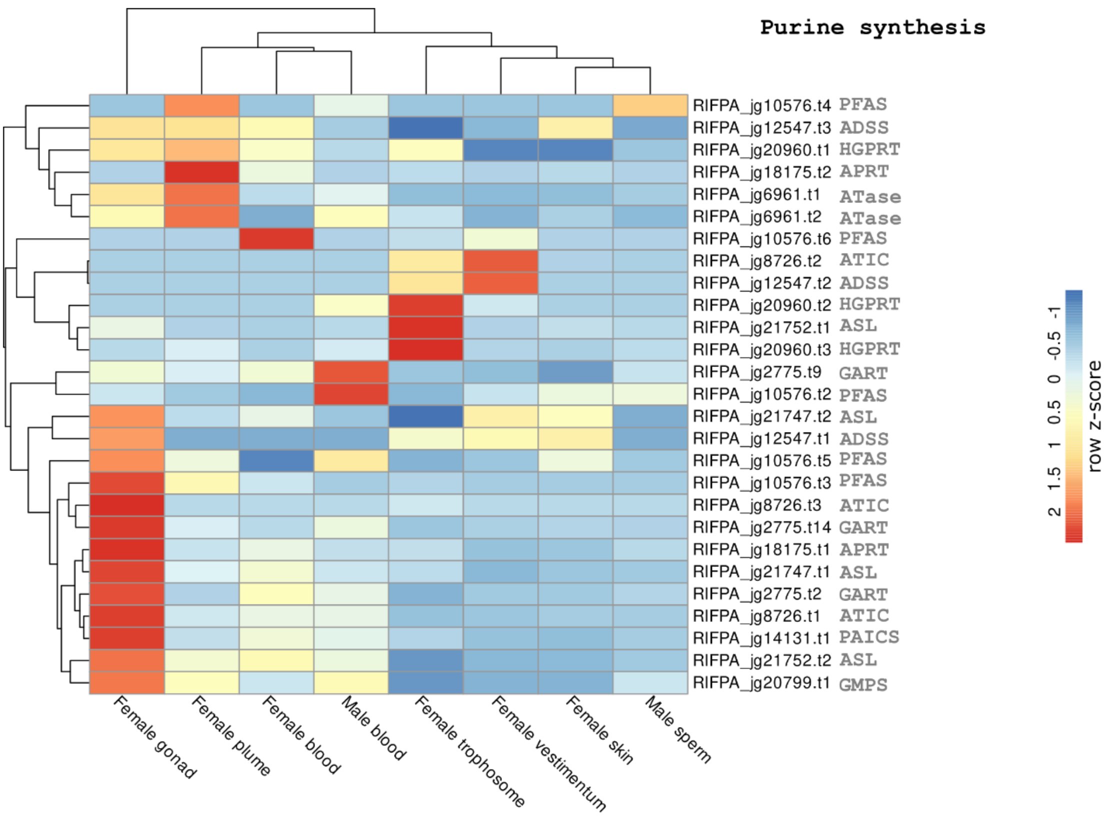

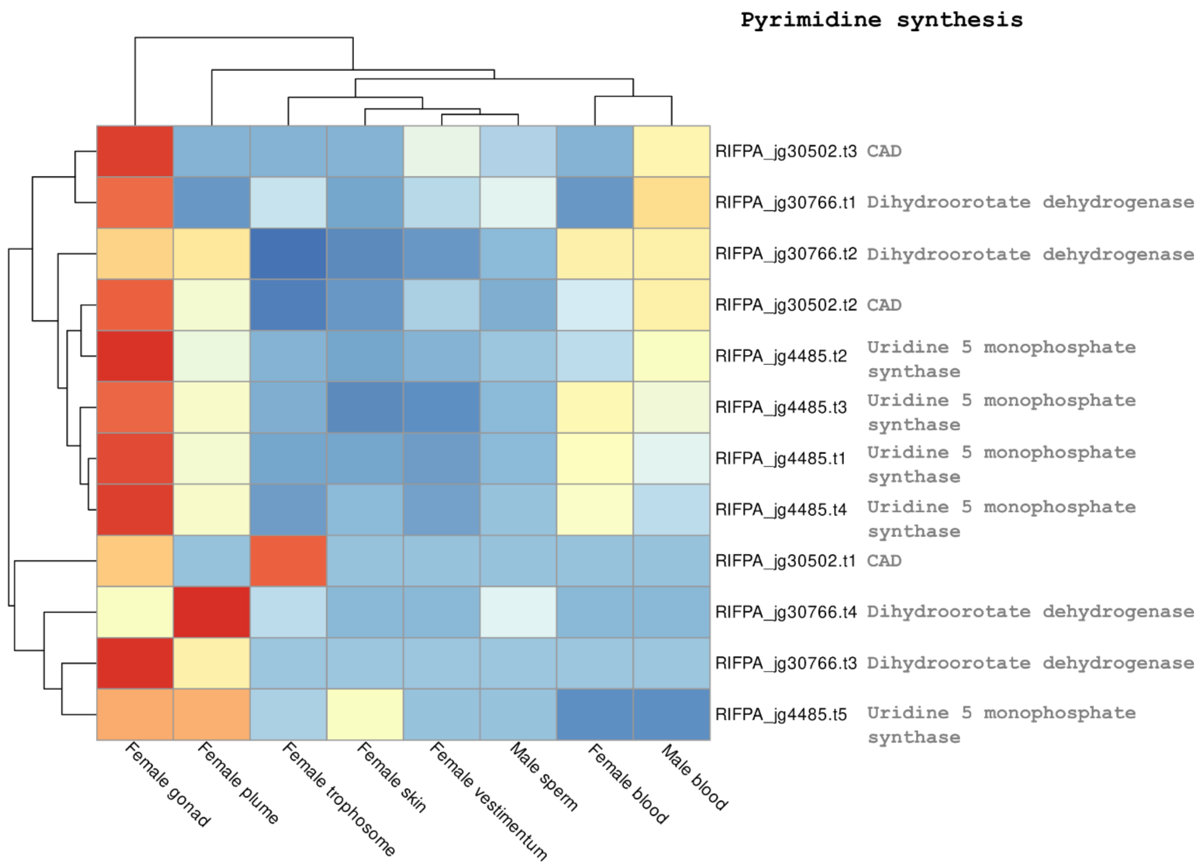

**Supplementary figure 42 | Gene expression of genes involved in the purine and pyrimidine pathways in *Riftia*.** Expression profile of different key enzymes related to the purine and pyrimidine biosynthesis pathways. Colour coding reflects the expression patterns based on row Z-score calculations. Abbreviations: PFAS: Phosphoribosylformylglycinamidine Synthase; ADSS: Adenylosuccinate Synthase; HGPRT: hypoxanthine phosphoribosyltransferase; APRT: adenine phosphoribosyltransferase; ATase: phosphoribosyl pyrophosphate amidotransferase; ATIC: 5-Aminoimidazole-4-Carboxamide Ribonucleotide Formyltransferase; ASL: Adenylosuccinate Lyase; ADSS: Adenylosuccinate Synthase; GART: Glycinamide Ribonucleotide Transformylase; PAICS: Phosphoribosylaminoimidazole Carboxylase And Phosphoribosylamino-imidazolesuccinocarboxamide Synthase; GMPS: Guanine Monophosphate Synthase; and CAD (Carbamoyl-Phosphate Synthetase 2, Aspartate Transcarbamylase, And Dihydroorotase)

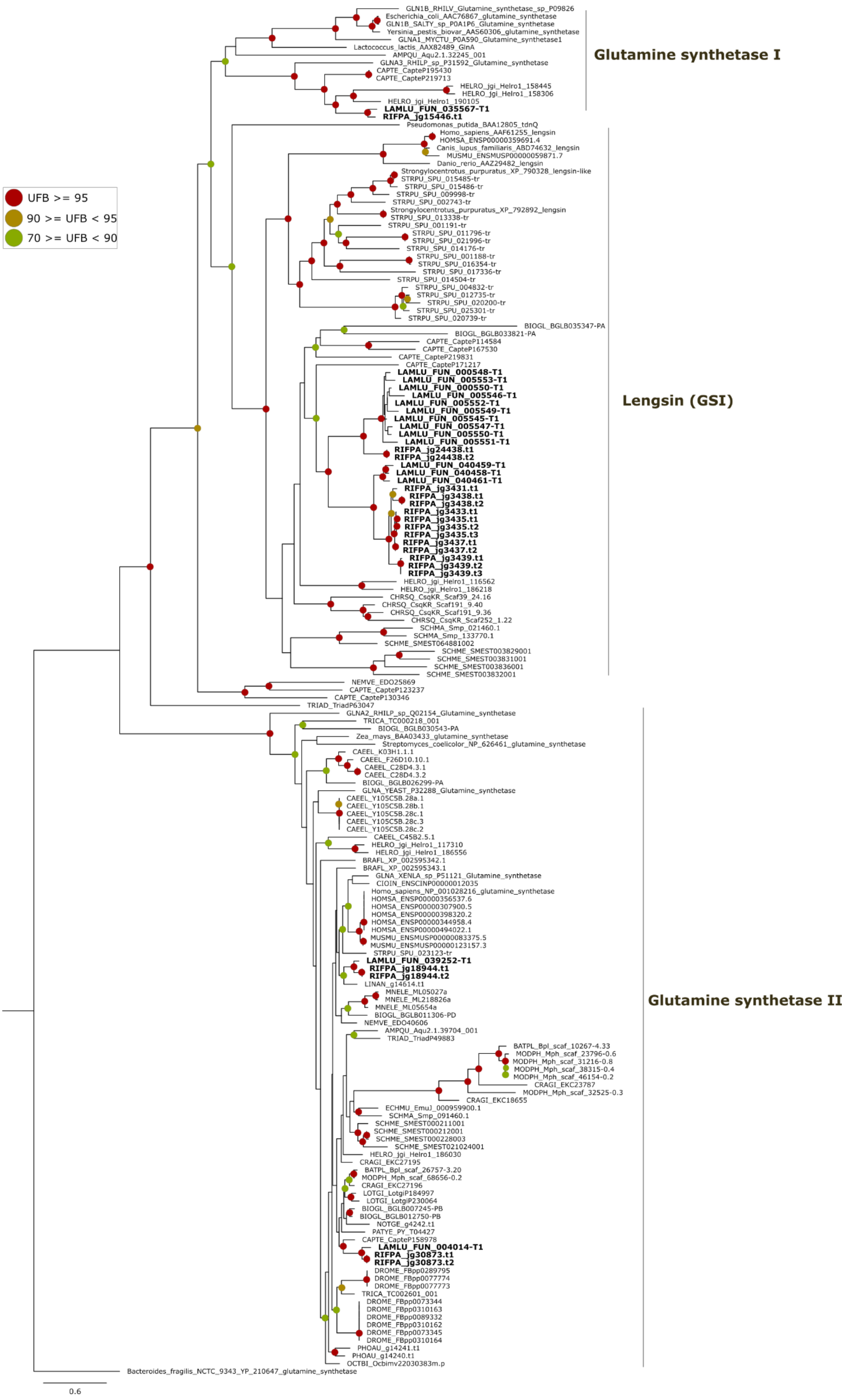

GLN1B_RHILV_Glutamine_synthetase_sp_P09826
Escherichia_coli_AAC76867_glutamine_synthetase
GLN1B_SALTY_sp_P0A1P6_Glutamine_synthetase
Yersinia_pestis_biovar_AAS60306_glutamine_synthetase
GLNA1_MYCTU_P0A590_Glutamine_synthetase1
Lactococcus_lactis_AAX82489_GlnA
AMPQU_Aqu2.1.32245_001
GLNA3_RHILP_sp_P31592_Glutamine_synthetase
CAPTE_CapteP195430
CAPTE_CapteP219713
HELRO_jgi_Helro1_158445
HELRO_jgi_Helro1_158306
HELRO_jgi_Helro1_190105
LAMLU_FUN_035567-T1
RIFPA_jg15446.t1
Glutamine synthetase I
Pseudomonas_putida_BAA12805_tdnQ
Homo_sapiens_AAF61255_lengsin
HOMSA_ENSP00000359691.4
Canis_lupus_familiaris_ABD74632_lengsin
MUSMU_ENSMUSP00000059871.7
Danio_rerio_AAZ29482_lengsin
Strongylocentrotus_purpuratus_XP_790328_lengsin-like
STRPU_SPU_015485-tr
STRPU_SPU_015486-tr
STRPU_SPU_009998-tr
STRPU_SPU_002743-tr
Strongylocentrotus_purpuratus_XP_792892_lengsin
STRPU_SPU_013338-tr
STRPU_SPU_001191-tr
STRPU_SPU_011796-tr
STRPU_SPU_021996-tr
STRPU_SPU_014176-tr
STRPU_SPU_001188-tr
STRPU_SPU_016354-tr
STRPU_SPU_017336-tr
STRPU_SPU_014504-tr
STRPU_SPU_004832-tr
STRPU_SPU_012735-tr
STRPU_SPU_020200-tr
STRPU_SPU_025301-tr
STRPU_SPU_020739-tr
BIOGL_BGLB035347-PA
BIOGL_BGLB033821-PA
CAPTE_CapteP114584
CAPTE_CapteP167530
CAPTE_CapteP219831
CAPTE_CapteP171217
LAMLU_FUN_000548-T1
LAMLU_FUN_005553-T1
LAMLU_FUN_000550-T1
LAMLU_FUN_005546-T1
LAMLU_FUN_005552-T1
LAMLU_FUN_005549-T1
LAMLU_FUN_005545-T1
LAMLU_FUN_005547-T1
LAMLU_FUN_005550-T1
LAMLU_FUN_005551-T1
RIFPA_jg24438.t1
RIFPA_jg24438.t2
LAMLU_FUN_040459-T1
LAMLU_FUN_040458-T1
LAMLU_FUN_040461-T1
RIFPA_jg3431.t1
RIFPA_jg3438.t1
RIFPA_jg3438.t2
RIFPA_jg3433.t1
RIFPA_jg3435.t1
RIFPA_jg3435.t2
RIFPA_jg3435.t3
RIFPA_jg3437.t1
RIFPA_jg3437.t2
RIFPA_jg3439.t1
RIFPA_jg3439.t2
RIFPA_jg3439.t3
HELRO_jgi_Helro1_116562
HELRO_jgi_Helro1_186218
CHRSQ_CsqKR_Scaf39_24.16
CHRSQ_CsqKR_Scaf191_9.40
CHRSQ_CsqKR_Scaf191_9.36
CHRSQ_CsqKR_Scaf252_1.22
SCHMA_Smp_021460.1
SCHMA_Smp_133770.1
SCHME_SMEST064881002
SCHME_SMEST003829001
SCHME_SMEST003831001
SCHME_SMEST003836001
SCHME_SMEST003832001
Lengsin (GSI)
NEMVE_EDO25869
CAPTE_CapteP123237
CAPTE_CapteP130346
TRIAD_TriadP63047
UFB >= 95
90 >= UFB < 95
70 >= UFB < 90
GLNA2_RHILP_sp_Q02154_Glutamine_synthetase
TRICA_TC000218_001
BIOGL_BGLB030543-PA
Zea_mays_BAA03433_glutamine_synthetase
Streptomyces_coelicolor_NP_626461_glutamine_synthetase
CAEEL_K03H1.1.1
CAEEL_F26D10.10.1
CAEEL_C28D4.3.1
CAEEL_C28D4.3.2
BIOGL_BGLB026299-PA
GLNA_YEAST_P32288_Glutamine_synthetase
CAEEL_Y105C5B.28a.1
CAEEL_Y105C5B.28b.1
CAEEL_Y105C5B.28c.1
CAEEL_Y105C5B.28c.3
CAEEL_Y105C5B.28c.2
CAEEL_C45B2.5.1
HELRO_jgi_Helro1_117310
HELRO_jgi_Helro1_186556
BRAFL_XP_002595342.1
BRAFL_XP_002595343.1
GLNA_XENLA_sp_P51121_Glutamine_synthetase
CIOIN_ENSCINP00000012035
Homo_sapiens_NP_001028216_glutamine_synthetase
HOMSA_ENSP00000356537.6
HOMSA_ENSP00000307900.5
HOMSA_ENSP00000398320.2
HOMSA_ENSP00000344958.4
HOMSA_ENSP00000494022.1
MUSMU_ENSMUSP00000083375.5
MUSMU_ENSMUSP00000123157.3
STRPU_SPU_023123-tr
LAMLU_FUN_039252-T1
RIFPA_jg18944.t1
RIFPA_jg18944.t2
LINAN_g14614.t1
MNELE_ML05027a
MNELE_ML218826a
MNELE_ML05654a
BIOGL_BGLB011306-PD
NEMVE_EDO40606
AMPQU_Aqu2.1.39704_001
TRIAD_TriadP49883
BATPL_Bpl_scaf_10267-4.33
MODPH_Mph_scaf_23796-0.6
MODPH_Mph_scaf_31216-0.8
MODPH_Mph_scaf_38315-0.4
MODPH_Mph_scaf_46154-0.2
CRAGI_EKC23787
MODPH_Mph_scaf_32525-0.3
CRAGI_EKC18655
ECHMU_EmuJ_000959900.1
SCHMA_Smp_091460.1
SCHME_SMEST000211001
SCHME_SMEST000212001
SCHME_SMEST000228003
SCHME_SMEST021024001
HELRO_jgi_Helro1_186030
CRAGI_EKC27195
BATPL_Bpl_scaf_26757-3.20
MODPH_Mph_scaf_68656-0.2
CRAGI_EKC27196
LOTGI_LotgiP184997
LOTGI_LotgiP230064
BIOGL_BGLB007245-PB
BIOGL_BGLB012750-PB
NOTGE_g4242.t1
PATYE_PY_T04427
CAPTE_CapteP158978
LAMLU_FUN_004014-T1
RIFPA_jg30873.t1
RIFPA_jg30873.t2
DROME_FBpp0289795
DROME_FBpp0077774
DROME_FBpp0077773
TRICA_TC002601_001
DROME_FBpp0073344
DROME_FBpp0310163
DROME_FBpp0089332
DROME_FBpp0310162
DROME_FBpp0073345
DROME_FBpp0310164
PHOAU_g14241.t1
PHOAU_g14240.t1
OCTBI_Ocbimv22030383m.p
Glutamine synthetase II
Bacteroides_fragilis_NCTC_9343_YP_210647_glutamine_synthetase
0.6

**Supplementary figure 43 | Phylogeny of glutamine synthetase genes in selected metazoans.** Maximum-likelihood phylogenetic tree inference of glutamine synthetase genes using 1000 ultrafast bootstrap replicates. The branch support values are represented by the coloured circles in the tree nodes. Red circles represent ultrafast bootstrap values >= 95. Yellow circles represent ultrafast bootstrap values >= 90 and < 95. Green circles represent ultrafast bootstrap values < 90 and >= 70. Ultrafast bootstrap values smaller than 70 are not shown. *Bacteroides fragilis* glutamine synthetase III gene was used as outgroup. Accession numbers for NCBI database are displayed after the species names. *Capitella, Helobdella* and *Lamellibrachia* gene identification are derived from the publicly available annotated genomes. Annelid sequences from *Riftia*, *Lamellibrachia, Capitella* and *Helobdella* are closely related to glutamine synthetase I bacterial genes.

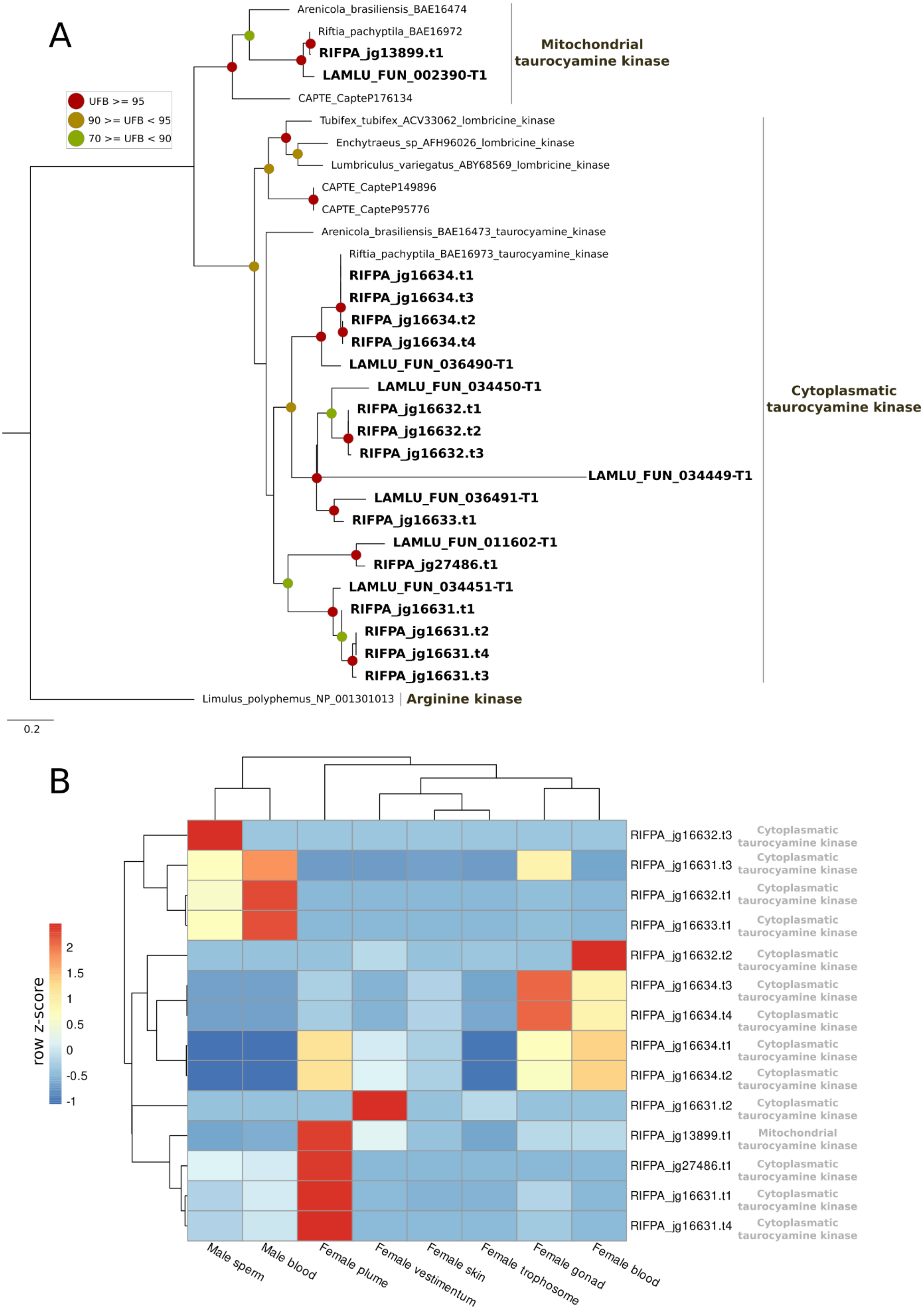

**Supplementary figure 44 | Phylogeny and gene expression of taurocyamine kinase genes. A.** Maximum-likelihood phylogenetic tree inference of the taurocyamine kinase genes using 1000 ultrafast bootstrap replicates. The branch support values are represented by the coloured circles in the tree nodes. Red circles represent ultrafast bootstrap values >= 95. Yellow circles represent ultrafast bootstrap values >= 90 and < 95. Green circles represent ultrafast bootstrap values < 90 and >= 70.

Ultrafast bootstrap values smaller than 70 are not shown. *Limulus* arginine kinase gene was used as outgroup. Accession numbers for NCBI database are displayed after the species names. *Capitella, Helobdella* and *Lamellibrachia* gene identification are derived from the publicly available annotated genomes. *Riftia* contains five copies of cytoplasmatic taurocyamine kinase genes surpassing previous estimates. **B,** Expression profile of cytoplasmatic and mitochondrial taurocyamine kinase genes in *Riftia*. Colour coding reflects the expression patterns based on row Z-score calculations.

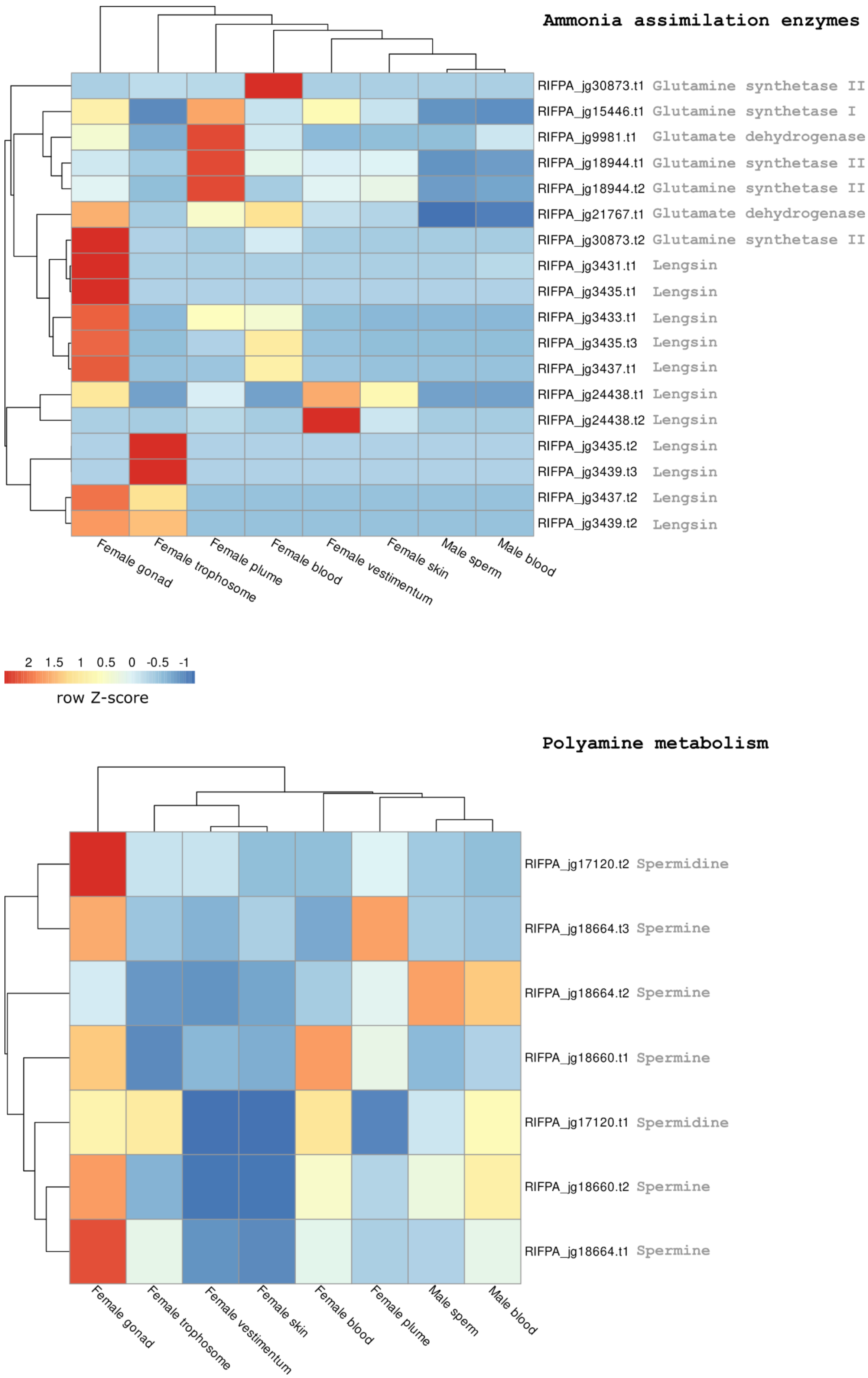

**Supplementary figure 45 | Gene expression of genes involved in the ammonia assimilation cycle and polyamine pathway in *Riftia*.** Expression profile of different key enzymes involved in the ammonia assimilation and polyamine pathway in *Riftia*. Colour coding reflects the expression patterns based on row Z-score calculations

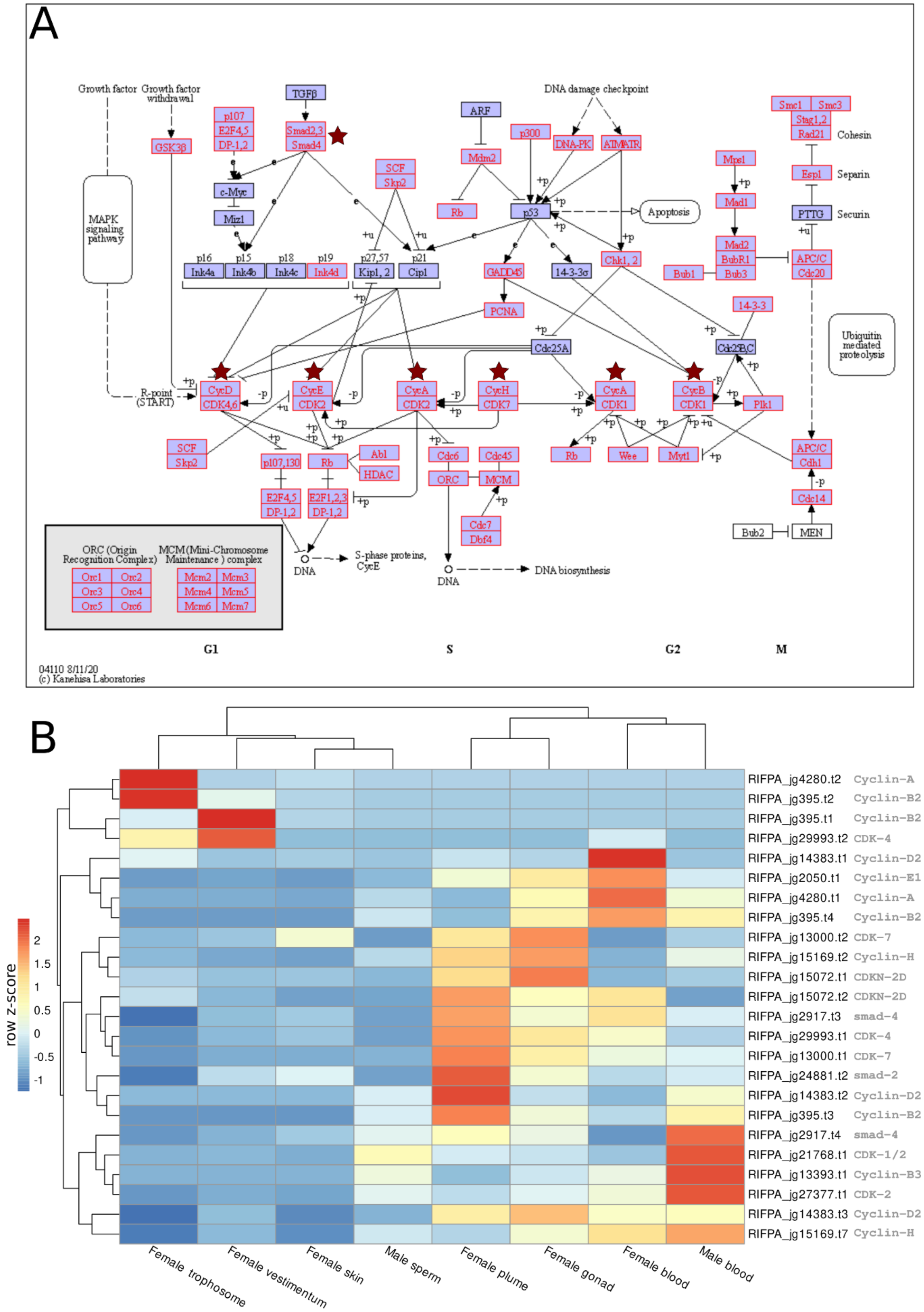

**Supplementary figure 46 – Cell cycle pathway and gene expression of cyclins and cyclin-dependent kinase genes. A**, Cell cycle pathway reference based on KEGG hsa04110 entry. Highlighted boxes correspond to the tubeworm genes identified in the reference pathway. Red stars indicate the genes used in the gene expression analyses. **B,** Expression profile of cyclin and cyclin-dependent kinase genes. Colour coding reflects the expression patterns based on row Z-score calculations.

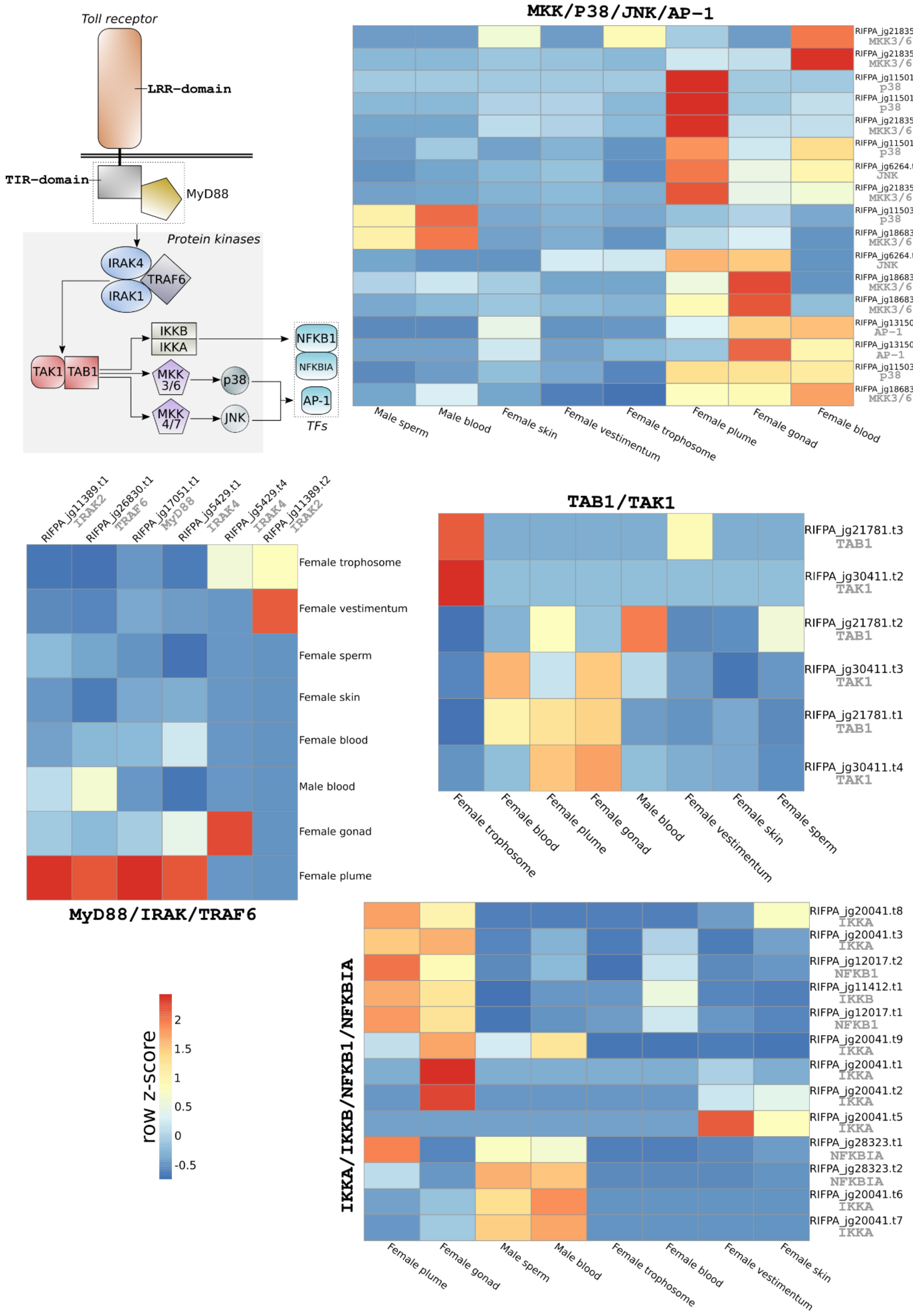

**Supplementary figure 47 | Overview of Toll-like receptor/MyD88 immune system pathway in *Riftia*.** Overview of the Toll-like receptor/MyD88 immune system pathway in *Riftia* (based on Luo et al., 2018). Expression profile of immune-related genes in the adult tissues of *Riftia pachyptila*. Colour coding in the different heatmaps reflects the expression patterns based on row Z-score calculations. It is possible to notice the high expression of immune-related genes in the plume and female gonad tissue.

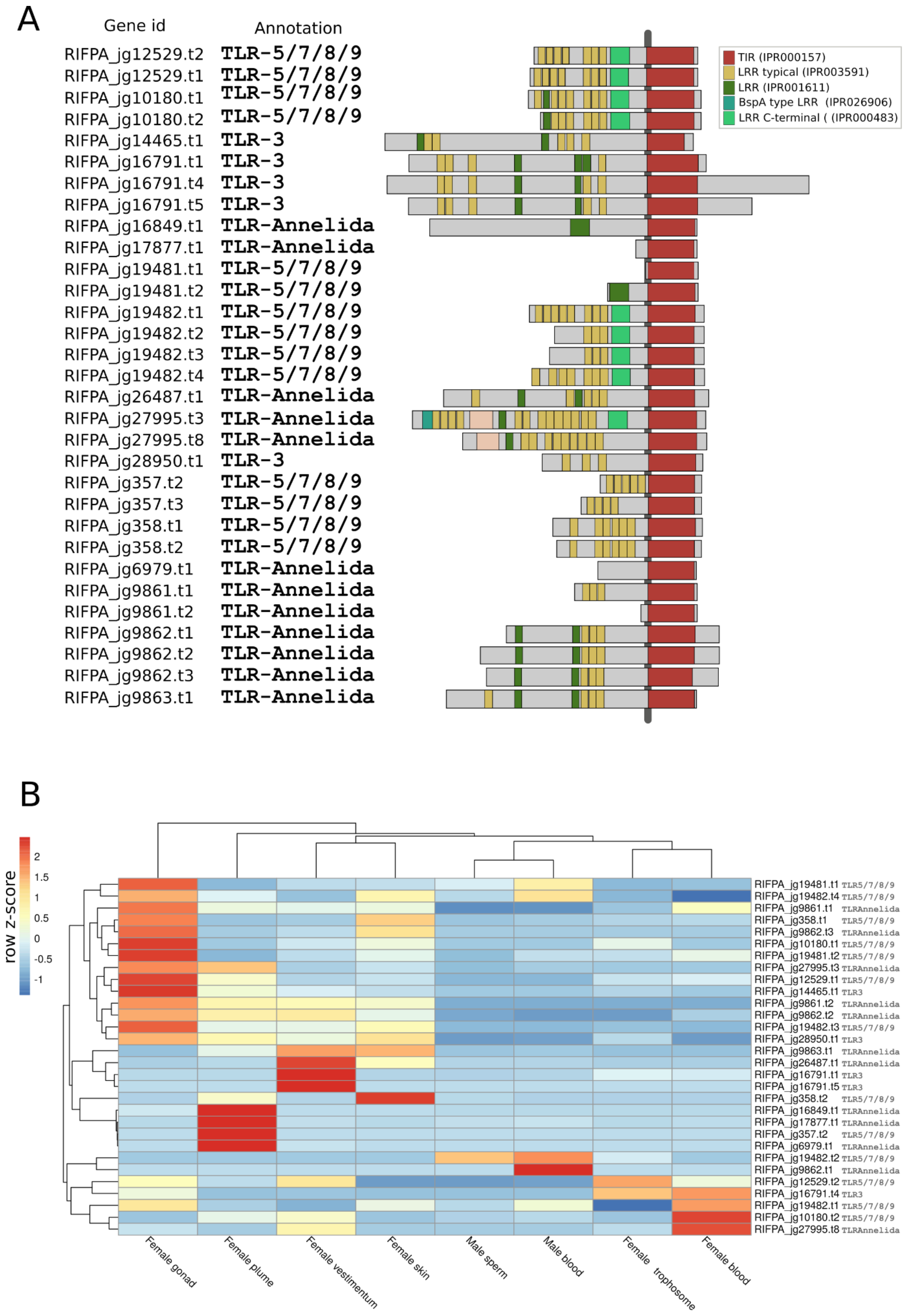

**Supplementary figure 48 | Toll-like gene domain composition and gene expression. A**, Domain composition of Toll-like genes based on PFAM database. Coloured boxes correspond to different protein domains found in the different Toll-like groups. **B,** Expression profile of Toll-like genes. Colour coding reflects the expression patterns based on row Z-score calculations. Toll-like genes are constitutively expressed in all *Riftia* adult tissues.

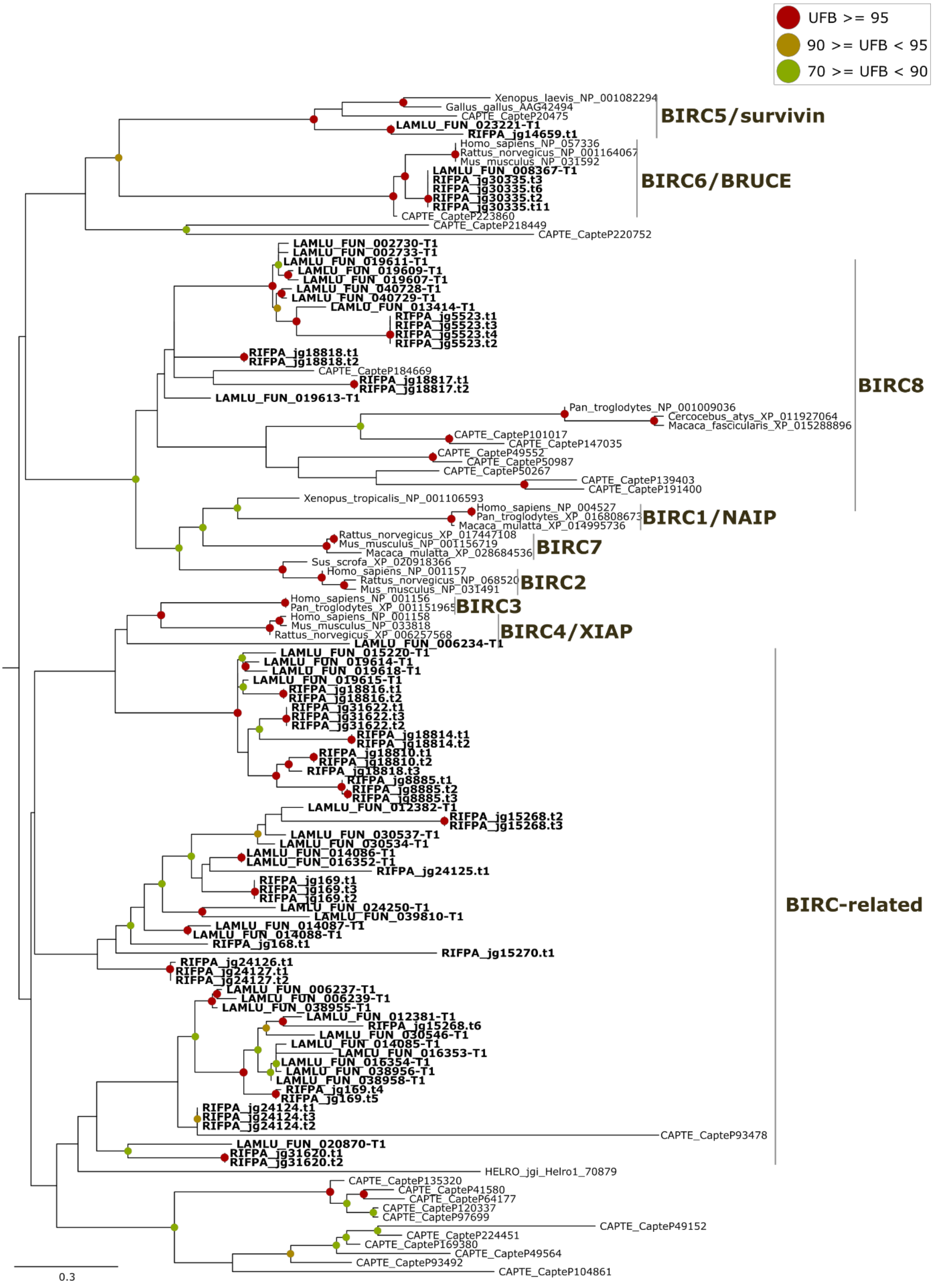

**Supplementary figure 49 - Phylogeny of Toll-like genes.** Maximum-likelihood phylogenetic tree inference of the Toll-like genes using 1000 ultrafast bootstrap replicates. The branch support values are represented by the coloured circles in the tree nodes. Red circles represent ultrafast bootstrap values >= 95. Yellow circles represent ultrafast bootstrap values >= 90 and < 95. Green circles represent ultrafast bootstrap values < 90 and >= 70. Ultrafast bootstrap values smaller than 70 are not shown. Vertebrate MyD88 genes were used as outgroup. Accession numbers for NCBI database are displayed after the species names. *Capitella, Helobdella* and *Lamellibrachia* gene identification are derived from the publicly available annotated genomes.

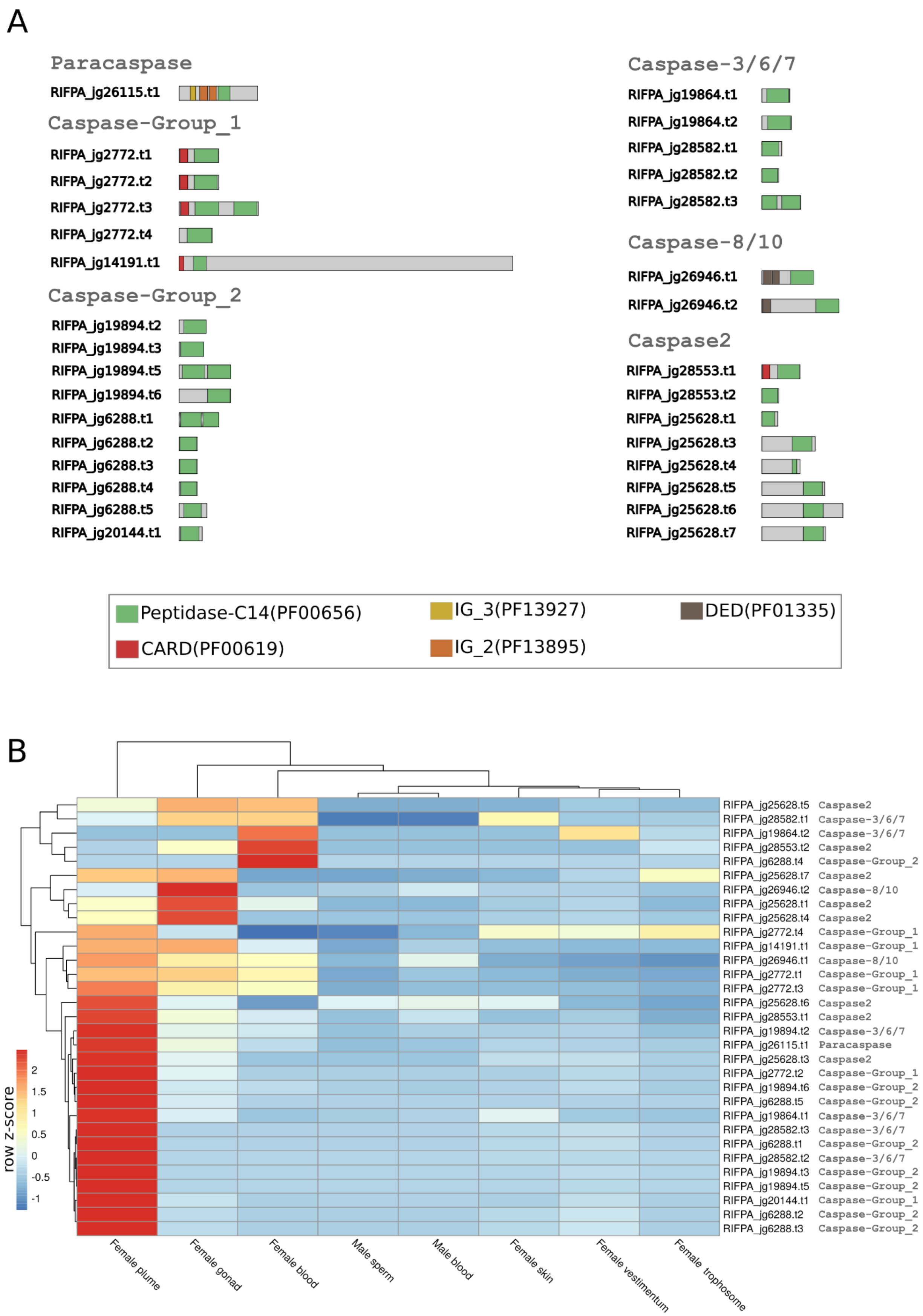

**Supplementary figure 50 | Caspase and paracaspase domain composition and gene expression. A,** Domain composition of caspases and paracaspases proteins based on PFAM database. Coloured boxes correspond to different protein domains found in the metacaspase and different caspase groups. **B,** Expression profile of caspase genes in the adult tissues of *Riftia*

*pachyptila*. Colour coding reflects the expression patterns based on row Z-score calculations. Caspases are highly expressed in the plume and female gonad.

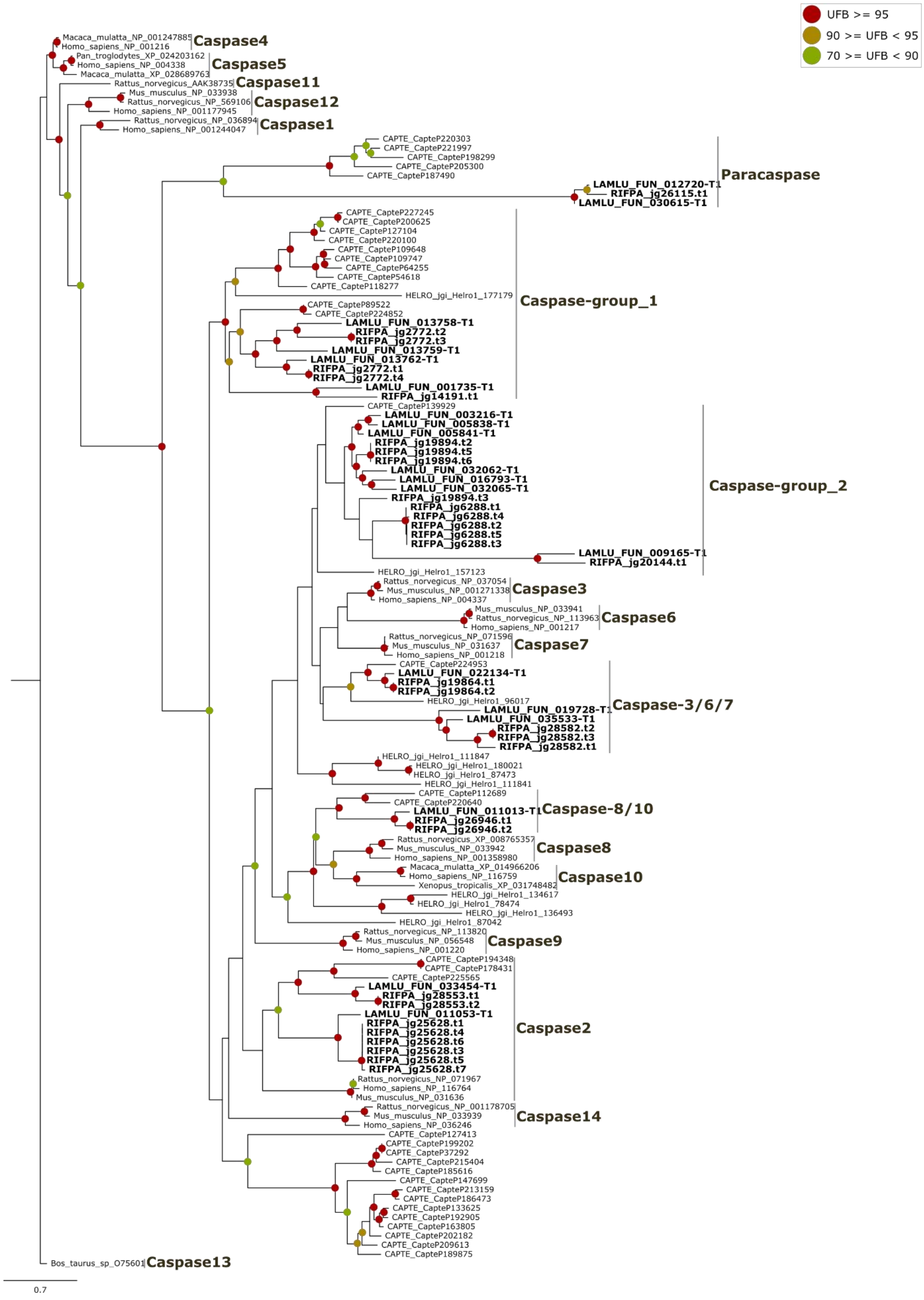

**Supplementary figure 51 | Phylogeny of caspases and paracaspases.** Maximum-likelihood phylogenetic tree inference of the caspases and paracaspases using 1000 ultrafast bootstrap replicates. The branch support values are represented by the coloured circles in the tree nodes. Red circles represent ultrafast bootstrap values >= 95. Yellow circles represent ultrafast bootstrap values >= 90 and < 95. Green circles represent ultrafast bootstrap values < 90 and >= 70. Ultrafast bootstrap values smaller than 70 are not shown. *Bos taurus* caspase-13 was used as outgroup. Accession numbers for NCBI database are displayed after the species names. *Capitella, Helobdella* and *Lamellibrachia* gene identification are derived from the publicly available annotated genomes.

A

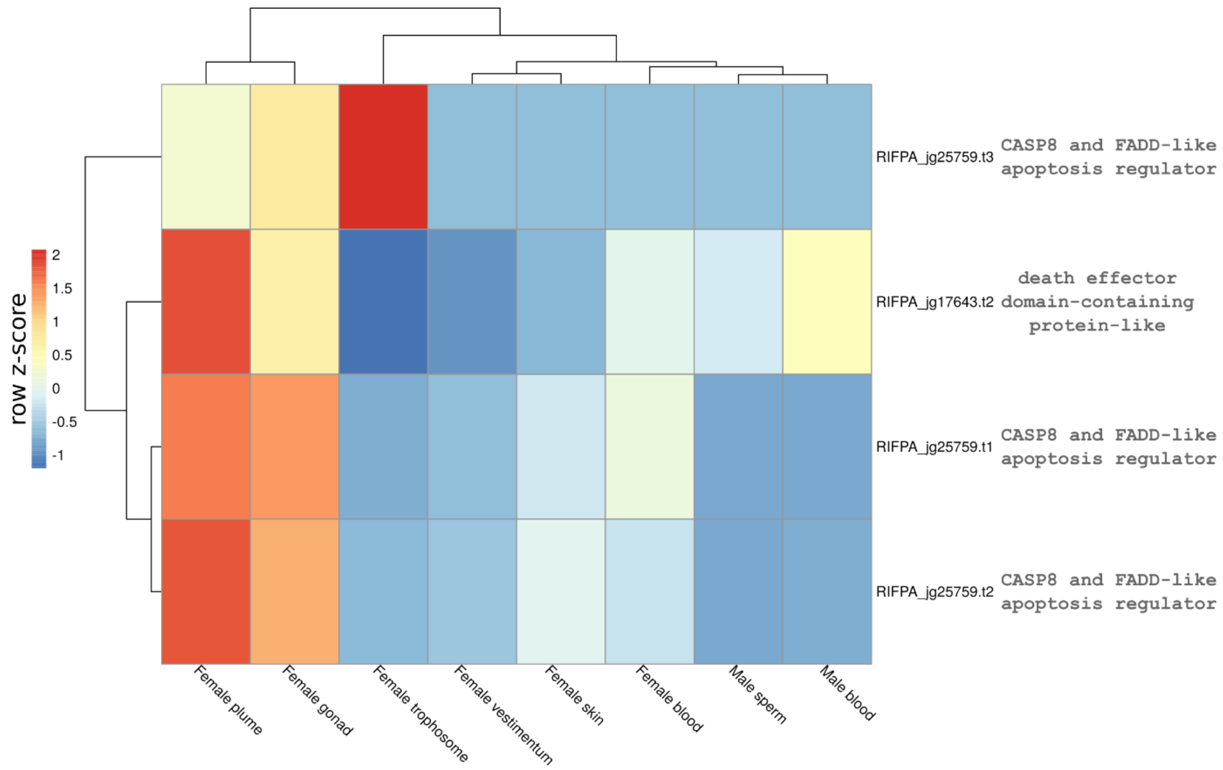

B

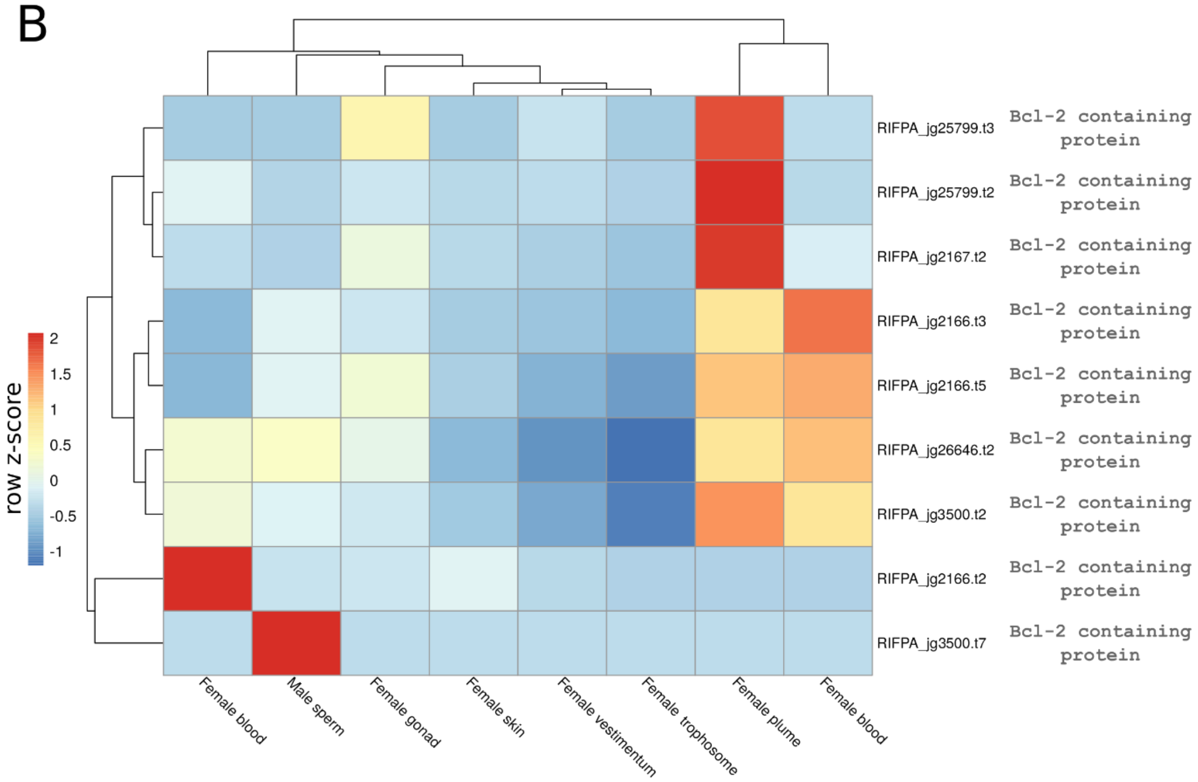

**Supplementary figure 52 | Gene expression of DED and BCL2 domain -containing proteins. A-B**, Expression profile of DED and BCL-2 domain-containing proteins obtained from the PFAM analysis. Colour coding reflects the expression patterns based on row Z-score calculations. DED- and BCL-2-domain containing proteins are highly expressed in the plume and female blood.

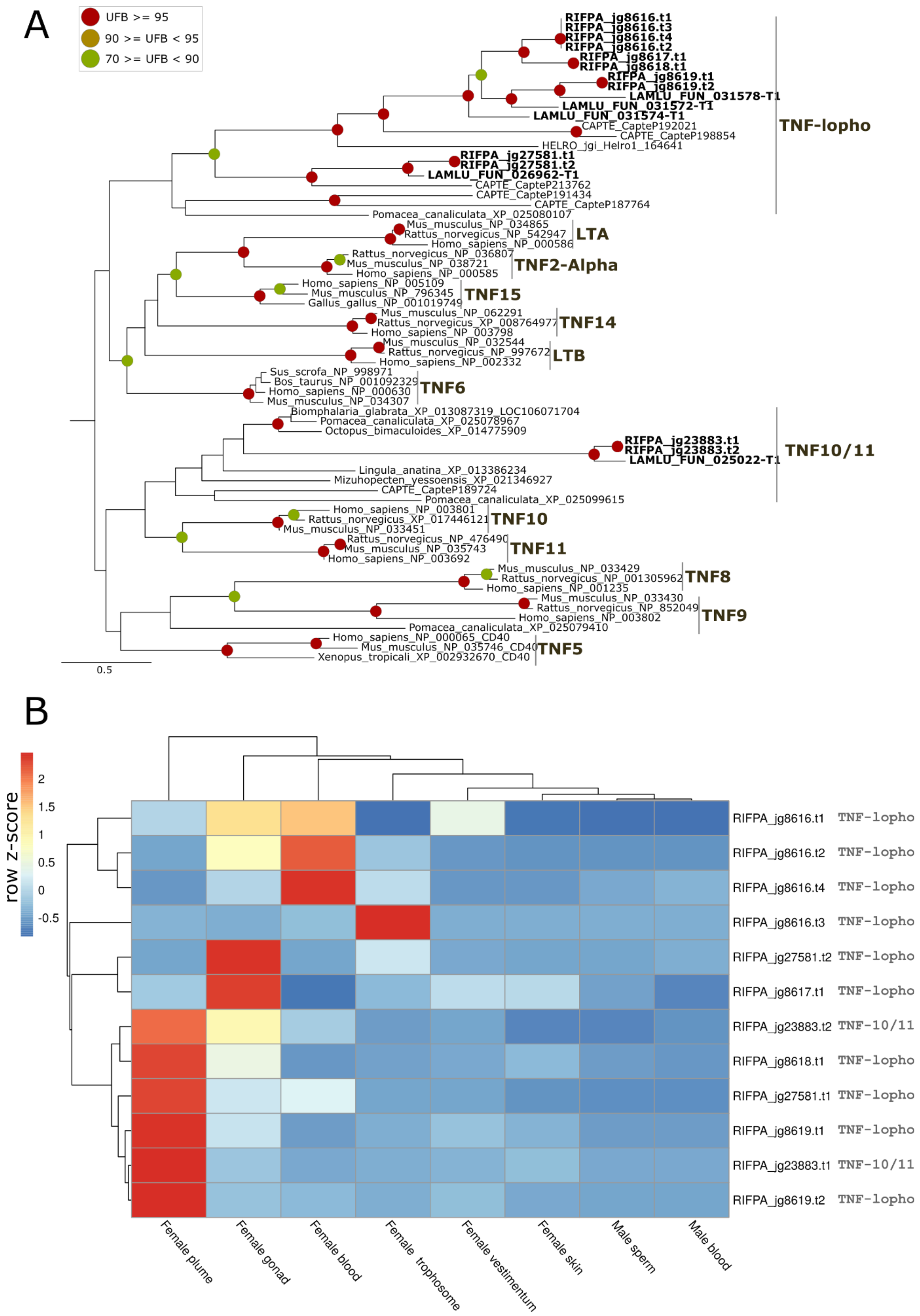

**Supplementary figure 53 | Phylogeny and gene expression of TNF ligands. A,** Mid-rooted maximum-likelihood phylogenetic tree inference of the TNF ligands using 1000 ultrafast bootstrap replicates. The branch support values are represented by the coloured circles in the tree nodes. Red circles represent ultrafast bootstrap values >= 95. Yellow circles represent ultrafast bootstrap values >= 90 and < 95. Green circles represent ultrafast bootstrap values < 90 and >= 70. Ultrafast bootstrap values smaller than 70 are not shown. Accession numbers for NCBI database are displayed after the species names. *Capitella, Helobdella* and *Lamellibrachia* gene identification are derived from the publicly available annotated genomes. **B,** Expression profile of TNF ligands. Colour coding reflects the expression patterns based on row Z-score calculations.

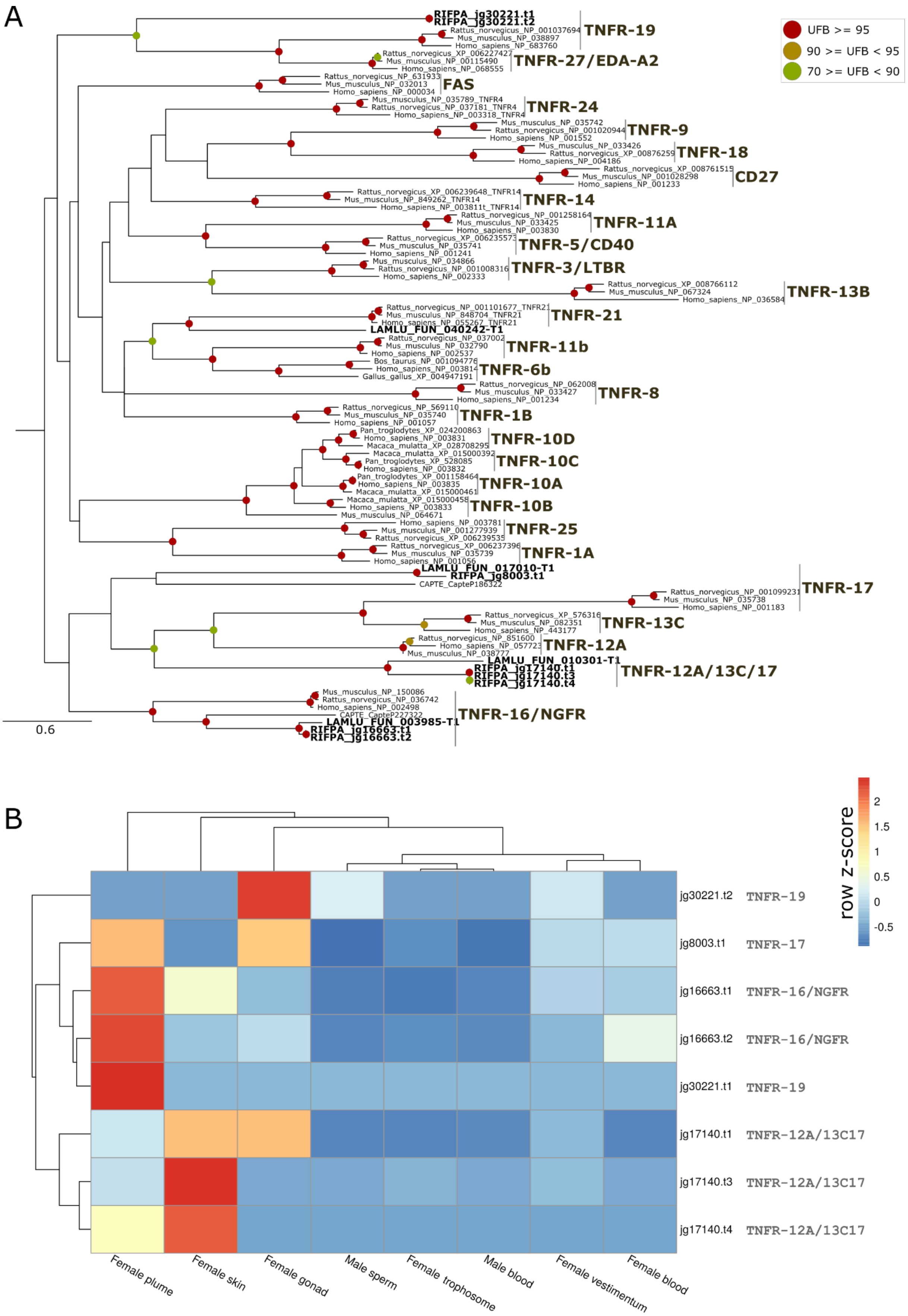

**Supplementary figure 54 | Phylogeny and gene expression of TNF receptors. A,** Mid-rooted maximum-likelihood phylogenetic tree inference of the TNF receptors using 1000 ultrafast bootstrap replicates. The branch support values are represented by the coloured circles in the tree nodes. Red circles represent ultrafast bootstrap values >= 95. Yellow circles represent ultrafast bootstrap values >= 90 and < 95. Green circles represent ultrafast bootstrap values < 90 and >= 70. Ultrafast bootstrap values smaller than 70 are not shown. Accession numbers for NCBI database are displayed after the species names. *Capitella, Helobdella* and *Lamellibrachia* gene identification are derived from the publicly available annotated genomes. **B,** Expression profile of TNF receptors. colour coding reflects the expression patterns based on row Z-score calculations.

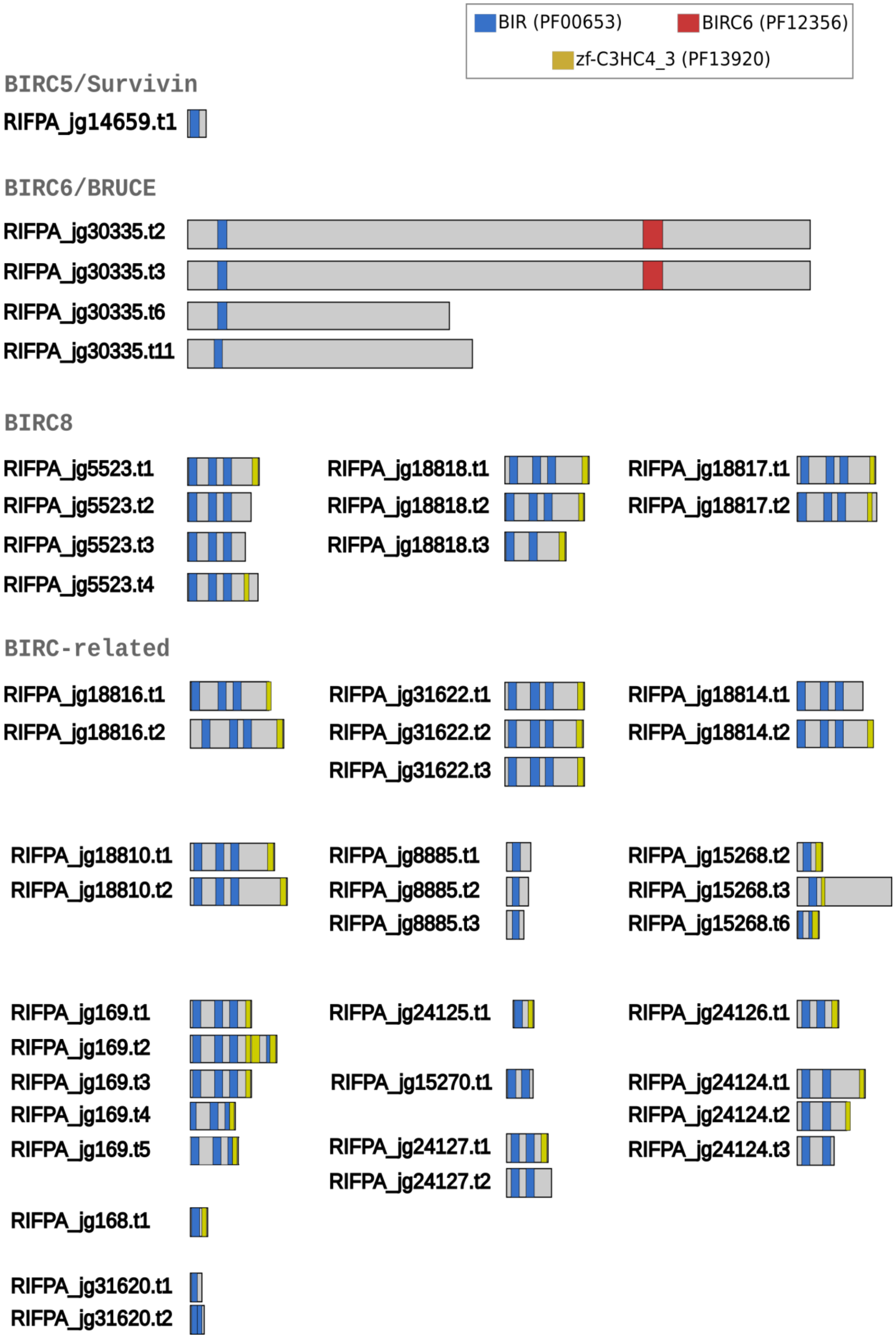
BIR (PF00653)
BIRC6 (PF12356)
zf-C3HC4_3 (PF13920)
BIRC5/Survivin
RIFPA_jg14659.t1
BIRC6/BRUCE
RIFPA_jg30335.t2
RIFPA_jg30335.t3
RIFPA_jg30335.t6
RIFPA_jg30335.t11
BIRC8
RIFPA_jg5523.t1
RIFPA_jg5523.t2
RIFPA_jg5523.t3
RIFPA_jg5523.t4
RIFPA_jg18818.t1
RIFPA_jg18818.t2
RIFPA_jg18818.t3
RIFPA_jg18817.t1
RIFPA_jg18817.t2
BIRC-related
RIFPA_jg18816.t1
RIFPA_jg18816.t2
RIFPA_jg31622.t1
RIFPA_jg31622.t2
RIFPA_jg31622.t3
RIFPA_jg18814.t1
RIFPA_jg18814.t2
RIFPA_jg18810.t1
RIFPA_jg18810.t2
RIFPA_jg8885.t1
RIFPA_jg8885.t2
RIFPA_jg8885.t3
RIFPA_jg15268.t2
RIFPA_jg15268.t3
RIFPA_jg15268.t6
RIFPA_jg169.t1
RIFPA_jg169.t2
RIFPA_jg169.t3
RIFPA_jg169.t4
RIFPA_jg169.t5
RIFPA_jg24125.t1
RIFPA_jg15270.t1
RIFPA_jg24127.t1
RIFPA_jg24127.t2
RIFPA_jg24126.t1
RIFPA_jg24124.t1
RIFPA_jg24124.t2
RIFPA_jg24124.t3
RIFPA_jg168.t1
RIFPA_jg31620.t1
RIFPA_jg31620.t2

**Supplementary figure 55 | Domain composition of IAP genes**. Domain composition of IAP genes based on PFAM database. Coloured boxes correspond to different protein domains found in the different IAP groups. We identified a complement of 19 IAP genes on *Riftia* genome.

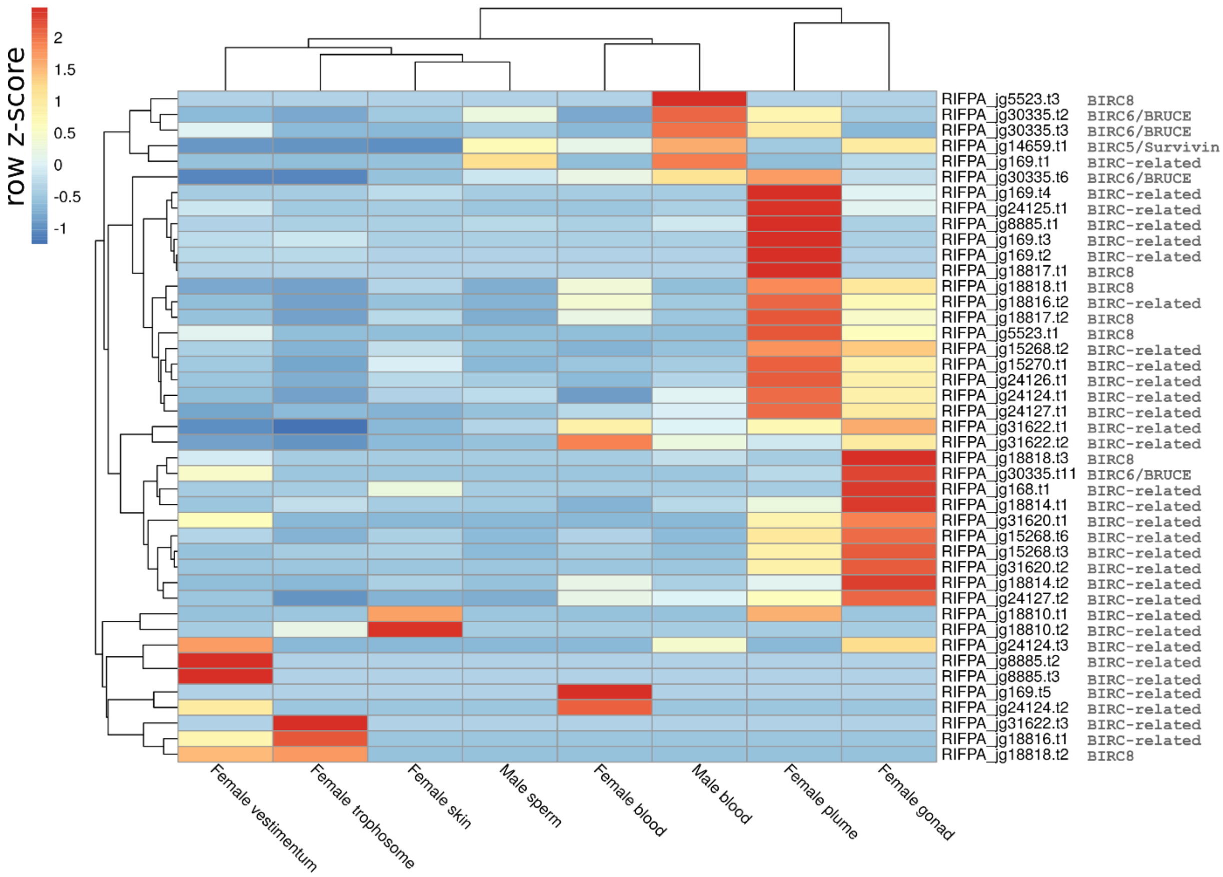

**Supplementary figure 56 | Gene expression of IAP domain-containing proteins.** Expression profile IAP domain-containing proteins obtained from the PFAM analysis. Colour coding reflects the expression patterns based on row Z-score calculations. IAP gene expression is present in all tissues, however, many IAP genes are highly expressed on the female gonad and plume tissues.

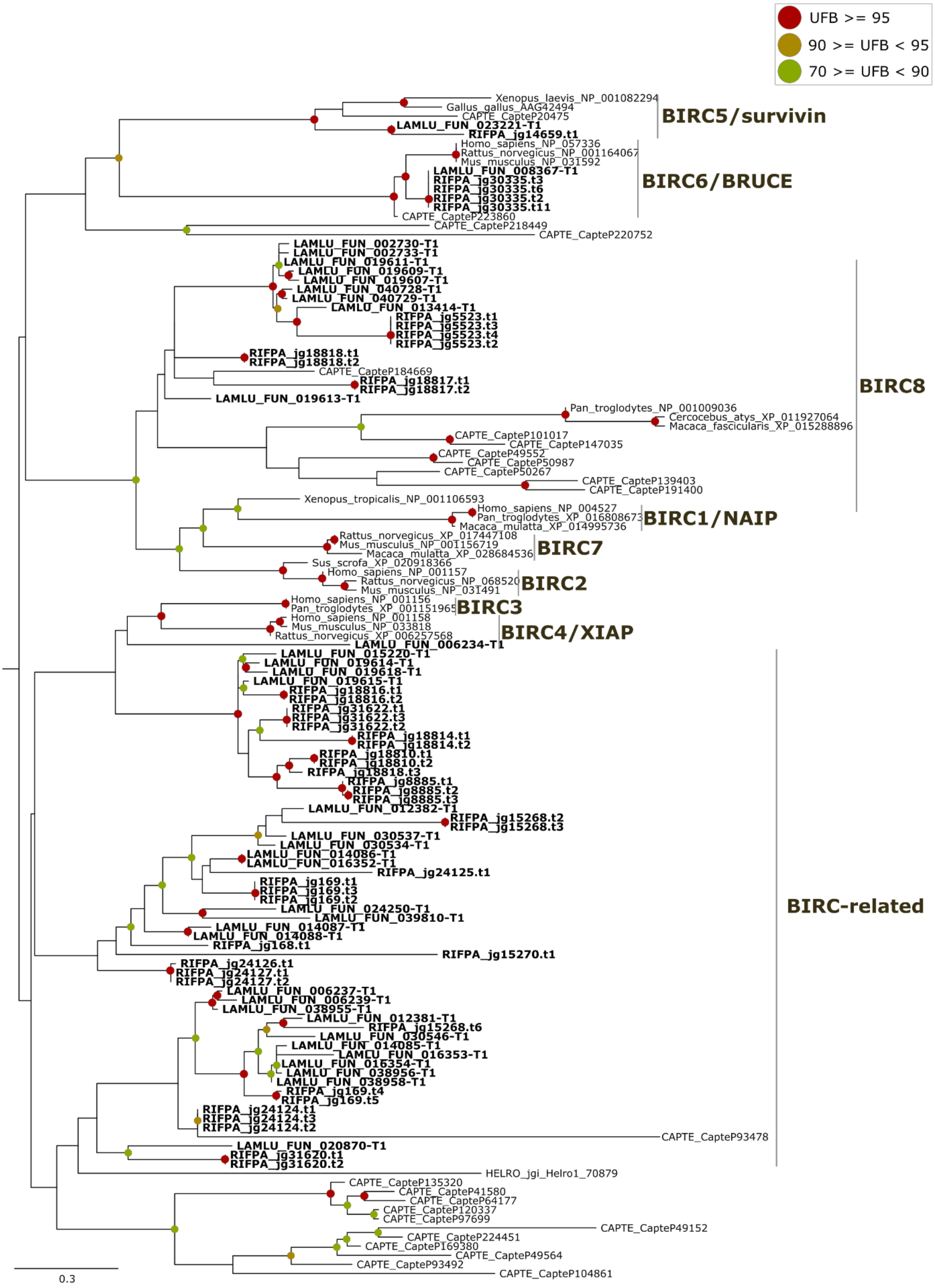

**Supplementary figure 57 | Phylogeny of IAP genes.** Mid-rooted maximum-likelihood phylogenetic tree inference of the IAP genes using 1000 ultrafast bootstrap replicates. The branch support values are represented by the coloured circles in the tree nodes. Red circles represent ultrafast bootstrap values >= 95. Yellow circles represent ultrafast bootstrap values >= 90 and < 95. Green circles represent ultrafast bootstrap values < 90 and >= 70. Ultrafast bootstrap values smaller than 70 are not shown. Accession numbers for NCBI database are displayed after the species names. *Capitella, Helobdella* and *Lamellibrachia* gene identification are derived from the publicly available annotated genomes.

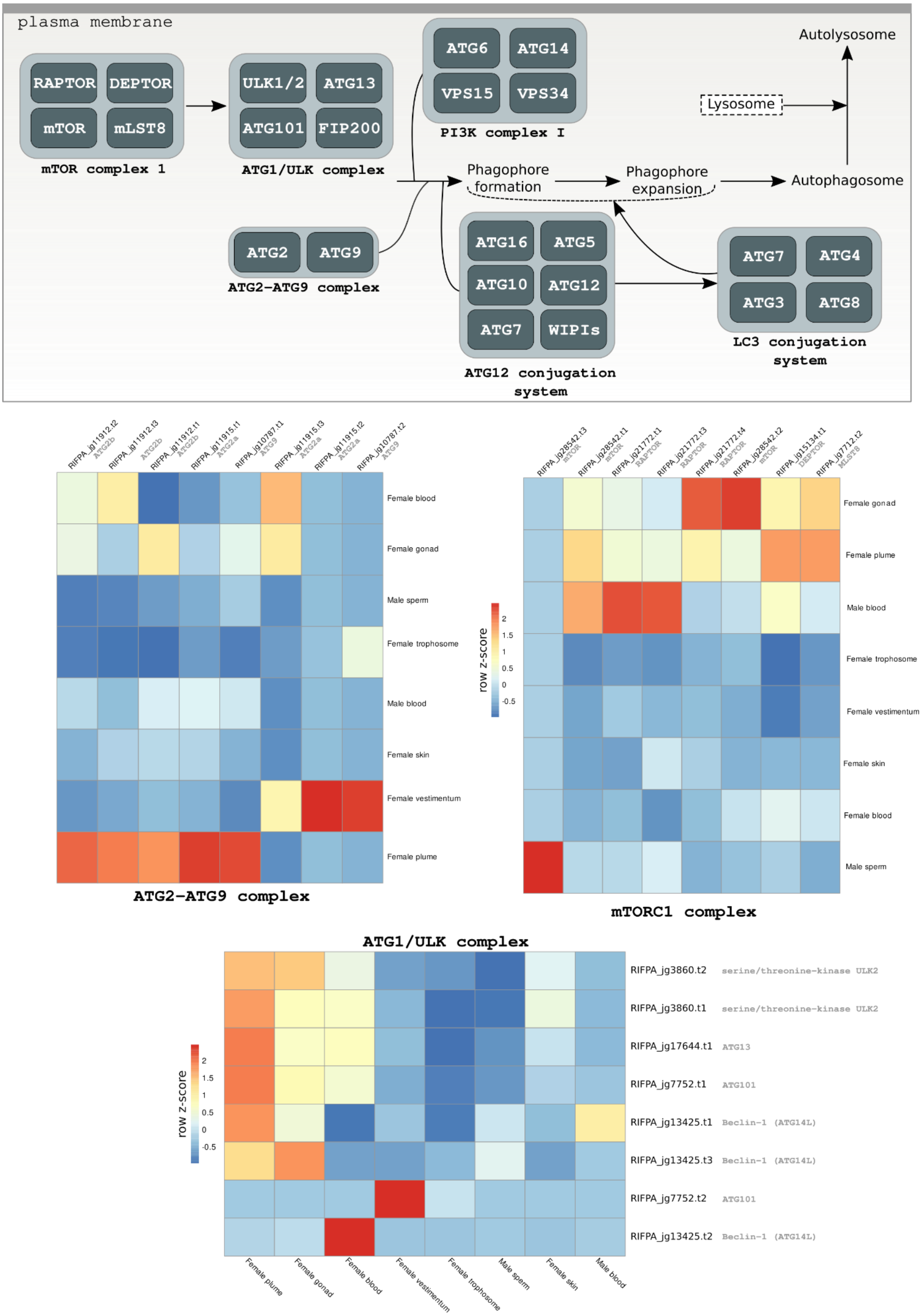

plasma membrane
RAPTOR
DEPTOR
mTOR
mLST8
mTOR complex 1
ULK1/2
ATG13
ATG101
FIP200
ATG1/ULK complex
ATG6
ATG14
VPS15
VPS34
PI3K complex I
Autolysosome
Lysosome
Phagophore formation
Phagophore expansion
Autophagosome
ATG2
ATG9
ATG2-ATG9 complex
ATG16
ATG5
ATG10
ATG12
ATG7
WIPIs
ATG12 conjugation system
ATG7
ATG4
ATG3
ATG8
LC3 conjugation system
RIFPA_jg11912.t2
RIFPA_jg11912.t3
RIFPA_jg11912.t1
RIFPA_jg11915.t1
RIFPA_jg10787.t1
RIFPA_jg11915.t3
RIFPA_jg11915.t2
RIFPA_jg10787.t2
Female blood
Female gonad
Male sperm
Female trophosome
Male blood
Female skin
Female vestimentum
Female plume
ATG2-ATG9 complex
row z-score
2
1.5
1
0.5
0
-0.5
RIFPA_jg28542.t3
RIFPA_jg28542.t1
RIFPA_jg21772.t1
RIFPA_jg21772.t3
RIFPA_jg21772.t4
RIFPA_jg28542.t2
RIFPA_jg15134.t1
RIFPA_jg7712.t2
Female gonad
Female plume
Male blood
Female trophosome
Female vestimentum
Female skin
Female blood
Male sperm
mTORC1 complex
ATG1/ULK complex
RIFPA_jg3860.t2 serine/threonine-kinase ULK2
RIFPA_jg3860.t1 serine/threonine-kinase ULK2
RIFPA_jg17644.t1 ATG13
RIFPA_jg7752.t1 ATG101
RIFPA_jg13425.t1 Beclin-1 (ATG14L)
RIFPA_jg13425.t3 Beclin-1 (ATG14L)
RIFPA_jg7752.t2 ATG101
RIFPA_jg13425.t2 Beclin-1 (ATG14L)
row z-score
2
1.5
1
0.5
0
-0.5
Female plume
Female gonad
Female blood
Female vestimentum
Female trophosome
Male sperm
Female skin
Male blood

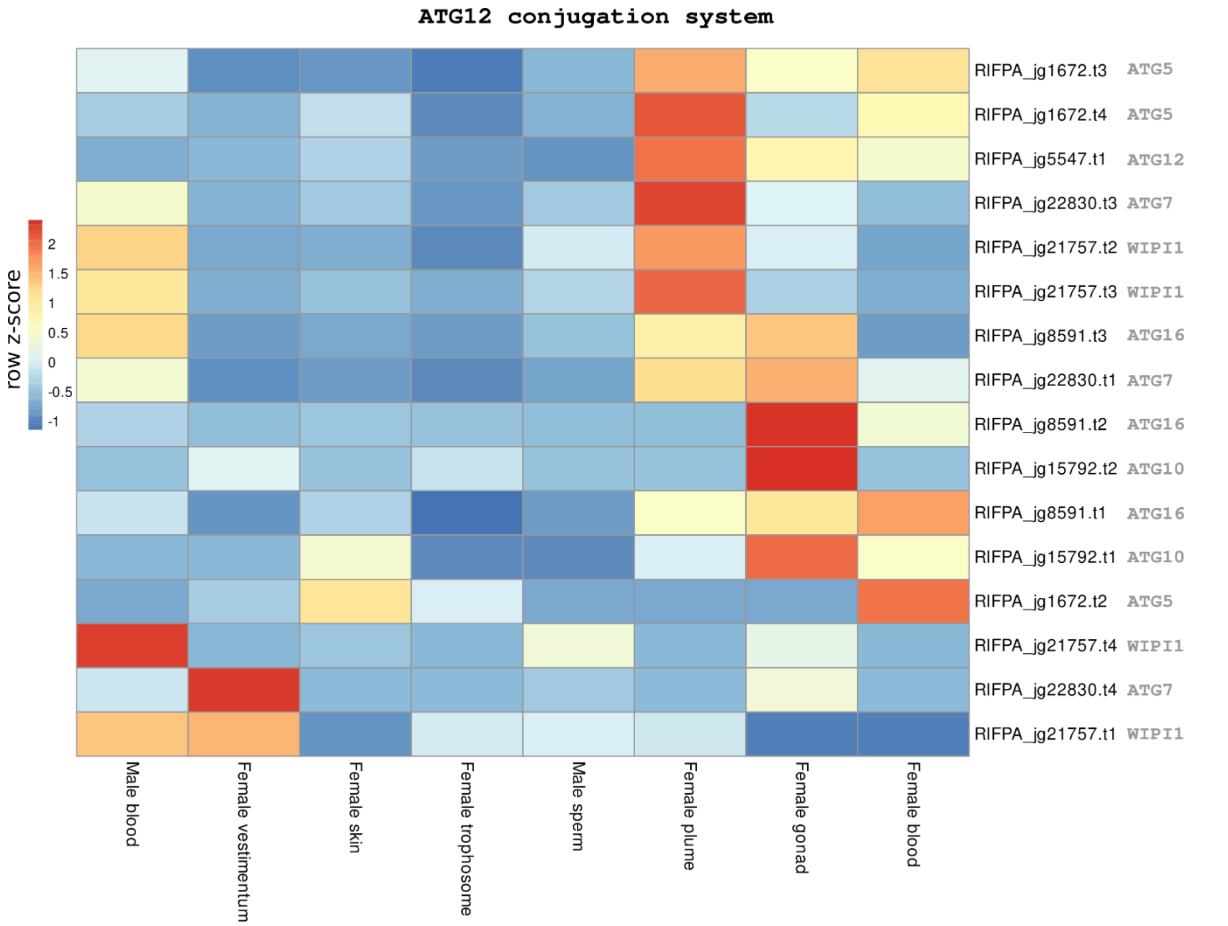
ATG12 conjugation system
row z-score
2
1.5
1
0.5
0
-0.5
-1
RIFPA_jg1672.t3 ATG5
RIFPA_jg1672.t4 ATG5
RIFPA_jg5547.t1 ATG12
RIFPA_jg22830.t3 ATG7
RIFPA_jg21757.t2 WIPI1
RIFPA_jg21757.t3 WIPI1
RIFPA_jg8591.t3 ATG16
RIFPA_jg22830.t1 ATG7
RIFPA_jg8591.t2 ATG16
RIFPA_jg15792.t2 ATG10
RIFPA_jg8591.t1 ATG16
RIFPA_jg15792.t1 ATG10
RIFPA_jg1672.t2 ATG5
RIFPA_jg21757.t4 WIPI1
RIFPA_jg22830.t4 ATG7
RIFPA_jg21757.t1 WIPI1
Male blood
Female vestimentum
Female skin
Female trophosome
Male sperm
Female plume
Female gonad
Female blood

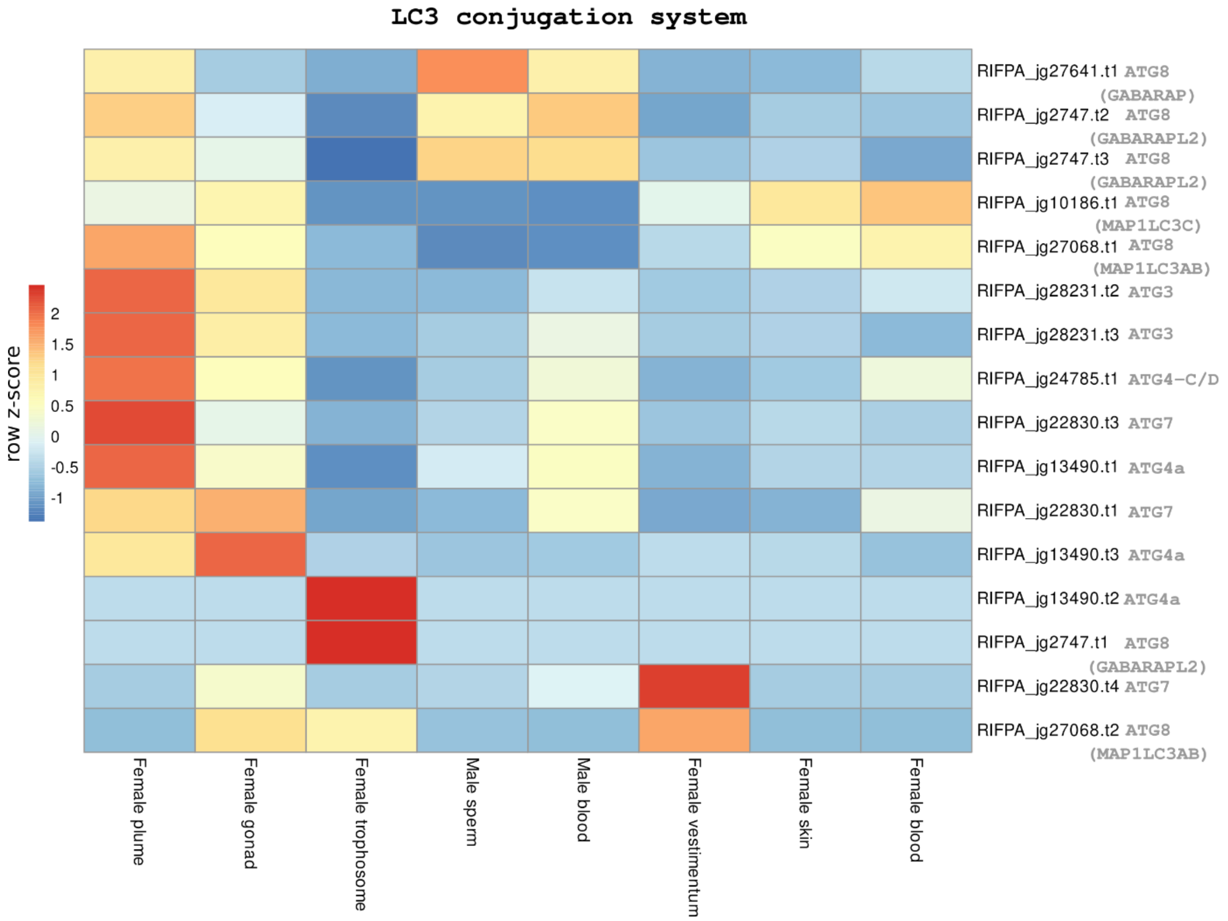
LC3 conjugation system
row z-score
2
1.5
1
0.5
0
-0.5
-1
RIFPA_jg27641.t1 ATG8 (GABARAP)
RIFPA_jg2747.t2 ATG8 (GABARAPL2)
RIFPA_jg2747.t3 ATG8 (GABARAPL2)
RIFPA_jg10186.t1 ATG8 (MAP1LC3C)
RIFPA_jg27068.t1 ATG8 (MAP1LC3AB)
RIFPA_jg28231.t2 ATG3
RIFPA_jg28231.t3 ATG3
RIFPA_jg24785.t1 ATG4-C/D
RIFPA_jg22830.t3 ATG7
RIFPA_jg13490.t1 ATG4a
RIFPA_jg22830.t1 ATG7
RIFPA_jg13490.t3 ATG4a
RIFPA_jg13490.t2 ATG4a
RIFPA_jg2747.t1 ATG8 (GABARAPL2)
RIFPA_jg22830.t4 ATG7
RIFPA_jg27068.t2 ATG8 (MAP1LC3AB)
Female plume
Female gonad
Female trophosome
Male sperm
Male blood
Female vestimentum
Female skin
Female blood

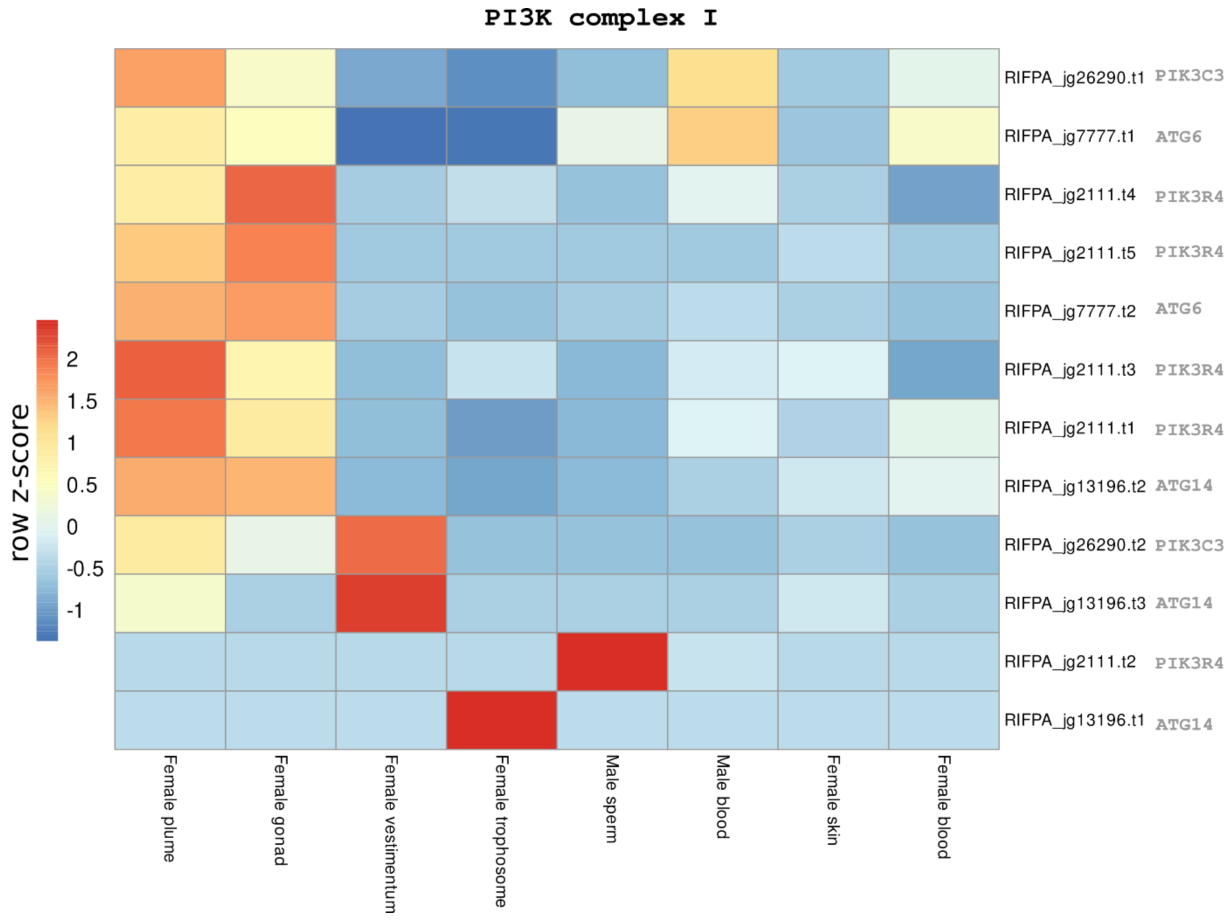

**Supplementary figure 58 | Overview of autophagy pathway in *Riftia* and gene expression of autophagy-related genes.** Overview of the core elements present in the autophagy pathway in *Riftia.* The giant tubeworm contains all the core elements commonly found in yeast, deuterostomes and lophotrochozoans. Expression profile of autophagy-related genes in the adult tissues of *Riftia pachyptila*. Colour coding in the different heatmaps reflects the expression patterns based on row Z-score calculations. It is possible to notice the high expression of autophagy-related genes in the plume and female gonad tissue.

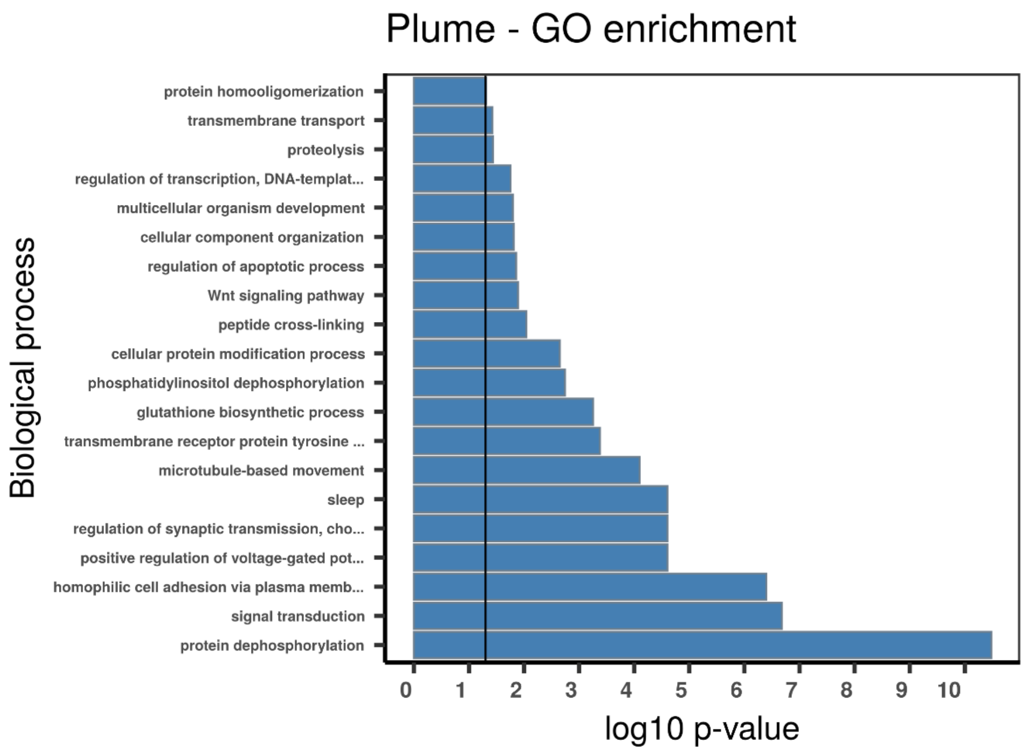
Plume - GO enrichment
Biological process
protein homooligomerization
transmembrane transport
proteolysis
regulation of transcription, DNA-templat...
multicellular organism development
cellular component organization
regulation of apoptotic process
Wnt signaling pathway
peptide cross-linking
cellular protein modification process
phosphatidylinositol dephosphorylation
glutathione biosynthetic process
transmembrane receptor protein tyrosine ...
microtubule-based movement
sleep
regulation of synaptic transmission, cho...
positive regulation of voltage-gated pot...
homophilic cell adhesion via plasma memb...
signal transduction
protein dephosphorylation
0 1 2 3 4 5 6 7 8 9 10
log10 p-value

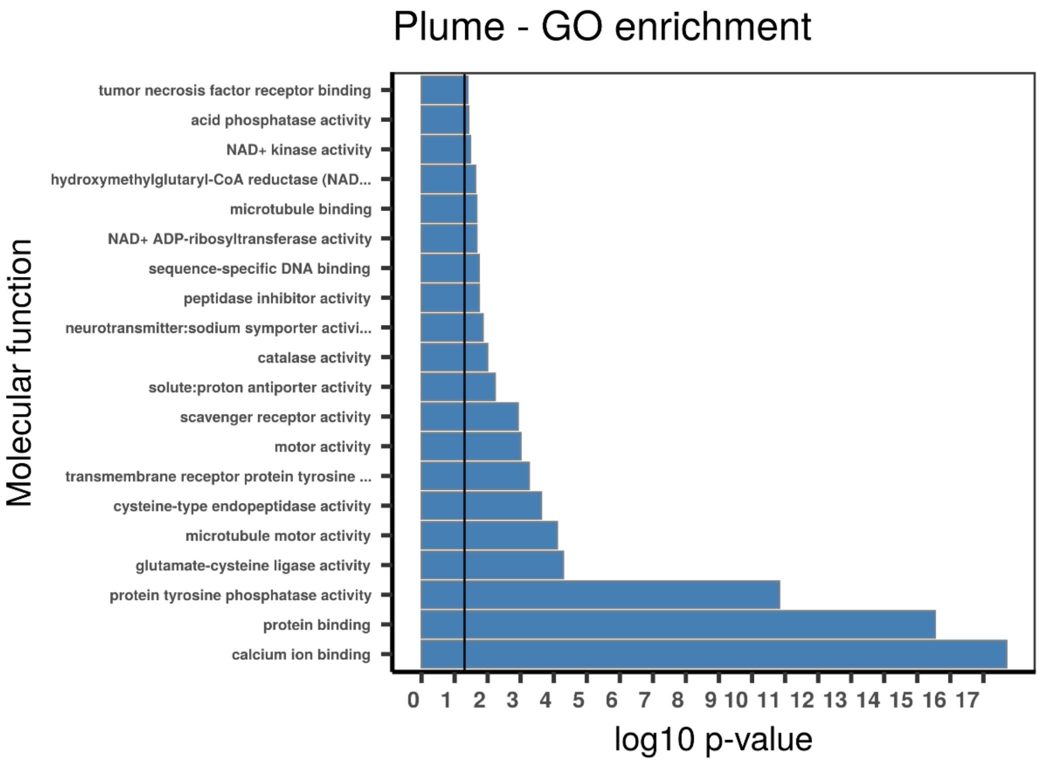
Plume - GO enrichment
Molecular function
tumor necrosis factor receptor binding
acid phosphatase activity
NAD+ kinase activity
hydroxymethylglutaryl-CoA reductase (NAD...
microtubule binding
NAD+ ADP-ribosyltransferase activity
sequence-specific DNA binding
peptidase inhibitor activity
neurotransmitter:sodium symporter activi...
catalase activity
solute:proton antiporter activity
scavenger receptor activity
motor activity
transmembrane receptor protein tyrosine ...
cysteine-type endopeptidase activity
microtubule motor activity
glutamate-cysteine ligase activity
protein tyrosine phosphatase activity
protein binding
calcium ion binding
0 1 2 3 4 5 6 7 8 9 10 11 12 13 14 15 16 17
log10 p-value

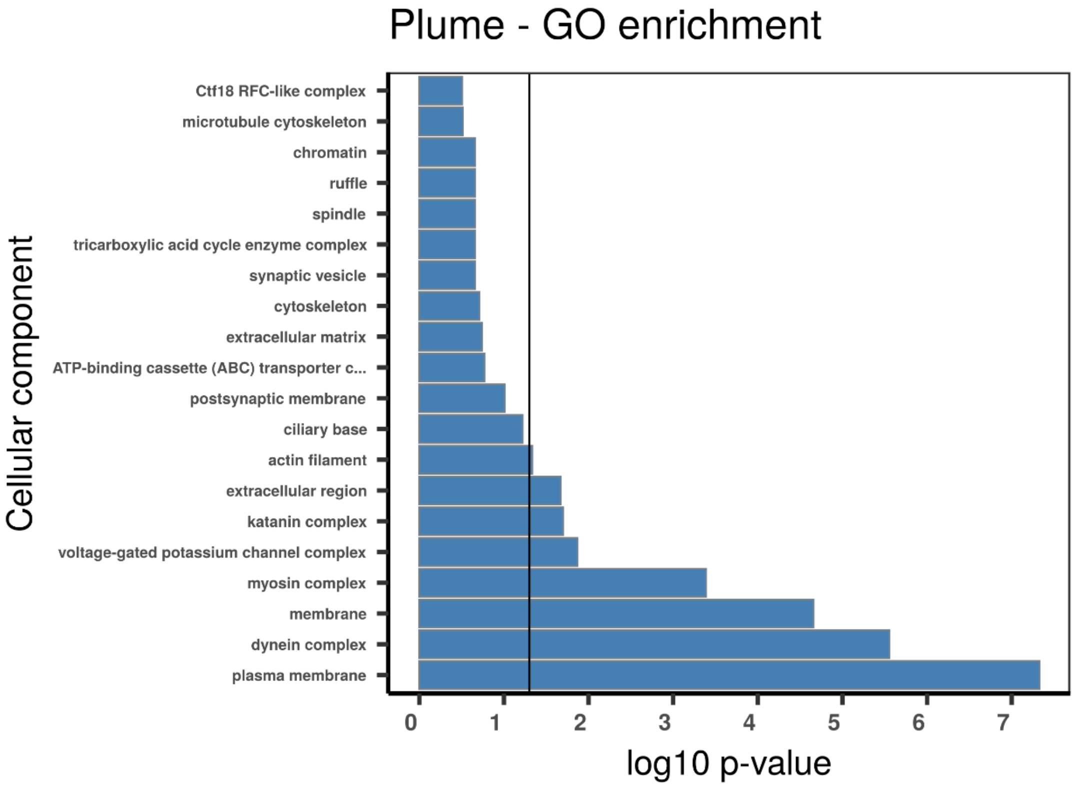

**Supplementary figure 59 | Gene set enrichment analysis with topGO using absolutely plume specific TAU genes**. Gene ontology (GO) enrichment analyses for absolutely plume specific TAU genes. The graphs correspond to the three domains of ontologies: biological process (BP), molecular function (MF) and cellular component (CC). The selected genes were analysed for enrichment in specific GO categories using the TopGO program against the background (all coding sequence genes). Y axis corresponds to enriched GO terms found in the respective domains (BP, MF and CC). X axis correspond to the log function of Fisher p-values obtained for each one of the enriched terms. The back line denotes a p-value = 0.05. P-values greater than 1,30 (log 0,05) indicate statistically significant enriched term. Genes involved in apoptosis, cytoskeleton, signal transduction and cell division are differentially expressed in the plume.

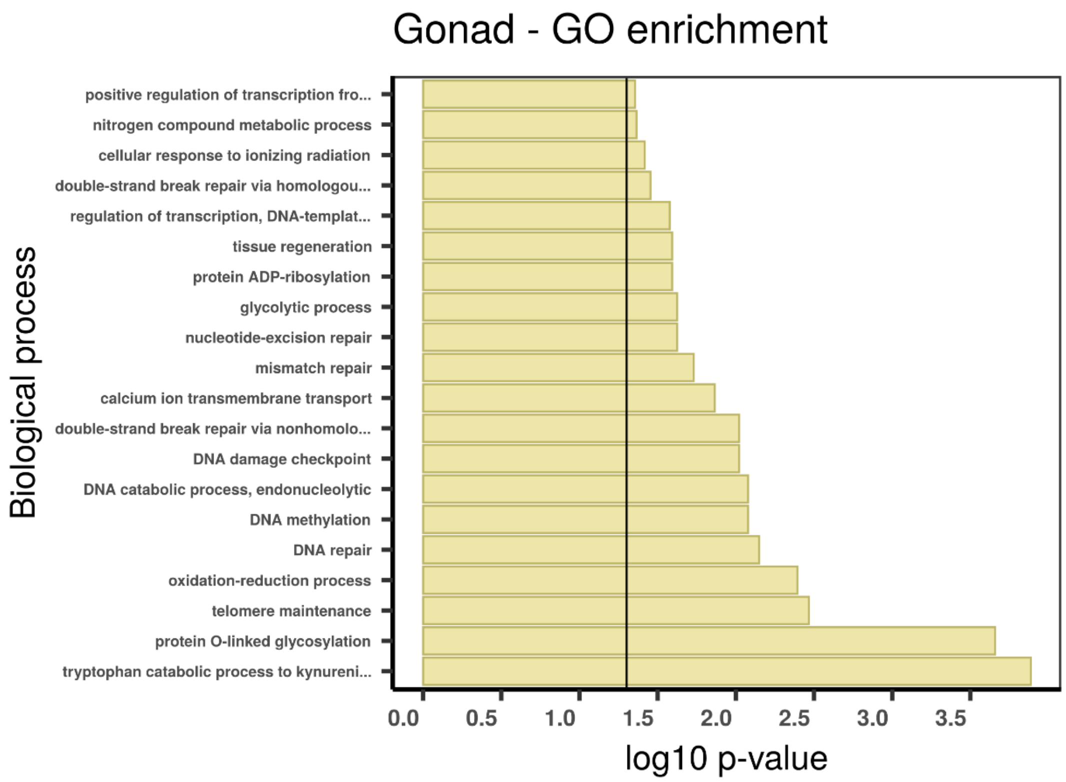

Gonad - GO enrichment
Biological process
positive regulation of transcription fro...
nitrogen compound metabolic process
cellular response to ionizing radiation
double-strand break repair via homologou...
regulation of transcription, DNA-templat...
tissue regeneration
protein ADP-ribosylation
glycolytic process
nucleotide-excision repair
mismatch repair
calcium ion transmembrane transport
double-strand break repair via nonhomolo...
DNA damage checkpoint
DNA catabolic process, endonucleolytic
DNA methylation
DNA repair
oxidation-reduction process
telomere maintenance
protein O-linked glycosylation
tryptophan catabolic process to kynureni...
0.0
0.5
1.0
1.5
2.0
2.5
3.0
3.5
log10 p-value

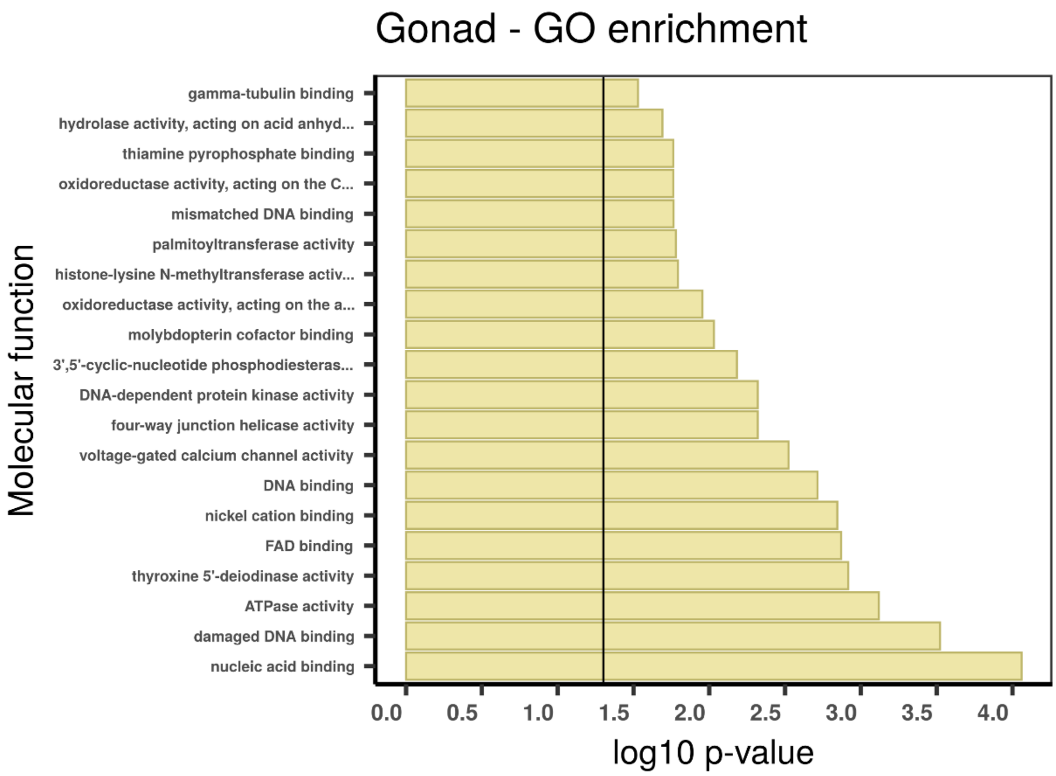

Gonad - GO enrichment
Molecular function
gamma-tubulin binding
hydrolase activity, acting on acid anhyd...
thiamine pyrophosphate binding
oxidoreductase activity, acting on the C...
mismatched DNA binding
palmitoyltransferase activity
histone-lysine N-methyltransferase activ...
oxidoreductase activity, acting on the a...
molybdopterin cofactor binding
3',5'-cyclic-nucleotide phosphodiesteras...
DNA-dependent protein kinase activity
four-way junction helicase activity
voltage-gated calcium channel activity
DNA binding
nickel cation binding
FAD binding
thyroxine 5'-deiodinase activity
ATPase activity
damaged DNA binding
nucleic acid binding
0.0
0.5
1.0
1.5
2.0
2.5
3.0
3.5
4.0
log10 p-value

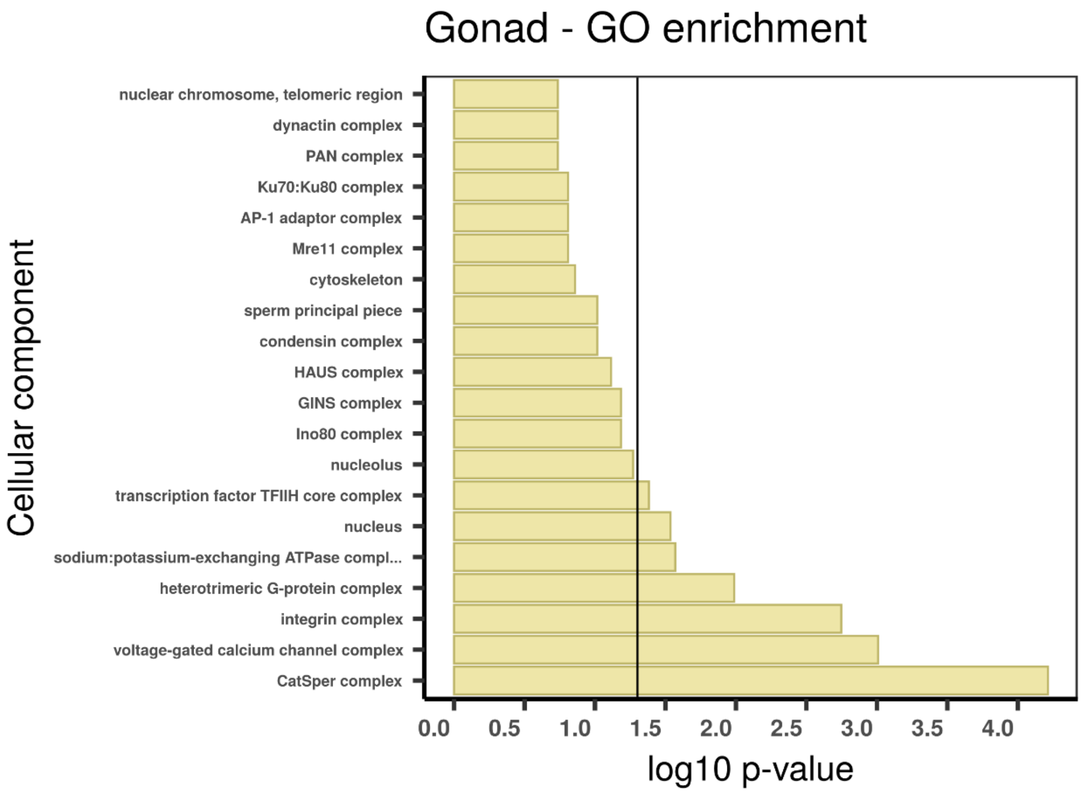

**Supplementary figure 60 | Gene set enrichment analysis with topGO using absolutely gonad specific TAU genes** Gene ontology (GO) enrichment analyses for absolutely female gonad specific TAU genes. The graphs correspond to the three domains of ontologies: biological process (BP), molecular function (MF) and cellular component (CC). The selected genes were analysed for enrichment in specific GO categories using the TopGO program against the background (all coding sequence genes). Y axis corresponds to enriched GO terms found in the respective domains (BP, MF and CC). X axis correspond to the log function of Fisher p-values obtained for each one of the enriched terms. The back line denotes a p-value = 0.05. P-values greater than 1,30 (log 0,05) indicate statistically significant enriched term. Genes involved in genome integrity are differentially expressed in female gonad tissue.

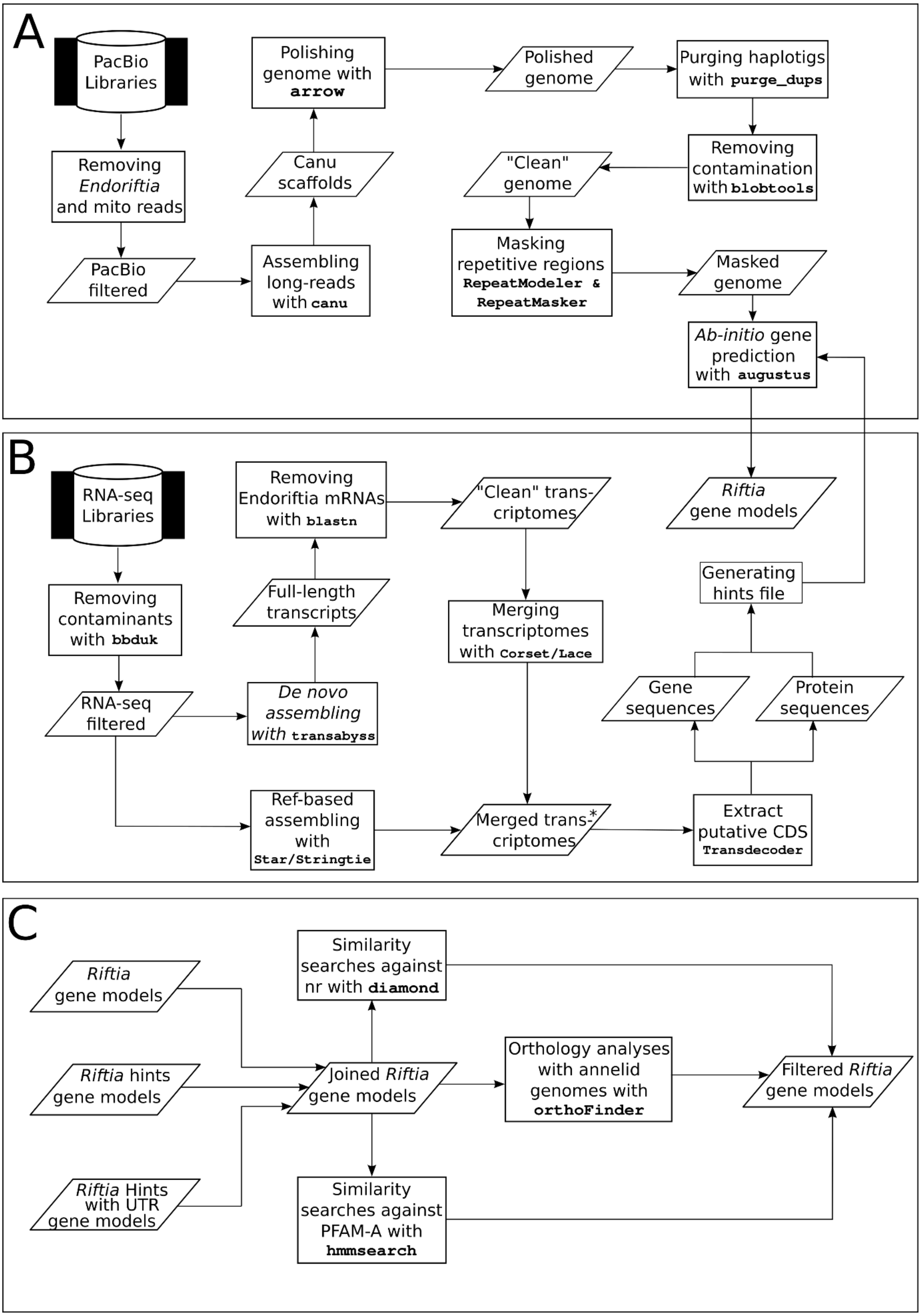

**Supplementary figure 61 | Bioinformatic workflow used in this study**. **A**, *Riftia pachyptila* genome pipeline for pre-processing, assembling and gene prediction. Mitochondrial and *Ca.* Endoriftia persephone reads were removed from the five PacBio libraries. The filtered long reads were assembled, polished and finally, the haplotigs purged from the final assembly. Contamination screening was performed, and the identification and masking of the repeat regions executed with RepeatModeler and RepeatMasker, respectively. Gene prediction with the polished, cleaned, soft-masked genome was carried out. **B**, *Riftia pachyptila* transcriptome pipeline. Illumina adaptors and low-quality bases were removed from the RNA-seq libraries. Pre-processed libraries were de-novo and reference-based assembled. Putative coding sequence regions were identified after the removal of contaminant transcripts. Hint files based on the transcriptome data were used to aid the gene prediction based on genome sequences. **C**, Gene models were joined and filtered based on homology searches, orthology identification and gene expression (not shown). *Two sets of merged transcriptomes were generated: one containing all de-novo transcriptomes, and other containing all reference-based transcriptome assemblies.

#### SUPPLEMENTARY NOTES

##### Supplementary note 1 | *Riftia* genome and transcriptome sequencing, assembling and assessment

We collected and then, using PacBio™ technology, sequenced the whole genome and eight tissue-specific transcriptomes of the giant tubeworm *Riftia pachyptila* (short *Riftia*) (Supplementary Figures 1-3; Supplementary Table 1). The long-read sequencing associated with the developed bioinformatic pipelines produced a highly contiguous and complete *Riftia pachyptila* draft genome. The results are comparable and, in some cases, surpass the overall quality (e.g., $N_{50}$, number of scaffolds) of other publicly available lophotrochozoan genomes (Simakov et al. 2013; Albertin et al. 2015; Luo et al. 2015; Sun et al. 2017; Wang et al. 2017; Luo et al. 2018; Belcaid et al. 2019; Calcino et al. 2019; Li et al. 2019; Sun et al. 2020). The assembled draft genome of the giant tubeworm was smaller than the predicted genome size studies (Dixon et al. 2001; Bonnivard et al. 2009) (Supplementary Figure 4). Specifically, the *Riftia* tubeworm genome is 128Mb smaller than the close relative *Lamellibrachia luymesi* (genome size of ~688Mb) (Li et al. 2019). To further validate the estimated genome size, a second tool called Flye (Kolmogorov et al. 2019), which implements a quite different long-read assembly algorithm compared to the primary assembler choice Canu (Koren et al., 2017), was employed. Additionally, a genome size estimation was performed with GenoneScope (http://qb.cshl.edu/genomescope/). The three independent analyses point to a giant tubeworm genome size ranging from ~510Mb to ~560Mb. Notably, though horizontal gene transfer in metazoan-bacteria endosymbiotic systems have been previously described (Nikoh and Nakabachi 2009; Husnik et al. 2013; Sloan et al. 2014; Ip et al. 2021), our analyses did not find any evidence of DNA transmission from *Candidatus* Endoriftia persephone to the giant tubeworm. However, more sensitive analyses should be employed (as described in Ip et al., 2021) to corroborate this hypothesis.

*Riftia pachyptila* genome contains the lowest repeat content among annelids with 29.9% of the genome composed of repeat regions (*Capitella*: 31%, *Helobdella*: 33%; *Lamellibrachia*: 36.92%) (Simakov et al. 2013; Li et al. 2019). As a matter of fact, *Riftia* repeat content is relatively smaller than most lophotrochozoans, being surpassed only by the scaly-foot snail *Chrysomallon squamiferum*, the brachiopod *Lingula anatina* and the limpet *Lottia gigantea* (25.5%, 22% and 21%, respectively) (Luo et al. 2015; Luo et al. 2018; Sun et al. 2020). Comparative genomic analyses showed different histories of repeat element expansions between the two tubeworms *Riftia* and *Lamellibrachia* (Li et al. 2019). The estimated proportion of repeat element classes varies significantly between the two annelids with *Lamellibrachia* showing a higher abundance of LINES, LTR and DNA elements. *Riftia* LTR and DNA repeat composition is more similar to the leech *Helobdella* and the polychaete *Capitella* than the closest relative *Lamellibrachia* suggesting an independent expansion of repeat elements in the *Lamellibrachia* lineage (Supplementary Figure 5).

With 25,984 protein coding genes and BUSCO4 (Simão et al. 2015) score of 99,37% (complete and partial) the giant tubeworm genome is the most complete annelid genome to date and ranks among the best lophotrochozoan genomes publicly available (Albertin et al., 2015; Belcaid et al., 2019; Calcino et al., 2019; Ip et al., 2021; Li et al., 2019; Luo et al., 2018, 2015; Simakov et al., 2013; Sun et al., 2020, 2017; Zhang et al., 2012). Compared to other animals that have an established symbiotic relationship with chemoautotrophic bacteria, *Riftia pachyptila* and *Chrysomallon squamiferum* (Sun et al. 2020) harbour the lowest number of coding sequence genes. The tubeworm *Lamellibrachia luymesi* (Li et al. 2019), and the bivalves *Bathymodiolus platifrons* and *Modiolus philippinarum* present between 33,584 and 38,998 coding sequence genes (Sun et al. 2017).

The number of transcripts obtained from the *de novo* assembled male (blood and sperm) and female (plume, vestimentum, trophosome, gonad, blood, and body wall) transcriptomes ranges between 122,284 (body wall - skin) and 279,949 (trophosome) (Supplementary Table 1D-F). Similarity searches with blastn (>90% identity) between the assembled transcriptomes and the genomes of *Riftia* and *Ca.* Endoriftia persephone identified the presence of endosymbiont reads across all tissues, with the majority located in the trophosome sample, as expected. The bacterial reads found in the non-trophosome tissues are certainly linked to cross-contamination during the sample extraction and collection.

The mapping rate for six out of eight assembled *de novo* transcriptomes against the unassembled transcriptome reads was higher than 95% attesting the quality of the *de novo* assemblies. Interestingly, the mapping rates for the trophosome and female blood tissues were ~62% and ~84% indicating misassemble errors and fragmentation during the *de novo* assembly procedure, since strict quality filtering were applied. This idea is further substantiated by the elevated number of Endoriftia reads in these two samples, which probably complicated the *de Bruijn* generation leading to the aforementioned problems.

An average of ~75% of the transcripts, with similarity against the *Riftia* reference genome, present in each of the eight individual *de novo* transcriptomes were annotated at a protein level against the nr database, suggesting novel uncharacterised genes present in the *Riftia* genome. A similar scenario was found in specific-tissue type transcriptomes of the scaly-foot snail *Chrysomallon squamiferum* (Sun et al. 2020). Whether these novel genes found in *Riftia* transcriptome tissues reflect the poor taxon coverage of closely related siboglinids/vestimentiferans in the protein database or lineage-specific genes, remains to be shown.

The reference-based transcriptome assemblies using the *Riftia* genome draft contain between 25,227 (skin – body wall) and 72,006 transcripts (trophosome) mirroring the trend obtained in the *de novo* reconstruction procedure. The elevated number of transcripts in the skin and trophosome compared to the other six tissues may indicate fragmentation of the reconstructed transcripts. The deep sequencing, tissue-specific transcriptomes and different gene prediction protocols enabled the identification of many alternative splice isoforms (average of ~2,2 isoforms per protein coding gene) in the *Riftia* genome.

**Supplementary note 2 | Developmental genes and signalling molecules**
During the early development, *Riftia* metatrochophore larvae are actively and specifically infected with *Ca*. Endoriftia persephone, which triggers the development of the trophosome and the reduction of the digestive system (Bright et al. 2013). The transient larval digestive system is divided into three distinct regions, foregut, midgut, and hindgut, with the foregut and hindgut formed by the ectoderm, whereas the midgut presents an endodermal origin (Jones and Gardiner 1989; Arendt et al. 2001). Many genes involved in the bilaterian gut development have been studied in annelids, such as *Platynereis*, *Capitella*, *Helobdella*, and *Hirudo medicinalis* (Rosa et al. 2005; Kulakova et al. 2007; Boyle and Seaver 2008; Hui et al. 2009). However, no available data focusing on the genes responsible for the development of the digestive tract are available for vestimentiferans. *Brachyury*, *goosecoid*, *fork head*, and the three Parahox genes *xlox*, *gsx* and *cdx* involved in the developing foregut, midgut and hindgut are present in the giant tubeworm genome. Gene expression quantification in *Riftia* showed little or no expression of these genes in the tubeworm adult tissues. The presence of *brachyury*, *fork head*, *goosecoid*, and all ParaHox genes in *Riftia* enable future comparative studies on the molecular mechanisms and evolution guiding the transient larval digestive system (Supplementary Figures 8-10)

Hox and ParaHox genes are two of the most investigated gene families in developmental biology (Supplementary Figures 8-9). They are remarkably conserved across invertebrates and vertebrates and are commonly organized in genomic clusters. By comparing the Hox gene distribution in Annelida, it can be inferred that the last common annelid ancestor had at least 11 Hox (*hox1-5, lox5*, *hox7*, *lox4*, *lox2*, *post1-2* (Kulakova et al. 2007; Simakov et al. 2013; Zwarycz et al. 2016). *Riftia* contains all Hox genes, except *hox7* (*ATNP)*. Our genomic screening did not identify *hox7*, *lox2* and *lox5* in the cold-seep tubeworm. Considering the high completeness of the *Lamellibrachia* genome (Li et al. 2019) (BUSCO score ~ 95%), possibly the Hox central class is disintegrated in the cold-seep tubeworm. Duplications and losses of Hox genes are a common feature in lophotrochozoans (e.g, *hox1-5* and *host-2* are duplicated in the nemertean *Notospermus geniculatus, hox2-4* is lost in the cephalopod *Octopus bimaculoides*) (Albertin et al., 2015; Luo et al., 2018). Within Annelida, duplications, and losses of Hox genes have also been reported. In the leech *Helobdella robusta*, *hox1* (*Lab*), *hox4* (*Dfd*), *hox5* (*Scr*) and *lox4* contains 2, 2, 4 and 2 copies, respectively, whereas *hox2* and *post1* are missing (Simakov et al. 2013). The polychaete annelids *Nereis virens* and *Capitella teleta* both contain the complete set, in contrast to *Platynereis dumerilii* (also a polychaete) which does not possess the *hox7* and *lox4* (Kulakova et al. 2007; Simakov et al. 2013). The *Riftia* genome does not show any sign of duplicated Hox and ParaHox genes, however the Hox-like elements *engrailed* (*En*) and *even-skipped* (*Eve*) present multi-copies in the giant tubeworm genome (two and four, respectively). Duplication of *en* is reported in the deep-vent snail *Chrysomallon squamiferum* (Sun et al. 2020). *Riftia* and *Lamellibrachia* genomes contain the three ParaHox genes. The Hox cluster in *Riftia* is almost intact, with only the *post1* located in a different scaffold. The presence of the almost complete Hox gene cluster and complement, as well as other developmental genes, attest for the quality and contiguity of the *Riftia* draft genome.

The transforming growth factor-β (TGFβ) family, responsible for cell fate specification and embryonic development, contains 33 members, including bone morphogenetic proteins (BMPs), growth and differentiation factors (GDFs), activins and *nodal* (Moustakas and Heldin 2009; Massagué 2012). *Riftia* genome contains 13 TGFβ members, including lefty, reported to be a deuterostome innovation (Simakov et al. 2015). *Lefty* has been reported to be present in the nemertean *Notospermus* and the brachiopod *Lingula* (Luo et al. 2018), our analyses found its presence, in addition to *Riftia*, in the *Lamellibrachia* and *Capitella* genomes (Supplementary Figure 11). The domain composition of *lefty* in all lophotrochozoans differs from the deuterostome ortholog, with only the presence of TGF_propeptide protein domain (PF00688). The *lefty* gene models on *Riftia*, *Lamellibrachia* and *Capitella* are complete and not fragmented as previously reported in the *Notospermus* and *Lingula* (Luo et al. 2018), suggesting that the TGF_beta domain (PF00019) is indeed not present in Lophotrochozoa. Broader taxon sampling and comparative genomics are required to elucidate whether both domains were present in the last common bilaterian ancestor.

The giant tubeworm genome contains a single copy of hedgehog and notch ligands, whereas the close relative *Lamellibrachia luymesi* contains two and one copy, respectively (Supplementary Figures 12-13). Additionally, our gene screening and phylogenetic analysis identified a single copy of hedgehog and notch ligands in the polychaete *Capitella* and the leech *Helobdella robusta*, corroborating previous studies on these annelid model species (Kang et al. 2003; Rivera et al. 2005; Seaver and Kaneshige 2006; Thamm and Seaver 2008). Overall, numbers of hedgehog and notch ligands in annelids are smaller than other lophotrochozoans, such as nermerteans, phoronids and molluscs. *Phoronis australis* contains, for example, 13 hedgehog and three notch ligand homologs (Luo et al. 2018). An expansion of the Hedgehog receptor dispatched, a multipass membrane protein that facilitates the transport of cholesterol modified Hedgehog (Burke et al. 1999), was identified in the Vestimentifera lineage compared to the other two annelids herein analysed.

The Wnt gene family is involved and cell fate specification and regulation of posterior growth during early animal embryogenesis (Niehrs 2012). The giant tubeworm, *Lamellibrachia* and *Capitella* genomes have all the 12 expected lophotrochozoan Wnt genes (*wnt-A*, *wnt1*, *wnt2*, *wnt4-11*, *wnt16*), indicating that the last common annelid ancestor harboured the full Wnt lophotrochozoan complement (Cho et al. 2010; Luo et al. 2018). *Wnt3* is lost in the Protostomia lineage (Holstein 2012). *Lamellibrachia luymesi* genome present two lineage-independent duplications of *wnt7* and *wnt1*. Independent losses (*wnt-A*, *wnt9-10*) and duplications of Wnt subfamilies are also found in *Helobdella robusta* (*wnt5*, *wnt11* and *wnt16*), as previously reported (Cho et al. 2010). We identified four Wnt-receptor Frizzled subfamilies in *Riftia*, with *fz-4* containing three paralogous copies in the giant tubeworm genome. The Wnt antagonist *sFRP3/4* (Cruciat and Niehrs 2013), despite being identified in lophotrochozoans (Luo et al. 2018), is missing from *Riftia* genome (Supplementary Figures 14-15).

**Supplementary note 3 | Orthology and gene family analyses**

The orthology analyses using 36 metazoan taxa (4 non-bilaterians, 19 lophotrochozoans, 6 ecdysozoans, 7 deuterostomes – Supplementary Table 2) showed that *Riftia* and *Lamellibrachia* present the highest number of orphans orthogroups within the annelid taxa herein analysed (*Riftia*: 8,132; *Lamellibrachia*: 10,262; *Capitella*: 4,300; *Helobdella*: 4,821). In agreement with recent studies (Fernández and Gabaldón 2020), the distribution of metazoan, bilaterian, protostomian, lophotrochozoan and lineage-specific orthogroups within Annelida follows a bimodal distribution, with a massive gain of genes at the deep (i.e., last common metazoan ancestor) and recent nodes (i.e., taxonomically restricted genes).

Analyses with topGO using *Lamellibrachia* and *Riftia* lineage-specific genes revealed different patterns of gene enrichment (Supplementary Figures 21-24; Supplementary Table 5). We found significant enrichment ($p < 0.05$) of terms involved in chitin synthesis and secretion in the *Riftia* genome. Chitin is the major component of the tube of *Riftia* and the most abundant biopolymer in nature (Zakrzewski et al. 2014). *Riftia* alone, during the development of its tube, is responsible for the highest production of chitin recorded in marine environments, producing ~100 times more than any other marine animal (Shillito et al. 1995; Gaill et al. 1997). Produced by specialised large multi-cellular gland cells in the plume, trunk and opisthosome region, the tubes of *Riftia* are highly stable structures, with a lower degradation rates than other vent animals, such as crabs (Shillito et al. 1993; Shillito et al. 1995; Ravaux et al. 2003). The expanded complement of chitin-related genes in *Riftia* further corroborates the important role of this complex macromolecule against toxic vent chemicals and predators in the vent ecosystem (Shillito et al. 1993; Shillito et al. 1995; Ravaux et al. 2000; Ravaux et al. 2003). These results open the possibility of comparative studies focused on the exoskeleton protein synthesis and secretion pathways on *Riftia* and other vent animals.

Among the lineage-specific genes in *Lamellibrachia*, we found enriched terms associated with G protein-coupled receptor (GPCRs) (Supplementary Table 6). GPCRs are evolutionary conserved protein families that trigger signal transduction pathways having important roles in physiological responses to hormones, neurotransmitters, and environmental inputs (Fredriksson and Schiöth 2005). Additionally, PFAM protein domain analysis showed that the complement size of GPCR (7tm_1; PF00001) proteins in *Lamellibrachia* greatly surpasses other (Albertin et al., 2015; Simakov et al., 2013) (*Lamellibrachia luymesi*: 631; *Octopus bimaculoides*: 328; *Lottia gigantea*: 267). As a matter of fact, Fisher's exact test showed that GPCR-containing proteins are expanded ($p < 0.05$) in the cold-seep tubeworm in relation to *Riftia pachyptila* (308). The presence of many lineage-specific GPCRs in *Lamellibrachia* raises interesting questions about the evolutionary history of this large repertoire and its association with sensory information and homeostatic regulation in the cold-seep tubeworm. Our PFAM protein domain analysis indicated that *Capitella teleta* genome harbours the highest number of GPCR genes in lophotrochozoans (963), results corroborated by previous studies (Simakov et al. 2013; Albertin et al. 2015; Ritschard et al. 2019). Further fine-grained analyses on the evolution of GPCR families are necessary to elucidate patterns of gene gains and losses in the different nodes of the Annelida tree.

PFAM enrichment analysis in the *Riftia* genome showed the expansion of protein domains present in high-molecular mass proteins such as collagen, laminin, nidogen (Supplementary Table 6). Clustering analysis with the expanded proteins involved in the production of the basement membrane in *Riftia* showed that these proteins are all connected to each other, indicating evolutionary relatedness (Supplementary Figure 24). However, due the presence of many tandem repeats within these proteins, such as EGF, the clustering probably reflects the reoccurring low complexity repeats in these proteins.

Pairwise comparisons of the enriched/contracted PFAM terms present in the genomes of symbiotic deep-sea animals, *Riftia pachyptila*, *Lamellibrachia luymesi*, *Bathymodiolus platifrons* and *Chrysomallon squamiferum*, against selected lophotrochozoans (N = 14; Supplementary Table 6; Supplementary Figure 24A) identified few shared protein domains. Among them, the zinc-finger H2C2/Integrase H2C2 DNA-binding domains (*Riftia*: 0/127; *Lamellibrachia*: 4/115; *Bathymodiolus*: 9/136 *Chrysomallon*: 1/99) and peptidase A17 (domain not found in any symbiotic deep-sea animal). Interestingly, C1q domain-containing proteins, regarded as key players in the innate immunity and pathogen recognition (Thielens et al. 2017), are contracted in *Riftia* (2), *Lamellibrachia* (3) and *Chrysomallon* (6) in relation to other lophotrochozoans (average = 69). The genome of *Bathymodiolus*, in contrast to the other deep-vent symbiotic animals, contains an expansion of C1q domains (386), as reported in the pacific oyster *Crassostrea gigas* and in the Mediterranean mussel *Mytilus galloprovincialis* (Gerdol et al. 2011; Gerdol et al. 2015; Gerdol et al. 2019).

The *Riftia* genome contains the smallest number of transcription factors (TFs) within Annelida with a complement of 414 genes (*Lamellibrachia luymesi*: 423; *Capitella teleta*: 551; *Helobdella robusta*: 568) (Supplementary Table 3; Supplementary Figure 19). The nuclear factor zf-C4, bHLH (helix-loop-helix) and the zinc finger C2H2 domain containing genes are underrepresented in the giant tubeworm genome in comparison to the other three annelid genomes. We did not identify any notable TF lineage-specific expansion in *Riftia*. Our analyses retrieved the exact same number of homeobox domain containing genes in *Riftia* and *Lamellibrachia* (122 genes), making the homebox complement in the vestimentiferans smaller than the other two annelids. A recent study on the homebox complement of the oligochaete *Eisenia fetida* showed that the earthworm experienced many gene gains events totaling 363 homebox genes, surpassing the numbers in many deuterostomes, ecdysozoans, lophotrochozoans and annelids (Zwarycz et al. 2016). The homeobox complement herein reported is in agreement with a previous study (Zwarycz et al. 2016) (*Capitella*: 178 / our study 160; *Helobdella*: 270 / our study 246), showing the robustness of our methods.

Loss and gains of TFs have been reported across many metazoan lineages, and as two examples we could cite the cephalopod *Octopus bimaculoides* and the parasite tapeworms (Tsai et al. 2013; Albertin et al. 2015). The first shows a massive expansion of the C2H2 superfamily of zinc-fingers transcription factors, which regulate neuronal development, and the later a broad reduction of the homebox complement. These changes in the TF complement sizes are related to the large and complex nervous system and adaptation to parasitic lifestyle in the cephalopod and tubeworm, respectively. More thorough investigations are required to establish if the reduced TF numbers in *Riftia* and *Lamellibrachia* is linked to the loss of anatomical structures in these two tubeworms.

One unexpected result derived from CAFE and PFAM analyses was the contraction of the histone gene families in the giant tubeworm genome in relation to other lophotrochozoans (N = 18) (Supplementary Tables 6). As histones are essential to the transcription machinery and condensation of eukaryotic DNA, their absence from *Riftia* genome caught our attention. Furthermore, few studies indicate their presence in the giant tubeworm (Rouse et al. 2015; Hinzke et al. 2019), challenging our initial results. To investigate this question, we performed additional local similarity searches with hmmscan (Mistry et al. 2013) using the histone HMM profile as query (PF00125) and the proteomes obtained from the de-novo transcriptomes as databases. Our similarity searches identified many copies of the core histones H2B, H2A and H3 in the *Riftia* tissues (vestimentum: 16; body wall (skin): 16; male blood:17; sperm: 18; female gonad: 17; female plume:15). Subsequent multiple sequence alignments, homology searches and phylogenetic inferences with the non-redundant histones set confirmed their presence in the giant tubeworm genome. The initial erroneous genome annotation of histone genes could be attributed to the over masking of repeat regions overlapping the histone coding sequence regions (Mario Stanke personal communication, May 2020).

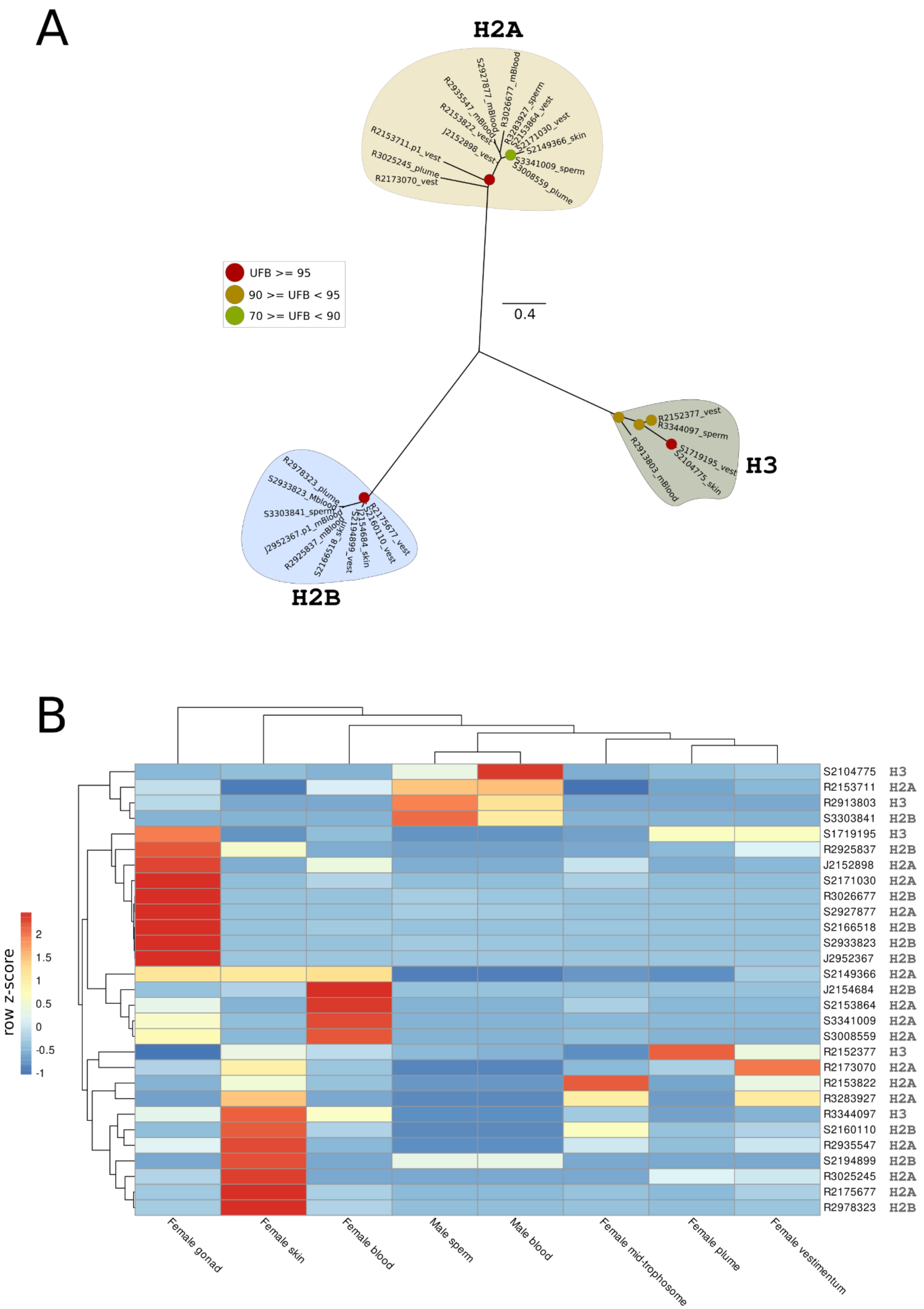

**Additional supplementary figure 1 – Histone phylogeny and gene expression analysis. A,** Unrooted maximum-likelihood phylogenetic inference of the non-redundant core histone set (i.e., H2A, H2B and H3) found in different *de novo* transcriptomes. Leaf names correspond the transcript name followed by the tissue in which they were found. The branch support values are represented by the coloured circles in the tree nodes. Red circles represent ultrafast bootstrap values >= 95. Yellow circles represent ultrafast bootstrap values >= 90 and < 95. Green circles represent ultrafast bootstrap values < 90 and >= 70. Ultrafast bootstrap values smaller than 70 are not shown. **B**, Expression profile of histone genes *Riftia pachyptila*. colour coding reflects the expression patterns based on row Z-score calculations. Despite not identified in the genome prediction, histone genes, as expected, are present in the giant tubeworm genome. Histones are highly expressed on gonad and body wall (skin) tissues.

Hinzke et al. (2019) speculated that histones may have antimicrobial properties functioning as mediators of host-symbiont interactions on the trophosome and defence against environmental microbes on the plume. This hypothesis was based on previous observations that histones and histone-derived peptides modulate immune system responses in vertebrates (Fr et al. 1998; Park et al. 1998; Cho et al. 2009; Bishop et al. 2017). Contrary to the results described by Hinzke et al.(2019), we did not identify a relative high abundance of histone genes in the trophosome compared to other tissues, challenging their findings. We found that histone genes are constitutively expressed on *Riftia* tissues, with the gonad and skin presenting an elevated number of highly expressed histone genes. Expression of core histones is required during the S phase of cell-division cycle where the DNA replication commences (Mei et al. 2017). The expression of histones in the aforementioned *Riftia* tissues probably reflects the high levels of cell proliferation, especially in the skin as shown in immunohistochemical and ultrastructural cell cycle studies (Pflugfelder et al. 2009).

The expanded gene families identified by CAFE point to molecular adaptations associated with sulphide rich environments and symbiotic lifestyle in *Riftia* (Supplementary Table 4; Supplementary Figure 20). The toxicity of $H_2S$ is primarily linked to the inhibition of the cytochrome-c oxidase in the mitochondrial respiratory chain, which limits most animals to survive and reproduce in sulphide rich environments (Cooper and Brown 2008). *Riftia* has many expanded families enriched with GO terms associated with sulphur metabolism and detoxification (e.g., carbohydrate-, galactose- and heparan-sulphate-sulfotransferases). Recent comparative transcriptome and differential gene expression analyses with the shrimp *Rimicaris* sp. (Zhang et al. 2017), predominant megafauna of deep-sea hydrothermal vents, recovered a similar gene toolkit responsible for detoxification in the arthropod, indicating similar molecular mechanisms in different animals that live in similar sulphide-rich vent ecosystem.

Three expanded rapidly evolving gene families (i.e., positively selected) were detected in the *Riftia* genome: sushi-domain, glycosyltransferase, and mucin-related families (Supplementary Figure 4). Host sushi-domain proteins have been hypothesized to present symbiosis specific roles in sponges and cephalopods (Collins et al. 2012; Pita et al. 2018). Sushi domains are involved with innate immunity and self/non-self-recognition (Kirkitadze and Barlow 2001). The expansion of this gene family in *Riftia*, as well as *Lamellibrachia* (Li et al. 2019), could be associated with the recognition of the endosymbionts by the tubeworm hosts (Supplementary Figure 27). Furthermore, the expansion of glycosyltransferases is probably coupled with the enriched number of genes in *Riftia* involved in the formation of the basement membranes, such as collagen. Finally, independent expansions of mucin-related genes have been identified in other lophotrochozoans, such as phoronids and brachiopods (Luo et al. 2018). In *Lingula* and *Phoronis*, mucin genes are highly expressed in the lophophores and in the phoronid lineage they might be related to protection against predators. The function of mucus production in *Riftia* adults is less clear. However, during the symbiont infection model suggested by Nussbaumer et al., (2006), the metatrocophore larvae actively secrete a mucous coat during settlement in which the environmental free-living bacteria attach, including the *Riftia* endosymbiont. Interestingly, gene family analysis in *Lamellibrachia* recovered the same expansion of mucin-related genes (Li et al. 2019). Based on these observations, we suggest that the expanded mucin-related proteins in *Riftia* might play a role in the acquisition of the free-living endosymbionts from the environment during the larval settlement, scenario that could be generalised to *Lamellibrachia*.

Vestimentiferan specific genes present are involved in a range of biological processes, including oxygen transport, protein dephosphorylation, G protein-coupled receptor signalling pathway and cyclic nucleotide biosynthetic process.

**Supplementary note 4 | Haemoglobin evolution**

*Riftia* possesses three complex multimeric haemoglobins (HBs), two of them dissolved in vascular blood (V1 and V2) and one in the coelomic fluid (C1) (Zal et al., 1996a, 1996a). The Hb complement of the giant tubeworm has been suggested to contain six distinct genes, from which three are classified as β1-Hb, and the remaining three belong to the paralog groups α1-, α2- and β2-Hbs(Bailly et al., 2002; Zal et al., 1997, 1996a). To further investigate the multigenic extracellular globin family in *Riftia*, we retrieved all Hb genes from the giant tubeworm genome and processed them through a phylogenetic pipeline. Using a large reference dataset (Belato et al., 2019), tree inferences and multiple sequence alignments we identified and annotated 26 extracellular Hbs in *Riftia* (Supplementary Figures 21 and 28; Figure 3). The paralog group β1-Hb is expanded on the giant tubeworm, whereas α1- and β2-Hbs groups contain only one copy each. The α2-Hb group contains two paralogous genes. The Hb complement of *Riftia* mirrored the complement of the cold-seep tubeworm *Lamellibrachia luymesi* (Li et al., 2019), suggesting an expansion of the β1-Hbs already at the base of Vestimentifera.

We propose a division of the β1-Hb chains into eight paralogous groups enriching the established classification of Baily et al (Bailly et al., 2002). In addition to the previous β1a-Hb, β1b-Hb and β1c-Hb groups, five more groups were recognised through phylogenetic analyses: β1d- to β1h-Hb. Seven out of the eight β1-Hb paralogous groups have recognisable orthologs in *Lamellibrachia luymesi*. In agreement with previous studies, the 12 conserved amino acid residues located in extracellular globins were present in the giant tubeworm and *Lamellibrachia* Hb genes (Belato et al., 2019; Gotoh et al., 1987; Li et al., 2019; Negrisolo et al., 2001; Shishikura et al., 1986; Yuasa et al., 1996) (Supplementary Figure 29) β1-Hb group members harbour a conserved motif, VNV[ADE] at positions 48-51 (considering the Cys-19, as the first residue of the alignment block), as previously reported (Bailly et al., 2002) (although variations were observed in *Riftia* and *Lamellibrachia* representatives belonging to β1d-, β1e-, β1h- and β1g-Hb groups). The residues 74-, 122- and 128-Ans, as well as 83-Glu, are relatively conserved in β-Hb groups. Free cysteine residues, associated with the sulphide binding function on vestimentiferans and other annelids living in permanent sulphide-rich environments (Bailly et al., 2002; Li et al., 2019; Pallavicini et al., 2001; Suzuki et al., 1995, 1989; Takagi et al., 1991; Yuasa et al., 1996; Zai et al., 1999; Zal et al., 1998, 1997), have been previously identified on the giant tubeworm α2- and β2-Hb chains. We found additional putative cysteine residues in β1e, β1f and β1g-Hb groups, totalling new seven Hb genes with the capability of carry H2S on the giant tubeworm. Our analyses revealed eight copies of *Lamellibrachia luymesi* genes containing putative free cysteine residues, as reported by Li et al (Li et al., 2019). Homology model generation reconfirmed the presence of free-cysteine residues, indicating the possible sulphide-binding capability of some members of the expanded tubeworm β1-Hb chain (Supplementary Figure 30). Comparisons between *Riftia* β1-Hb containing proteins and the two highest ranked templates (3WCT_D - deoxygenated haemoglobin from the tubeworm *Lamellibrachia satsuma*; 1YHU_C C1 haemoglobin from the tubeworm *Riftia pachyptila*) showed sequence similarity values ranging from 35.57% - 48.30% and 39.42% - 54.29%, respectively.

Tandem gene duplication, which is the main driving force behind the rise of multigene families, is an important mutational process in evolutionary adaptation and the evolution of eukaryotic genomes (Friedman and Hughes, 2001; Loehlin and Carroll, 2016). In lophotrochozoans, lineage-specific expansions through tandem duplications related to defence mechanisms, immune responses to pathogens and neuronal development, have been reported in nemerteans, phoronids and cephalopods, respectively (Albertin et al., 2015; Luo et al., 2018). In *Riftia*, the close chromosomal proximity of the expanded β1 chain genes and their phylogenetic relationship indicate that this multigenic family was originated through a series of tandem duplications. We identified seven genomic Hb-containing clusters on the giant tubeworm genome, from which six correspond to the β1 chain genes (Fig 3B). The position of some Hb genes in the end of scaffolds and their nested phylogetic pattern revealed in the tree inference, point to a more contiguous cluster containing the Hb chains. Additional sequencing (e.g., Hi-C method) and subsequent genome scaffolding are necessary to confidently elucidate the exact chromosomal organisation of the Hb complement in the *Riftia* genome.

Deep-sea hydrothermal vents are unstable environments marked by small-scale disturbances of the physicochemical conditions (Bright and Lallier 2010). The changes in pH, sulphide, oxygen, and temperature pose a challenge for the vent fauna, that must cope with these variations to thrive under these unpredictable conditions. As *Riftia* is extremely dependent upon its endosymbionts, the trophosome needs to be constantly nurtured with the necessary metabolites (e.g., carbon dioxide, sulphide, and oxygen) required from the bacteria to fix carbon through autotrophic pathways (Fisher et al. 1989). The transport of $H_2S$ and $O_2$ substrates to the trophosome is carried out through the circulatory system and multimeric haemoglobin complexes. As expected, we found that representatives of all four Hb groups (α1-2, β1-2) are highly expressed in the trophosome tissue. Notably, gene expression quantification using publicly available transcriptome datasets (Hinzke et al. 2019) obtained from sulphide rich, medium, and depleted trophosome samples showed great variation of Hb expression. Many members of expanded β1 group are highly expressed on medium and sulphur-rich trophosome samples (β1a, b, c, d, e, f, g), including the Hbs containing the free-cysteine residues. The presence of highly expressed β1 genes in the trophosome has also been reported in the cold-seep tubeworm *Lamellibrachia luymesi* (Li et al. 2019). Despite the clear distinct expression patterns of the Hbs in the different trophosome samples, it is hard to associate these changes in expression levels with the exact environmental conditions which they were sampled. The qualitative classification of depleted, medium, and sulphur-rich employed by Hinzke et al. (2019) was based on the trophosome colour, which is directly linked to presence of elemental sulphur vesicles in the endosymbionts. As shown by previous studies (Fisher et al. 1988; Wilmot and Vetter 1990; Pflugfelder et al. 2005), variability of colours may occur in the same trophosome, as well as differences in metabolism of the distinct endosymbiont subpopulations within a single animal (Hinzke et al. 2021), making it difficult to clearly state the physicochemical conditions from which the trophosome samples were obtained. However, irrespective of the exact environment conditions, we could observe clearly distinct patterns of Hb gene expression, indicating a more specialised role of these genes in *Riftia*. The ubiquitous expression of the six linker genes in the sulphur-rich, medium, and depleted samples is consistent with the molecular organisation of the Hb complexes in *Riftia*, in which the linker genes act as structural non-globin component of the multimeric haemoglobin. We were unable to recover the monophyly of the linker groups, corroborating previous studies (Belato et al. 2019).

**Supplementary note 5 | Comparative tissue-specific transcriptomics and gene expression**

Comparative transcriptome analysis of eight different *Riftia* tissues (plume, vestimentum, trophosome, body wall (skin), female gonad, sperm, and male/female blood) revealed distinct transcriptional landscapes (Supplementary Table 9). The vestimentum and body wall tissues (Supplementary Figures 25 e 26) harbour several tissue specific genes (TSGs) mainly involved in chitin metabolism responsible for the tube production and growth (Gaill et al. 1992; Shillito et al. 1993; Shillito et al. 1995). These results are not surprising, since these two organs contain conspicuous pyriform glands responsible for the chitin production and degradation (Jones 1981; Bright and Lallier 2010).

The branchial plume contains many TSGs involved with cell division and growth, signal transduction, immune system, and apoptosis. We also identified a high expression of immune response, endosomal, cell cycle, autophagy, and (anti-) apoptotic proteins in the plume (discussed into more details in the next section).

The female gonad tissue is enriched with proteins involved in genome integrity (e.g., DNA repair and damage checkpoint, telomerase maintenance and nucleotide excision repair), an important factor for subsequent internal fertilisation and healthy offspring (Additional supplementary figure 2). Interestingly, we found many TSGs in the female gonad tissue associated with methyltransferase activity indicating potential epigenetic regulation in the tubeworm female gonadal development. The sperm tissue is characterised by genes enriched with ion transmembrane transport functions, such as CatSper (Rahban and Nef 2020), which has been shown essential to mammalian sperm flagellum mobility, chemotaxis towards the egg, capacitation, and acrosome reaction (Brown et al. 2019).

The trophosome tissue is characterised by many TSGs involved with mitochondrial activity (Supplementary Figure 33) (e.g., oxidoreductase activity, mitochondrial fission, mitochondrial ribosomal translation), beta oxidation of fatty acids, oxygen transport (e.g., oxygen biding), haemoglobin metabolism (e.g., iron ion binding, iron-sulphur cluster biding, heme biding, porphyrin and tetrapyrrole metabolism, 5-aminolevulinate synthase), lysosomal activity (e.g., threonine-type endopeptidase activity, peptidyl-dipeptidase activity), protein translation (e.g., ribosome, structural constituent of ribosome), and urea cycle (e.g., hydroxyisourate hydrolase activity).

The presence of TSGs in the trophosome related to 5-aminolevulinate synthase, porphyrin metabolism, and metal ion biding indicates that this tissue harbours the enzymatic machinery necessary for haem biosynthesis. Haem is an integral part of haemoglobin molecules, which is synthesized in a multistep pathway that begins and ends in the mitochondrion. The process starts with the formation of δ-aminolevulinic acid (ALA) catalysed by the enzyme 5-aminolevulinate synthase from glycine and succinyl-CoA (Malik and Djaldetti 1979). ALA migrates to the cytoplasm and through a series of enzymatic reactions involving seven universally conserved enzymes it is converted into haem. The last two steps mediated by the enzymes protoporphyrinogen oxidase (PPOX) and ferrochelatase (FeCH), which are responsible for the dehydrogenation of trotoporphyrinogen IX and its chelation with iron to produce the haem molecule, is located inside the mitochondria (Supplementary Figure 40) (Ajioka et al. 2006; Kořený et al. 2013; Celis and DuBois 2019).

TSGs belonging to the mitochondrial carrier family were found in the trophosome, including the mitochondrial coenzyme a transporter slc25a42, solute carrier family 25 (carnitine/acylcarnitine translocase member 20), mitochondrial ornithine transporter; and tricarboxylate transport protein (Additional supplementary figure 4; Supplementary Table 11). Mitochondrial carriers are widespread in eukaryotic organisms and lineage-specific variations have been reported (e.g., 35, 58 and 67 mitochondrial carriers are reported in the yeast *Saccharomyces cerevisae,* the plant *Arabidopsis thaliana* and in humans, respectively) (Wohlrab 2006; Monné et al. 2015). Similarity searches using the PFAM model PF00153 (mitochondrial carrier protein domain) against the giant tubeworm genome identified the presence of 45 mitochondrial carriers in *Riftia* (Additional supplementary figure 4). The giant tubeworm mitochondrial carriers are involved in the transport of different substrates such as amino acids, coenzymes, and nucleotides.

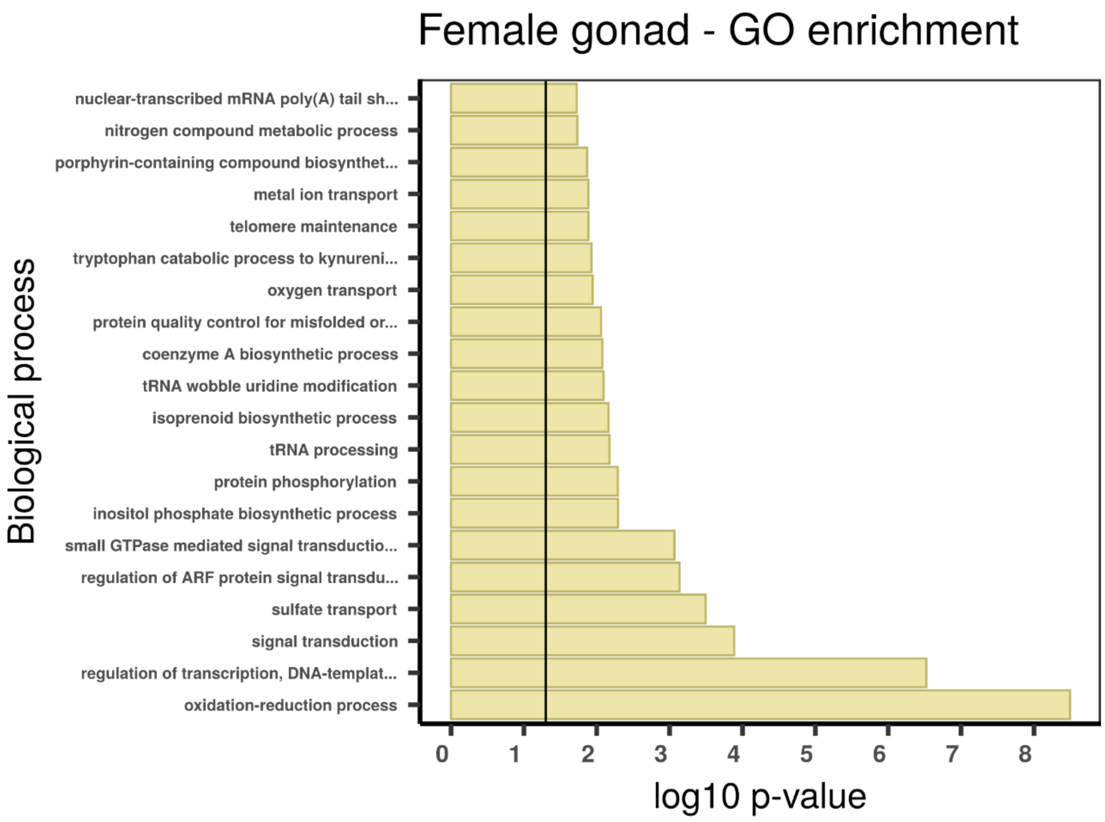

Female gonad - GO enrichment
Biological process
nuclear-transcribed mRNA poly(A) tail sh...
nitrogen compound metabolic process
porphyrin-containing compound biosynthet...
metal ion transport
telomere maintenance
tryptophan catabolic process to kynureni...
oxygen transport
protein quality control for misfolded or...
coenzyme A biosynthetic process
tRNA wobble uridine modification
isoprenoid biosynthetic process
tRNA processing
protein phosphorylation
inositol phosphate biosynthetic process
small GTPase mediated signal transductio...
regulation of ARF protein signal transdu...
sulfate transport
signal transduction
regulation of transcription, DNA-templat...
oxidation-reduction process
0 1 2 3 4 5 6 7 8
log10 p-value

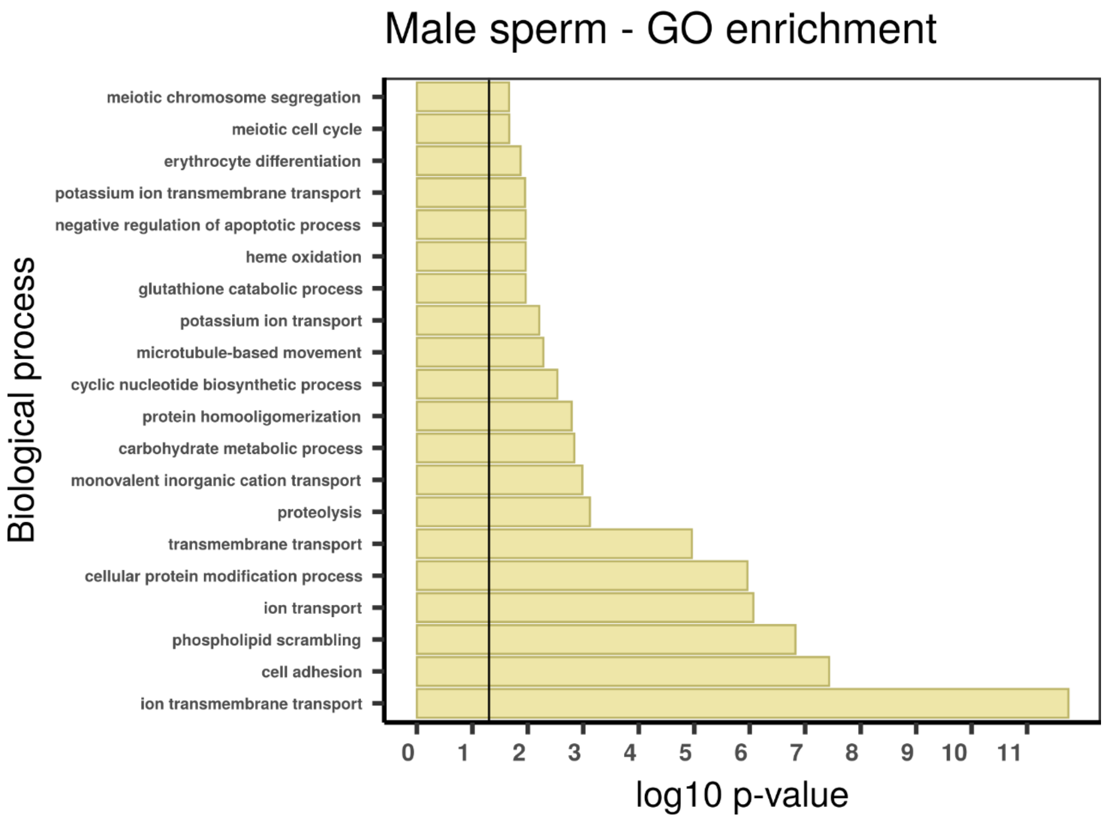

Male sperm - GO enrichment
Biological process
meiotic chromosome segregation
meiotic cell cycle
erythrocyte differentiation
potassium ion transmembrane transport
negative regulation of apoptotic process
heme oxidation
glutathione catabolic process
potassium ion transport
microtubule-based movement
cyclic nucleotide biosynthetic process
protein homooligomerization
carbohydrate metabolic process
monovalent inorganic cation transport
proteolysis
transmembrane transport
cellular protein modification process
ion transport
phospholipid scrambling
cell adhesion
ion transmembrane transport
0 1 2 3 4 5 6 7 8 9 10 11
log10 p-value

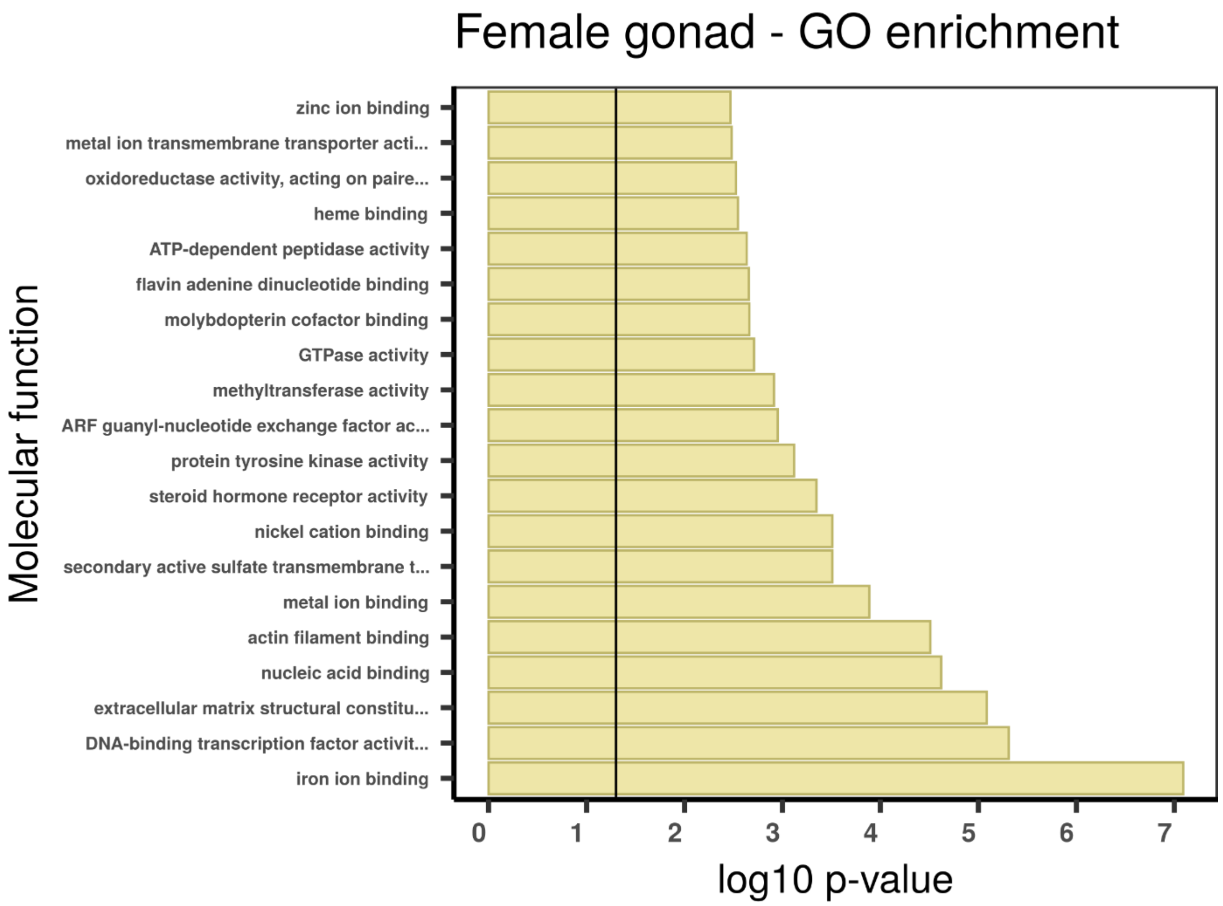

Female gonad - GO enrichment
Molecular function
zinc ion binding
metal ion transmembrane transporter acti...
oxidoreductase activity, acting on paire...
heme binding
ATP-dependent peptidase activity
flavin adenine dinucleotide binding
molybdopterin cofactor binding
GTPase activity
methyltransferase activity
ARF guanyl-nucleotide exchange factor ac...
protein tyrosine kinase activity
steroid hormone receptor activity
nickel cation binding
secondary active sulfate transmembrane t...
metal ion binding
actin filament binding
nucleic acid binding
extracellular matrix structural constitu...
DNA-binding transcription factor activit...
iron ion binding
0 1 2 3 4 5 6 7
log10 p-value

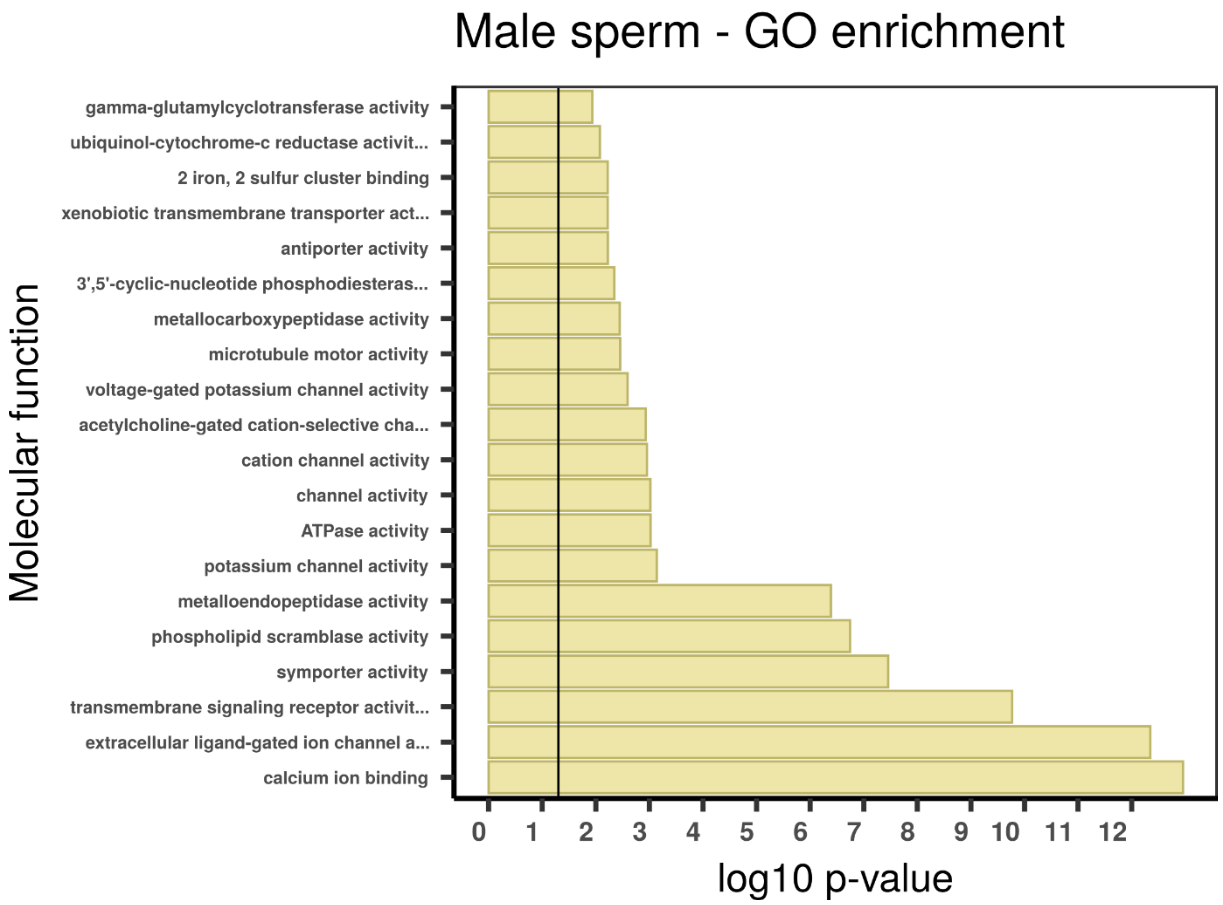

Male sperm - GO enrichment
Molecular function
gamma-glutamylcyclotransferase activity
ubiquinol-cytochrome-c reductase activit...
2 iron, 2 sulfur cluster binding
xenobiotic transmembrane transporter act...
antiporter activity
3',5'-cyclic-nucleotide phosphodiesteras...
metallocarboxypeptidase activity
microtubule motor activity
voltage-gated potassium channel activity
acetylcholine-gated cation-selective cha...
cation channel activity
channel activity
ATPase activity
potassium channel activity
metalloendopeptidase activity
phospholipid scramblase activity
symporter activity
transmembrane signaling receptor activit...
extracellular ligand-gated ion channel a...
calcium ion binding
0 1 2 3 4 5 6 7 8 9 10 11 12
log10 p-value

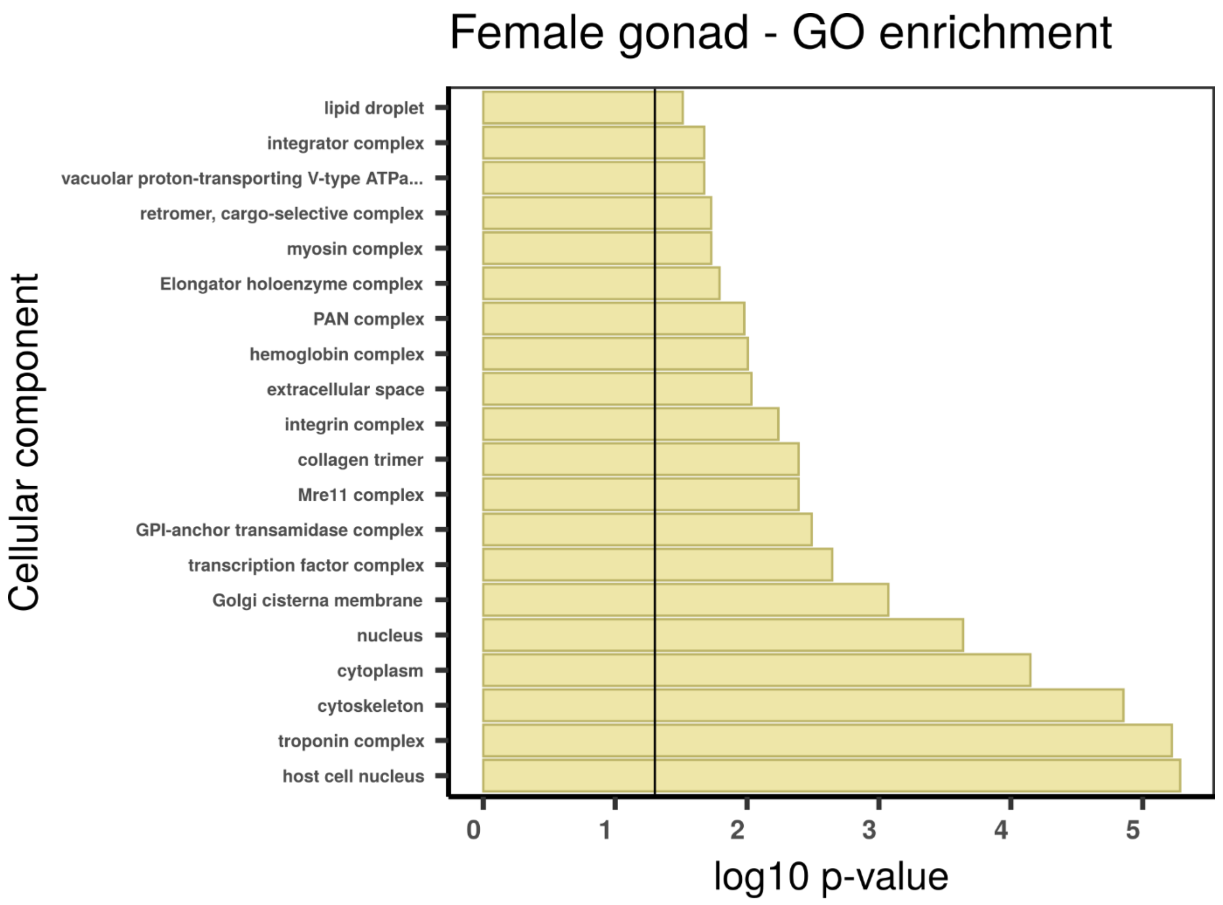
Female gonad - GO enrichment
Cellular component
lipid droplet
integrator complex
vacuolar proton-transporting V-type ATPa...
retromer, cargo-selective complex
myosin complex
Elongator holoenzyme complex
PAN complex
hemoglobin complex
extracellular space
integrin complex
collagen trimer
Mre11 complex
GPI-anchor transamidase complex
transcription factor complex
Golgi cisterna membrane
nucleus
cytoplasm
cytoskeleton
troponin complex
host cell nucleus
0
1
2
3
4
5
log10 p-value

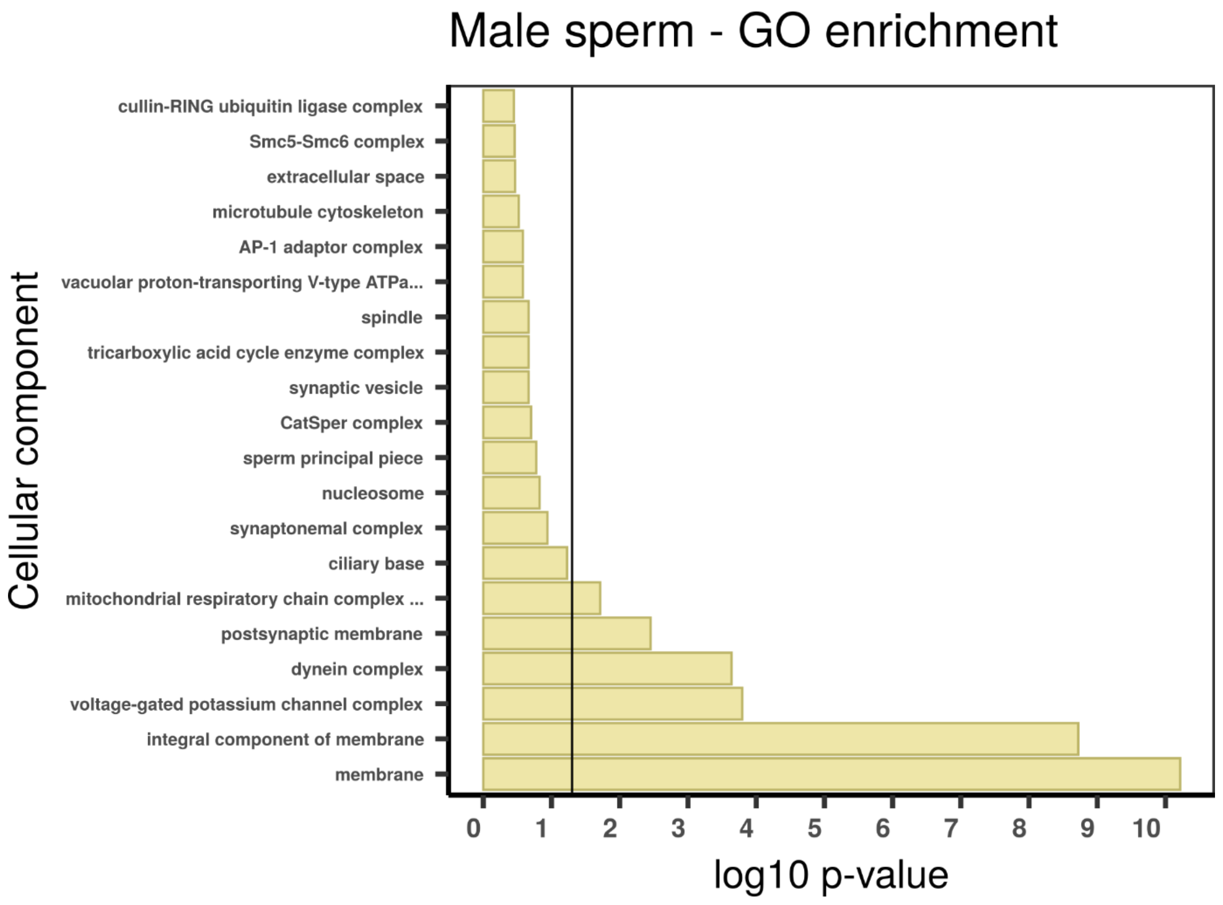
Male sperm - GO enrichment
Cellular component
cullin-RING ubiquitin ligase complex
Smc5-Smc6 complex
extracellular space
microtubule cytoskeleton
AP-1 adaptor complex
vacuolar proton-transporting V-type ATPa...
spindle
tricarboxylic acid cycle enzyme complex
synaptic vesicle
CatSper complex
sperm principal piece
nucleosome
synaptonemal complex
ciliary base
mitochondrial respiratory chain complex ...
postsynaptic membrane
dynein complex
voltage-gated potassium channel complex
integral component of membrane
membrane
0
1
2
3
4
5
6
7
8
9
10
log10 p-value

**Additional supplementary figure 2 -** Gene ontology (GO) enrichment analyses for absolutely blood specific TAU genes. The graphs correspond to the three domains of ontologies: biological process (BP), molecular function (MF) and cellular component (CC). The selected genes were analysed for enrichment in specific GO categories using the TopGO program against the background (all coding sequence genes). Y axis corresponds to enriched GO terms found in the respective domains (BP, MF and CC). X axis correspond to the log function of Fisher p-values obtained for each one of the enriched terms. The back line denotes a p-value = 0.05. P-values greater than 1,30 (log 0,05) indicate statistically significant enriched term.

Important genes part of the mitochondrial β-oxidation pathway involved in the catabolism of fatty acids were highly expressed in the trophosome, including the mitochondrial trifunctional protein and acyl-CoA dehydrogenases. The degradation of fatty acids by mitochondrial β-oxidation in trophosomal tissues has also been reported by Hinzke et al. (2019). Since acyl-CoA dehydrogenases (ACADs) constitute a multigene family, we performed similarity searches followed by phylogenetic inferences of well-annotated metazoan/lophotrochozoan ACAD proteins using homologs retrieved from the giant tubeworm and other annelid genomes (Swigoňová et al. 2009). Our analyses revealed that the *Riftia* genome contains 12 genes belonging to nine recognised ACAD subfamilies: ACAD-8 (isobutyryl-CoA dehydrogenase), -9, -10, SCAD (short-chain acyl-CoA dehydrogenase), MCAD (medium-chain acyl-CoA dehydrogenase), SBCAD (short/branched-chain acyl-CoA dehydrogenase), GCD (glutaryl-CoA dehydrogenase), IVD (isovaleryl-CoA dehydrogenase), and VLCAD (very long-chain acyl-CoA dehydrogenase) (Figure 4B, C and D). The subfamilies SCAD, MCAD, VLCAD, ACAD-9 and -10 are involved in the catabolism of fatty acids (Ye et al. 2004; Swigoňová et al. 2009), the remaining 4 families participate in the degradation of amino acids. We did not identify in *Riftia* any gene belonging to the LCAD subfamily (long-chain acyl-CoA dehydrogenase).

Genes involved with the TCA cycle (e.g., succinyl-CoA synthetase, dihydrolipoyl lysine-residue acetyltransferase) and oxidative phosphorylation process (e.g., NADH Dehydrogenase, ATP synthase) (Supplementary Figure 9) also present tissue-specificity in the trophosome. The intense activity of the mitochondrial respiratory chain in the trophosome is corroborated by the high expression of superoxide dismutase 2 genes (*sod2*; located in the mitochondrial matrix – Supplementary Figure 38; Supplementary Table 9) which neutralise highly reactive superoxide radicals protecting the tissue against oxidate stress (Miao and St. Clair 2009). Other antioxidant system, methionine sulfoxide reductase (Supplementary Table 9) (Sreekumar et al. 2011) also plays a role in the prevention of oxidative damage in the trophosome.

As *Riftia* is nutritionally dependent on its chemolithoautotrophic endosymbionts, which are deficient in polyunsaturated fatty acids (PUFAs), it has been questioned how the giant tubeworm obtains these biomolecules. PUFAs are precursors of a number of molecules and have important roles in inflammatory/immune responses, and membrane fluidity (Wallis et al. 2002). Surprisingly, contrary to most vertebrates which lack the enzymatic machinery required to synthesise PUFAs, *Riftia* contains the ω3- desaturase gene and is capable of desaturate and elongate fatty acids from the endosymbionts to gain access to PUFAs (Phleger et al. 2005; Liu et al. 2017). As fatty acids provide useful insights into the vent trophodynamics, nutritional strategies, and adaptation to abiotic factors (e.g., homeoviscous adaptation (Sinensky, 1974)) we screened the giant tubeworm, and closely related annelid taxa, for fatty acid desaturase genes. We identified eight fatty acid desaturases in *Riftia*, including the ω3-desaturase gene previously reported (Liu et al., 2017) (Additional supplementary figure 6A). Interestingly, based on our phylogenetic inferences, a clade has been recovered containing two *Riftia* paralog sequences classified as ω3-desaturase with a one-to-one orthology relationship with the cold seep tubeworm *Lamellibrachia.* The gene model RIFPA_jg1449 is 100% identical to the previously described ω3-desaturase gene and 84% identical to the *Lamellibrachia* ortholog. The second putative ω3-desaturase gene (RIFPA_jg32120) is 47% similar to the ω3-desaturase described by Liu et al. (2017). Whether this newly identified paralog present a ω3-desaturase activity remains to be shown. The remaining six genes and their isoforms are involved in sphingolipid, stearoyl-CoA, and acyl-CoA desaturase activities. Despite desaturases are ubiquitously expressed in tubeworm tissues, tissue-specificity was observed (Additional supplementary figure 6B). The presence of ω3-desaturase genes in annelids (including vestimentiferans), molluscs, and nematodes are in accordance with recent reports that invertebrates have the ability to produce PUFAs (Kabeya et al. 2018).

We identified in the *Riftia* genome the presence of 17 distinct cathepsin genes (Supplementary Figures 34-35) (cathepsin-C:1; cathepsin-Unc:1; cathepsin-B:2; cathepsin-L:4; cathepsin-Z:3; cathepsin-O:1; cathepsin-F:1; cathepsin-S:4), whereas the close relative *Lamellibrachia* contains 26 representatives (cathepsin-C:1; cathepsin-Unc:1; cathepsin-B:5; cathepsin-L:7; cathepsin-Z:3; cathepsin-O:1; cathepsin-F:2; cathepsin-S:6). Members of five out of the eight different cathepsins family are highly expressed in the trophosome (cathepsins-C/L/B/S/Z). The high expression of cathepsins in different trophosome samples belonging to *Riftia* and *Lamellibrachia* has been described also by Hinzke et al. (2019) and Li et al. (2019), respectively. Additionally, we identified moderate/high expression of several other lysosomal proteins, such as glycosidases (hexosaminidase alpha - *hexA*), sulphatases (heparan N-sulfatase - *sgsh*), proteases (legumain - *lgmn*), phosphatases (acid phosphatase 5, tartrate resistant – *acp5*), sphingomyelinases (sphingomyelin phosphodiesterase 1 – *smpd1*), and lipases (phospholipase A2 group XV – *lypla3*) in trophosome, which are probably also involved in the endosymbiont digestion.

Surprisingly, the male and female blood transcriptomes present distinct transcriptional landscapes, as showed by the few shared GO terms (Additional supplementary figure 4). The shared terms are related to cation/anion transport and cell adhesion, probably reflecting the transfer of metabolites and transport of coagulation factors through the circulatory system, respectively. The female blood TSGs are involved with haem-, oxygen-, and insulin-like binding activities, in agreement with the transport of gaseous substances and hormones in this tissue. Many male blood TSGs, however, point to cell divisions events. Gene expression analyses of key elements involved in the cell cycle showed that several cyclins and cyclin-dependent kinases are expressed in the blood samples, in agreement with the GO enrichment analysis (Supplementary Figure 54). The enrichment of cell cycle genes in the blood transcriptomes might be explained by the sampling of multipotential hematopoietic stem cells during the tissue collection. As the blood samples were obtained after the dissection of the trunk region of the tubeworms, we speculate that hematopoietic cells present either in the blood vessels and/or trophosomal peritoneal membrane were sampled (Southward et al. 2005; Hartenstein 2006; Nakahama et al. 2008).

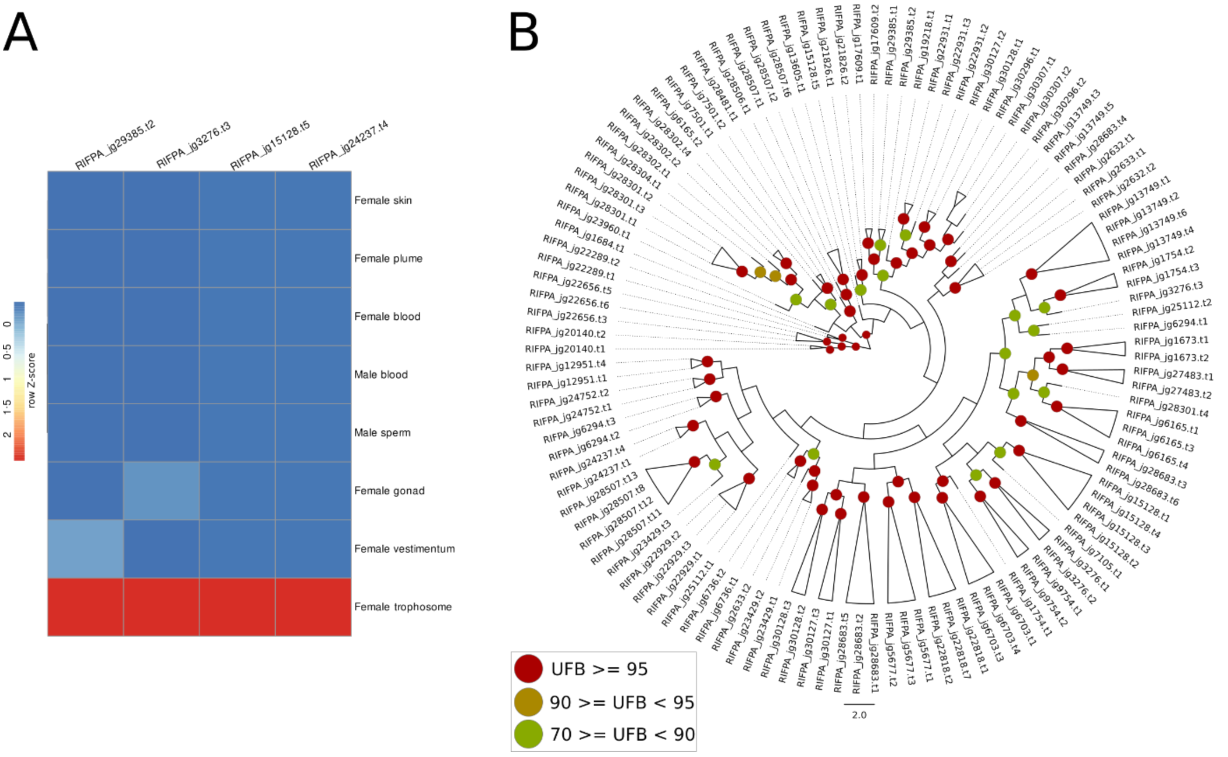

**Additional supplementary figure 3** – Gene expression and phylogeny of mitochondrial carriers present in the giant tubeworm genome. **A**, expression profile of four selected mitochondrial carrier genes in *Riftia pachyptila*. Colour coding reflects the expression patterns based on row Z-score calculations. B, phylogeny of 45 mitochondrial carriers found in the giant tubeworm genome. The branch support values are represented by the coloured circles in the tree nodes. Red circles represent ultrafast bootstrap values >= 95. Yellow circles represent ultrafast bootstrap values >= 90 and < 95. Green circles represent ultrafast bootstrap values < 90 and >= 70. Ultrafast bootstrap values smaller than 70 are not shown.

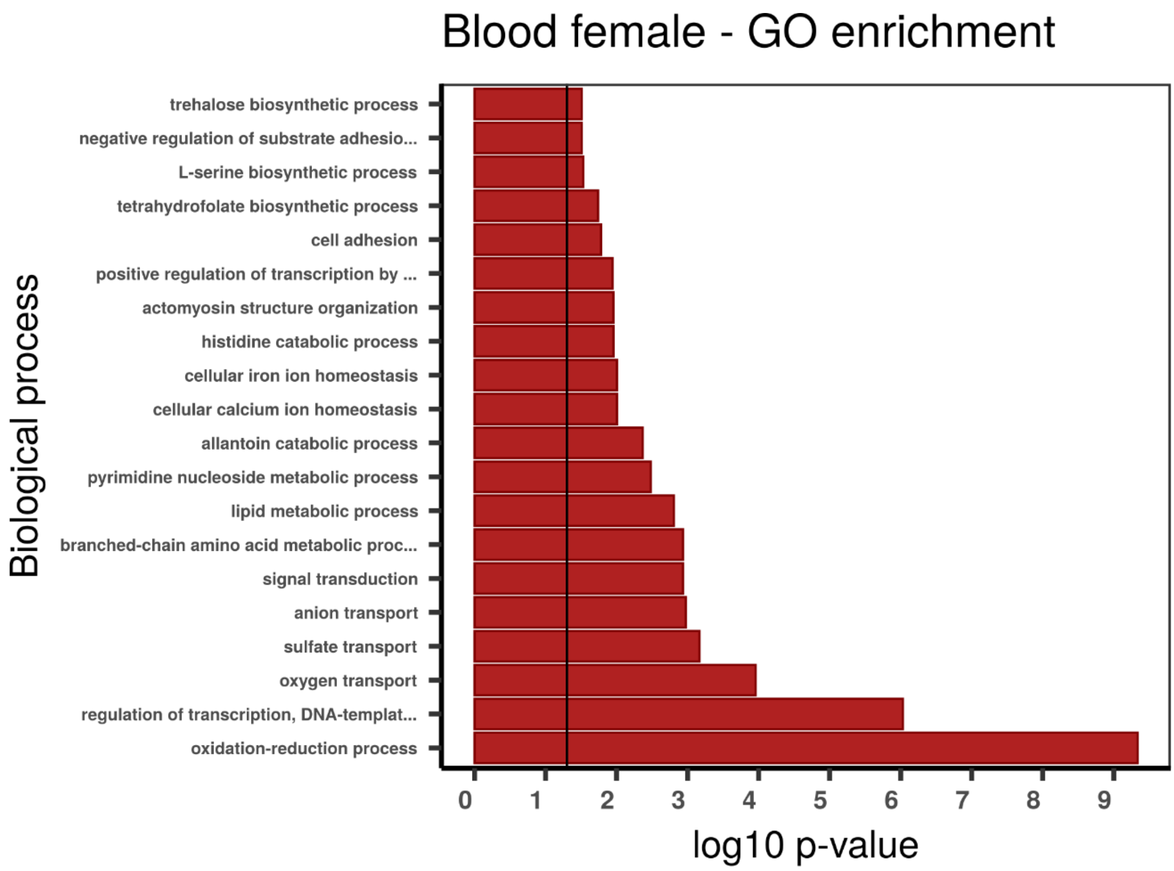
Blood female - GO enrichment
Biological process
trehalose biosynthetic process
negative regulation of substrate adhesio...
L-serine biosynthetic process
tetrahydrofolate biosynthetic process
cell adhesion
positive regulation of transcription by ...
actomyosin structure organization
histidine catabolic process
cellular iron ion homeostasis
cellular calcium ion homeostasis
allantoin catabolic process
pyrimidine nucleoside metabolic process
lipid metabolic process
branched-chain amino acid metabolic proc...
signal transduction
anion transport
sulfate transport
oxygen transport
regulation of transcription, DNA-templat...
oxidation-reduction process
0 1 2 3 4 5 6 7 8 9
log10 p-value

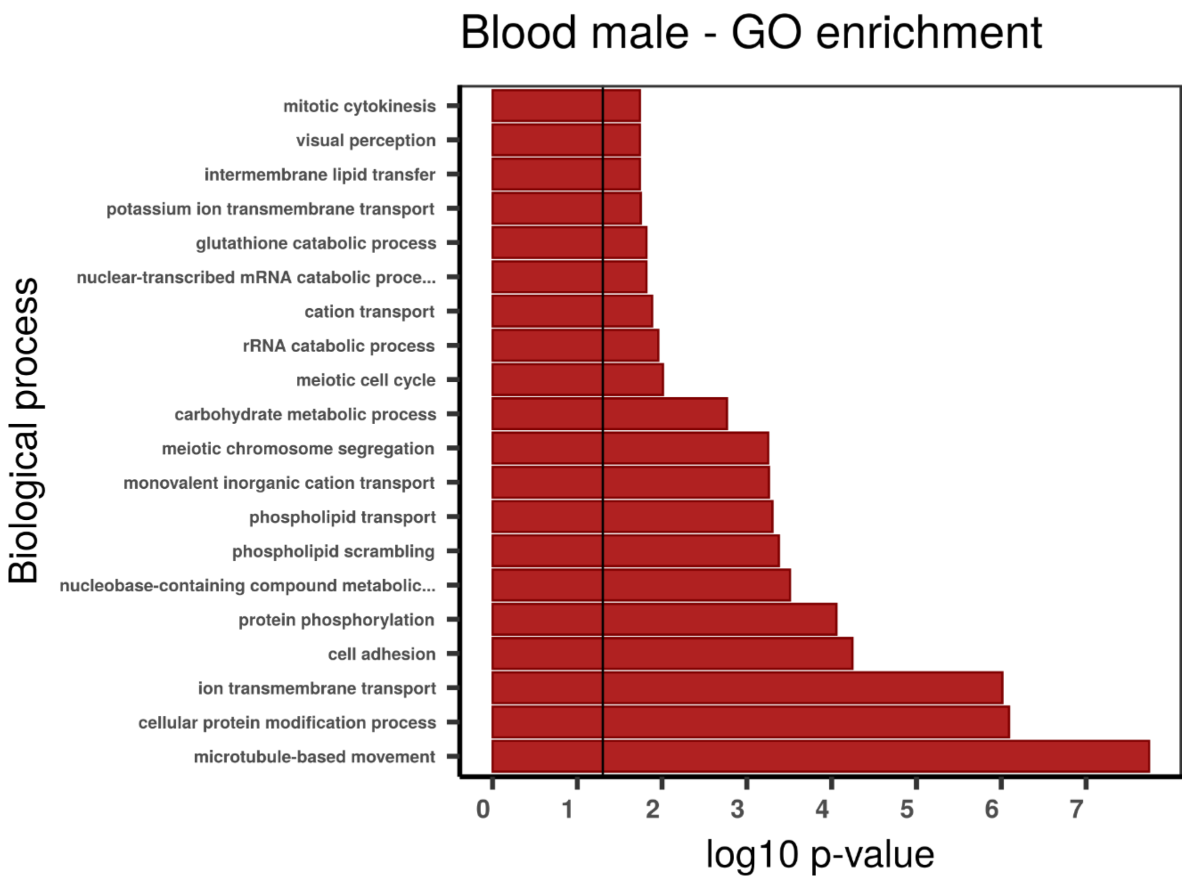
Blood male - GO enrichment
Biological process
mitotic cytokinesis
visual perception
intermembrane lipid transfer
potassium ion transmembrane transport
glutathione catabolic process
nuclear-transcribed mRNA catabolic proce...
cation transport
rRNA catabolic process
meiotic cell cycle
carbohydrate metabolic process
meiotic chromosome segregation
monovalent inorganic cation transport
phospholipid transport
phospholipid scrambling
nucleobase-containing compound metabolic...
protein phosphorylation
cell adhesion
ion transmembrane transport
cellular protein modification process
microtubule-based movement
0 1 2 3 4 5 6 7
log10 p-value

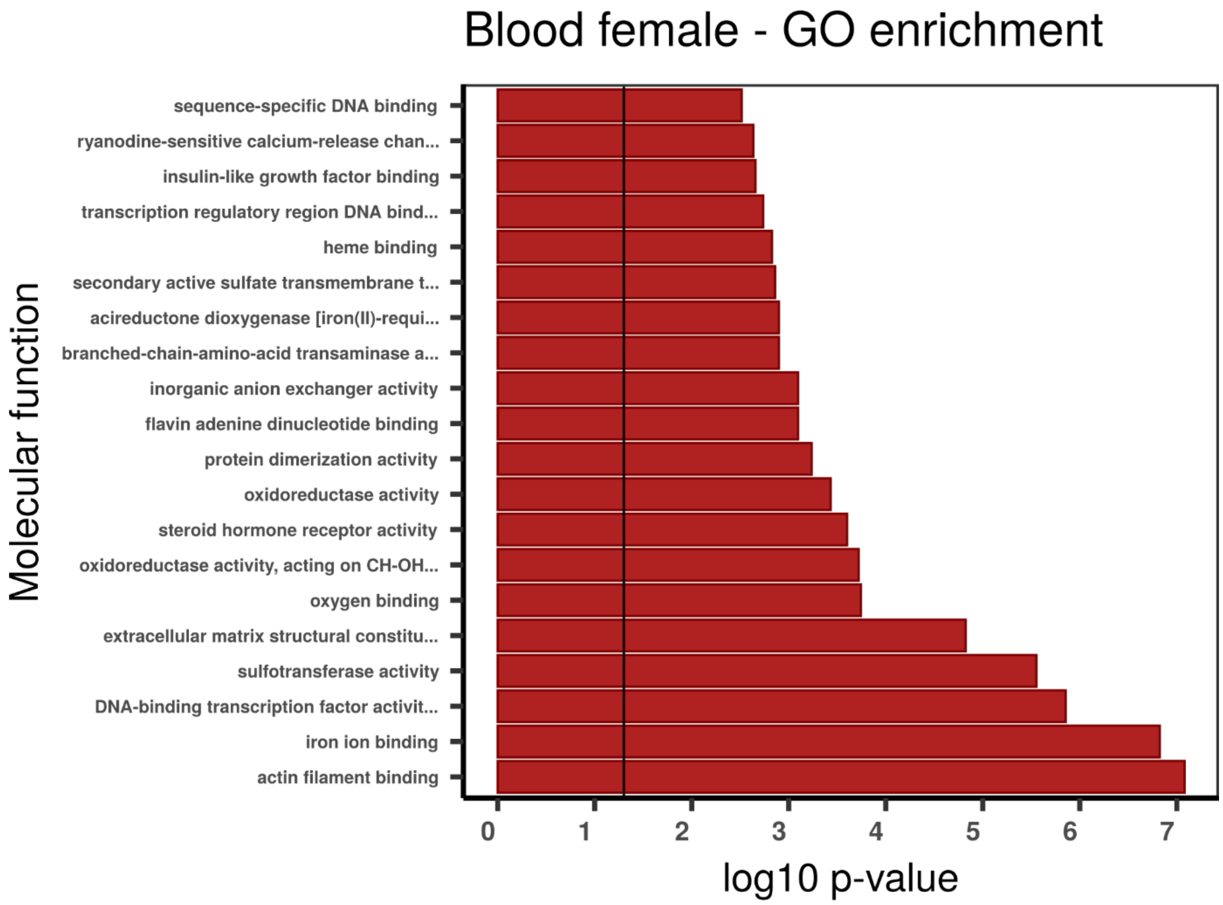

Blood female - GO enrichment
Molecular function
sequence-specific DNA binding
ryanodine-sensitive calcium-release chan...
insulin-like growth factor binding
transcription regulatory region DNA bind...
heme binding
secondary active sulfate transmembrane t...
acireductone dioxygenase [iron(II)-requi...
branched-chain-amino-acid transaminase a...
inorganic anion exchanger activity
flavin adenine dinucleotide binding
protein dimerization activity
oxidoreductase activity
steroid hormone receptor activity
oxidoreductase activity, acting on CH-OH...
oxygen binding
extracellular matrix structural constitu...
sulfotransferase activity
DNA-binding transcription factor activit...
iron ion binding
actin filament binding
0 1 2 3 4 5 6 7
log10 p-value

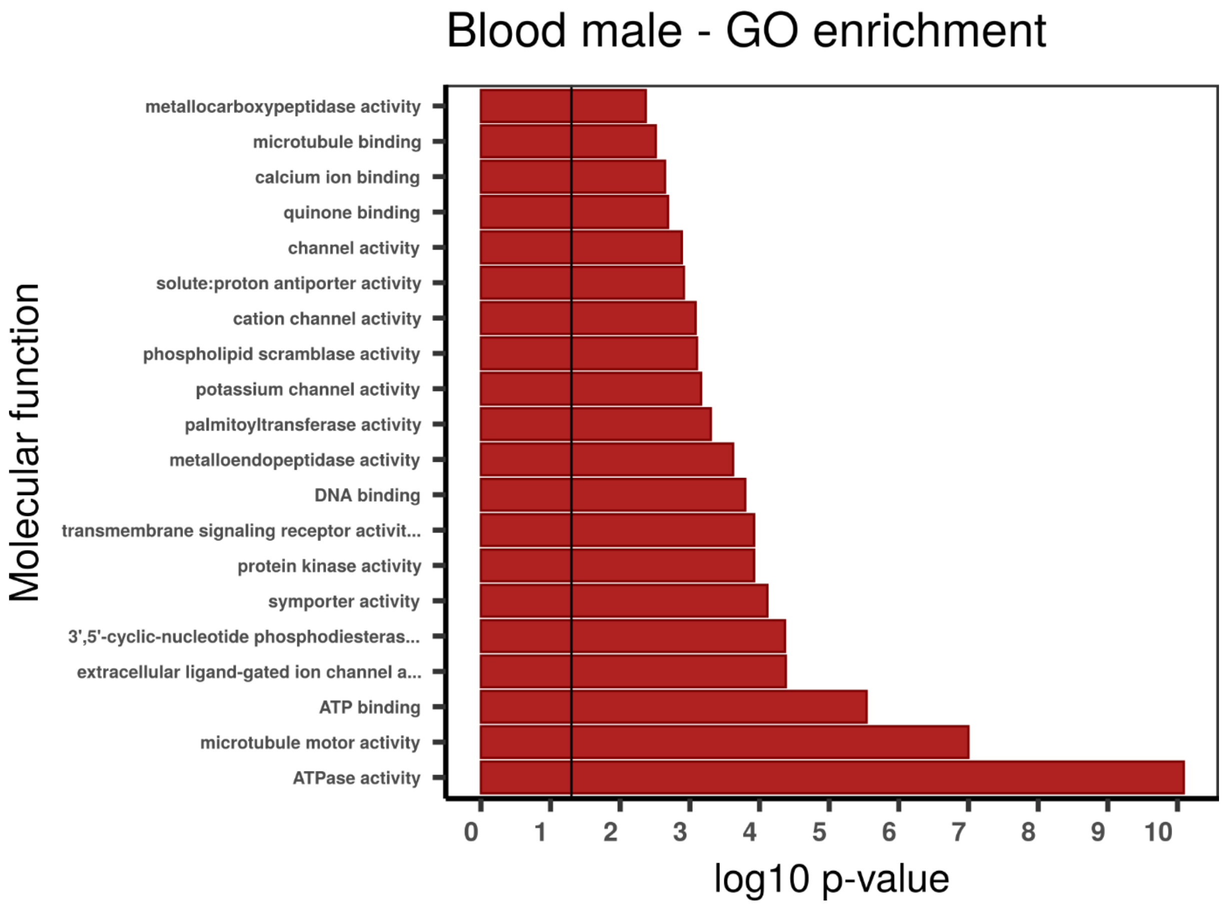

Blood male - GO enrichment
Molecular function
metallocarboxypeptidase activity
microtubule binding
calcium ion binding
quinone binding
channel activity
solute:proton antiporter activity
cation channel activity
phospholipid scramblase activity
potassium channel activity
palmitoyltransferase activity
metalloendopeptidase activity
DNA binding
transmembrane signaling receptor activit...
protein kinase activity
symporter activity
3',5'-cyclic-nucleotide phosphodiesteras...
extracellular ligand-gated ion channel a...
ATP binding
microtubule motor activity
ATPase activity
0 1 2 3 4 5 6 7 8 9 10
log10 p-value

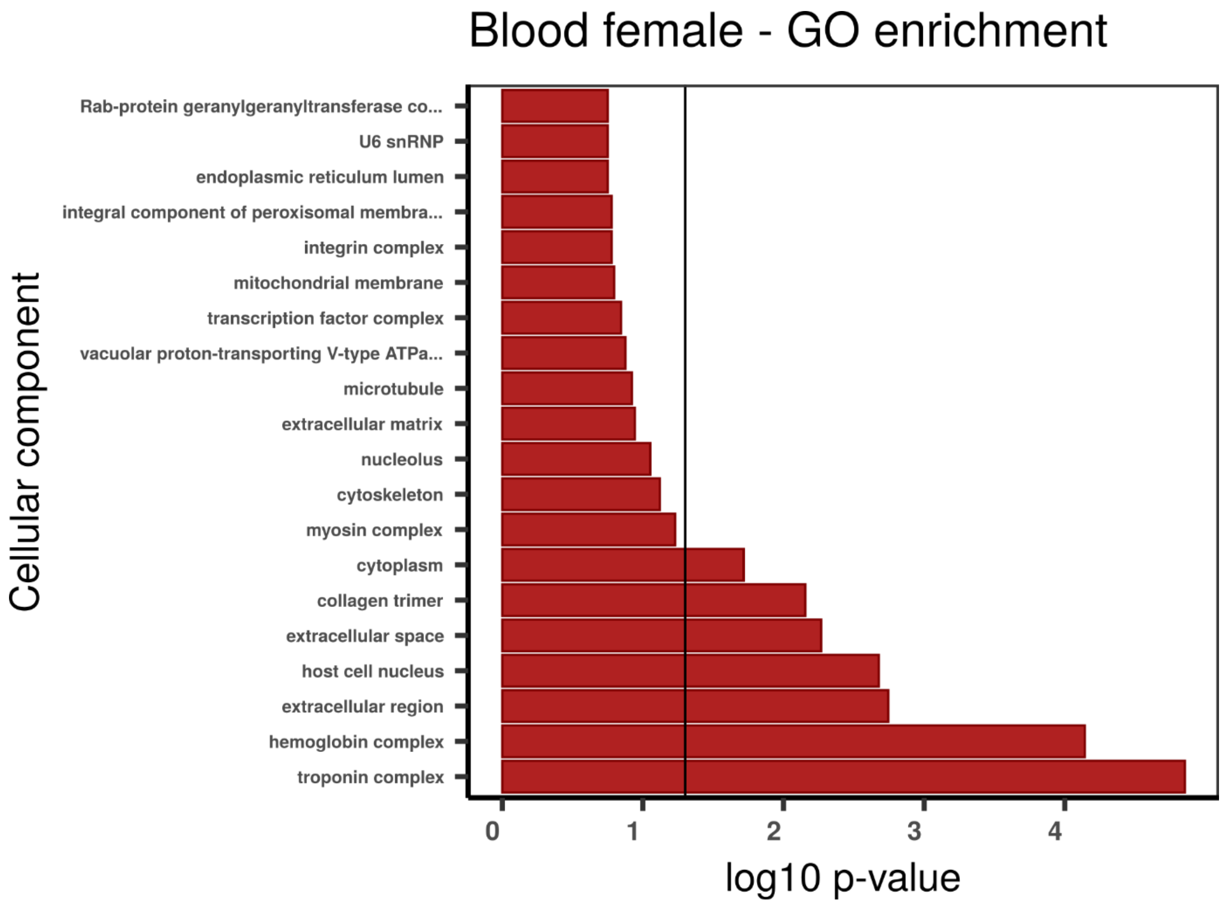

Blood female - GO enrichment
Cellular component
Rab-protein geranylgeranyltransferase co...
U6 snRNP
endoplasmic reticulum lumen
integral component of peroxisomal membra...
integrin complex
mitochondrial membrane
transcription factor complex
vacuolar proton-transporting V-type ATPa...
microtubule
extracellular matrix
nucleolus
cytoskeleton
myosin complex
cytoplasm
collagen trimer
extracellular space
host cell nucleus
extracellular region
hemoglobin complex
troponin complex
0
1
2
3
4
log10 p-value

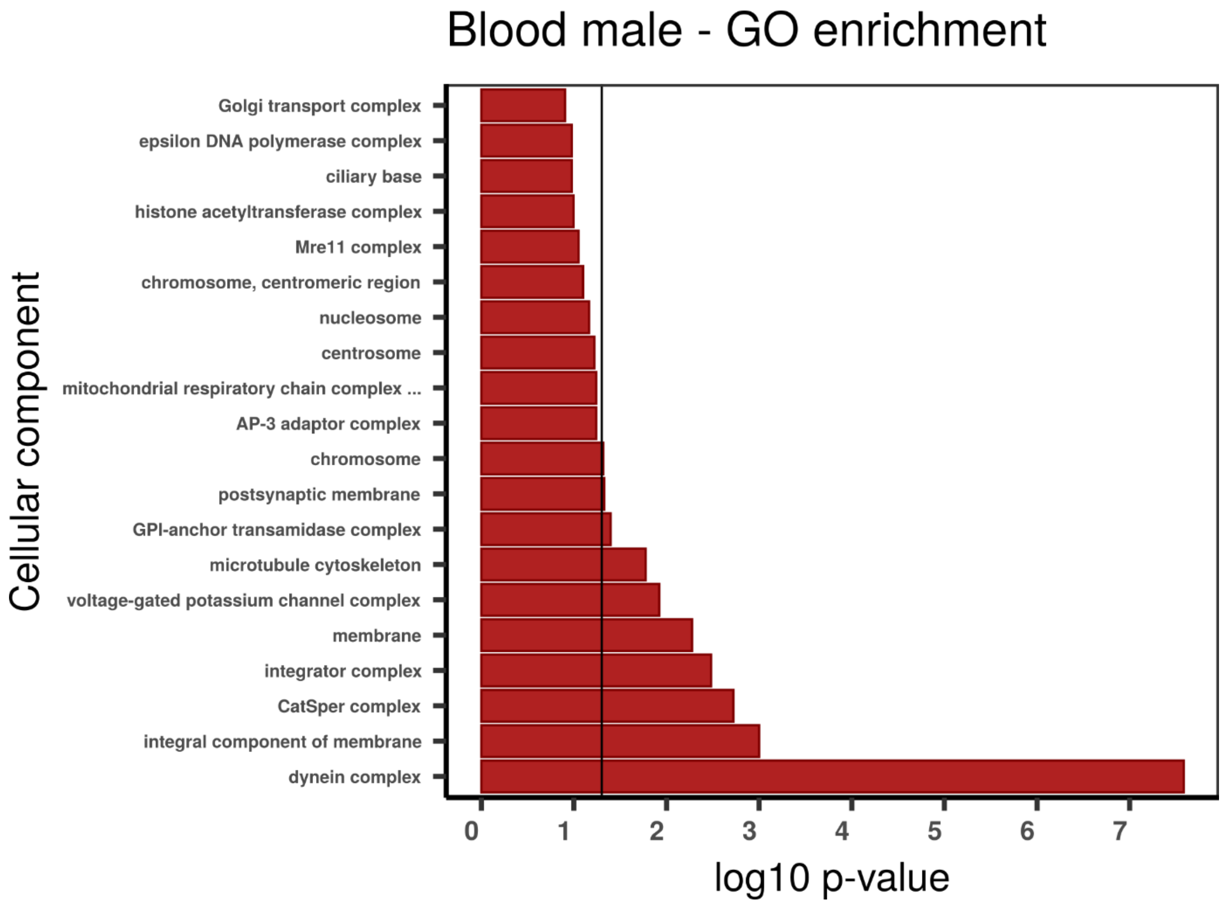

Blood male - GO enrichment
Cellular component
Golgi transport complex
epsilon DNA polymerase complex
ciliary base
histone acetyltransferase complex
Mre11 complex
chromosome, centromeric region
nucleosome
centrosome
mitochondrial respiratory chain complex ...
AP-3 adaptor complex
chromosome
postsynaptic membrane
GPI-anchor transamidase complex
microtubule cytoskeleton
voltage-gated potassium channel complex
membrane
integrator complex
CatSper complex
integral component of membrane
dynein complex
0
1
2
3
4
5
6
7
log10 p-value

**Additional supplementary figure 4 -** Gene ontology (GO) enrichment analyses for absolutely blood specific TAU genes. The graphs correspond to the three domains of ontologies: biological process (BP), molecular function (MF) and cellular component (CC). The selected genes were analysed for enrichment in specific GO categories using the TopGO program against the background (all coding sequence genes). Y axis corresponds to enriched GO terms found in the respective domains (BP, MF and CC). X axis correspond to the log function of Fisher p-values obtained for each one of the enriched terms. The back line denotes a p-value = 0.05. P-values greater than 1,30 (log 0,05) indicate statistically significant enriched term.

#### Supplementary note 6 | Nitrogen metabolism and excretion

To investigate the nitrogen metabolism in *Riftia*, we identified and quantified the gene expression of several enzymes related to the purineolytic (AMP deaminase), uricolytic (urease, uricase, xanthine dehydrogenase, OHCU decarboxylase, HIU hydrolase, allantoinase, and allantoicase), taurine/hypotaurine (taurocyamine kinase), purine (phosphoribosyl pyrophosphate amidotransferase, glycinamide ribonucleotide transformylase, phosphoribosylformylglycinamidine synthase, phosphoribosylaminoimidazole carboxylase and phosphoribosylamino-imidazolesuccinocarboxamide synthase, adenylosuccinate lyase, 5-Aminoimidazole-4-carboxamide ribonucleotide formyltransferase, guanine monophosphate synthase, adenylosuccinate synthase, hypoxanthine phosphoribosyltransferase, and adenine phosphoribosyltransferase), pyrimidine (CAD, dihydrooratate dehydrogenase, uridine-5 monophosphate synthase), polyamine pathways (spermine, spermidine), as well as the urea (arginase, argininosuccinate lyase, argininosuccinate synthase, and ornithine carbamoyltransferase) and ammonia cycles (glutamate dehydrogenase, glutamine synthetase) (Supplementary Figures 41-45).

The genes AMP deaminase, urease, OHCU decarboxylase, HIU hydrolase, allantoinase, allantoicase, mitochondrial taurocyamine kinase, and the four investigated enzymes in the urea cycle are present as single copy in the giant tubeworm genome, whereas the genes xanthine dehydrogenase and the cytoplasmatic taurocyamine kinase contain four and five distinct copies each, respectively. Three xanthine dehydrogenases/oxidases paralogs (RIFPA_jg12112, RIFPA_jg12113 and RIFPA_jg12115) and four cytoplasmatic taurocyamine kinase genes (RIFPA_jg16631, RIFPA_jg16632, RIFPA_jg16633, RIFPA_jg16634) are located in chromosomal clusters in *Riftia* (Figure 5A). All orthologs of the aforementioned genes were also identified in the cold seep tubeworm. The genes allantoinase, urease and the cytoplasmatic taurocyamine kinase contain one additional copy in *Lamellibrachia* compared to *Riftia,* totalling two (LAMLU_FUN_033197-T1, LAMLU_FUN_020286-T1), two (LAMLU_FUN_032122-T1, LAMLU_FUN_032120-T1) and six paralogs (Supplementary Figure 43), respectively.

Overall, the genes involved in the urea cycle and uricolytic pathway are ubiquitously expressed in all tubeworm tissues. Particularly in the trophosome, the urea cycle enzymes argininosuccinate synthase/lyase, arginase, ornithine carbomoyltransferase, and key components of the uricolytic pathway (urease, 5-hydroxyisourate hydrolase, uricase, allantoicase, allantoinase, OHCU decarboxylase, xanthine dehydrogenase) are highly/moderately expressed (Figure 41). These results agree with biochemical analyses which show an elevated concentration of uric acid, urea, and ammonia in the trophosome in comparison to other symbiont-free tissues (e.g., plume, blood, body wall) (Cian et al. 2000), and with a more recent metaproteomic and comparative transcriptomic study (Hinzke et al. 2019). These results point to different metabolic processes concerning the nitrogen metabolism in the trophosome and their importance in the host biology and/or endosymbiotic association.

Phosphagen kinases (PKs) constitute an evolutionary conserved family of phosphoryl transfer enzymes with important roles in energy homeostasis (Conejo et al., 2008; Ellington, 2001). PKs catalyse the reversible transfer of gamma phosphory group of ATP to guanidino compounds, such as taurocyamine and arginine (Uda et al. 2005; Uda et al. 2006; Suzuki et al. 2009). Hitherto in *Riftia*, one cytoplasmatic and one mitochondrial phosphotaurocyamine kinase have been described as the major phosphagen kinase system of the energy metabolism (Uda et al., 2005). We identified four additional copies of taurocyamine kinase in the *Riftia* genome. They are highly expressed in all tubeworm tissues, with the exception of the trophosome and skin (body wall) (Supplementary Figure 44). The cold seep tubeworm genome harbours six copies (five cytosolic and one mitochondrial) of taurocyamine kinases, whereas *Capitella* contains one and two copies of the mitochondrial and cytosolic types, respectively. We did not identify any taurocyamine kinase gene in the leech *Helobdella*. The finding of additional taurocyamine kinase genes in *Riftia* could clarify the regulation of glycogenolysis and intracellular energy transport, which are important factors that account for the tubeworm large size and rapid growth.

In a series of biochemical and thorough studies Minic et al. demonstrated that *Riftia* lacked the initial enzymes involved in the pyrimidine biosynthetic pathway relying on the endosymbiont for the *de novo* synthesis of pyrimidines and polyamine production (Minic et al. 2001; Minic et al. 2002; Minic and Hervé 2003; Minic and Hervé 2004). These results were challenged recently by Hinzke et al. (2019) which identified the trifunctional protein CAD (carbamoyl-phosphate synthetase 2, aspartate transcarbamylase, and dihydroorotase) and key genes in the polyamine synthesis, i.e., spermidine and spermine synthases, in the host metatranscriptome. We, through similarity searches and phylogenetic inferences, identified the CAD and the polyamine-related genes in the giant tubeworm genome, confirming that *Riftia* is indeed less dependent on the symbiont regarding the nitrogen metabolism. Additionally, in agreement with Hinzke et al. (2019), that did not detect the expression of CAD on a protein level, we observed that the CAD is lowly expressed in the eight analysed tissues (Supplementary Table 8 – Supplementary Figures 42 and 44), which probably is the explanation for Minic et al. (2002) results. In fact, all the genes involved in the pyrimidine biosynthesis, with the exception of uridine 5 monophosphate synthase, present a low expression level in the giant tubeworm tissues (Supplementary Table 8). Spermine and spermidine have been implicated in host-symbiont interactions in *Riftia*, due the restricted distribution of the enzymes catalysing these polyamines solely in trophosome metaproteomes (Hinzke et al. 2019). We, however, observed a high expression of these enzymes in all *Riftia* tissues (Supplementary Table 8; Supplementary Figures 42 and 44), corroborating their important roles in the host overall homeostasis (e.g., protein and nucleic acid synthesis, protection from oxidative damage, cell proliferation, apoptosis, and differentiation (Pegg, 2016)). Both vestimentiferans, *Riftia* and *Lamellibrachia*, contain one single copy of the CAD and spermidine synthase gene, whereas a duplication of the gene spermine synthase is observed in tubeworms (Supplementary Table 8; Supplementary Figures 42 and 44). These results show that the *de novo* pyrimidine biosynthesis and polyamine production is also maintained in the cold seep tubeworm. The key enzymes involved in the *de novo* purine synthesis are present in the giant tubeworm genome (Supplementary Table 8; Supplementary Figure 42).

*Riftia* requires a high demand of nitrogen because of the high biomass and growth rates (Lutz et al. 1994). These requirements are usually achieved by the level of environmental nitrate and ammonia in the environment (Johnson et al. 1988; Lilley et al. 1993), or by the active digestion of the symbionts (Girguis et al. 2000). We identified and quantified the gene expression of important genes containing the glutamine synthetase and glutamate dehydrogenase protein domains in the giant tubeworm genome and closely related annelid specifies and lophotrochozoans (Supplementary Table 8; Supplementary Figure 45; Figure 5). These enzymes are required to incorporate ammonium into organic matter. Glutamine synthetase and glutamate dehydrogenase show an elevated expression in eight tubeworm tissues herein investigated, especially in the plume and gonad tissues. The relative contribution of ammonium derived from the diffusion into the host by the endosymbionts (i.e., reduction of nitrate into nitrate and then ammonia by Endoriftia (Girguis et al. 2000)), or from the degradation of symbionts by lysosomal activity is still elusive. We identified in annelids the presence of genes belonging to two distinct clades of glutamine synthetase group I: the lengsins and a group so far containing only prokaryotic sequences. (Figure 5C; Supplementary Figure 43). Lengsins constitute a group of glutamine synthetase genes which were co-opted to non-enzymatic roles in the vertebrate eye lens (Wyatt et al. 2006). The giant tubeworm and *Lamellibrachia* genomes contain seven and 13 copies of lengsins, respectively. Surprisingly, only annelid sequences and poriferan sequences clustered together with hitherto prokaryotic glutamine synthetase I sequences in the phylogenetic tree (Figure 5). The glutamine synthetase group II, traditionally found in eukaryotes, includes two paralog copies of *Riftia* and *Lamellibrachia* genes. Vestimentiferans possess the highest number of glutamine synthethase-containing genes in all 18 investigated lophotrochozoans, highlighting the importance of the ammonia assimilation in the tubeworms. Lengsins are highly expressed in the *Riftia* trophosome.

In polychaetes, the site of the ammonia excretion and/or the mechanisms in which this nitrogenous waste is secreted are still poorly understood, with few studies publicly available (Smith et al. 1987; Smart and Von Dassow 2009; Thiel et al. 2016; Rimskaya-Korsakova et al. 2018; Weihrauch and Allen 2018; Rimskaya-Korsakova et al. 2020). A recent study investigated the expression important structural genes (*nephrin*, *kirrel* and *zo1*) responsible for the formation of filtering sites in the excretory organs of lophotrochozoans, including the polychaete *Owenia fusiformis,* ecdysozoans and deuterostomes taxa (Gąsiorowski et al. 2021). We screened the genome of the giant tubeworm for the presence of these three structural genes and quantified their gene expression in the eight tissue-specific transcriptomes (Additional supplementary figure 5).

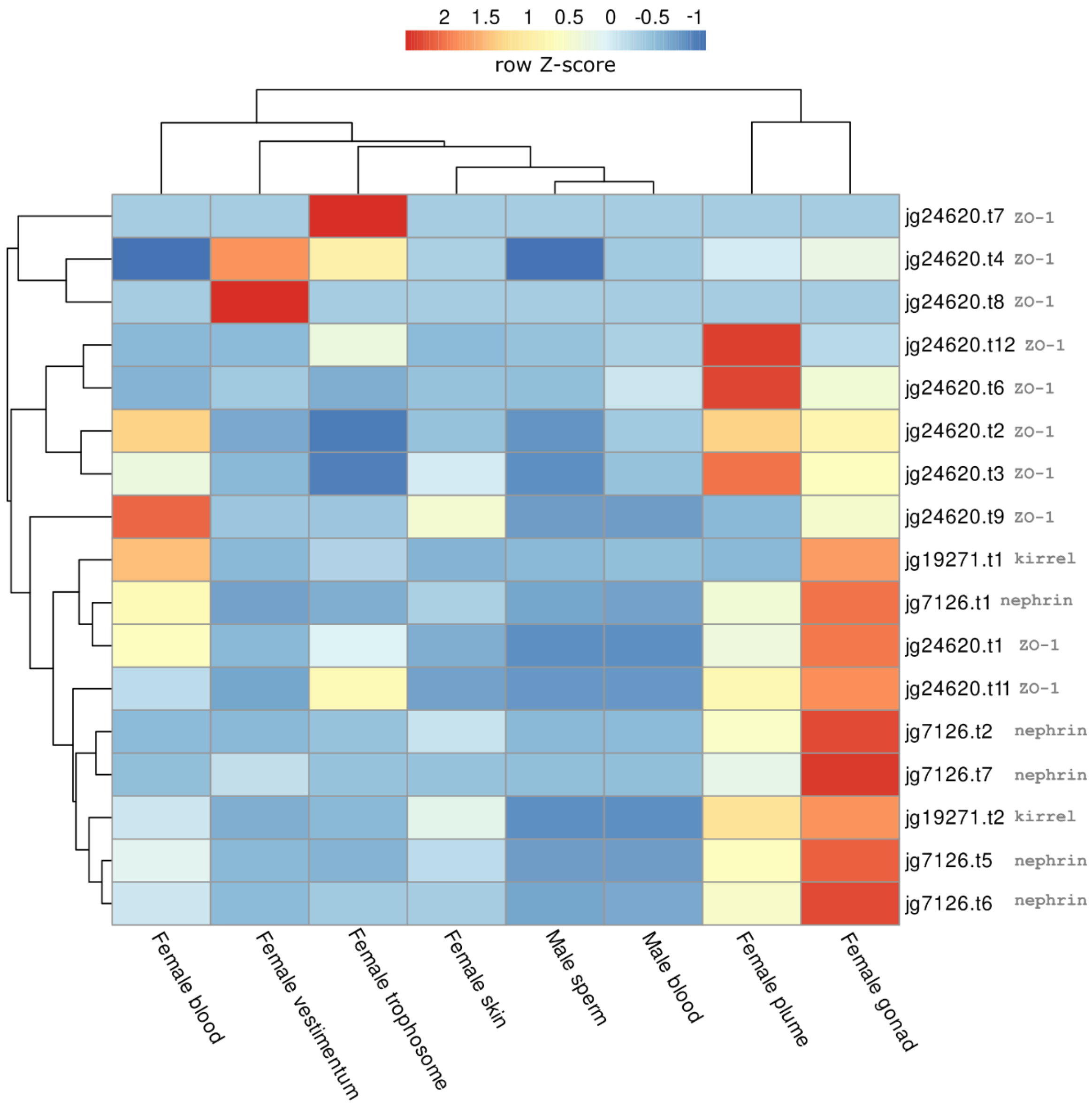

**Additional supplementary figure 5.** Expression profile of three important structural genes (*nephrin, kirrel* and *ZO-1*) related to excretory organs in *Riftia pachyptila*. Colour coding reflects the expression patterns based on row Z-score calculations. Desaturases are ubiquitously expressed in all tubeworm tissues.

We identified in *Riftia* and *Capitella* a unique copy of each one of the structural genes *nephrin* (RIFPA_jg7126, CAPTE_P184830), *kirrel* (RIFPA_jg19271, CAPTE_P184834), and *ZO-1* (RIFPA_jg2460, CAPTE_P221876). Duplications of *kirrel* (LAMLU_FUN_007962-T1, LAMLU_FUN_024093-T1) and *ZO-1* (LAMLU_FUN_029123-T1, LAMLU_FUN_033606-T1) were identified in the *Lamellibrachia,* and a possible secondary loss of *kirrel* and *nephrin* is present in the *Helobdella* genome. *Nephrin* and *ZO-1* are present as single copies in the *Lamellibrachia* and *Helobdella* genomes, respectively (LAMLU_FUN_008794-T1, HELRO_176655).

All three genes are highly expressed mainly in the female gonad, plume, and blood tissues of *Riftia*, in agreement previous studies that show that these proteins interact together in the filtering cells of ultrafiltration excretory organs (Gerke et al. 2003; Huber et al. 2003; Liu et al. 2003; Gąsiorowski et al. 2021). The gene expression results, especially in the plume region, are surprising, since the putative site of ultrafiltration in vestimentiferans is located in the large protonephridial excretory tree part of the vestimental region (Schulze 2001; Bright and Lallier 2010; Rimskaya-Korsakova et al. 2018).

The storage of nitrogenous waste in the trophosome may be explained, at least in part, by the unusual excretory organ found in vestimentiferans. Recently, a microanatomical study identified protonephridia in *Ridgeia pisceseae* (Schulze 2001; Rimskaya-Korsakova et al. 2018). In fact, it is highly unusual that these giant tubeworms do not develop metanephridia, known to be the dominant form of excretory organ in large polychaetes with a well-developed blood vascular system (Ruppert and Smith 1988; Bartolomaeus and Quast 2005). While in metanephridium the podocytes facilitate filtration from the blood to the coelomic fluid, in protonephridia only coelomic fluid is filtered (Ruppert and Smith 1988).

**Supplementary note 7 | Cell proliferation, innate immune system, apoptosis, and autophagy**

Immunohistochemical and ultrastructural cell cycle investigations have shown that in vestimentiferans, including *Riftia*, cell proliferation activities are higher than in any other characterised invertebrate (Pflugfelder et al. 2009). We investigated key regulatory enzymes present in the cell cycle (G1, S, G2 and M) with special focus on cyclin-dependent kinases (CDKs), *smad4* and *smad2* (Supplementary Figure 46). CDKs are serine/threonine kinases which depend on their regulatory subunits, cyclins, to perform catalytic activities (Lim and Kaldis 2013; Malumbres 2014). *Smad4* and *smad2* serve as the central mediator of TGF- β signalling pathway and are involved in a wide range of cellular processes, including uncontrolled proliferation leading to cancer initiation (Nakao et al. 1997; Zhao et al. 2018). Additionally, *smad4* has been shown to undergo an independent expansion and present signs of positive selection in the cold seep tubeworm *Lamellibrachia luymesi* (Li et al. 2019), which may play important roles controlling the high proliferation rates in vestimentiferans.

We identified in the *Riftia* genome through similarity searches and phylogenetic inferences all the expected CDKs involved in the G1, S, G2 and M phases of the cell cycle and the *smad4* gene. All genes are present as single copies in the giant tubeworm genome. KEGG pathway analysis reconfirmed our findings. We identified three copies of *smad4* in *Lamellibrachia*, two in *Helobdella robusta* and one in *Capitella*, contradicting Li et al. (2019) results regarding the number of homologues in the cold-seep tubeworm and the leech. Gene expression analysis showed that the cell cycle genes are ubiquitously expressed across the studied tubeworm tissues, with non-symbiotic tissues (i.e., female gonad, blood, plume and vestimentum) showing the highest expression levels (Supplementary Figure 46). Cyclin A is highly expressed in the female gonad and blood, indicative of DNA synthesis, and G2 phase, whereas cyclin B2 is highly expressed, in vestimentum and plume, indicative of mitosis. In the trophosome, the high expression of cyclins A and B2 marking the S, G2 and M phase of mitosis have been identified, despite the overall low expression of CDKs in this tissue. These results indicate active DNA synthesis and proliferation events in this tissue, which were also detected with immunocytochemistry in the unipotent stem cells in the central and the semi-differentiated cells in the median zone of the trophosome lobule (Pflugfelder et al. 2009). The transcription factors *smad4* and *smad2* seem to be down regulated in the trophosome, an indicative of high cell proliferation activity (Samanta and Datta 2012).

Innate immunity activation is facilitated by pattern recognition receptors such as the Toll-like receptors (TLRs) and peptidoglycan receptor proteins (PRPs) to recognize microbe associated molecular patterns (Nyholm and Graf 2012). The TLRs constitute an evolutionary conserved transmembrane family of proteins involved in innate immune responses in metazoans (Janssens and Beyaert 2003; Kawai and Akira 2006; Song et al. 2012; Kawasaki and Kawai 2014; Liu et al. 2020) (Supplementary

Figures 47-57). Briefly, when TLRs recognise pathogen-associated molecular patterns (PAMPs), the myeloid differentiation primary response protein 88 (*MyD88* gene) recruits various adaptor molecules (e.g., IRAK, TAB1/TAK1 complexes) that trigger the activation of the *AP1* and *NF-κB* transcription factors, that in turn, control the outcome of the innate immune responses. In lophotrochozoans, lineage-specific expansions (phoronids, bivalves) and gene loss (rotifers, planarians, and blood flukes) of TLR genes have been reported (Sun et al. 2017; Luo et al. 2018). We identified 17 distinct TLR genes in the giant tubeworm genome, with orthologous gene counterparts in the close relative *Lamellibrachia luymesi*, which contains 28 TLRs genes (Supplementary Figures 47-49). We could not clearly assign the tubeworm TLRs into the different vertebrate classes, probably due the lineage-specific duplications of these proteins in vertebrates and vestimentiferans. Additionally, *Riftia* contains all expected core proteins involved in the Toll-like receptor/MyD88 pathway, and gene expression analyses showed that TLRs and other components involved in the innate immune system are mainly expressed in the female gonad, plume, vestimentum and skin tissues. This suggests that TLR does not play a decisive role in the trophosome.

The cellular response upon innate immune system recognition and tissue homeostasis is driven either by apoptosis or autophagy. Alternatively, apoptosis (Supplementary Figures 50-57) and autophagy (Supplementary Figure 58) play important roles in host-symbiont interactions acting in the regulation of the endosymbiont populations in the deep-sea bivalve *Bathymodiolus platifrons* and the cereal weevil *Sitophilus*, respectively (Vigneron et al. 2014; Sun et al. 2017). Firstly, we screened the giant tubeworm genome for known protein domains involved with apoptotic process: caspases (Supplementary Figures 50-51), B-cell lymphoma 2 (Bcl-2; Supplementary Figure 52), the tumour necrosis factor ligands (TNF; Supplementary Figure 53) and receptors (TNFR; Supplementary Figure 54), and inhibitor of apoptosis proteins (IAP; Supplementary Figures 55-57). Caspases are intracellular cysteine proteases responsible for the controlled degradation of cells during apoptosis (Julien and Wells 2017). They are activated by extracellular ligands, such as TNF death receptor superfamily, and are involved in the extrinsic apoptosis pathway (Park et al. 2007). We identified in the *Riftia* genome 11 caspases and one para-caspase, making the giant tubeworm complement the smallest amongst annelids and molluscs. The cold-seep tubeworm contains 15 caspases and two para-caspases. Caspase expression in *Riftia* is stronger in the plume, and female gonad than in the skin and vestimentum, and in the trophosome. The Bcl-2 proteins, which are involved in the intrinsic apoptosis pathway (Czabotar et al. 2014) are highly/moderately expressed in the plume, female gonad, but not in the trophosome. *Riftia* contains a Bcl-2 complement of five genes.A total of four and six TNFRs/TNF ligands were found in the tubeworm genome, with overlapping gene expression in the plume and female gonadal tissues. In the trophosome, however, we found highly expressed a TNF ligand, but not the required corresponding TNFRs. TNFs (as well as extrinsic factors) activate caspases, intracellular cysteine proteases responsible for the controlled degradation of cells during apoptosis (Julien and Wells 2017). The IAPs, which negatively regulates caspases and cell death (Silke and Meier 2013), has been shown to be expanded in the deep-sea/seep mussel bivalves *Bathymodiolus* (130 genes) and *Modioulus* (95 genes), respectively (Sun et al. 2017). *Riftia* and *Lamellibrachia,* however, contains a relatively simple anti-apoptotic system with only 19 and 36 genes, respectively. IAP gene expression was upregulated in all giant tubeworm tissues, but mainly highly expressed in the plume and female gonad tissues. Only a few IPA genes were highly expressed in the vestimentum, skin and trophosome tissues.

Overall, tissues exposed to the environment exhibited high expression of apoptosis-related genes, while the skin and the vestimentum (protected by the tube) and the internally located trophosome appears much less involved in apoptosis. Despite a great variability in the number upregulated genes involved in the apoptotic events, all studied tubeworm tissues showed high expression of IAPs. This may be due to the fact that all samples consisted of multiple different cell types some of which might be upregulated in apoptosis, others inhibited (e.g., skin is composed of the epidermis, gland, nervous, and muscle cells). Even in the trophosome, despite the presence of blood vessels and aposymbiotic sheet cells, only the peripheral region of the tissue's lobules was found to die during terminal differentiation through apoptotic processes, while the more central region was not involved in massive apoptosis (Bright and Sorgo 2003; Pflugfelder et al. 2009). These differences in the morphological and ultrastructural organisation of the trophosome is certainly followed by variation in gene expression levels of apoptotic genes.

While phagocytes are described to clear the tissue from apoptotic cell remains (deCathelineau and Henson 2003), the terminal step of autophagy is lysosomal degradation of either sequestered cytosolic material present in autolysosomes (membrane-bound vesicles fused with lysosomes), or the direct uptake of components by the lysosomes, as found in macro- and microautophagic processes (Glick et al. 2010; Hansen et al. 2018), respectively. An additional and more complex type of autophagy, named chaperone-mediated autophagy, involves the degradation of targeted proteins through their translocation across the lysosomal membrane with the aid of chaperone proteins (Glick et al. 2010; Hansen et al. 2018). To date, the complete autophagic pathway has not been identified in any annelid genome and remains unknown. Furthermore, this study and others show that *Riftia* (Hinzke et al. 2019), and a closely related vestimentiferan (Li et al. 2019), present a high expression of lysosomal hydrolase genes in the trophosome, implicating that the tubeworms digest their endosymbionts for nutrition. To elucidate the autophagy pathway in *Riftia*, we identified through similarity searches and phylogenetic analyses the core genes (Atg proteins – autophagy related proteins (Klionsky 2012)) involved in macroautophagy process (hereafter termed autophagy), since this is the predominant and most studied form of autophagy (Mizushima and Komatsu 2011) (Supplementary Figure 58).

This type of programmed cell death starts with the key upstream regulators present in the mTORC1 complex, which phosphorylates the Atg1/ULK1 complex initiating the autophagy (Wong et al. 2013). The membrane nucleation, phagophore formation and expansion are mediated by Atg proteins present in the PI3K complex I, ATG12 and LC3/GABARAP conjugation systems (Glick et al., 2010). *Atg2*, and *atg9* also participate in the autophagosome formation by transferring phospholipids and expanding the newly synthesized phagophore (Velikkakath et al. 2012; Zhou et al. 2017; Gómez-Sánchez et al. 2018; Kotani et al. 2018; Osawa et al. 2019). Lastly, the autophagosome maturation is achieved with the fusion of the autophagosome with lysosomes to form the autolysosome. *Riftia* contains all the core elements commonly found in yeast, human and bivalve autophagy pathways, demonstrating the high conservation of this mechanism in distantly related eukaryotic taxa (Ohsumi 2014; Picot et al. 2020).

The gene family Atg2 contains two paralogs in the giant tubeworm genome (*atg2a* and *atg2b*) and the same scenario is found mammals (Velikkakath et al. 2012; Tamura et al. 2017). Two paralogous sequences of Atg4 (*atg4a, atg4cd*) are present in *Riftia.* The cold-seep tubeworm contains three paralogs, whereas the worm *Caenorhabditis elegans* and humans present two and four copies, respectively. These results indicate different patterns of gene duplication in different closely related species, as well as in protostome, and deuterostome lineages. Atg8 gene family contains four members in *Riftia*, *Lamellibrachia*, *Capitella* and *Helobdella* (*gabarap*, *gabarapl2*, *map1lc3ab*, *map1lc3c*), indicating a conservation of gene numbers of this autophagic gene family in annelids. The Atg8 family has undergone different gene gain and loss events throughout the metazoan tree, with noticeable expansions of two (GABARAP and LC3/MAP1LC3) out of the three subfamilies in vertebrates (Shpilka et al. 2011). The Gate16/GABARAPL2 subfamily is represented by a single copy gene in most of the animals, except for sponges (*Amphimedon queenslandica*) and echinoderms (*Stronglycentrotus purpuratus*) which contains two paralogous genes (Shpilka et al. 2011). All the remaining autophagy-related genes are present as single copy in the vestimentiferans *Riftia* and *Lamellibrachia*. Gene expression analyses showed that Atg genes present different levels of expression in the eight tubeworm tissues, with the female plume, gonad and blood samples harbouring most of the highly expressed genes. The trophosome, however, only shows upregulation of a few genes only, indicating that autophagy is not a widespread mechanism of control death in this tissue.

Lysozymes have an important dual role in invertebrate-endosymbiont interactions, not only controlling the intracellular symbiont population, but also accessing nutritional resources during the process (Nakabachi et al. 2005; Nishikori et al. 2009; Xue et al. 2010; Futahashi et al. 2013; Kim et al. 2014; Detree et al. 2016). As previously mentioned, lysosomal degradation is involved in distinct controlled cell death mechanisms involving recognition and elimination of self and nonself (i.e., apoptosis and autophagy). However, other different pathways, such as the endocytic, employ the lysosomal system for the digestion of macromolecules (Bröker et al. 2004; Guicciardi et al. 2004; Mariño et al. 2014; Hu et al. 2015; de Castro et al. 2016; Man and Kanneganti 2016; Wong et al. 2017). To identify important early, late and recycling endosomal genes in the giant tubeworm genome, we performed multiple sequence similarity searches using metazoan Rab GTPases (Stenmark 2009), and other key genes (transferrin, EEA1, M6PR) (Brown et al. 1986; Mu et al. 1995; Mayle et al. 2012) as queries, followed by phylogenetic inferences. We found that key genes involved in the different stages of endosome maturation are present as single copy in the giant and cold-seep tubeworm genomes (Supplementary Figure 37). The endosomal genes are expressed in all *Riftia* tissues, with the plume and the trophosome harbouring distinct highly expressed early and late endosomal genes. This suggest that endosomal degradation is prevalent in the trophosome but also in aposymbiotic hosts tissue, most strongly expressed in the plume and the female gonad.

Taking into consideration all the aforementioned results, it is hard to definite state, solely based on gene expression analyses and orthology inferences, the exact scenario behind the endosymbiont digestion and trophosome tissue homeostasis in *Riftia*. To make matters worse, autophagic and apoptotic events may occur simultaneously within the same cell and the trophosome contains distinct proliferative and degenerative regions (Bright and Sorgo 2003; Pflugfelder et al. 2009; Mariño et al. 2014). Nonetheless, as previously mentioned, we did not identify an abundant expression of autophagic, apoptotic, and immune-related genes in the trophosome, but only a high expression of lysosomal hydrolases, and moderate/highly expression of endosomal genes in this tissue. These results point that endosymbiont digestion in *Riftia* is probably independent of innate immune system recognition and programmed cell death events, in a process similar to endosome maturation, as suggested by Hinzke et al. (2019). We hypothesized a mechanism that involves the fusion of the double membrane symbiosomes with lysosomes (coined here as symbio-lysosome structure) followed by the direct digestion of the endosymbionts.

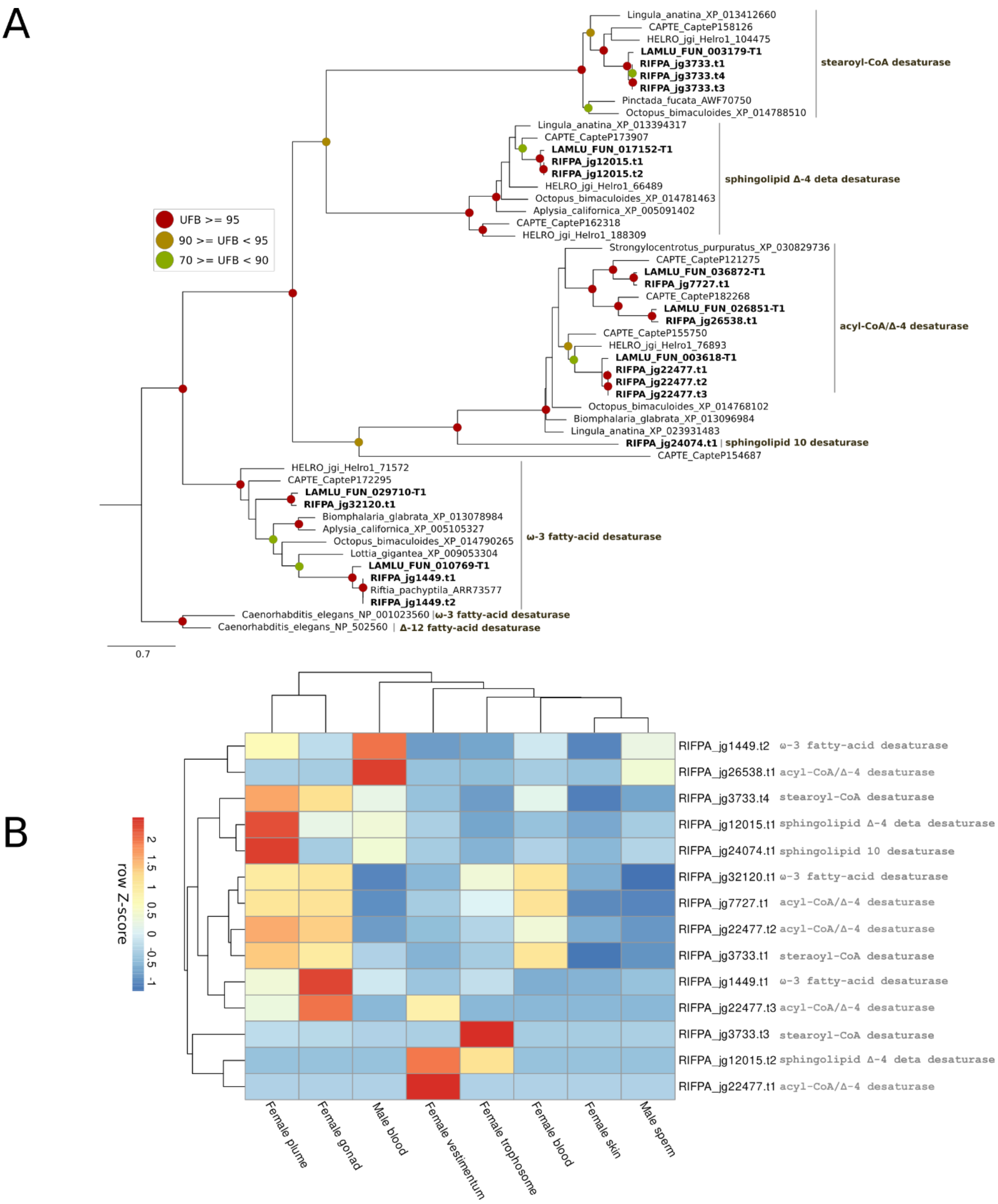

**Additional supplementary figure 6** – Phylogeny and gene expression of fatty acid desaturases in the giant tubeworm genome. **A**, Phylogeny of eight desaturases identified in the the giant tubeworm genome. The branch support values are represented by the coloured circles in the tree nodes. Red circles represent ultrafast bootstrap values >= 95. Yellow circles represent ultrafast bootstrap values >= 90 and < 95. Green circles represent ultrafast bootstrap values < 90 and >= 70. Ultrafast bootstrap values smaller than 70 are not shown. Accession numbers for NCBI database are displayed after the species names. *Capitella, Helobdella* and *Lamellibrachia* gene identification are derived from the publicly available annotated genomes. **B**, expression profile of fatty acid desaturases in the genome of *Riftia pachyptila*. Colour coding reflects the expression patterns based on row Z-score calculations. Desaturases are ubiquitously expressed in all tubeworm tissues.

The high expression of programmed cell-death, immune system and cell cycle genes in the plume and female gonad of the giant tubeworm raises interesting questions about the proposed model of symbiosis between the deep-sea *Bathymodiolus platifrons* and its methane oxidising endosymbiont. Sun et al. (2017) showed high expression levels of gene families related to immune recognition, endocytosis, and caspase-mediated apoptosis in the gills of *B. platifrons*, suggesting evolutionary adaptations of the deep-sea mussel towards its endosymbionts. Our analysis, however, did not identify an abundant expression of proliferation, immune system, and controlled cell death (i.e., apoptosis and autophagy) gene markers in the trophosome. Two alternative hypotheses can be drawn from these results. First, the similar transcriptional landscapes of the gills and the plume, in *B. platifrons* and *Riftia*, respectively, indicate similar defence mechanisms against pathogens and homeostasis of the respiratory organs, rather than the control of the endosymbionts in the bivalve. Second, these two lophotrochozoans display distinct mechanisms to control, digest and maintain the symbiont populations within their respective endosymbiont harbouring tissues. We argue, that as the trophosome is internally located in the trunk region of the giant tubeworm not contacting the deep-sea vent fluids, it represents a more “sterile” and controlled environment to understand the molecular dynamics behind the host-endosymbiont mutualism. Cell atlases of vestimentiferan trophosome tissues obtained from single cell transcriptomic analyses (Aldridge and Teichmann 2020) are required to completely unravel the biology of this tissue.

**SUPPLEMENTARY MATERIAL AND METHODS**

#### 1 GENOME SEQUENCING AND ASSEMBLY

##### 1.1 Sample collection, genomic DNA extraction and Sequencing Strategy

*Riftia* genomic DNA was obtained from a piece of vestimentum tissue belonging to single worm collected at the hydrothermal vent site Tica, East Pacific Rise (Alvin dive 4839, 9º 50.398 N, 104º 17.506 W, 2514 m depth, 2016) (Supplementary Figures 1, 2). The preparation of the vestimentum sample was performed by grinding 100mg of frozen tissue with a pre-chilled mortar. After a fine powder was obtained, the tissue was added into a 50 mL conical tube with 9.5 mL of Buffer G2, 19 µL of RNAse A (200 µg/mL final concentration), and 0.5 mL of QIAGEN protease (all reagents except for RNAse A are part of the QIAGEN Blood and Cell Culture Midi Kit (catalog #13343)). The tissue and the reagents were thoroughly mixed in a vortexer, and subsequently incubated at 50 °C for two hours. Lysate was immediately loaded onto an equilibrated QIAGEN Genomic-tip upon completion of incubation step. The Genomic-tip was double washed with 7.5 mL of Buffer QC and then genomic DNA was eluted with 5 mL of Buffer QF (pre-warmed to 50 °C). High-molecular weight DNA was precipitated using 3.5 mL of isopropanol and centrifuged at 5,000x g for 15 minutes at 4 °C. The supernatant was carefully removed without disturbing the pellet. The sample was air-dried for ten min and resuspended in in 200 µL of Tris-HCl, pH 8.5. DNA was dissolved overnight a shaker. Finally, the DNA was quantified and stored at -80 °C. PacBio libraries were generated with Sequel technology for a *de novo* large insert library (30kb) using the SMRTbell Express Template Kit 2.0 and purified *Riftia* DNA by the University of Minnesota Genomics Center (St. Paul, MN, USA).

##### 1.2 Genome pre-processing

The five PacBio libraries were individually converted from the native BAM to fastq file format using bam2fastq tool v1.3.0 present in the "PacBio Secondary Analysis Tools on Bioconda" (https://github.com/PacificBiosciences/pbbioconda). The resulting libraries were then mapped with minimap v2.17-r941 (Li 2018) against a custom database of contaminants composed of the closed genome of the *Cand.* Endoriftia persephone and the *Riftia pachyptila* reference mitochondrial genome (Accession number: NC_026860). All PacBio read sequences that failed to align against any of the references were stored in a file and used in the assembly procedure.

##### 1.3 Genome assemblies

The long filtered PacBio reads were assembled with canu v1.8 (Koren et al. 2017) using the parameters optimised for Sequel chemistry and heterozygosity (see https://github.com/marbl/canu/issues/1470 for details). An alternative assembly, as a purpose of benchmarking and quality control, was produced using the tool flye v2.5 under default parameters (Kolmogorov et al. 2019). The quality assessment of the draft genomes was performed with quast v5.0.2 (Gurevich et al. 2013).

#### 1.4 Genome post-processing

##### 1.4.1 Polishing

To obtain a high-quality consensus draft genome and call variant types (e.g., insertion and deletions), the native PacBio BAM files were aligned against the genome assemblies with pbmm2 tool v1.0.0 (https://github.com/PacificBiosciences/pbmm2), a minimap2 SMRT wrapper for PacBio data present in the "PacBio Secondary Analysis Tools on Bioconda" (https://github.com/PacificBiosciences/pbbioconda). The mapped results together with the assembly file, were submitted to arrow v2.3.3 (https://github.com/pacificbiosciences/genomicconsensus/) for genome polishing and variant calling. The quality assessment of the polished high-quality draft genomes was performed with quast v5.0.2 (Gurevich et al. 2013).

##### 1.4.2 Purging haplotigs and contig overlaps

The removal of haplotigs and contig overlaps was performed manually with purge_dups using the recommended pipeline available at: https://github.com/dfguan/purge_dups. The read depth cutoffs were manually adjusted based on histogram plots (e.g., lower, and upper bounds for read depth) in order to avoid overpurging. The read depth was calculate aligning the PacBio original data against the polished genomes with minimap v2.17-r941 (Li 2018). The quality assessment of the purged polished high-quality draft genome was performed with quast v.5.0.2 (Gurevich et al. 2013).

##### 1.4.3 Contamination screening with blobtools

To further assess microbial contamination, a blobplot was generated following the recommended "Workflow A" available at https://blobtools.readme.io/docs/the-blobtools-workflows#section-workflow-a. The blobtools v1.1.1 (Laetsch and Blaxter 2017) was executed with: (1) a coverage file obtained by mapping the five filtered long-read PacBio libraries (i.e., libraries without mitochondrial and *Cand.* Endoriftia persephone bacterial reads) against the purged polished genome; (2) the *Riftia pachyptila* purged polished high-quality draft genome; and (3) a hit file containing NCBI TaxIDs resulted from the alignment of the draft tubeworm genome against the nt database (57,030,965 sequences) using the blastn v2.8.1+ (Camacho et al. 2009). Two scaffolds assigned to bacterial TaxIDs were carefully inspected using three additional methods:

1. Similarity search approach: the predicted protein coding sequences from the two suspicious scaffolds were aligned against the NCBI nr database (229,636,095 sequences) with diamond blastp v0.9.25.126 (Buchfink et al. 2015). The TaxIDs from the best diamond blastp hits, were checked for consistent and narrow patterns of bacterial phyletic distribution with MEGAN v6.18.5 (Huson et al. 2016).
2. Gene architecture and gene density approach: the presence of intronless genes (i.e., single exon proteins) and the gene density in the suspicious

scaffolds were analysed, as prokaryotic genomes present a much higher gene density than eukaryotes and do not contain any introns.

3. Genomic completeness approach: the suspicious scaffolds were checked for bacterial marker genes with checkM v1.1.2 (Parks et al. 2015).

##### 1.5 Mitochondrial genome assembly and annotation

The PacBio reads mapped against the *Riftia* reference mitochondrial genome were assembled using flye v2.5 (Kolmogorov et al. 2019). The polishing of the extrachromossomal genome was performed as described in section 1.4.1. The fully reconstructed mitochondrial genome was annotated with MITOS2 (Bernt et al. 2013: 2) (http://mitos2.bioinf.uni-leipzig.de/index.py) and GeSeq (Tillich et al. 2017) (https://chlorobox.mpimp-golm.mpg.de/geseq.html) online servers. Manual curation of the mitogenome was performed using the previous *Riftia* mitogenomes available at NCBI as references. The circular plots generated with ConcatMap (https://github.com/darylgohl/ConcatMap) and circos (Krzywinski et al. 2009).

#### 2 TRANSCRIPTOME SEQUENCING AND ASSEMBLY

##### 2.1 Sample collection, total RNA extraction and Sequencing Strategy

One female *Riftia* (24 cm length) was collected from the hydrothermal vent Rebecca's Roost (27 0.64548684 N, 111 24.41801816 W, 2012 m depth), Guaymas Basin, Feburary 27 2019 during SuBastian dive 231. One male (75 cm length) was collected from a vent close to Big Pagoda (27 0.82596514 N, 111 24.66462214 W, 2028 m depth), Guaymas Basin, March 1 2019 during SuBasian dive 233. Tissue samples from one female (plume, vestimentum, body wall (skin), trophosome, blood and gonad) and one male individual (sperm and blood) were obtained, immerged in RNAprotect Bacterial Reagent (Quiagen) prior freezing at -80 degrees. The RNA from the RNAprotect stabilized tissue samples was purified using RNeasy Plus Mini Kit (Qiagen Cat. No. 74134). Sample disruption and homogenization was performed as follows: not more than 30mg of RNAlater stabilized tissue was transferred to a sterile RNase free 2ml reaction tube, which was pre-cooled by liquid nitrogen. The tissue samples were disrupted by grinding them to a fine powder under liquid nitrogen; 600ul of RTL buffer was added and the lysate was transferred to a QIAshredder spin column (Qiagen Cat. No. 79656) for homogenization. The following gDNA elimination and RNA purification steps were performed following the suppliers instructions, and finally, the purified RNA (30ul) was immediately stored -80° C.The RNA samples were sent to the Vienna Biocenter Core Facility (VBCF: https://www.viennabiocenter.org/facilities/next-generation-sequencing/) and stranded paired-end RNA-seq libraries (2x150 pb) of the eight adult tissues were constructed using the NEB/poly-A kit followed by an ultra-deep sequencing with the Illumina NovaSeq SP platform.

##### 2.2 Transcriptome pre-processing

The quality of the RNA-seq libraries was assessed with fastqc v0.11.8 (https://www.bioinformatics.babraham.ac.uk/projects/fastqc/). Based on the reported statistics obtained from bbduk v38.42 (https://sourceforge.net/projects/bbmap/) we selected the most appropriated parameters to filter the raw paired-end RNA-seq libraries. Illumina adapter sequences as well as poor quality bases/reads were trimmed from the tubeworm transcriptomic libraries and the final high-quality RNA-seq libraries used in the subsequent analyses.

##### 2.3 Transcriptome assembly

The eight high-quality paired-end RNA-seq libraries were individually assembled using two distinct approaches: (1) *de novo* strategy with transabyss v2.0.1 (Robertson et al. 2010), and (2) reference-based strategy using the draft *Riftia pachyptila* genome, STAR aligner v2.7.1a and Stringtie v2.0.6 (Dobin et al. 2013; Kovaka et al. 2019).

###### 2.3.1 *De novo* transcriptome assembly

The filtered and trimmed RNA-seq libraries were assembled using transabyss v.2.0.1 (Robertson et al. 2010) with default parameters, paired-end and strand-specific modes activated and the minimum transcript length defined as 200 base pairs. The quality assessment of the *de novo* transcriptome was performed with quast v5.0.2 (Gurevich et al. 2013).

###### 2.3.2 Reference-based assembly

The filtered and trimmed RNA-seq libraries were mapped against the hard-masked draft genome of *Riftia pachyptila* with STAR v2.7.1a (Dobin et al. 2013). The mapping results in BAM format were sorted and index with samtools v1.9-138 (Li et al. 2009) and the transcriptomes reconstructed with Stringtie v2.0.6 (Kovaka et al. 2019). To obtain a global non-redundant transcriptome, Stringtie v2.0.6 was executed again with the “transcript merge mode” invoked, using as input the eight reference-based reconstructed tissue specific transcriptomes. The quality assessment of the global non-redundant transcriptome was performed with quast v5.0.2 (Gurevich et al. 2013).

##### 2.4 *De novo* transcriptome post-processing

###### 2.4.1 Removal of endosymbiont contamination

To remove any possible endosymbiont contamination, the *de novo* transcriptome assemblies were submitted to blastn v2.8.1+ (Camacho et al. 2009) similarity searches against a custom database composed of the combined *Riftia pachyptila* hard-masked draft genome and the closed genome of *Cand.* Endoriftia Persephone (De Oliveira, under revision). Transcripts that aligned exclusively to the reference bacterial genome or mapped to both bacterial and annelid genomes (tagged as "suspicious transcripts") were removed from the transcriptomes.

##### 2.4.2 Generating a *de novo* global non-redundant transcriptome

In order to obtain a global *de novo* non-redundant transcriptome, a "SuperTranscript" approach was employed (Davidson et al. 2017a; Davidson et al. 2017b) using the tools Bowtie v2.3.4.3, Corset v1.09 (https://github.com/Oshlack/Corset) and Lace v1.13 (https://github.com/Oshlack/Lace) (Langmead and Salzberg 2012; Davidson and Oshlack 2014). Briefly, the trimmed high-quality RNA-seq paired-end reads were multi-mapped against the *de novo* transcriptomes using Bowtie v2.3.4.3 and the mapping results used to cluster the transcripts into genes with Corset. Finally, Lace v1.13 was used to remove the redundancy and build the global *de novo* "SuperTranscriptome". The quality assessment of the *de-novo* global non-redundant transcriptome was performed with quast v5.0.2 (Gurevich et al. 2013).

##### 2.5 Prediction of the coding sequence regions

The global *de novo* and reference-based non-redundant transcriptomes, as well as the individual tissue-specific *de novo* transcriptomes were searched for candidate coding sequence regions (CDS) with TransDecoder.LongOrfs and TransDecoder.Predict v.5.5.0 (https://github.com/TransDecoder/TransDecoder). TransDecoder runs were carried out following three criteria: (1) the coding sequence regions must contain a minimum of 100 amino acids; (2) only the top strand must be analysed, due the strand-specificity of the RNA-seq libraries; and (3) CDS with homology against PFAM-A v32.0 (17,929 pHMMs - http://hmmer.org/) and/or Swiss-Prot v2019_11 (561,568 sequences - "UniProt," 2019) databases were retained in the final output. The homology searches were performed with hmmscan v3.1b2 and blastp v2.8.1+ (Camacho et al. 2009; Mistry et al. 2013), and all the file conversions required from TransDecoder to work properly were achieved following the TransDecoder online tutorial (https://github.com/TransDecoder/TransDecoder/wiki). The quality of the final predicted proteomes obtained from the global non-redundant *de novo* and reference-based transcriptomes was measured with BUSCO4 (Simão et al. 2015).

#### 3 GENOME ANNOTATION

##### 3.1 Identification of interspersed repetitive regions and low complexity DNA

*De novo* transposable element families were modeled and identified in the tubeworm genome with RepeatModeler v2.0 (A.F.A Smit, R. Hubley & P. Green, *RepeatMasker Open-4.0*), which employs three complementary *de novo* repeat finding software RECON, RepeatScout and LtrHarvest/Ltr_retriever (Bao 2002; Price et al. 2005). The masking of the interspersed repetitive regions and low complexity DNA in the *Riftia pachyptila* genome was performed with RepeatMasker v4.0.9 using the custom database of repetitive sequences generated by RepeatModeler v2.0. The Repeat landscape was generated using the accessory scripts calcDivergenceFromAlign.pl and createRepeatLandscape.pl available at the RepeatMasker v4.0.9 package.

##### 3.2 *Ab initio* gene predictions

The *ab initio* gene predictions were performed with AUGUSTUS v3.3.3 (Stanke and Morgenstern 2005; Hoff and Stanke 2018). The *Riftia* gene models required for training AUGUSTUS were obtained using an unsupervised procedure implemented by GeneMark-ES version 4.48_3.60_lic (Lomsadze et al. 2014) followed by homology searches with diamond v0.9.25.126 (Buchfink et al. 2015) against nr database. The hints files, i.e., extrinsic structural/positional information obtained from the RNA-seq data about gene, intron, and exon boundaries, as well as the correct coding reading frame, were obtained using the combined *de novo* and reference based TransDecoder results (see section 2.5) and a comprehensive bioinformatic protocol (Hoff and Stanke 2018). Finally, the *Riftia* UTR gene models were retrieved from the Transdecoder gff3 genome annotation files and included only genes with complete coding sequence regions and both 3'- and 5' UTR features annotated. These genes were further filtered through sequence similarity searches against the nr database using diamond blastp v0.9.25.126. A total of three gene predictions were performed: (1) gene prediction without any hints files; (2) gene prediction with hints file without UTR model, and; (3) gene prediction with hints file and UTR model activated. The three gene predictions were merged with the tool joingenes, and the coding sequence regions (in both amino acid and nucleotide format) were extract with getAnnoFastaFromJoingenes.py (both softwares available as auxiliary tools in the AUGUSTUS V3.3.3 package).

##### 3.3 Filtering AUGUSTUS gene model predictions

The merged AUGUSTUS prediction was further submitted to homology searches against the PFAM-A v32.0 and nr databases using hmmsearch v3.1b2 and diamond blastp v0.9.25.126 tools. Additionally, the eight trimmed and filtered tubeworm tissue specific transcriptomes were mapped against the merged AUGUSTUS gene models with kallisto v0.46.1 (Bray et al. 2016), and orthology inferences were performed with orthoFinder v2.3.8 (Emms and Kelly 2019) using the *Riftia pachyptila* AUGUSTUS predicted proteins and other four publicly available annelid genomes (Simakov et al. 2013; Paul et al. 2018; Li et al. 2019). Only gene models with sequence similarity against the protein databases, and/or orthology and evidence of gene expression were retained. The quality of the combined AUGUSTUS gene prediction was measured with BUSCO4 (Simão et al. 2015).

##### 3.4 Protein annotation

The predicted protein sequences were scanned for protein domains and functional sites using Interproscan 5.39-77.0 (Jones et al. 2014) and 16 publicly available databases (TIGRFAM v15.0, SFLD v4, Phobius v1.01, SUPERFAMILY v1.75,

Gene3D v4.2.0, Hamap v2019_01, ProSiteProfiles v2019_01, Coils v2.2.1, SMART v7.1, CDD v3.17, PRINTS v42.0, ProSitePatterns v2019_01, Pfam v32.0, MobiDBLite v2.0, PIRSF v3.02, TMHMM v2.0c). The prediction of tRNA and signal peptide sequences were performed with tRNAscan-SE 2.0.5 and signalP v5.0b, respectively (Nawrocki and Eddy 2013; Lowe and Chan 2016; Almagro Armenteros et al. 2019).

##### 3.5 Protein family analyses (PFAM)

The pfam_scan.pl script (ftp://ftp.ebi.ac.uk/pub/databases/Pfam//Tools/PfamScan.tar.gz) was used to align selected lophotrochozoan protein sequences against a database of HMM profiles (Pfam-A v.33.0) using the standard hmmscan program (v3.1b2) (Mistry et al. 2013). This software, in additional to the standard hmmscan searches, clusters query proteins which share a common evolutionary ancestor whilst showing only the most significant match within the given PFAM clan.

##### 3.6 *Riftia* gene toolkits essential for development, homeostasis, and body patterning

###### 3.6.1 Antennapedia class

Using blastp v2.8.1+ the predicted *Riftia pachyptila* and *Lamellibrachia luymesi* protein sequences were searched against a well-curated catalog of metazoan Antennapedia class genes (Simakov et al. 2013; Zwarycz et al. 2016; Luo et al. 2018; Li et al. 2019). Additionally, Antennapedia gene ortholog representatives were downloaded from the NCBI protein database when necessary. The tubeworms protein candidates were aligned together with the metazoan homologues with mafft v7.450 (Katoh and Standley, 2013) and the multiple sequence alignments were manually trimmed to remove gappy and dissimilar regions. Independent molecular phylogenetic analyses were performed with iqtree v1.6.11 combining ModelFinder, tree search, 1000 ultra-fast bootstrap and SH-aLRT test replicates (Nguyen et al. 2015; Kalyaanamoorthy et al. 2017; Hoang et al. 2018).

###### 3.6.2 Transcription factors

Unique lophotrochozoan non-overlapping genes containing protein domains with similarity to the largest transcription factor families (bZIP, p53, homeobox, NuclearFactor, bHLH and zfc2h2) (Degnan et al. 2009; Vaquerizas et al. 2009; Schmitz et al. 2016) were quantified based on the results of the pfam_scan.pl analyses (section 3.5).

###### 3.6.3 Signalling, amino/fatty acid biosynthesis, endocytosis-, autophagy-, apoptosis- and immune-related gene toolkits

*Riftia pachyptila* genes containing protein domains associated with known signalling molecules (Wnt ligand and their Frizzled receptors, Hedgehog, Patched, Notch and TGF-beta), amino/fatty acid biosynthesis (mitochondrial beta oxidation), endocytosis

(RAB), autophagy (ATG, ULK), apoptosis-, and immune-related genes (Caspases, pro-caspases, Tumor Necrosis Factor ligands/receptor, Inhibitor of apoptosis, BCL, Toll-like, Superoxide dismutase) were retrieved from the pfam_scan.pl results (section 3.5). The orthology of the selected proteins were confirmed through similarity searches against the NCBI nr database (229,636,095 sequences) using diamond blastp v0.9.25.126 and phylogenetic inferences with iqtree v1.6.11, as described previously (section 3.6.1). The protein diagrams were drawn using IBS v.1.0.3 software (Liu et al. 2015) and the clustered heatmaps generated with the R package pheatmap (v1.0.12) (https://www.rdocumentation.org/packages/pheatmap/versions/1.0.12/topics/pheatmap).

##### 3.6.4 Hemoglobin gene identification, characterisation, phylogenetic inferences and gene expression quantification

Using blastp v2.8.1+ (Camacho et al. 2009) the predicted *Riftia pachyptila* protein sequences were searched against a well-curated publicly available metazoan hemoglobin and linker sequences. The tubeworm hemoglobin candidates were interrogated for the presence of the globin domain (PF00042) with hmmalign v3.1b2 and proteins without a hit were excluded from the analyses. Manual inspection and characterisation of the signature diagnostic residues/motifs in the hemoglobin chain and linker sequences were performed following previous works (Zal et al. 1996; Zal et al. 1997; Bailly et al. 2002; Bailly et al. 2003; Flores et al. 2005; Waits et al. 2016; Li et al. 2019). Multiple sequence alignments, trimming and phylogenetic analyses were carried out as described in the section 3.5. The resulting trees were midpoint rooted using Figtree (http://tree.bio.ed.ac.uk/software/figtree/). Additionally, to investigate the haemoglobin gene expression across different environmental conditions (sulphur rich and sulphur depleted) we downloaded six publicly available trophosome transcriptomes from SRA (https://www.ncbi.nlm.nih.gov/sra) (accession numbers: SRR8949066 to SRR8949071). The transcriptome libraries were pre-processed as described in section 2.2 (Transcriptome pre-processing). Quantification was performed with kallisto tool (Bray et al. 2016). The clustered heatmap was generated as previously described (section 3.6.3).

##### 3.6.5 Homology model generation of *Riftia* Hb

RIFPA_jg20259.t3-HB1c-SG1 mature sequence (Met63-Pro213), was modelled after the deoxygenated 400 kda hemoglobin structure from *Lamellibrachia satsuma* (pdb ID: 3WCT, resolution 2.4Å, sequence identity: 35.6%). Template was selected based on sequence and secondary structure similarity using HH-Pred online. Three-dimensional protein structures for the full complex were generated by homology modelling using the Prime program implemented in the Schrödinger Drug Discovery (v2020.2) software suite with standard options using secondary-structure prediction option to guide the query – template alignment. Protein complexes were prepared using the protein preparation tool and further refined with a restrained minimization step, allowing the heavy atoms to vary within 0.5 Å of limit, to fix sterical clashes. All models were inspected for Ramachandran outliers, which were fixed by energy minimization. All illustrations of structures were made with PyMol v2.4 (https://pymol.org/2/).

#### 4 COMPARATIVE GENOMICS AND GENE FAMILIES ANALYSES

##### 4.1 Orthology

36 representative animal species broadly distributed across the metazoan tree were selected for gene family analyses: *A. queenslandica*, *M. leidyi*, *N. vectensis*, *T. adhaerens* (non-bilaterian group); *A. vaga*, *B. platifrons*,  *B. glabrata*, *C. teleta*, *C. squamiferum*, *C. gigas*, *E. multilocularis*, *H. robusta*, *L. luymesi, L. anatina*, *L. gigantea, M. philippinarum*, *N. geniculatus*, *O. bimaculoides*, *M. yessoensis*, *P. australis*, *R. pachyptila*,  *S. mansoni*, *S. mediterranea* (lophotrochozoan group); *B. floridae*, *C. intestinalis*, *D. rerio*, *G. gallus*, *H. sapiens*,  *M. musculus*, *S. purpuratus*, *X. tropicalis* (deuterostomian group); *A. gambiae*, *C. elegans*, *D. pulex*, *D. melanogaster*, *T. castaneum* (ecdysozoan group). Redundancy was removed from the gene sets based on the gene coordinates specified in the gff3 files and only the longest isoform for each gene was kept. Orthofinder v2.3 (Emms and Kelly 2019) was used to define the orthogroups, and groups belonging only to *Capitella+Helobdella+Lamelibrachia+Riftia, Lamellibrachia+Riftia* and only *Riftia* were considered: Annelida-, Vestimentifera- and *Riftia*-specific, respectively. Enrichment analysis for Gene Ontology and protein annotation with PANTHER HMM scoring tool were performed with the *Riftia*-, *Lamellibrachia* and Vestimentifera-specific gene families as described in the section 4.2. To identify statistically significant gene family expansions/contractions in *Riftia* compared to other lophotrochozoans a second round of Orthofinder v2.3.8 was performed using 18 lophotrochozoan representatives (all except the bdelloid rotifer *Adineta vaga*, which was removed due the tetraploidy nature of its genome (Flot et al. 2013; Nowell et al. 2018), and *Tribolium castaneum* as outgroup. Finally, to identify the Annelida gene family core and the shared orthogroups only among annelids, a final instance of orthofinder was invoked using *C. teleta, H. robusta, L. luymesi* and *R. pachyptila.*

##### 4.2 Gene family expansions/contractions with CAFE

The 67 single-copy ortholog groups shared among the 18 lophotrochozoans + *T. castaneum* were individually aligned with mafft v7.450 (Katoh and Standley 2013). The resulting 67 multiple sequence alignment files were concatenated into a protein supermatrix using FASconCAT-G v1.04 (Kück and Meusemann 2010), and subsequently trimmed with BMGE v1.12 (Criscuolo and Gribaldo 2010). The final concatenated trimmed alignment with 23,975 positions was submitted to Bayesian phylogenetic inferences using Phylobayes v4.1b with CAT-GTR model defined and a starting lophotrochozoan tree generated by FastTree version 2.1.3-SSE3 (De Bie et al. 2006; Lartillot and Philippe 2006; Benton et al. 2009; Price et al. 2010; Han et al. 2013).Three calibrations points based on fossil data were used based on Benton et al., (2009): (1) *C. teleta* – *H. robusta* = 581 MYA – 305 MYA; (2) *C. teleta* – *L.*

*gigantea* = 581 MYA – 531 MYA; (3) *L. gigantea* – *B. glabrata* = 531 MYA – 470 MYA. The age of the root (i.e., Protostomia split) was set as 600 MYA. The ultrametric tree was obtained after 31,665 rounds discarding the initial 7,916 rounds as burn-in (25%). Gene family expansions and contractions were statistically analysed using the time tree generated by Phylobayes v4.1b and the lophotrochozoan orthology inference described in the section 4.1 with the CAFE v4.2.1 (De Bie et al. 2006). CAFE was executed with four different lambda values and the appropriate error model. The contracted/expanded gene families were annotated with Interproscan 5.39-77.0 (Jones et al. 2014) and the Enrichment analysis for Gene Ontology was performed as described in the section 5.2. Rapidly evolving gene families in *R. pachyptila* were annotated using PANTHER HMM scoring tool v2.2 (ftp://ftp.pantherdb.org/hmm_scoring/current_release/) with PANTHER_hmmscore database v15 (Thomas 2003; Han et al. 2013; Mi et al. 2017).

##### 4.3 Gene family expansion/contraction with PFAM

Annotation of the PFAM domains was performed with 18 selected lophotrochozoan taxa (section 4.1) using the pfam_scan.pl (section 3.5). Protein domains associated with transposable elements (e.g., Helicase, Helitron, DDE_Tnp) and identified as DUF (domain of unknown function) were removed from the analysis. As described in Albertin et al. (2019) and Sun et al. (2017), reoccurring identical domains present in the same protein sequence were counted only once. An iterative two-tailed Fisher's exact test was implemented in R (https://www.r-project.org/) to identify protein domain contractions/expansions. The obtained p-values were corrected using Benjamini and Hochberg method (Benjamini and Hochberg 1995) and only domains with a significant p-value of $< 0.01$ were further investigated. Four sets of comparisons were performed:

1. Set 1: *Riftia* vs. *Lamellibrachia*;
2. Set 2: *Riftia* vs. remaining non-vestimentiferan lophotrochozoans (background average domain counts of 16 taxa – all except *L. luymesi* and *R. pachyptila*);
3. Set 3: *Lamellibrachia* vs. remaining non-vestimentiferan lophotrochozoans (background average domain counts of 16 taxa – all except *L. luymesi* and *R. pachyptila*);
4. Set 4: Hydrothermal vents lophotrochozoans with symbiotic association (i.e., *B. platifrons, C. squamiferum*, *R. pachyptila* and *L. luymesi*) vs. remaining lophotrochozoan species (background average domain counts of 12 taxa).

##### 4.4 Clustering of *Riftia* expanded PFAM-containing genes with clans

A dataset composed of the *Riftia pachyptila* expanded PFAM-containing genes was clustered during approximately 21,000 rounds with the program clans, a Java application for visualising protein families based on all-against all comparisons (ftp://ftp.tuebingen.mpg.de/pub/protevo/CLANS/) (Frickey and Lupas 2004). The final 3D maps were collapsed to 2D after the clustering for easier visualisation.

##### 4.5 Synonymous and non-synonymous substitution rates analyses

Non-synonymous (Ka) and synonymous (Ks) substitution rates were calculated with the stand-alone version of KaKs_calculator v.2 using the orthofinder v2.3.8 results for the Lophotrochozoa and Annelida set (Wang et al. 2010; Emms and Kelly 2019). A modified version of pal2nal.pl (Suyama et al. 2006) software, called Epal2nal v.13 and available at the ParaAT v2.0 package (https://bigd.big.ac.cn/tools/paraat/) (Z. Zhang et al. 2012), was used to generate codon alignments in axt format required by KaKs_calculator. Additionally, potential genes under positive selection in *R. pachyptila* genome were screened in the Annelida and Lophotrochozoa orthogroups using the adaptive branch site-random effects likelihood (aBS-REL) model implemented in HyPhy v. 2.5.15 (Pond et al. 2005; Smith et al. 2015). Only single-copy genes (1:1 orthologs) without any inconsistencies between the nucleotide and protein sequences were used in the analyses.

#### 5 GENE EXPRESSION ANALYSES

##### 5.1 Gene expression quantification and identification of absolutely tissue specific genes

The eight pre-processed *Riftia pachyptila* transcriptome libraries were pseudoaligned against the merged filtered AUGUSTUS gene models with kallisto v.0.46.1 to collect the gene expression data expressed as TPM counts (transcripts per million) (Bray et al. 2016; Hoff and Stanke 2018). Normalisation within and across tissues were independently performed before calculating the tissue specificity tau values (see https://rdrr.io/github/roonysgalbi/tispec/f/vignettes/UserGuide.Rmd) (Yanai et al. 2005; Kryuchkova-Mostacci and Robinson-Rechavi 2016). To mitigate possible sex-specific differences in the gene expression levels, tau calculations were performed using only the tubeworm female tissues.

##### 5.2 Gene set enrichment analyses

The absolutely tissue specific genes (genes expressed only in a single tissue defined by a tau value of 1) were submitted to enrichment analyses for Gene Ontology with topGO v2.36.0 (https://bioconductor.org/packages/release/bioc/html/topGO.html) using Fisher's exact test against the *R. pachyptila* background (i.e., complete set of *Riftia* genes) coupled with weight01 algorithm (Alexa et al., 2006).

**LIST OF BIOINFORMATIC COMMANDS**
### 1 GENOME PIPELINE

#### 1.1 Pre-processing of databases

##### 1.1.1 Converting native BAM PacBio files to fastq

- bam2fastq -c 9 -o PacBio_library.fastq.gz PacBio_library.bam

##### 1.1.2 Removal of endosymbiont and tubeworm mitochondrial reads

- minimap2 -ax map-pb Endoriftia_plus_Riftia_mito.fasta PacBio_library.fastq.gz | samtools sort -o PacBio_library_mapped_against_contaminants.bam

##### 1.1.3 Extraction of unmapped reads from BAM results

- samtools view -f 4 PacBio_library_mapped_against_contaminants.bam > PacBio_library_mapped_against_contaminants_unmapped_reads.bam
- bam2fastq -c 9 -o PacBio_library_mapped_against_contaminants_unmapped_reads.fastq.gz PacBio_library_mapped_against_contaminants_unmapped_reads.bam

#### 1.2 Genome assemblies

##### 1.2.1 *Riftia* genome assembly: canu

- canu corMhapSensitivity=normal corOutCoverage=200 correctedErrorRate=0.105 “batOptions=-dg 3 -db 3 -dr 1 -ca 500 -cp 50” genomeSize=700m -pacbio-raw PacBio_libraries.fastq.gz

##### 1.2.2 *Riftia g*enome assembly: flye

- flye -g 772m --debug --pacbio-raw PacBio_libraries.fastq.gz -o Rpa-flye

#### 1.3 Genome post-processing

##### 1.3.1 Creating pbmm2 reference databases with genome assemblies

- pbmm2 index assembly_result.fasta assembly_result.mmi

##### 1.3.2 Alignment of PacBio native data against draft genomes with pbmm2

- pbmm2 align --log-level DEBUG --sort assembly_result.mmi list_of_PacBio_bam_files.fofn assembly_result_mapped_against_PacBio_native.bam

###### 1.3.3 Polishing and variant calling with arrow

- arrow assembly_result_mapped_against_PacBio_native.bam -r assembly_result.fasta -o assembly_result_arrow.fastq -o assembly_result_arrow.fasta -o assembly_result_arrow.gff3 --annotateGFF --algorithm arrow --log-level INFO --reportEffectiveCoverage

###### 1.3.4 Purging haplotigs and contig overlaps with purge_dups: minimap aligment and read, base-level read depth calculations

- for i in list_of_PacBio_databases.txt; do minimap2 -xmap-pb assembly_result_arrow.fasta $i | gzip -c - > $i.paf.gz; done
- pbcstat *.paf.gz
- calcuts -l 15 -u 130 -m 50  PB.stat > cutoffs 2>calcults.log

###### 1.3.5 Purging haplotigs and contig overlaps with purge_dups: self-alignment

- split_fa  assembly_result_arrow.fasta >  assembly_result_arrow.fasta.split
- minimap2 -xasm5 -DP assembly_result_arrow.fasta.split assembly_result_arrow.fasta.split | gzip -c - > assembly_result_arrow.fasta.split.self.paf.gz

###### 1.3.6 Purging haplotigs and contig overlaps with purge_dups: purge haplotigs and overlaps

- purge_dups -2 -T cutoffs -c PB.base.cov assembly_result_arrow.fasta.split.self.paf.gz > dups.bed 2> purge_dups.log

###### 1.3.7 Purging haplotigs and contig overlaps with purge_dups: get purged haplotig sequences

- get_seqs dups.bed  assembly_result_arrow.fasta > assembly_result_arrow_purged.fasta 2> assembly_result_arrow_haplotigs.fasta

###### 1.3.8 Contamination screening with blobtools: generating coverage file with minimap2

- minimap2 -ax map-pb  assembly_result_arrow_purged.fasta PacBio_libraries_no_contamination.fastq.gz | samtools sort -o assembly_result_arrow_mapped.bam

**1.3.9 Contamination screening with blobtools: generating hit file**

- blastn -max_target_seqs 10 -max_hsps 1 -query assembly_result_arrow_purged.fasta  -outfmt '6 qseqid staxids bitscore std' -db nt -evalue 1e-25 -out hit_file_tabular.txt

**1.3.10 Contamination screening with blobtools: generating blobplot**

- blobtools create -i assembly_result_arrow_purged.fasta -b assembly_result_arrow_mapped.bam -t hit_file_tabular.txt -o blobplot --taxrule "bestsumorder"

**1.3.11 Contamination screening: diamond searches against suspicious scaffold**

- diamond blastp -d nr.dmnd --outfmt 0 -q list_of_genes_tig00019702.fasta --evalue 1e-10 --verbose --out tig00019702_blastp_vs_diamond_1e10.txt
- diamond blastp -d nr.dmnd --outfmt 0 -q list_of_genes_tig00002255.fasta --evalue 1e-10 --verbose --out tig00002255_blastp_vs_diamond_1e10.txt

**1.3.12 Contamination screening: identifying bacterial gene markers with checkm**

- checkm lineage_wf -x fasta -g -t 10 ./ output/ >checkM.log

**2.1 Mitochondrial genome assembly and annotation**

**2.1.1 Flye assembly**

- flye --genome-size 15k --pacbio-raw reads_that_mapped_against_mit_ref.fastq.gz  -o RIFPA_mitochondrion

**2.1.2 Coverage plot with ConcatMap: minimap2**

- minimap2 -t 12 -ax map-pb -N 5 --secondary=no Riftia_pachyptila_mito.fasta m54098_190810_mapped_mito.fastq.gz > Riftia_pachyptila_mito_vs_reads.sam

**2.1.3 Coverage plot with ConcatMap: ConcatMap**

- ConcatMap_v1.2.py -m 500 –input_file Riftia_pachyptila_mito_vs_reads.sam --reference_length 15406 --figure_format "svg" -s 15 -l 0.002

###### 2.1.4 Circular plot with circos

- circos circos.conf

###### 2.1.5 Synteny analysis with Sibelia

- Sibelia -s far -o synteny_Riftia_mitochondria –gff all_mitochondria_refs.fasta

#### 2 TRANSCRIPTOME PIPELINE

##### 2.1 Quality assessment of the raw transcriptome databases

- fastqc transcriptome_1.fastq.gz transcriptome_2.fastq.gz

##### 2.2 Adapter removal and quality trimming

###### 2.2.1 Adapter removal

- bbduk ktrim=r ziplevel=9 ref=bbmap/resources/adapters.fa in=transcriptome_#.fastq.gz out=transcriptome_#_no_adapters.fastq.gz k=23 mink=11 hdist=1 tpe tbo

###### 2.2.2 Quality trimming

- bbduk ziplevel=9 qtrim=rl trimq=30 minlength=30 in=transcriptome_#_no_adapters.fastq.gz out=transcriptome_#_no_adapters_filtered_trimmed.fastq.gz

##### 2.3 Transcriptome assembly

###### 2.3.1 *De novo* transabyss assembly

- transabyss -SS -pe transcriptome_1_no_adapters_filtered_trimmed.fastq.gz transcriptome_2_no_adapters_filtered_trimmed.fastq.gz --length 200 --cleanup 0

###### 2.3.2 Reference-based assembly: Generating genome database with STAR

- STAR --runMode genomeGenerate --genomeDir ./ --genomeFastaFiles Riftia_genome_hard_masked.fasta --genomeSAindexNbases 13

###### 2.3.3 Reference-based assembly: RNA-seq alignment with STAR

- STAR --runMode alignReads --outFilterScoreMinOverLread 0.1 --outFilterMatchNminOverLread 0.1 --genomeDir 001-Database/ --readFilesIn transcriptome_1_no_adapters_filtered_trimmed.fastq transcriptome_2_no_adapters_filtered_trimmed.fastq --outFileNamePrefix transcriptome_STAR_mapped_outFilter0.1 --outSAMtype BAM Unsorted --outBAMcompression 10

###### 2.3.4 Reference-based assembly: Sorting BAM files with samtools

- samtools sort transcriptome_STAR_mapped_outFilter0.1-Aligned.out.bam

###### 2.3.5 Reference-based assembly: Indexing sorted BAM files with samtools

- samtools index transcriptome_STAR_mapped_outFilter0.1-Aligned.out.bam

###### 2.3.6 Reference-based assembly: Assembling with Stringtie

- stringtie transcriptome_STAR_mapped_outFilter0.1-Aligned.out.bam.sorted --conservative --rf -m 150 -o transcriptome.gtf -A transcriptome_gene_abundances.tab -l RPA

###### 2.3.7 Reference-based assembly: Merging Stringtie assemblies

- stringtie --merge -m 150 -l RPA_merged -o Riftia_pachyptila_STAR_transcriptomes.gtf list_of_stringtie_gtf_files_to_merge.txt

##### 2.4 *De novo* transcriptome post-processing

###### 2.4.1 Removal of endosymbiont contamination: creating BLAST database

- makeblastdb -dbtype nucl -hash_index -parse_seqids -in Riftia_masked_plus_Endoriftia_db.fasta

###### 2.4.2 Removal of endosymbiont contamination: running blastn

- blastn -evalue 1e-05 -num_descriptions 5 -num_alignments 5  -perc_identity 90 -db Riftia_masked_plus_Endoriftia_db.fasta -query transabyss_transcriptome.fasta

###### 2.4.3 Generating a *de-novo* global non-redundant transcriptome: creating Bowtie database

- bowtie2-build transcriptome_transabyss.fasta  transcriptome_transabyss.fasta

###### 2.4.4 Generating a *de-novo* global non-redundant transcriptome: multi-mapping of RNA-reads with Bowtie

- bowtie2 -k 40 --no-unal -1 transcriptome_1_no_adapters_filtered_trimmed.fastq.gz -2 transcriptome_2_no_adapters_filtered_trimmed.fastq.gz -x transcriptome_transabyss.fasta | samtools sort -o transcriptome_mapped.bam

###### 2.4.5 Generating a *de-novo* global non-redundant transcriptome: clustering transcripts into genes with Corset

- corset all_bam_files_generated_by_bowtie2.bam

###### 2.4.6 Generating a *de-novo* global non-redundant transcriptome: building the global *de novo* non-redundant "SuperTranscriptome" with Lace

- Lace.py –outputDir lace_results_from_transcriptome transcriptome_transabyss.fasta transcriptome_clusterFile_generated_by_corset.txt

##### 2.5 Prediction of the coding sequence regions with TransDecoder and Stringtie results

###### 2.5.1 Retrieving complete genes from Stringtie GTF files

- TransDecoder-v5.5.0/util/gtf_genome_to_cdna_fasta.pl Riftia_pachyptila_transcriptomes.gtf Riftia_genome_hard_masked.fasta  >Riftia_pachyptila_transcriptome_transdecoder.fasta

###### 2.5.2 Converting Stringtie GTF to GFF3 file

- TransDecoder-v5.5.0/util/gtf_to_alignment_gff3.pl Riftia_pachyptila_transcriptomes.gtf >Riftia_pachyptila_transcriptomes_transdecoder.gff3

###### 2.5.3 Extracting all candidate coding sequence regions from Stringtie global transcriptome

- TransDecoder.LongOrfs -t Riftia_pachyptila_transcriptomes_transdecoder.fasta -S
- TransDecoder.Predict -t Riftia_pachyptila_transcriptomes_transdecoder.fasta --retain_pfam_hits longest_orfs.pep-vs-pfam.domtblout.txt --retain_blastp_hits longest_orfs.pep-vs-uniprot-blastp.outfmt6.txt --output_dir RPA_transdecoder/

##### 2.5.4 Generating a genome-based Stringtie TransDecoder annotation file

- TransDecoder-v5.5.0/util/cdna_alignment_orf_to_genome_orf.pl Riftia_pachyptila_transcriptomes_transdecoder.fasta.transdecoder.gff3 Riftia_pachyptila_transcriptomes_transdecoder.gff3 Riftia_pachyptila_transcriptomes_transdecoder.fasta

##### 2.5.5 Homology searches with candidate CDS against PFAM-A and SwissProt

- blastp -query longest_orfs.pep -db uniprot_sprot.fasta -max_target_seqs 1 -outfmt 6 -evalue 1e-06 longest_orfs.pep-vs-uniprot-blastp.outfmt6.txt
- hmmscan -E 1e-06 --domtblout longest_orfs.pep-vs-pfam.domtblout Pfam-A.hmm longest_orfs.pep

#### 2.6 Prediction of the coding sequence regions with TransDecoder and Lace results

##### 2.6.1 Generating a GTF file for Lace SuperTranscriptome: create a paf file

- minimap2 -cx splice -C5 --secondary=no -uf -t 1 --cs Riftia_genome_hard_masked.fasta Riftia_pachyptila_SuperTranscriptomes.fasta > Riftia_pachyptila_SuperTranscriptomes_vs_Riftia_genome.paf

##### 2.6.2 Generating a GTF file for Lace SuperTranscriptome: convert paf to bed file

- paftools.js splice2bed Riftia_pachyptila_SuperTranscriptomes_vs_Riftia_genome.paf > Riftia_pachyptila_SuperTranscriptomes_vs_Riftia_genome.bed

##### 2.6.3 Generating a GTF file for Lace SuperTranscriptome: convert bed to GTF file with bedToGenePred and genePredToGtf tools

- bedToGenePred Riftia_pachyptila_SuperTranscriptomes_vs_Riftia_genome.bed /dev/stdout |

genePredToGtf file /dev/stdin
Riftia_pachyptila_SuperTranscriptomes_vs_Riftia_genome.gtf

##### 2.6.4 Retrieving complete genes from Lace GTF files

- TransDecoder-v5.5.0/util/gtf_genome_to_cdna_fasta.pl Riftia_pachyptila_SuperTranscriptomes_vs_Riftia_genome.gtf Riftia_genome_hard_masked.fasta >Riftia_pachyptila_SuperTranscriptomes_transdecoder.fasta

##### 2.6.5 Converting Lace GTF to GFF3 file

- TransDecoder-v5.5.0/util/gtf_to_alignment_gff3.pl Riftia_pachyptila_SuperTranscriptomes_vs_Riftia_genome.gtf > Riftia_pachyptila_SuperTranscriptomes_transdecoder.gff3

##### 2.6.6 Extracting all candidate coding sequence regions from Lace SuperTranscriptome

- TransDecoder.LongOrfs -t Riftia_pachyptila_SuperTranscriptomes_transdecoder.fasta -S
- TransDecoder.Predict -t Riftia_pachyptila_SuperTranscriptomes_transdecoder.fasta --retain_pfam_hits longest_orfs.pep-vs-pfam.domtblout.txt --retain_blastp_hits longest_orfs.pep-vs-uniprot-blastp.outfmt6.txt --output_dir RPA_Super_transdecoder/

##### 2.6.7 Generating a genome-based Lace TransDecoder annotation file

- TransDecoder-v5.5.0/util/cdna_alignment_orf_to_genome_orf.pl Riftia_pachyptila_SuperTranscriptomes_transdecoder.fasta.transdecoder.gff3 Riftia_pachyptila_SuperTranscriptomes_transdecoder.gff3 Riftia_pachyptila_SuperTranscriptomes_transdecoder.fasta

##### 2.6.8 Homology searches with candidate CDS against PFAM-A and SwissProt

1. blastp -query longest_orfs.pep -db uniprot_sprot.fasta -max_target_seqs 1 -outfmt 6 -evalue 1e-06 longest_orfs.pep-vs-uniprot-blastp.outfmt6.txt
2. hmmscan -E 1e-06 --domtblout longest_orfs.pep-vs-pfam.domtblout Pfam-A.hmm longest_orfs.pep

#### 2.7 Quality assessment of the final predicted proteomes with BUSCO

- busco -i Riftia_pachyptila_SuperTranscriptomes_transdecoder.fasta.transdecoder.pep -l metazoa_odb10 –offline -m protein
- busco -i Riftia_pachyptila_transcriptomes_transdecoder.fasta.transdecoder.pep -l metazoa_odb10 --offline -m protein

**3 ANNOTATION PIPELINE**

**3.1 Identification of the repetitive regions with RepeatModeler/RepeatMasker**

**3.1.1 Generating custom repeat library with RepeatModeler**

- RepeatModeler -genomeSampleSizeMax 560783187 -LTRStruct -database Riftia_pachyptila_genome.fasta

**3.1.2 Hard masking the *Riftia* genome with RepeatMasker**

- RepeatMasker -s -e rmblast -dir . -gff -html -a -lib Riftia_pachtyptila_repetitive_db.fasta Riftia_pachyptila_genome.fasta

**3.1.3 Soft masking the *Riftia* genome with RepeatMasker**

- RepeatMasker -s -e rmblast -dir . -gff -html -a -lib Riftia_pachtyptila_repetitive_db.fasta -xsmall Riftia_pachyptila_genome.fasta

**3.1.4 Generation of *Riftia* repeat landscape**

- calcDivergenceFromAlign.pl -s Riftia_div.txt Riftia_pachyptila_genome_masked.fasta.align
- createRepeatLandscape.pl -div Riftia_div.txt -g 560783187

**3.2 *Ab initio* gene prediction with augustus**

**3.2.1 Generating *Riftia pachyptila* intron hints file: merging STAR mapped transcriptomes**

- samtools merge -b all_mapped_star_transcriptomes.fofn -O BAM -l 9 Rpa_merged_transcriptomes_STAR.bam

**3.2.2 Generating *Riftia pachyptila* intron hints file: sorting BAM files by name**

- samtools sort -n Rpa_merged_transcriptomes_STAR.bam -O BAM -o Rpa_merged_transcriptomes_STAR.sortedByName.bam
- samtools sort Rpa_merged_transcriptomes_STAR.sortedByName.filtered.bam -o Rpa_merged_transcriptomes_STAR.sortedByName.filtered.sorted.bam

##### 3.2.3 Generating *Riftia pachyptila* intron hints file: filtering aligned reads from BAM

- filterBam --uniq --paired --pairwiseAlignments --in Rpa_merged_transcriptomes_STAR.sortedByName.bam --out Rpa_merged_transcriptomes_STAR.sortedByName.filtered.bam

##### 3.2.4 Generating *Riftia pachyptila* intron hints file: extracting intron information and removing inappropriate splice site information from BAM file

- bam2hints --intronsonly --in=Rpa_merged_transcriptomes_STAR.sortedByName.filtered.bam --out=Riftia_introns_hints.gff
- filterIntronsFindStrand.pl  Riftia_genome_soft_masked.fasta Riftia_introns_hints.gff --score > Riftia_introns_filtered_hints.gff

##### 3.2.5 Generating training set gene structure: running unsupervised GeneMark-ET

- gmes_petap.pl --verbose –sequence=Riftia_genome_soft_masked.fasta --ET=Riftia_introns_filtered_hints.gff --soft_mask 1000 --max_intron 349151 --max_contig 13119000 --min_contig 2273

##### 3.2.6 Generating training set gene structure: filtering GeneMark-ET predictions

- filterGenemark.pl genemark.gtf  Riftia_introns_filtered_hints.gff

##### 3.2.7 Generating training set gene structure: computing flanking region size of the genes

- computeFlankingRegion.pl genemark.f.good.gtf

##### 3.2.8 Generating training set gene structure: converting GeneMark structures to GenBank

- gff2gbSmallDNA.pl genemark.f.good.gtf Riftia_genome_hard_masked.fasta 124 tmp.gb

**3.2.9 Generating training set gene structure: filtering good gene models from bonafide.gtf**

- filterGenesIn_mRNAname.pl genemark.gtf tmp.gb > GeneMark_models_filtered.gb

**3.2.10 Generating *Riftia pachyptila* exonpart hints from RNA-seq data**

- bam2wig Rpa_merged_transcriptomes_STAR.sortedByName.filtered.bam > RNASeq_exonpart_Rpa_merged_transcriptomes_STAR.wig
- cat RNASeq_exonpart_Rpa_merged_transcriptomes_STAR.wig | wig2hints.pl --width=10 --margin=10 --minthresh=2 --minscore=4 --prune=0.1 --src=W --type=ep -UCSC=unstranded.track --radius=4.5 --pri=4 --strand="." > Riftia_exonpart_hints.gff

**3.2.11 Generating *Riftia pachyptila* exonpart, intron, acceptor (3') splice site, donor splice site (5') and exon hints file from Cdna data: running blat**

- blat -noHead -mask=lower soft -minIdentity=92 Riftia_genome_soft_masked.fasta Riftia_pachyptila_non_redundant_de_novo_reference_based_cDNA_from_transdecoder.fasta cdna_vs_Riftia_genome_soft_masked.psl

**3.2.12 Generating *Riftia pachyptila* exonpart, intron, acceptor (3') splice site, donor splice site (5') and exon hints file from Cdna data: filtering and sorting blat alignments**

- pslCDnaFilter -ignoreNs -localNearBest=0.005 -minId=0.9 -bestOverlap cdna_vs_Riftia_genome_soft_masked.psl > cdna_vs_Riftia_genome_soft_masked.filtered.psl
- cat cdna_vs_Riftia_genomes_soft_masked.filtered.psl |sort -n -k 16,16 |sort -s -k 14,14 > cdna_vs_Riftia_genomes_soft_masked.filtered.sorted.psl

**3.2.13 Generating *Riftia pachyptila* exonpart, intron, acceptor (3') splice site, donor splice site (5') and exon hints file from Cdna data: converting psl file to hints format**

- blat2hints.pl --in=cdna_vs_Riftia_genomes_soft_masked.filtered.sorted.psl --minintronlen=31 --maxintronlen=349151 --trunkSS --ssOn --remove_redundant --out=Riftia_cDNA_hints.gff

##### 3.2.14 Generating *Riftia pachyptila* protein hints file from protein data: running genomeThreader

- startAlign.pl --prg=gth --prot=Riftia_pachyptila_denovo_plus_reference_based_transcriptomes_clustered90.pep --genome= Riftia_genome_hard_masked.fasta

##### 3.2.15 Generating *Riftia pachyptila* protein hints file from protein data: converting alignment output to protein hints file:

- align2hints.pl --minintronlen=31 --maxintronlen=349151 --in=gth.concat.aln --out=Riftia_protein_hints.gff --prg=gth

##### 3.2.16 Training augustus for *Riftia pachyptila*: creating a template for a new species

- new_species.pl --species=riftia_pachyptila
- randomSplit.pl Riftia_pachyptila_gene_models_filtered.gb 200
- etraining –species=riftia_pachyptila Riftia_pachyptila_gene_models_filtered.gb.train > etrain.out

##### 3.2.17 Training augustus for *Riftia pachyptila*: meta parameter optimisation

- optimize_augustus.pl –species=riftia_pachyptila Riftia_pachyptila_gene_models_filtered.gb.train
- optimize_augustus.pl –species=riftia_pachyptila Riftia_pachyptila_UTR_gene_models_filtered.gb.train --UTR=on --metapars=Augustus/config/species/riftia_pachyptila/riftia_pachyptila_metapars.utr.cfg --trainOnlyUtr=1

##### 3.2.18 Running *Riftia pachyptila* gene prediction with augustus

- augustus --species=riftia_pachyptila --UTR=no --softmasking=on Riftia_soft_masked_genome.fasta –min_intron_len=31 > Riftia_pachyptila_no_hints_predictions.gtf
- augustus --species=riftia_pachyptila Riftia_soft_masked_genome.fasta --extrinsicCfgFile=extrinsic.M.RM.E.W.P.cfg --

hintsfile=Riftia_pachtypila_hints_no_EP.gff --allow_hinted_splicesites=atac --alternatives-from-evidence=on --UTR=no --softmasking=on --min_intron_len=31 > Riftia_pachyptila_hints_no_UTR_predictions.gtf

- augustus –species=riftia_pachyptila Riftia_soft_masked_genome.fasta --extrinsicCfgFile=extrinsic.M.RM.E.W.P.cfg –hintsfile=Riftia_pachtypila_hints.gff --allow_hinted_splicesites=atac --alternatives-from-evidence=on --UTR=yes --softmasking=on --min_intron_len=31 > Riftia_pachyptila_hints_UTR_predictions.gtf

##### 3.3 Augustus post-processing results

###### 3.3.1 Joining gtf annotations

- joingenes --alternatives --genesets=Riftia_pachyptila_no_hints_predictions.gtf,Riftia_pachyptila_hints_no_UTR_predictions.gtf,Riftia_pachyptila_hints_UTR_predictions.gtf –priorities=1,2,3 --output=Riftia_pachyptila_predictions_combined.gtf

###### 3.3.2 Extracting coding sequence regions in amino acid and nucleotide formats

- getAnnoFastaFromJoingenes.py -g Riftia_pachyptila_soft_masked.genome --filter_out_invalid_stops=yes -f Riftia_pachyptila_predictions_combined.gtf -o Riftia_pachyptila_predictions_combined

###### 3.3.3 Removing non supported gene models: PFAM searches

- hmmsearch --cpu 12 --domtblout Riftia_pachyptila_predictions_combined_vs_PFAM-1e10_tabular.txt -E 1e-10 Pfam-A.hmm Riftia_pachyptila_predictions_combined.aa

###### 3.3.4 Removing non supported gene models: diamond blastp searches

- diamond blastp --db nr.dmnd --outfmt 6 --query Riftia_pachyptila_predictions_combined.aa --out diamond_augustus_predicted_vs_nr_1e10.txt --evalue 1e-10 --max-target-seqs 1

3.3.5 Removing non supported gene models: orthofinder orthology inferences

- orthofinder -M msa -t 12 -T iqtree -f directory_with_protein_files/

###### 3.3.6 Removing non supported gene models: gene expression evidence

- kallisto index -i Riftia_pachyptila_predictions_combined.codingseq.idx Riftia_pachyptila_predictions_combined.codingseq
- kallisto quant –bias -i Riftia_pachyptila_predictions_combined.codingseq.idx --rf-stranded -b 100 -o ./ Trimmed_filtered_transcriptomes_1.fastq.gz Trimmed_filtered_transcriptomes_2.fastq.gz

##### 3.4 Protein annotation

###### 3.4.1 Interproscan

- interproscan --iprlookup --goterms -i Riftia_pachyptila_predictions_combined_filtered.aa -f TSV,HTML,GFF3,XML --disable-precalc --pathways -d interpro_results

###### 3.4.2 signalp

- signalp-v5 -batch 20000 -fasta Riftia_pachyptila_predictions_combined_filtered.aa -format long -gff3 -mature

###### 3.4.3 tRNA-scan

- tRNAscan-SE -HQ -o# -f# -m# -s# -a# --detail -p Riftia_pachyptila_hard_masked Riftia_pachyptila_hard_masked.fasta

###### 3.4.4 Individual gene identification and phylogeny: Antennapedia cluster-, Wnt ligands/receptors-, apoptosis-, hemoglobin-related, immune-related genes

- blastp -db tubeworms_proteins.fasta -query metazoan_queries.fasta -outfmt 6 -max_target_seqs 10 > tubeworms_candidates_10best_tabular.txt
- for i in `cat tubeworms_candidates_10best_tabular.txt|sort -u`; do blastdbcmd -entry $i -db tubeworms_proteins.fasta >> tubeworms_candidates_10best_tabular.fasta; done;
- cat tubeworms_candidates_10best_tabular.fasta metazoan_homologues.fasta > tubeworms_candidates_plus_metazoa.fasta
- mafft --maxiterate 1000 --localpair tubeworms_candidates_plus_metazoa.fasta > tubeworms_candidates_plus_metazoa.fasta.aln
- iqtree -nt AUTO -s tubeworm_best_candidates_plus_metazoa_trimmed.fasta.aln -alrt 1000 -bb 1000

###### 3.4.5 Transcription factor families (TFFs) identification

- pfam_scan.pl -e_seq 1e-05 -e_dom 1e-05 -fasta proteome.fasta -dir PFAM-A/ -outfile output.txt

#### 4 GENE FAMILY ANALYSIS

##### 4.1 Adding unique identifier and removing redundancy with gff3 files coordinates

- sed -i 's/>/>ID_/' metazoan_db.fasta
- pymakeMap.py -gff metazoan.gff3 -p ID -f CDS -d -k protein_id -o metazoan.chrom

##### 4.2 Orthofinder

- orthofinder -f metazoan_db_directory/
- orthofinder -f lophotrochozoan_db_directory/
- orthofinder -f annelida_db_directory/

##### 4.3 Synonymous and non-synonymous substitution rates and positive selection analysis

###### 4.3.1 KaKs_calculator

- for i in `cat list_of_OGs.txt`; do mafft --maxiterate 1000 --localpair $i_AA.fasta.aln > $i_AA.fasta.aln; done;
- for i in `cat list_of_OGs.txt`; do Epal2nal.pl $i_AA.fasta.aln $i_CDS.fasta -output axt > $i_CDS.fasta.axt; done;
- for i in `cat list_of_OGs.txt`; do KaKs_calculator -i $i_CDS.fasta.axt -o $i_CDS.fasta.axt.kaks;

###### 4.3.2 Hyphy aBSREL

- for i in `cat list_of_OGs.txt`; do pal2nal.pl $i_AA.fasta.aln $i_CDS.fasta -output fasta > $i_CDS.fasta.aln; done;
- for i in `cat list_of_OG.txt`; do iqtree -st CODON -s $i_CDS_fixed.fasta.aln -m TESTONLY -nt AUTO; rm $i_CDS_fixed.fasta.aln.iqtree; rm $i_CDS_fixed.fasta.aln.mldist; rm $i_CDS_fixed.fasta.aln.log; rm $i_CDS_fixed.fasta.aln.ckp.gz; rm $i_CDS_fixed.fasta.aln.model.gz; rm $i_CDS_fixed.fasta.aln.bionj; done;

- for i in `cat list_of_OG.txt`; do perl -p -e "s/(RIFPA_Rpa_jg\d+.t\d+)/\$1\{tubeworm\}/" $i_CDS_fixed.fasta.aln.treefile.renamed > $i_CDS_fixed.fasta.aln.treefile.renamed.branch.label; done;
- for i in `cat list_of_OGs.txt`; do hyphy aBSREL --alignment $i_CDS_fixed.fasta.aln --tree \ $i_CDS_fixed.fasta.aln.treefile.renamed --branches tubeworm
- for i in `cat list_of_OGs_tubeworm_positive_selected.txt`; do hyphy aBSREL --alignment $i_CDS_fixed.fasta.aln --tree \$i_CDS_fixed.fasta.aln.treefile --branches All

###### 4.3.4 Annotation of positive selected genes: pantherScore2.0.pl

- pantherScore2.2.pl -l PANTHER14.1/ -D B -V -i positive_selected_genes_lophotrochozoa.fasta -o output.panther -n -s
- pantherScore2.2.pl -l PANTHER14.1/ -D B -V -i positive_selected_genes_annelida.fasta -o output.panther -n -s

##### 4.4 Phylogenomics

###### 4.4.1 Unifying header names across orthogroups

- for i in `ls -1 *fa`; do perl -pe 's/(>\S{5}).*/$1/' $i > $i.renamed.fasta; done;

###### 4.4.2 Multiple sequencing alignments

- for i in `cat list.txt`; do mafft --maxiterate 1000 --localpair $i.renamed.fasta > $i.renamed.fasta.aln; done;

###### 4.4.3 FASconCAT and multiple sequencing alignment trimming

- FASconCAT-G_v1.04.pl
- BMGE.jar -i Lophotrochozoa_supermatrix.fasta -m BLOSUM30 -h 1 -b 1 -t AA -of Lophotrochozoa_supermatrix.trimmed.fasta

###### 4.4.4 Bayesian inference: generating Lophotrochozoa starting tree and phylogenetic run

- FastTree Lophotrochozoa_supermatrix.trimmed.fasta > Lophotrochozoa.tree

- pb -d Lophotrochozoa_supermatrix.trimmed.phy -T Lophotrochozoa.tree -r outgroup.txt -ln -cal calibrations.txt -gtr -cat -rp 600 0 -prior lophotrochozoa_run
- readdiv -x 7916 100 lophotrochozoa_run

##### 4.5 Expanded/contracted gene families

##### 4.5.1. CAFE analysis: preparing orthogroups for CAFE

- python cafetutorial_clade_and_size_filter.py -i Orthogroups.GeneCount.tsv -o Orthogroups.GeneCount.filtered.tsv -s
- ./cafe_filtered_families.sh
- ./cafe_large_families.sh
- ./cafe_filtered_families_lambda.sh
- ./caferror.sh
- ./cafe_filtered_families_error_many_lambdas.sh
- cafetutorial_report_analysis.py -r 0 -i report_file.cafe -l large_families.cafe -o output_report

##### 4.5.2 Annotation of the rapidly evolving gene families: pantherScore2.0.pl

- pantherScore2.2.pl -l PANTHER14.1/ -D B -V -i rapidly_evolving_gene_families.fasta -o output.panther -n -s

##### 4.6 Clustering of *Riftia pachyptila* expanded PFAM-domain containing genes

- clans -in PFAM_domain_containing_genes.fasta -eval 1e-5 -cpu 10 -blastpath "blastp"

##### 4.6 Global view of *Riftia pachyptila* proneuropeptide and prohormone complement

- signalp -fasta Riftia_pachyptila_proteins.fasta -gff3 -mature
- hmmsearch -E 1e-10 –domtblout Riftia_pachyptila_mature_with_cleavage_vs_PFAM-A.txt Pfam-A.hmm Riftia_pachyptila_proteins_mature_with_cleavage_sites.fasta
- clans -in Riftia_general_cluster.fasta -eval 1e-10 -blastpath psiblast
- clans -in Riftia_central_cluster.fasta -eval 1e-10 -blastpath blastp

### 5 GENE EXPRESSION QUANTIFICATION

#### 5.1 Kallisto: creating index

- kallisto index -i Riftia_pachyptila_predictions_combined.codingseq.idx Riftia_pachyptila_predictions_combined.codingseq

#### 5.2 Kallisto: quantification

- for i in `cat list_of_trimmed_filtered_transcriptomes.txt`; do kallisto quant --bias -i Riftia_pachyptila_predictions_combined.codingseq.idx --rf-stranded –plaintext -b 100 -o ./ $i_trimmed_filtered_1.fastq.gz $i_trimmed_filtered_2.fastq.gz

#### 5.3 Tissue specificity quantification: tau values

- Rscript tau_female_tissues.R

#### 5.4 GO enrichment analyses

- Rscript goTerms_female_tissues.R

### 6 ADDITIONAL COMMAND LINE APPLICATIONS USED

#### 6.1 Quality assessment with quast

- quast.py -m 1 file1.fasta file2.fasta

#### 6.2 Generating Cdna, cds and protein datasets for augustus

##### 6.2.1 Combining transdecoder predicted protein files from *de novo* and reference-based approaches

- cat predicted_proteins_transdecoder_from_de_novo_assemblies.pep predicted_proteins_transdecoder_from_reference_based_assemblies.pep > combined_proteins_from_de_novo_reference_based_assemblies_transdecoder.pep

##### 6.2.2 Removing redundancy from combined protein dataset

- cd-hit -i combined_proteins_from_de_novo_reference_based_assemblies_transdecoder.pep -o

Riftia_pachyptila_denovo_plus_reference_based_transcriptomes_clustered90.pep -c 0.9

##### 6.2.3 Retrieving matching cDNA fasta files: creating blast database cDNA files

- makeblastdb -dbtype nucl -hash_index -parse_seqids -in Riftia_pachyptila_denovo_plus_reference_based_transcriptomes_cDNA.fasta

##### 6.2.3 Retrieving matching cDNA fasta files: retrieving sequences with blastdbcmd

- blastdbcmd -entry_batch list_of_fasta_headers_to_retrieve.txt -db Riftia_pachyptila_denovo_plus_reference_based_transcriptomes_cDNA.fasta > Riftia_pachyptila_non_redundant_de_novo_reference_based_cDNA_from_transdecoder.fasta

#### 6.3 Removing redundant gene structures from augustus training sets

##### 6.3.1 Converting training gene structure gtf file to a protein sequences

- gtf2aa.pl Riftia_genome_hard_masked.fasta training_genes.gtf proteins.fasta

##### 6.3.2 Removing redundancy from training gene amino acid sequences

- aa2nonred.pl proteins.fasta proteins.nr.fasta

##### 6.3.3 Filtering and retrieving non-redundant loci

- diamond blastp nr.dmnd --outfmt 6 --query  proteins.nr.fasta --out nr-proteins-vs-diamond-1e10.full.txt --evalue 1e-10
- cut -f1 nr-proteins-vs-diamond-1e10.full.txt |sort -u > nonred.lst
- cat  training_genes.gb | perl -ne 'if ($_ =~ m/LOCUS\s+(\S+)\s/) {$txLocus = $1;} elsif ($_ = m/\/gene=\"(\S+)\"/) { $txInGb3{$1} = $txLocus } if(eof()) { foreach (keys %txInGb3) { print "$_\t$txInGb3{$_}\n"; } }' > loci.lst
- grep -f nonred.lst loci.lst | cut -f2 > nonred.loci.lst
- filterGenesIn.pl nonred.loci.lst training_genes.gb > training_genes_filtered.gb
